# Brain expression quantitative trait locus and network analysis reveals downstream effects and putative drivers for brain-related diseases

**DOI:** 10.1101/2021.03.01.433439

**Authors:** Niek de Klein, Ellen A. Tsai, Martijn Vochteloo, Denis Baird, Yunfeng Huang, Chia-Yen Chen, Sipko van Dam, Patrick Deelen, Olivier B. Bakker, Omar El Garwany, Zhengyu Ouyang, Eric E. Marshall, Maria I. Zavodszky, Wouter van Rheenen, Mark K. Bakker, Jan Veldink, Tom R. Gaunt, Heiko Runz, Lude Franke, Harm-Jan Westra

## Abstract

Gaining insight into the downstream consequences of non-coding variants is an essential step towards the identification of therapeutic targets from genome-wide association study (GWAS) findings. Here we have harmonized and integrated 8,727 RNA-seq samples with accompanying genotype data from multiple brain-regions from 14 datasets. This sample size enabled us to perform both *cis*- and *trans*-expression quantitative locus (eQTL) mapping. Upon comparing the brain cortex *cis*-eQTLs (for 12,307 unique genes at FDR<0.05) with a large blood *cis-*eQTL analysis (n=31,684 samples), we observed that brain eQTLs are more tissue specific than previously assumed.

We inferred the brain cell type for 1,515 *cis*-eQTLs by using cell type proportion information. We conducted Mendelian Randomization on 31 brain-related traits using *cis*-eQTLs as instruments and found 159 significant findings that also passed colocalization. Furthermore, two multiple sclerosis (MS) findings had cell type specific signals, a neuron-specific *cis-*eQTL for *CYP24A1* and a macrophage specific *cis*-eQTL for *CLECL1*.

To further interpret GWAS hits, we performed *trans*-eQTL analysis. We identified 2,589 *trans*-eQTLs (at FDR<0.05) for 373 unique SNPs, affecting 1,263 unique genes, and 21 replicated significantly using single-nucleus RNA-seq data from excitatory neurons.

We also generated a brain-specific gene-coregulation network that we used to predict which genes have brain-specific functions, and to perform a novel network analysis of Alzheimer’s disease (AD), amyotrophic lateral sclerosis (ALS), multiple sclerosis (MS) and Parkinson’s disease (PD) GWAS data. This resulted in the identification of distinct sets of genes that show significantly enriched co-regulation with genes inside the associated GWAS loci, and which might reflect drivers of these diseases.

## Introduction

Diseases of the brain manifesting as psychiatric or neurological conditions continue to be a massive global health burden: The World Health Organization estimates that in 2019 globally 280 million individuals were affected by depression, 39.5 million by bipolar disorder, and 287.4 million by schizophrenia^1^. Likewise, the fraction of 50 million people living with dementia today is expected to rise to 152 million by 2050^2^, with similar trajectories for other neurodegenerative diseases. While substantial progress has been made in uncovering the genetic basis of psychiatric and neurological diseases through genome-wide association studies (GWAS), much of how the identified genetic variants impact brain function is still unknown.

To translate from genetic signals to mechanisms, associations with gene expression levels, or expression quantitative trait loci (eQTL) have shown great potential. eQTLs can be divided in direct effects of local genetic variants (*cis*-eQTLs) and indirect effects of distal variants (*trans*-eQTLs). *Cis*-eQTLs and *trans*-eQTLs can aid interpretation of GWAS loci in several ways. *Cis*-eQTLs aid interpretation by identifying direct links between genes and phenotypes through causal inference approaches such as Mendelian randomization (MR) instrumented on QTLs and genetic colocalization analysis, whereas *trans*-eQTLs expose sets of downstream genes and pathways on which the effects of disease variants converge.

eQTLs are dynamic features and vary with tissue, cell type and additional factors such as response to stimulation. For an optimal interrogation of GWAS loci, it is therefore desirable to perform eQTL analyses in disease-relevant tissues^3^. To help interpret GWAS of neurodegenerative and psychiatric diseases, several brain-derived eQTL studies have been published, including meta-analyses by the PsychENCODE^4^ and AMP-AD^5^ consortia, which cover 1,866 and 1,433 individuals, respectively. However, to yield reliable results, statistical approaches such as MR and colocalization require robust effect size estimates from even larger carefully curated eQTL datasets. Large sample sizes are better suited to decompose eQTL effects to specific cell types.

To maximize the potential of eQTL-based analyses in brain, we here combined and rigorously harmonized brain RNA-seq and genotype data from 15 different cohorts, including 8,727 RNA-seq samples from all major brain eQTL studies and publicly available samples from the European Nucleotide Archive (ENA). By leveraging the statistical power across these datasets, we created a gene coregulation network based on 8,544 RNA-seq samples covering different brain regions and performed *cis*- and *trans*-eQTL analysis in up to 2,970 individuals of European descent, with replication in up to 420 individuals of African descent. This sample size enabled us to make inferences on the brain cell types in which eQTLs operate, and to systematically conduct Mendelian Randomization and colocalization analyses to find shared genetic effects between eQTLs and GWAS traits. This prioritized likely causal genes from GWAS loci for 31 brain-related traits, including neurodegenerative and psychiatric conditions. Additionally, this identified cell type dependent eQTLs that may be associated with disease risk (Figure 1).

**Figure 1.**
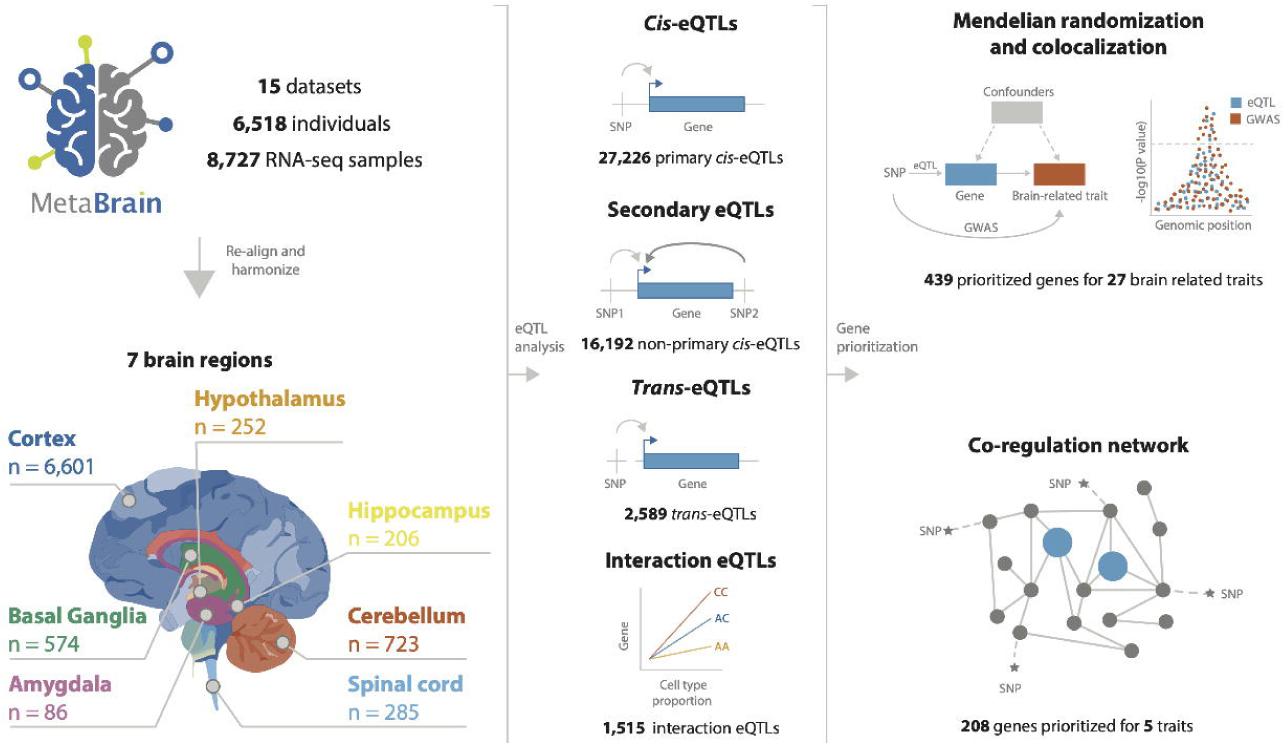
Overview of the study. We downloaded publicly available RNA-seq and genotype data from 15 different datasets consisting 8,727 RNA-seq measurements from 7 main brain regions in 6,518 individuals. We performed *cis-, trans-* and interaction*-*eQTL analysis, built a brain-specific gene coregulation network and prioritized genes using Mendelian randomization, colocalization and the co-regulation network.

## Results

### Leveraging public RNA-seq and genotype data to create large, harmonized brain eQTL and gene co-regulation datasets

We combined 15 eQTL datasets into the ‘*MetaBrain*’ resource to maximize statistical power to detect eQTLs and to create a brain specific gene coregulation network (Figure 2**; Supplementary Table 1, Supplementary Figures 1-5**). *MetaBrain* includes 7,604 RNA-seq samples and accompanying genotypes from the AMP-AD consortium^6^ (AMP-AD MAYO^6^, ROSMAP^6^ and MSBB^6^), Braineac^7^, the PsychENCODE consortium^8^ (Bipseq^4^, BrainGVEX^4^, CMC^9^, GVEX and UCLA_ASD^4^), BrainSeq^10^, NABEC^11^, TargetALS^12^, and GTEx^3^. Additionally, we carefully selected 1,759 brain RNA-seq samples from the European Nucleotide Archive (ENA)^13^, calling and imputing genotypes based on the RNA-seq alignment (**Supplementary Note, Supplementary Figure 1)**. There were 8,727 RNA-seq samples remaining after realignment and stringent quality control (**Methods and Supplementary Note, Supplementary Figure 2-3**). Using slightly different quality control measures, we created a gene network using 8,544 samples (**Supplementary Note**). We corrected the RNA-seq data for technical covariates and defined 7 major tissue groups (amygdala, basal ganglia, cerebellum, cortex, hippocampus, hypothalamus and spinal cord): Principal Component Analysis (PCA) on the RNA-seq data showed clear clustering by these major tissue groups, resembling brain physiology (Figure 2D**, Supplementary Figure 4**). Genotype data revealed individuals from different ethnicities (Figure 2B**; Supplementary Figure 5**), including 5,138 samples from European descent (EUR) and 805 samples from African descent (AFR). We created 6 *cis-*eQTL discovery datasets: Basal ganglia-EUR (n=208), Cerebellum-EUR (n=492), Cortex-EUR (n=2,970), Cortex-AFR (n=420), Hippocampus-EUR (n=168) and Spinal cord-EUR (n=108; **Supplementary Table 1,** Figure 2C). *Cis-*eQTLs were not calculated for amygdala and hypothalamus tissue groups due to the small sample size (n<100).

**Figure 2.**
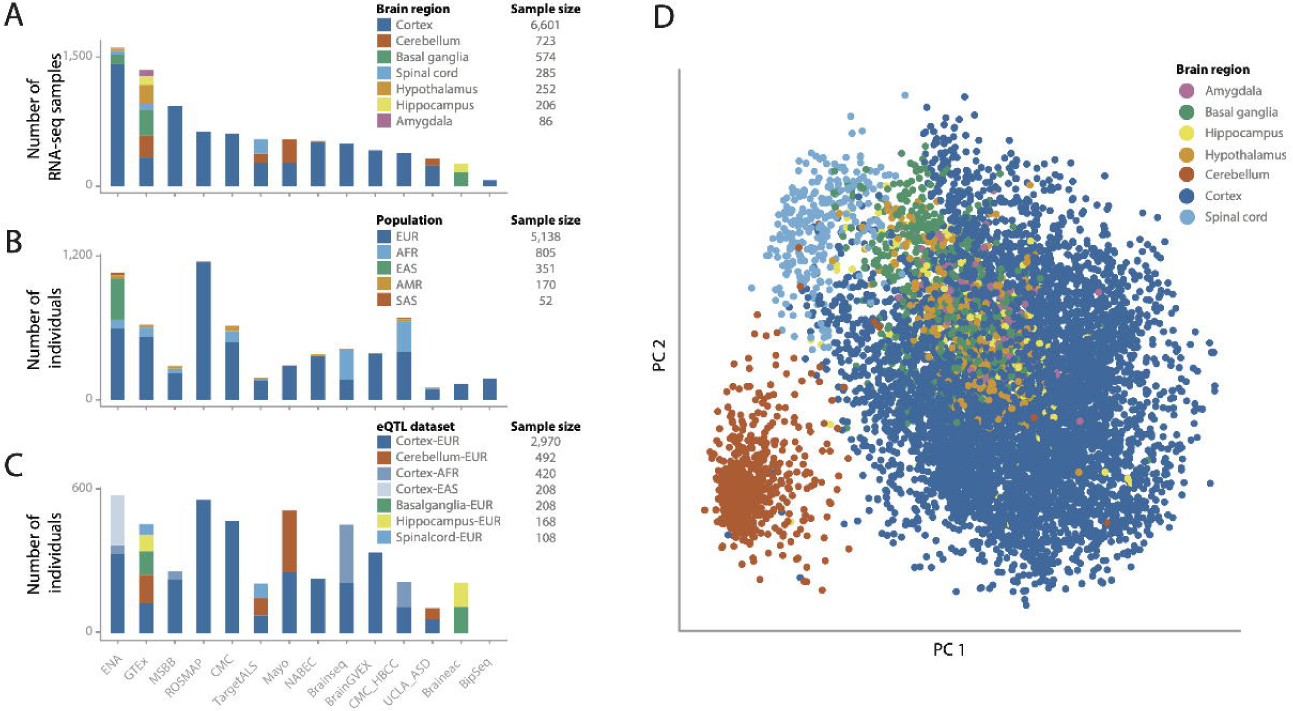
Overview of the datasets. **(A)** The number of samples per included cohort, with each color representing one of the 7 major brain regions. **(B)** The number of genotypes per cohort, with each color representing a population. **(C)** The number of individuals per cohort, with each color representing an eQTL dataset. The number of individuals is different from the intersection between the number of RNA-seq samples and number of genotypes, because not all samples with genotypes have RNA-seq samples and vice-versa, and some individuals with genotypes have multiple RNA-seq measurements. **(D)** PCA dimensionality reduction plot of the normalized expression data after covariate correction. Each dot represents an RNA-seq sample and is colored by brain region. The figure shows that the samples cluster mainly on brain region.

### 41% of the cortex *cis-*eQTL genes are regulated by multiple independent variants

Within each discovery dataset, we performed a sample-size weighted *cis*-eQTL meta-analysis on common variants (MAF>1%), within 1 megabase (Mb) of the transcription start site (TSS) of a protein-coding gene. We identified 1,317 (Basal ganglia-EUR), 6,865 (Cerebellum-EUR), 5,440 (Cortex-AFR), 11,803 (Cortex-EUR), 990 (Hippocampus-EUR), and 811 (Spinal cord-EUR) *cis*-eQTL genes (FDR<0.05; Figure 3A**; Supplementary Table 2**). *Cis*-eQTL effect directions were highly concordant between datasets included in the Cortex-EUR meta-analysis (median Spearman r=0.80; median allelic concordance=89%; **Supplementary Figure 6**), indicating robustness of the identified effects across datasets. We observed that significant *cis*-eQTL findings were sensitive to RNA-seq alignment strategies, and it is difficult to confidently ascertain *cis*-eQTLs in regions with multiple haplotypes represented on patch chromosomes, like the *MAPT* locus on 17q21 (**Supplementary Note, Supplementary Figures 7-9**). We next performed conditional analysis to identify independent associations in each *cis*-eQTL locus (e.g., secondary, tertiary and quaternary eQTLs). In Cortex-EUR, 4,791 genes had a significant secondary *cis*-eQTL (41% of *cis*-eQTL genes identified in this dataset). 1,658 genes had tertiary and 598 had quaternary *cis*-eQTLs. We also identified secondary associations for the other discovery datasets albeit to a lesser extent (Figure 3A**; Supplementary Table 2 and 3**).

**Figure 3.**
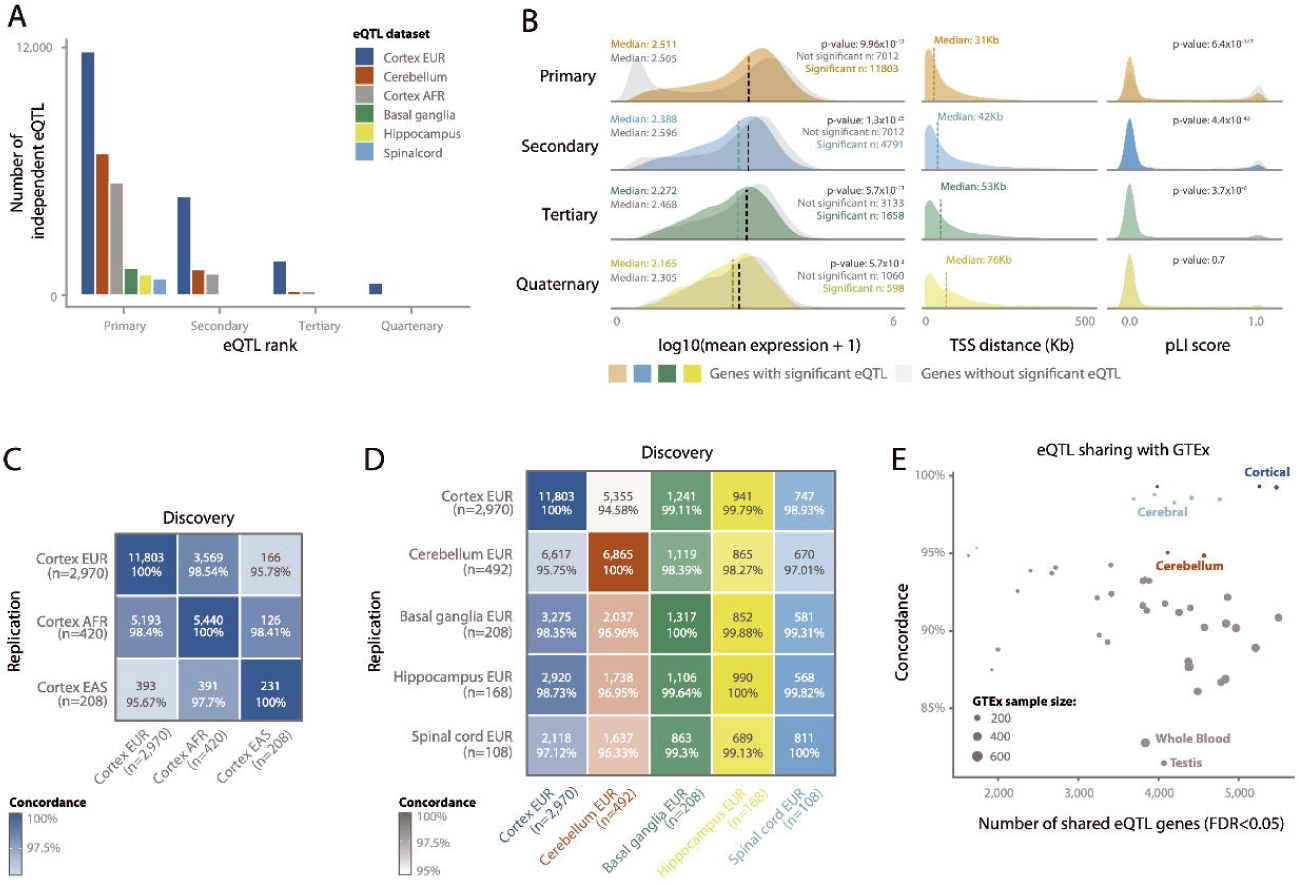
Conditional *cis*-eQTLs. **(A)** The number of conditional *cis-*eQTLs per eQTL dataset. **(B)** Comparison of characteristics between primary and non-primary eQTLs, where each row compares the eQTL genes for that rank with eQTL genes from the previous rank. P-values are calculated using a Wilcoxon test between significant and non-significant genes. (left) The difference in mean gene expression levels; (middle) the difference in distance between the most significant SNP-gene combination and the transcription start site (TSS); (right) the difference in probability for loss of function intolerance (pLI) score. For primary, secondary and quaternary eQTLs, non-significant eQTLs have higher pLI scores. **(C)** Replication of primary *cis*-eQTLs between the cortex eQTLs of different ethnicities and (**D**) the different brain regions for the European datasets. n indicates sample size of each dataset. Numbers in boxes indicate the number of eQTLs that are significant in both the discovery and the replication dataset, and the percentage of those that shows the same direction of effect. **(E)** Replication of primary *cis*-eQTLs of Cortex-EUR (discovery) in all the GTEx tissues (replication). Each dot is a different GTEx tissue, the x-axis is the number of eQTLs that is significant in both discovery and replication, and the y-axis is the percentage that shows the same direction of effect.

The properties of the Cortex-EUR *cis*-eQTLs conform to studies performed earlier in blood^14^ and brain^15^ (Figure 3B): primary lead *cis*-eQTL SNPs were generally located close (median distance: 31 kilobase; kb) to the transcription start site (TSS; Figure 3B) and *cis*-eQTL genes had a lower probability for loss of function intolerance (pLI; χ^2^ p=6.35×10^-147^). Genes with a *cis-*eQTL generally had a higher median expression than those without (Wilcoxon p-value: 9.96×10^-12^). Contrary to blood, where genes in the highest expression decile are the most likely to have a *cis*-eQTL, the third decile of gene expression had the most *cis*-eQTLs in cortex, and higher deciles had increasingly lower proportions of eQTLs (**Supplementary Note, Supplementary Figure 10A**). This could suggest that highly expressed genes in the cortex have tighter genetic regulation than highly expressed genes in the blood, although we did not observe differences when comparing variance per gene expression decile between blood and brain (**Supplementary Note, Supplementary Figure 10B**). Cortex-EUR *cis*-eQTL genes showed limited functional enrichment for human phenotype ontologies (HPO), GO ontologies and TRANSFAC^16^ transcription factor motifs (**Supplementary Figure 10C and D, Supplementary Table 4**). We observed similar patterns for secondary, tertiary and quaternary *cis-*eQTLs (**Supplementary Note).**

We investigated differences in *cis*-eQTLs due to ancestry, brain region, data sets and tissue type. We compared Cortex-EUR, Cortex-AFR and a smaller, East Asian cortex dataset (Cortex-EAS; n=208, limited to the ENA cohort; Figure 2C) and observed high concordance between the different ethnicities (>95.67%; Figure 3C). There was high concordance between different brain regions overall (>94.58%), though the cerebellum showed lower concordance with the cerebral brain regions (Figure 3D). Despite the limited sample size compared to Cortex-EUR, we identified 846 *cis*-eQTLs that were unique to Cerebellum-EUR (**Supplementary Figure 11A**). Of the 846 Cerebellum-EUR unique *cis*-eQTL genes, 184 had low gene expression levels in cortex, which may explain why they did not have a *cis*-eQTL in that tissue (**Supplementary Figure 11B, C, Supplementary Note**). For the remaining 662 genes that were highly expressed in both cortex and cerebellum, we performed functional enrichment of transcription factor binding sites (TFBS; **Supplementary Table 5, Supplementary Note**) and determined that these genes were enriched for TFBS of 101 distinct transcription factors. Five of these transcription factors had low gene expression in cortex and high expression in cerebellum (*EOMES, TFAP2B, TFAP2A, IRX1* and *IRX5,* **Supplementary Figure 11D**). These transcription factors might explain the difference in *cis*-eQTL genes found in cerebellum but not in cortex, while many of these *cis*-eQTL genes are expressed in both tissues. Next, we compared Cortex-EUR *cis*-eQTLs with different tissues from the GTEx project (Figure 3E**; Supplementary Figure 12, Supplementary Table 6**). There was high concordance in brain-related tissues (cerebral tissues, >98% and cerebellar tissues, >94%) compared to other tissue types, and the lowest concordance rates were observed in testis (84%) and whole blood (85%). We also compared Cortex-EUR *cis*-eQTLs with eQTLGen^17^, a large blood-based eQTL dataset (n=31,684; majority EUR ancestry) and observed a 76% concordance rate (**Supplementary Figure 13; Supplementary Table 7**) with a moderate correlation of *cis*-eQTL effect sizes (R_b_=0.54 including all eQTLs, or R_b_=0.62 when pruning genes within 1Mb)^18^, supporting the lower concordance observed in GTEx-blood. Since we found that 24% of the shared *cis*-eQTLs between blood and brain showed opposite allelic effects, these results suggest that with larger sample sizes, more tissue specific regulatory variants can be identified. If a causal tissue-specific regulatory variant resides on a haplotype that also contains a variant that is specific for another tissue, it is well conceivable that opposite allelic effects are going to be observed when contrasting eQTLs for these two tissues^19^. Since the procedures for eQTL mapping were identical between *MetaBrain* and eQTLGen, our results highlight the relevance of tissue-specific eQTL mapping to accurately assess the directionality of eQTLs, which can elucidate eQTLs with opposite allelic effects^20^. This direct comparison illustrates the importance of investigating the appropriate tissue type for the interpretation of GWAS signals.

### 8% of Cortex *cis*-eQTLs are mediated by cell type proportion differences

Cell type dependent eQTLs can be identified in bulk RNA-seq data by performing cell type deconvolution and determining cell type interaction eQTLs (ieQTLs)^3, 21, 22^. We predicted five major cell types using single cell RNA-seq derived signature profiles^23^. Of these, neurons were the most abundant cell type (median cell proportion: 32.8%), followed by endothelial cells (24.9%), macrophages (17.8%), oligodendrocytes (12.4%) and astrocytes (12.1%; **Supplementary Figure 14**). We predicted similar proportions for cerebellum as well as other brain regions. We observed that predicted cell proportions are different for spinal cord, showing a relatively low proportion of neuronal cells and high proportions of macrophage and oligodendrocytes compared to other brain tissues, as was previously reported^24^ (**Supplementary Figures 15 and 16)**. Predicted neuron proportions in both cortex and cerebellum were negatively correlated with the predicted proportions of other cell types, and predicted endothelial cell proportions were negatively correlated with predicted macrophage proportions (Figure 4A). Predicted cell type proportions were positively correlated with immunochemistry (IHC) counts from the ROSMAP cohort^25^, both overall (Spearman r=0.71; Figure 4B) and per individual cell type (Spearman r>0.1; Figure 4B). It is difficult to validate these cell type proportion predictions due to the small scale of the IHC experiment, but also because IHC and bulk RNA-seq reflect different aspects of gene or protein expression. Thus, there is a level of uncertainty for the expected proportion for each cell type^26, 27^.

**Figure 4.**
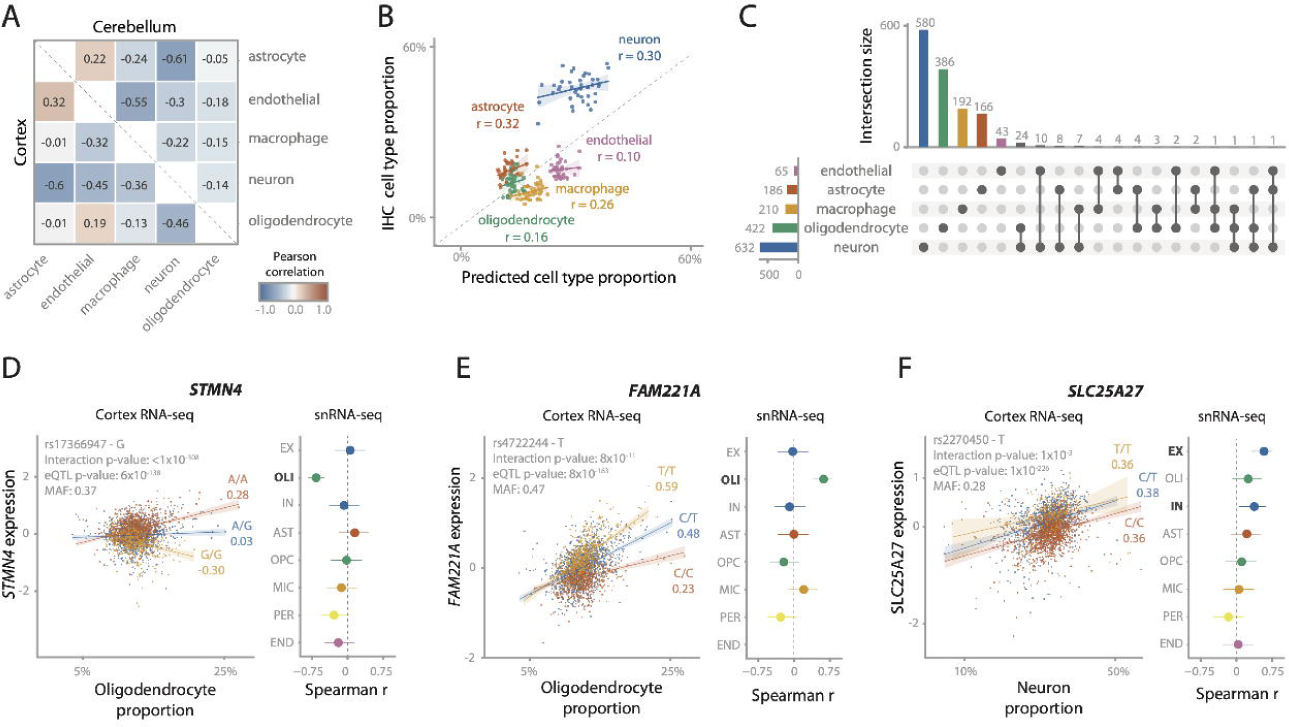
Cell type interacting eQTLs. **(A)** Spearman correlations between the 5 predicted cell count proportions. Lower triangle is within cortex samples, upper triangle is within cerebellum samples. **(B)** Predicted cell type proportions (x-axis) compared to cell type proportions measured using immunohistochemistry (IHC; y-axis) for 42 ROSMAP samples. Values in the plot are Pearson correlation coefficients. Cell count predictions for most cell types closely approximates actual IHC cell counts, although neurons are underestimated. **(C)** Number of cell type interacting eQTLs for Cortex-EUR deconvoluted cell types. The majority of interactions are with neurons and oligodendrocytes. Notably, most interactions are unique for one cell type in 90% of the cases. (**D, E, F**) Replication of cell type interacting eQTLs for *STMN4* (**D**), *FAM221A* (**E**) and *SLC25A27* (**F**), consisting of the scatterplot of the interaction eQTL in *MetaBrain* Cortex-EUR bulk RNA-seq (left) and a forest plot for the eQTL effect in the ROSMAP snRNA-seq data (right). Scatterplot: the x-axis shows the estimated cell type proportion, the y-axis shows the gene expression, each dot represents a sample. Colors indicate SNP genotype, with yellow being the minor allele. Values under the alleles are Spearman correlation coefficients. Forest plot: Spearman coefficients with effect direction relative to the minor allele when replicating the eQTL effect in ROSMAP single nucleus data (n=38). Error bars indicate 95% confidence interval. Each row denotes a cell type specific dataset: excitatory neurons (EX), oligodendrocytes (OLI), inhibitory neurons (IN), astrocytes (AST), oligodendrocyte precursor cells (OPC), microglia (MIC), pericytes (PER) and endothelial cells (END). Cell types highlighted in bold reflect the equivalent to the cell type used in the interaction eQTL.

With these predicted cell type proportions, we used DeconQTL^22^ to identify interaction-eQTLs (ieQTLs) by testing 18,850 *cis*-eQTLs in Cortex-EUR and 8,347 *cis*-eQTLs in cerebellum (including primary, secondary, tertiary and quaternary eQTLs). We identified 1,515 significant ieQTLs (8%) in at least one cell type (Benjamini-Hochberg; BH FDR<0.05) for Cortex-EUR (**Supplementary Table 8**). Of these, 632 (42%) were an ieQTL in neurons, likely because this is the most prevalent cell type. The majority of the ieQTLs (90.2%) were uniquely mapped to one cell type (Figure 4C). Although we observed a lower proportion of ieQTLs in cerebellum (126; 1.5%, **Supplementary Figure 17, Supplementary Table 8**), this is likely a power issue due to the smaller sample size. While we observed the most ieQTLs for neurons in cortex, the majority (n=106; 84%) of ieQTLs in cerebellum were mediated by astrocytes and macrophages.

We compared the allelic direction of the identified ieQTLs for each cell type with matching cell types from a single nucleus RNA-seq (snRNA-seq) dataset (ROSMAP cohort, n=39; **Supplementary Table 9**)^28^. When filtering on cell type mediated eQTLs by Decon-QTL (FDR<0.05), we observed a high average concordance in allelic direction for both the eQTL main effect (68%), as well as the direction of the interaction (68%; **Supplementary Figure 18B**). 106 of the cortex *cis*-ieQTLs were also significant (BH FDR<0.05) in the snRNA-seq datasets (63 in excitatory neurons and 43 in oligodendrocytes). Of these, 13 excitatory neuron and 21 oligodendrocyte ieQTLs were cell type mediated by the corresponding cell type in bulk with 100% allelic concordance (Decon-QTL; BH FDR<0.05; **Supplementary Figure 18D**). The ieQTLs replicating in oligodendrocytes included *STMN4, NKAIN1,* and *FAM221A* (Figure 4D **and E and Supplementary Figure 19A-C**), which have previously been identified as oligodendrocyte specific^29^. Additionally, this set of ieQTLs included *AMPD3* (rs11042811) and *CD82* (rs2303865), genes involved in the white matter microstructure^30^, suggesting a role for oligodendrocytes in this pathway. The ieQTLs replicating in excitatory neurons included *SLC25A27* (alias *UCP4;* Figure 4F **and Supplementary Figure 19D**), a gene principally expressed in neurons^31^ that modulates neuronal metabolism^32^. The eQTL SNP for this gene, rs2270450, is in high LD (r^2^=0.71) with a variant previously associated with schizophrenia^33^. Previous work has suggested a possible role of this gene in Parkinson’s disease^34, 35^. These results suggest that the decomposition of eQTLs to their relevant cell types in *MetaBrain* yields additional valuable information about the underlying biological mechanisms of genes and cell types of interest for genes associated with disease.

### Shared genetic effects between Cortex-EUR *cis-*eQTLs and brain-related traits

As one application of the *MetaBrain* resource, we linked *cis*-eQTLs to variants associated with brain-related traits and diseases. For this, we first evaluated linkage disequilibrium (LD) between the Cortex-EUR *cis*-eQTL SNPs with the strongest association signals and index variants identified in 1,057 GWASs of brain-related traits (**Supplementary Note, Supplementary Table 10**). We observed that 10% of brain-related trait SNPs for 242 eQTL genes were in LD with *cis*-eQTL SNPs (r^2^>0.8). This percentage marginally increased to 12% when secondary, tertiary and quaternary eQTL SNPs were included, indicating that the majority of LD overlap is driven by primary eQTL effects: primary eQTLs were 3.3-fold more likely to be in LD with a GWAS SNP (Fisher exact test p-value = 6.2×10^-16^; **Supplementary Note**).

To more formally test for overlap between GWAS and *cis*-eQTL signals, we conducted Mendelian randomization (MR) to test for a causal effect between gene expression and 31 neurological traits using *cis*-eQTLs as instruments (**Supplementary Table 11**). We computed a Wald ratio for each eQTL instrument, from which 1,192 Wald ratios out of 268,030 tested in total passed a suggestive p-value threshold (p<5×10^-5^ **Supplementary Table 12).** 120 of the *cis*-eQTL instruments from these suggestive findings were also cell type ieQTLs. We further prioritized our list of genes with evidence of Wald ratio effects by determining genetic colocalization between GWAS and *cis*-eQTL signals using coloc^36^. There were 159 significant Wald ratios that passed a strict Bonferroni correction (p<1.87×10^-7^) where the GWAS SNP and eQTL colocalized (PP4>0.7; Figure 5A**; Supplementary Figure 20**). 69 of these prioritized findings were associated with neurological and neuropsychiatric disease risk (**Table 1**). Three examples where MR and colocalization pointed to likely causal GWAS genes are reported below, for others, see **Supplementary Note, Supplementary Tables 11-16 and Supplementary Figures 21 and 22**.

**Figure 5.**
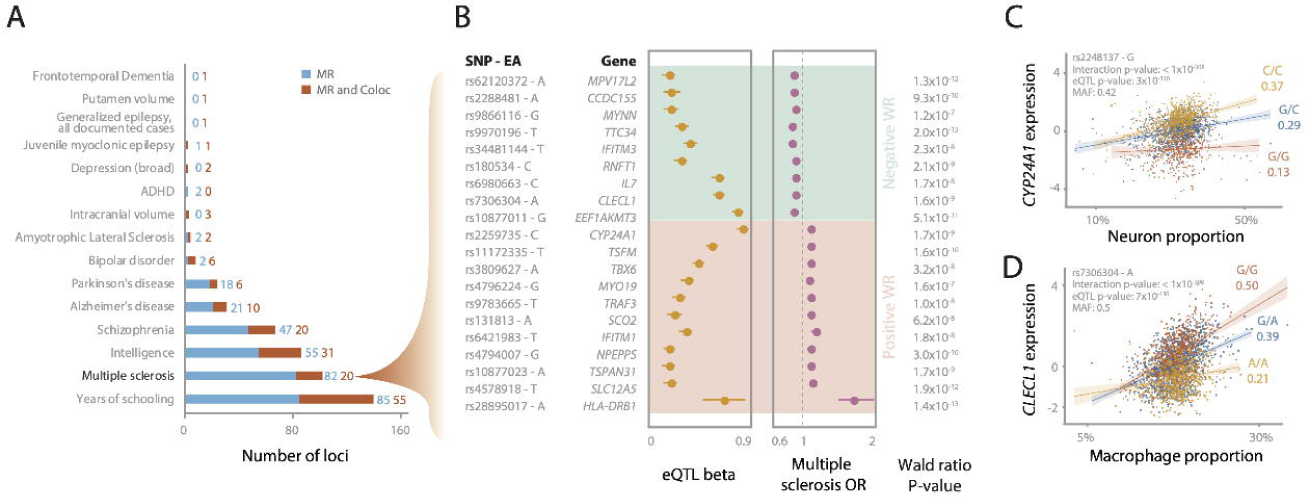
Mendelian randomization and colocalization of brain-related traits. **(A)** Number of significant Mendelian randomization (MR) signals (blue) and those with both MR and Coloc significant signals for 15 brain-related traits. **(B)** SNP and effect allele (EA), eQTL beta and GWAS odds ratio for 20 multiple sclerosis (MS) genes that are both MR and Coloc significant, and their Wald ratio p-value. Cell type interaction eQTL for *CYP24A1* (**D**) and *CLECL1* (**E**), showing interactions with predicted neuron, and macrophage proportions respectively. The x-axis shows the estimated cell type proportion, the y-axis shows the gene expression, each dot represents a sample. Colors indicate SNP genotype, with yellow being the MS risk allele. Values under the alleles are Spearman correlation coefficients.

**Table 1.** Prioritized genes from the Mendelian Randomization analysis on MetaBrain eQTLs versus brain related outcomes. Harmonized eQTL and GWAS SNP effects and single SNP Wald Ratio estimates are reported in the table for all genes with Wald Ratio effects at P<1.865×10^-7^. Columns are **genomic position**, **rsid** and **alleles** for SNP instrument (**EA**: Effect allele. **NONEA**: non-effect allele. **proxy SNP**: rsid of proxy SNP replacement used for outcome if instrument not present in GWAS), the SNP effects (**beta**, **SE**, **p**) for the MetaBrain eQTLs followed by the SNP effects for the brain related outcomes and then the Wald Ratio effects.

### MR comparison between blood and brain eQTL datasets

MR analysis for multiple sclerosis (MS)^37^ identified 102 instruments in 83 genes that passed the Bonferroni-adjusted p-value threshold (**Supplementary Table 12**). 20 of these findings passed colocalization (**Table 1**; Figure 5B). This included 11 genes for which MR suggested that increased gene expression and 9 genes where decreased gene expression may confer MS risk. Systematic comparison of the Wald ratio estimates for MS of 5,919 shared *cis-*eQTL genes between Cortex-EUR and eQTLGen (where the same gene was instrumented but could be with different SNPs)^17^ showed opposite directions of effect for 2,291 (38.7%) genes (**Supplementary Figure 23, Supplementary Table 14**). Agreement improved when the same SNP instrument was compared between studies, but discordance still remained high with 1,891 (26%) out of 7,274 *MetaBrain* Wald ratios showing opposite directionality to eQTLGen (**Supplementary Table 15)**. The notable discordance in the directionality of the blood and brain eQTLs underscore the importance of tissue-specific differences when interpreting transcriptomics data.

Of the 135 genes with MR findings in Cortex-EUR for MS, there were 28 genes without a significant eQTLGen instrument, including 3 genes (*SLC12A5*, *CCDC155* and *MYNN*) for which we found both MR significance and colocalization in *MetaBrain* (**Supplementary Note; Supplementary Table 16**. Comparing blood and brain gene expression levels for these genes in GTEx, *SLC12A5* had almost no expression in blood, while expression was comparable between tissues for *CCDC155* and *MYNN* (**Supplementary Note, Supplementary Figure 24**). The discrepancy in MR findings observed between Cortex-EUR and eQTLGen suggest tissue-dependent genetic effects for MS.

### MR and colocalization analysis links multiple sclerosis GWAS loci to cell type specific eQTLs for *CYP24A1* and *CLECL1*

Two MS genes, *CYP24A1* and *CLECL1*, showed cell type specific *cis-*eQTLs (Figure 5C and D). Another gene that was previously suggested to be neuron specific^38^, *SLC12A5*, did not show a significant ieQTL in our data. Our analysis used rs2259735 as the Cortex-EUR eQTL instrument variable and suggested that higher expression of *CYP24A1* is associated with increased MS risk (MR Wald ratio=0.13, p=1.7×10^-9^). We also observed colocalization of the *cis*-eQTL and the MS GWAS signal at this region (coloc PP4=0.99), suggesting the same underlying genetic signal. Furthermore, ieQTL analysis showed increasing expression of *CYP24A1* with increasing neuronal proportions for the MS risk allele rs2248137 (interaction beta=2.85; interaction FDR=1×10^-308^; Figure 5C). Rs2248137 has previously been associated with MS^39^ and is in strong LD with SNP rs2259735 (r^2^=0.9). *CYP24A1* is a mitochondrial cytochrome P450 hydroxylase that catalyzes the inactivation of 1,25-dihydroxyvitamin D_3_ (calcitriol), the active form of vitamin D^40^. Loss of function mutations in *CYP24A1* increase serum calcitriol and cause hereditary vitamin D-mediated PTH-independent hypercalcemia^41, 42^. In the brain, vitamin D plays vital functions in regulating calcium-mediated neuronal excitotoxicity, reducing oxidative stress and regulating synaptic activity^43^. Epidemiological studies have proposed vitamin D deficiency as a risk factor for MS^44, 45^, which has recently been validated through MR^46–48^. Our findings are consistent with a previous report of a shared MS GWAS signal and *CYP24A1 cis*-eQTL signal with frontal cortex but not white matter, using a brain eQTL dataset derived from expression microarrays to confirm the findings in the same direction of effect^49^.

As another MS signal that passed MR and colocalization, decreased expression of *CLECL1* was associated with increased MS risk (MR Wald ratio=-0.16, p=1.58×10^-9^, coloc PP4>0.92). The ieQTL analysis indicated that the rs7306304 allele increased expression of *CLECL1* with increasing macrophage proportion (interaction beta=-3.65; interaction FDR=1×10^-308^, Figure 5D), confirming a previous finding of a microglia cell-type specific *cis*-eQTL for *CLECL1* at this MS risk locus^39^. Rs7306304 is in strong LD with the MS lead SNP, rs7977720 (r^2^=0.84)^39^. *CLECL1* encodes a C-type lectin-like transmembrane protein highly expressed in dendritic and B cells that has been proposed to modulate immune response^50^. *CLECL1* was previously found to be lowly expressed in cortical bulk RNA-seq data, while having a 20-fold higher expression in a purified microglia dataset^39^, suggesting that decreased *CLECL1* expression increases MS susceptibility through microglia-mediated dysregulation of immune processes in the brain.

### *MetaBrain* allows for the identification of *trans*-eQTLs

*Trans*-eQTL analysis can identify the downstream transcriptional consequences of disease associated variants. However, we have previously observed in blood that *trans*-eQTL effect-sizes are usually small. Here we studied whether this applies to brain as well. In order to maximize sample size and statistical power, we performed a *trans*-eQTL analysis in 3,111 unique individuals. We reduced the number of tests performed by limiting this analysis to 130,968 unique genetic variants: these include variants that have been previously found to be associated with diseases and complex traits through GWAS and variants that were primary, secondary, tertiary or quaternary lead *cis*-eQTL SNPs from any of the aforementioned discovery datasets.

We identified 3,940 *trans*-eQTLs (FDR<0.05), of which 2,589 (66%) were significant after removing *trans*-eQTLs for which the gene that partially map within 5Mb of the *trans*-eQTL SNP (**Supplementary Note**; Figure 6A**; Supplementary Table 17**). These 2,589 eQTLs reflect 373 unique SNPs, and 1,263 unique genes. 222 (60%) of the *trans*-eQTL SNPs were a *cis*-eQTL SNP, of which 42 (19%) were a *cis*-eQTL index SNP in Cortex-EUR, and 22 (10%) in tissues other than cortex. This suggests that *trans*-eQTLs can also be observed for *cis*-eQTLs index SNPs identified in other tissues (**Supplementary Table 17**).

**Figure 6.**
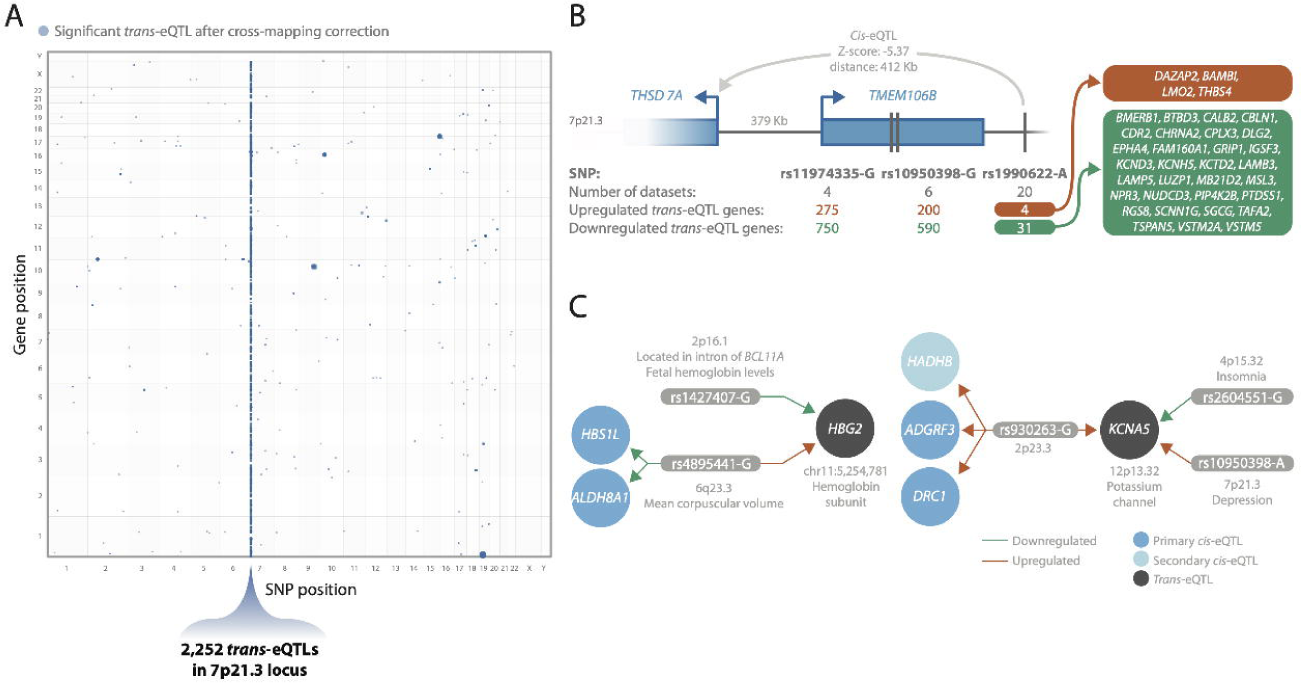
*Trans*-eQTLs in brain. **(A)** Location of identified *trans*-eQTLs, with the SNP position (x-axis) and gene position (y-axis) in the genome. Size of the dots indicate the p-value of the *trans*-eQTL (larger is more significant). 7p21.3, the locus with most (83%) of the *trans*-eQTLs, is highlighted. **(B)** Three SNPs in the 7p21.3 locus and the number of datasets and number of up- and down-regulated *trans*-eQTL genes each SNP has. For rs1990622, a SNP associated with frontotemporal lobar degeneration, the 35 genes it affects in *trans* and the 1 gene it affects in *cis* are shown. **(C)** Two examples of convergent effects, where multiple independent SNPs affect the same genes in *trans*. Left: *trans-*eQTLs of rs1427407 and rs4895441 on *HBG2* and right *trans-*eQTL of rs930263, rs2604551, and rs10950398 on *KCNA5*.

1,060 (83%) of the observed *trans*-eQTL genes were affected by 3 variants at 7p21.3 (rs11974335, rs10950398 and rs1990622, LD r^2^>0.95; Figure 6A **and B**; **Supplementary Table 17**). This locus is associated with several brain-related traits, including frontotemporal lobar degeneration^51^ and major depressive disorder^52^ (**Supplementary Table 17**). The *trans*-eQTL SNP rs1990622 in this locus is the lead GWAS SNP for the TDP-43 subtype of frontotemporal lobar degeneration (FTLD-TDP)^53^, just downstream of *TMEM106B*. Matching previous reports^54, 55^, we observed that this locus was associated with predicted neuron proportions (**Supplementary Tables 18-20).** Moreover, the predicted neuronal proportions were lower in AD cases than controls (**Supplementary Figure 25**), which may explain why a large number of *trans*-eQTLs signals at this region were most pronounced in the AMP-AD datasets and had stronger effect sizes in AD samples (**Supplementary Figure 26 and 27**). We performed functional enrichment on the *trans*-eQTL genes using g:Profiler^56^and observed that upregulated *trans*-eQTL genes were enriched for neuron related processes such as synaptic signaling (p=1.3×10^-28^) and nervous system development (p=2.9×10^-21^). Downregulated genes were enriched for gliogenesis (p=1.6×10^-8^) and oligodendrocyte differentiation (p=3.1×10^-6^; **Supplementary Table 21**). Surprisingly, 21 of these *trans*-eQTLs were also significant (BH FDR<0.05) in the snRNA-seq data of excitatory neurons with 100% allelic concordance (**Supplementary Figure 28; Supplementary Table 22**), suggesting that some of these *trans*-eQTLs might not be driven by differences in neuron proportions. A detailed description of this locus can be found in the **Supplementary Note**.

We observed *trans*-eQTLs from multiple independent genomic loci for 14 genes, suggesting convergent *trans*-eQTL effects (**Supplementary Table 17)**. The genes with these convergent *trans*-eQTL effects were previously associated with immunological phenotypes (*HBG2, PIWIL2,* and *SVEP1*), brain-related phenotypes (*DAZAP2*), immunological and brain-related phenotypes (*HMCES, KCNA5, MBTPS1, PRPF19, PTH2R* and *RFPL2*) or other phenotypes (*ANKRD2, PEX12, PROM1* and *ZNF727*).

Encouragingly, some of these convergent *trans*-eQTLs have already been previously identified in blood. For example, two independent variants (rs1427407 on 2p16.1 and rs4895441 on 6q23.3) affected hemoglobin subunit gamma-2 (*HBG2*) on 11p15.4 in *trans* (Figure 6C). These variants have previously been associated with fetal hemoglobin levels^57–59^ and various blood cell counts.

We also observed converging effects that were not identified in blood. For instance, *KCNA5* (12p13.32) was affected by variants from three independent loci at 2p23.3 (rs930263), 4p15.32 (rs2702575 and rs2604551) and 7p21.3 (rs10950398 and rs11974335) as described in Figure 6C; **Supplementary Table 17**. *KCNA5* encodes the potassium voltage-gated channel protein Kv1.5. Potassium voltage-gated channels regulate neuron excitability among other functions, and blockers for these channels have been suggested as a therapeutic target for multiple sclerosis patients^60^. Furthermore, *KCNA5* has previously been associated with cardiovascular disease^61^, and has been suggested to modulate macrophage and microglia function^62^. Three *cis*-eQTLs were associated with rs930263, including *ADGRF3*, *DRC1*, and a secondary eQTL on *HADHB*. rs930263 was previously associated with sleep dependent LDL levels^63^ and several blood metabolite levels^64–67^. The 4p15.32 locus was previously associated with insomnia and adult height^68^ and the 7p21.3 locus with depression and blood protein levels. These results thus suggest that several sleep related variants affect potassium voltage-gated regulation of neuron excitability.

This is the first report of *trans*-eQTLs in the brain cortex for many of the variants identified, and our results indicate that many of these signals are brain-specific. We observed the *trans*-eQTL effect-sizes in brain are usually small, similar to what we previously observed in blood, emphasizing the importance of increasing the sample-size of brain eQTL studies.

### Brain co-regulation networks improve GWAS interpretation

We generated brain-region specific co-regulation networks based on the RNA-seq data from 8,544 samples **(Supplementary Note, Supplementary Figures 29-30)**. We previously have done this for a heterogenous set of RNA-seq samples spanning across all available tissue types and cell lines (n=31,499)^69, 70^, which showed that such a co-regulation network can be informative for interpreting GWAS studies^69^ and helpful in the identification of new genes that cause rare diseases^70^.

We applied a new approach (*’Downstreamer’,* in preparation, see **Supplementary Note**) that improves upon DEPICT, our previously published post-GWAS pathway analysis method^69^. *Downstreamer* can systematically determine which genes are preferentially co-regulated with genes that reside within GWAS loci. It does not use a significance threshold for a GWAS, but instead uses all SNP information. In addition, *Downstreamer* accounts for LD and uses rigorous permutation testing to determine significance levels and control for Type I errors.

We applied *Downstreamer* to schizophrenia (SCZ)^71^, PD^72^, MS^37^, AD^73^ and ALS GWAS summary statistics (**Supplementary Table 23-27**), using three different brain-derived co-regulation networks: one based on all 8,544 brain samples, one limited to 6,527 cortex samples and one limited to 715 cerebellum samples. We observed that there were multiple sets of genes that showed strong co-regulation with genes inside the GWAS loci for these diseases. For MS and AD, these were mostly immune genes, whereas for PD, ALS and SCZ these were genes that are specifically expressed in brain (**Supplementary Table 23-27**).

For ALS, we applied *Downstreamer* to summary statistics from a recent meta-analysis in individuals from European ancestry (**Supplementary Table 28**), and a trans-ethnic meta-analysis including European and Asian individuals (EUR+ASN; **Supplementary Table 23;** van Rheenen *et al*., manuscript in preparation). To look for contributions of non-neurological cell types and tissues, we first used the previously published heterogenous network^70^ that comprises many different tissues and cell types, but did not identify genes that were significantly enriched for co-regulation with genes inside ALS loci. However, when we applied our method to the different brain co-regulation networks, we identified a set of 27 unique co-regulated genes (EUR+ASN summary statistics; Figure 7A**; Supplementary Table 23**), depending on the type of brain co-regulation network used. *HUWE1* was shared between the brain and cortex co-regulation network analysis, while *UBR4* was shared between the cortex and cerebellum analysis. *UBR4* is a ubiquitin ligase protein expressed throughout the body. A private *UBR4* mutation, segregated with episodic ataxia in a large three-generation Irish family, implicates its role in muscle coordination^74^. *UBR4* interacts with the Ca^2+^ binding protein, calmodulin and Ca^2+^ dysregulation has been linked to proteins encoded by ALS disease genes and motor neuron vulnerability^75^. We observed in the *Downstreamer* findings that many of these prioritized genes are co-regulated with each other (Figure 7B), and using our recently developed clinical symptom prediction algorithm^70^, there was an enrichment of genes implicated in causing gait disturbances (Figure 7C). These genes are associated with ALS (highlighted in blue), brain-related disorders (including *DNAJC5*, *HTT*, *HUWE1*, *TSC1* and *YEATS2*) or muscle-related disorders (including *KMT2B*). While various loci have been identified for both familial and sporadic forms of ALS, the function of the positional candidate genes within these loci is still unclear. Our *Downstreamer* analysis identified genes that show strong coregulation with positional candidate genes inside ALS loci, suggesting that these positional candidates must have a shared biological function.

**Figure 7.**
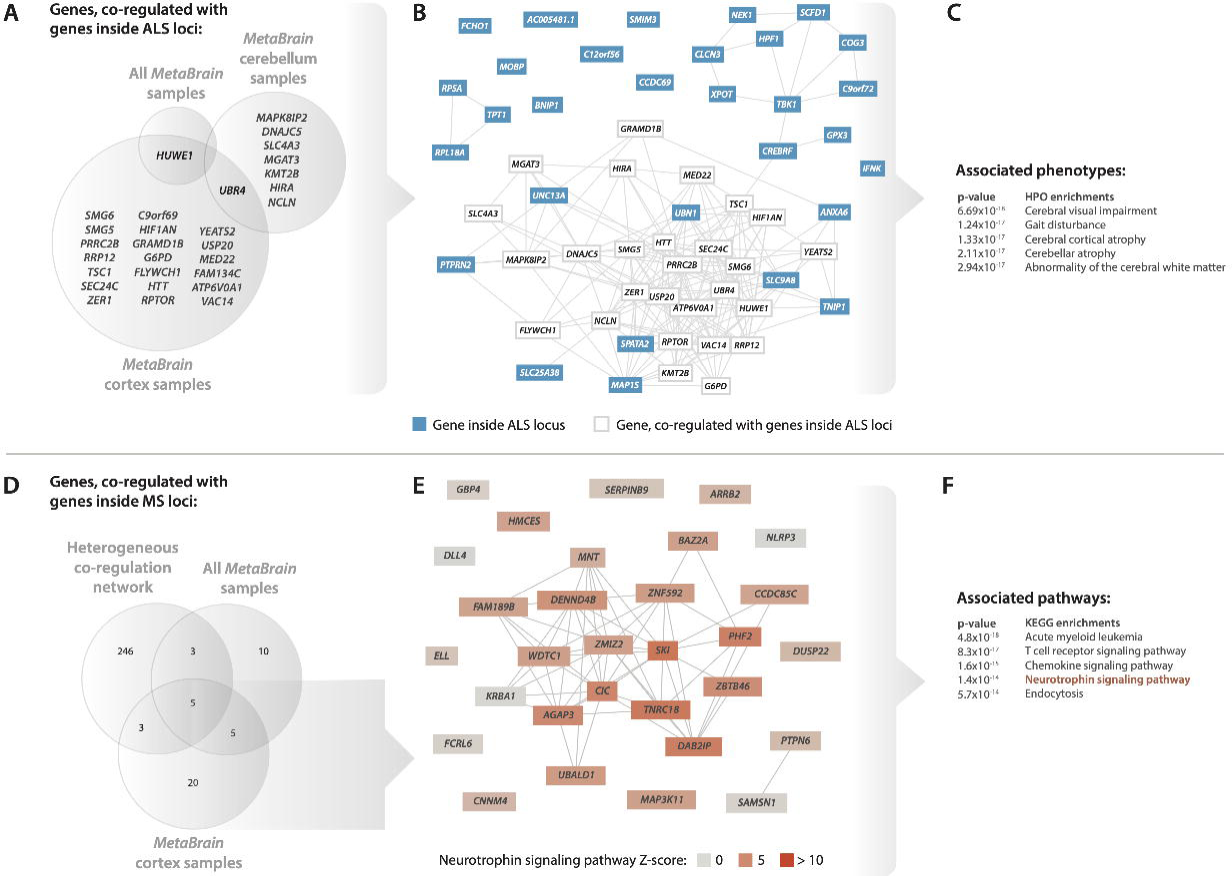
Gene co-regulation. **(A)** Genes that are co-regulated with genes that are within amyotrophic lateral sclerosis (ALS) loci. Co-regulation scores between genes are calculated using all *MetaBrain* samples, *MetaBrain* cerebellum samples, or *MetaBrain* cortex samples. Except for *URB4, c*ortex and cerebellum networks find different co-regulated genes for ALS. **(B)** Co-regulation network using all *MetaBrain samples* for all genes prioritized for ALS by *Downstreamer*. **(C)** Top 5 Human Phenotype Ontology (HPO) enrichments for the *Downstreamer* prioritized ALS genes. **(D)** Genes that are co-regulated with genes that are within multiple sclerosis loci. Co-regulation scores between genes are calculated using a heterogeneous multi-tissue network, *MetaBrain* cerebellum samples, or *MetaBrain* cortex samples. Most genes are found using a large heterogenous co-regulation network. **(E)** Co-regulation network of all *MetaBrain* samples for 33 genes prioritized by *Downstreamer* in cortex. Colors indicate the neutrophin signaling pathway enrichment Z-scores. **(F)** Top 5 KEGG enrichments for the *Downstreamer* prioritized multiple sclerosis genes in cortex.

For MS, the heterogeneous network, including many blood and immune cell type samples, identified 257 unique genes that showed significantly enriched co-regulation with genes inside MS loci (Figure 7D; **Supplementary Table 27**), and many were immune genes, which is also expected for this disease. However, when we applied the brain co-regulation networks, we identified a much smaller set of genes, and these genes showed strong enrichment for genes involved in the neurotrophin signaling pathway (Figure 7E and F). Neurotrophins are polypeptides secreted by immunological cell types. In the brain, neurotrophin concentrations are important to promote the survival and proliferation of neurons as well as synaptic transmission. In MS patients, neurotrophin reactivity is higher in MS plaques, whereby neurotrophins are released by peripheral immune cells directly to the inflammatory lesions, suggesting a protective role of this signaling process^76, 77^. Neutrophins are also released by glial cells in the brain, including microglia and astrocytes, and their role in stimulating neuronal growth and survival could also contribute to an overall neuroprotective effect^78^. In the heterogeneous network, we observed high expression for these genes in immune-related tissues (**Supplementary Figure 31A**), supporting the “outside-in hypothesis” that the immune system may be a potential trigger for MS^37, 79^. The brain specific network showed high expression in spinal cord and cerebellum but lower expression in cortex samples (**Supplementary Figure 31B**), which could be highlighting the specific biological processes taking place in these CNS regions that lead to disease. For example, the cerebellum is responsible for muscle coordination and ataxia occurs in approximately 80% of MS patients with symptoms^80^. We speculate that both dysregulation of the immune system and dysregulation of certain neurological processes is a prerequisite for developing MS.

## Discussion

We here describe an integrated analysis of the effects of genetic variation on gene expression levels in brain in over 3,000 unique individuals. This sample size yielded sufficient statistical power to identify robust *cis*-eQTLs and to our knowledge for the first-time brain *trans*-eQTLs that emanate from SNPs previously linked to neurodegenerative or psychiatric diseases.

We compared *cis*-eQTLs in *MetaBrain* to *cis*-eQTLs in eQTLGen from a set of 31,684 blood samples. We observe a large proportion of shared *cis*-eQTLs between brain and blood, most of which have the same allelic direction of effect. Our analysis also permitted us to identify *cis-*eQTL effects that are independent of the primary *cis*-eQTLs. Some of these independent effects reflect SNPs that are also the index variants for several neurological and psychiatric disorders, making them particularly interesting for subsequent follow-up. Recent observations have revealed that SNPs with the strongest *cis*-eQTL effects are depleted for GWAS associations^81^. Thus, secondary, tertiary or quaternary *cis*-eQTL SNPs could potentially be even more interesting to follow-up than certain primary *cis*-eQTL SNPs to link association signals to function.

We studied different regions in the brain, permitting us to identify brain-region specific eQTLs. For this, to exclude spurious differences that may arise from different cell type proportions across brain regions, we first inferred cell type percentages for the major brain cell types. We then applied an eQTL interaction model (i.e., using the cell type percentage x genotype as interaction term), permitting us to identify 1,515 *cis*-eQTLs that show cell type specificity. Most of these cell type dependent effects were observed for oligodendrocytes and neurons, the two most common cell types in the brain for which statistical power to observe such effects was the strongest. Still, we could identify 461 cell type dependent eQTLs also for macrophages, endothelial cells, or astrocytes. While we found strong concordance with immunohistochemistry results, our findings are largely based on a deconvolution approach, which in future studies will benefit from validation in purified cell types, e.g. using population-based single-cell RNA-seq datasets as they are now becoming available^82, 83^. Such single-cell eQTL studies can gain substantial statistical power by limiting analyses to the large set of primary, secondary, tertiary and quaternary *cis*-eQTLs our study reveals for bulk brain samples.

To our knowledge, this is the best powered Mendelian randomization and colocalization analysis using brain *cis*-eQTLs as instruments for bipolar disease, epilepsy, frontotemporal dementia, multiple sclerosis, cognitive function and years of schooling GWAS outcomes. Interestingly, also for schizophrenia three signals for *CILP2*, *MAU2* and *TM6SF2* met our criteria that had not been reported in a recent psychiatric genomics consortium study^84^, further emphasizing the value of our well-harmonized, large eQTL data set in the tissue type of interest (**Supplementary Note**). Our results also identify increased *CYP24A1* expression as associated with multiple sclerosis risk and propose neurons as the most susceptible cell type to *CYP24A1* expression changes and likely active vitamin D levels. The potentially novel role of *CYP24A1* in brain could play an important role in MS etiology, as may lowered expression of *CLECL1* in microglia.

The 2,589 identified *trans*-eQTLs allowed us to gain insights into downstream molecular consequences of several disease-associated genetic variants. Our *trans*-eQTL analysis focused on a single brain region and SNPs with a known interpretation (i.e. trait-associated variants and *cis*-eQTL SNPs). We therefore expect that a genome-wide approach will identify many more *trans*-eQTLs. 2,218 of the *trans*-eQTLs were located in a 7p21.3 locus and the genes were strongly correlated with neuron proportions, indicating that cell type proportions can heavily impact *trans*-eQTL identification. However, 21 of these *trans*-eQTLs replicated in snRNA-seq data, suggesting that some of these *trans*-eQTLs may also exist in single cells. Excluding the 7p21.3 locus, we identified 371 *trans*-eQTLs located elsewhere in the genome, which are less likely due to neuron proportion differences. For several neurological and psychiatric conditions, our analyses indicate pathways that may help to elucidate disease causes and putative intervention points for future therapies.

We used the brain-specific co-regulation networks to study several brain-related GWAS studies, with the aim to prioritize genes that show significantly enriched co-regulation with genes inside the associated GWAS loci. For ALS this revealed a limited, but significant set of genes which do not map within associated ALS loci, but that link genes within multiple ALS loci. Follow-up research on these prioritized genes might therefore help to better understand the poorly understood causal pathways that cause ALS. While it is tempting to speculate that these prioritized genes might represent genes that could serve as potential targets for pharmaceutical intervention, follow-up research is needed in order to establish whether these genes play a relevant role in ALS.

Our study had several limitations. For instance, we performed single tissue eQTL analyses that were limited to a single RNA-seq sample per individual, excluding many RNA-seq samples from the analysis. A joint analysis across tissues, including multiple RNA-seq samples per individual using for example random effects models would further improve power^85, 86^, which would be especially useful for the future identification of *trans*-eQTLs. Additionally, LD overlap analysis, Mendelian randomization and colocalization are sensitive to many factors, including eQTL and GWAS study sample size, effect size, variant density, LD structure and imputation quality. Differences between study designs may consequently influence the results of such analyses. For example, our colocalization and LD overlap analysis did not include the *MAPT* gene for Alzheimer’s disease. The effect sizes of the *cis*-eQTLs for this gene were limited in our study, since our alignment strategy could not account for the different long-range haplotypes in this locus causing the H1/H2 haplotype separating SNP rs8070723 to have a p-value of 0.2 (**Supplementary Note**). We note that this might be an issue for other genes as well. Future studies using graph-based alignment tools or long read sequencing methods would be required to ultimately determine the true effects on such genes. Our approach combined Mendelian randomization and colocalization, as it is possible for the *cis*-eQTL instrument to coincidentally share association with the GWAS trait due to surrounding LD patterns in the genomic region. We opted to perform single SNP MR because other approaches, such as inverse variance weighted^87^ (IVW) MR, pool the estimates across many SNP instruments, which for many genes were not available. Potentially, methods such as IVW could be applied to our dataset in the future when genome-wide *trans*-eQTL analysis would identify many more independent instruments per gene. However, MR analyses using QTLs could be susceptible to confounding because of horizontal pleiotropy^88^, where a single gene is affected by multiple indirect effects, which is likely to be exacerbated by including *trans*-eQTLs. Our colocalization analysis used a more lenient posterior probability (PP4) threshold of >0.7, which we selected because we performed colocalization only in loci having a significant MR signal, limiting potential false positives. However, our colocalization approach assumed the presence of a single association in each locus, which might not be optimal for *cis*-eQTL loci harboring multiple independent variants, such as for the *TREM2* gene (**Supplementary Note**). Consequently, our approach may have not detected colocalizing signals in some loci. Recently, colocalization methods were published^89^ that do not have this assumption, and consequently may improve future colocalization results.

With the numbers of GWAS loci for brain-related traits and diseases steadily climbing, we expect that our resource will prove itself as a highly valuable toolkit for post-GWAS brain research and beyond. Among others, we demonstrate how our dataset can be utilized to disambiguate GWAS loci, point to causal pathways and prioritize targets for drug discovery. To our knowledge, this is the largest non-blood eQTL analysis ever conducted, providing insights into the functional consequences of many disease associated variants. We expect that through future integration with single-cell eQTL studies that have higher resolution but lower power, our results will help to pinpoint transcriptional effects in specific brain cell types for many disease-associated genetic variants.

## Methods

### Dataset collection and description

We collected human brain bulk RNA-seq datasets from different resources. Briefly, we collected previously published samples from the AMP-AD consortium^6^ (AMP-AD MAYO^6^, ROSMAP^6^ and MSBB^6^), Braineac^7^, the PsychENCODE consortium^8^ (Bipseq^4^, BrainGVEX^4^, CMC^9^, GVEX, and UCLA_ASD^4^) from Synapse.org using the Python package synapseclient^90^. The NABEC and GTEx datasets were retrieved from NCBI dbGaP, and TargetALS data was provided directly by the investigators. For an overview of the number of samples per dataset, see **Supplementary Table 1**.

Additionally, we collected public brain bulk RNA-seq samples from the European Nucleotide Archive (ENA; **Supplementary Table 28**). To select only the brain samples, we first downloaded the SkyMap database^91^, which provides readily mapped read counts and sample annotations. We performed rigorous quality control on this dataset, and selected ENA, excluding for example brain cell lines, brain cancer samples, and samples with RNA spike ins (See **Supplementary Note** for more details on this method, **Supplementary Figure 1**), resulting in 1,759 samples, and 9,363 samples when combined with the previously published datasets (**Supplementary Table 1**).

### RNA-seq data

RNAseq data was processed using a pipeline built with molgenis-compute^92^. FASTQ files were aligned against the GENCODE^93^ v32 primary assembly with STAR^94^ (version 2.6.1c), while excluding patch sequences (see **Supplementary Note**) with parameter settings: outFilterMultimapNmax = 1, twopassMode Basic, and outFilterMismatchNmax = 8 for paired-end sequences, outFilterMismatchNmax = 4 for single-end sequences. Gene quantification was performed by STAR, similar to gene quantification using HTSeq^95^ with default settings. The gene counts were then TMM^96^ normalized per cohort using edgeR^97^ (version 3.20.9) with R^98^ (version 3.5.1).

To measure FASTQ and alignment quality we used FastQC^99^ version 0.11.3), STAR metrics, and Picard Tools^100^ (version 2.18.26) metrics (MultipleMetrics, and RNAseqMetrics). Samples were filtered out if aligned reads had <10% coding bases (**Supplementary Figure 3A**), <60% reads aligned (**Supplementary Figure 3B**), or <60% unique mapping. 117 of the RNA-seq samples did not pass this filter, mostly from GTEx^97^. The other quality measurements were visually inspected but contained no outliers.

RNA-sequencing library preparation, and other technical factors can greatly influence the ability to quantify of gene expression. Therefore, for a given sample such factors often influence the total variation. For example, such issues can be caused by problems during RNA-seq library preparation that led to an increased number of available transcripts to quantify, or conversely, a lack of variation in quantified transcripts (compared to other samples in the dataset). We therefore opted to identify RNA-seq outliers that were not explained by poor RNA-seq alignment metrics. For this purpose, we performed PCA on the RNA data prior to normalization: we reasoned that the first two components capture excess or depletion of variation caused by technical problems. We identified 20 samples that were outliers in the PCA plot of the RNA-seq data, where PC1 was more than 4 standard deviations from the mean (**Supplementary Figure 3A**). Twenty outlier samples were removed and the principal components were recalculated (**Supplementary figure 3B**). We detected and removed 45 additional outlier samples. We confirmed no additional outlier samples in the third iteration and principal component calculation, (**Supplementary Figure 3C**) and 8,868 samples were taken through additional QC.

We next removed genes with no variation and then log2-transformated, quantile normalized and Z-score transformed the RNA-seq counts per sample. PCA on the normalized expression data showed that datasets strongly cluster together (**Supplementary Figure 4A**), likely due to dataset specific technical differences (e.g., single-end versus paired-end sequencing). To correct for this, the normalized expression data was correlated against 77 covariates from different QC tools (FastQC^99^, STAR^94^, and Picard Tools^100^), such as percent protein coding, GC content, and 5’ prime/3’ prime bias. The top 20 correlated technical covariates (% coding bases, % mRNA bases, % intronic bases, median 3’ prime bias, % usable bases, % intergenic bases, % UTR bases, % reads aligned in pairs, average mapped read length, average input read length, number of uniquely mapped reads, % reads with improper pairs, number of reads improper pairs, total sequences, total reads, % chimeras, number of HQ aligned reads, number of reads aligned, HQ aligned Q20 bases, HQ aligned bases) were regressed out of the expression data using a linear model. After covariate correction, clustering of datasets in PC1 and PC2 were no longer present (**Supplementary Figure** 4**B**).

Our collection of RNA-seq samples consisted of 36 different tissue labels, many of which were represented by only a few samples. Therefore, we next defined major brain regions present in our dataset, including samples from amygdala, basal ganglia, cerebellum, cortex, hippocampus and spinal cord. We noted that some samples (especially from ENA) were not annotated with a specific major brain region. To resolve this, we performed PCA over the sample correlation matrix and then performed k-nearest neighbors on the first two PCs (k=7) to classify samples to the major brain regions. Using this approach, we defined a set of 86 amygdala, 574 basal ganglia, 723 cerebellum, 6,601 cortex, 206 hippocampus, 252 hypothalamus and 285 spinal cord samples (**Supplementary Table 1,** Figure 2A).

### Genotype data and definition of eQTL datasets

The genotype data for the included datasets was generated using different platforms, including genotypes called from whole genome sequencing (WGS; AMP-AD, TargetALS^12^, GTEx^3^), genotyping arrays (NABEC^11^, Braineac^7^), and haplotype reference consortium (HRC)^101^ imputed genotypes (PsychENCODE datasets), or were called from RNA-seq directly (ENA dataset; see **Supplementary Note**). In total, 22 different genotyping datasets were available, reflecting 6,658 genotype samples (**Supplementary Table 1)**.

We performed quality control on each dataset separately, using slightly different approaches per platform. For the array-based datasets, we first matched genotypes using GenotypeHarmonizer^102^ using 1000 genomes phase 3 v5a (1kgp) as a reference, limited to variants having MAF >1%, <95% missingness and Hardy-Weinberg equilibrium p-value <0.0001. Genotypes were then imputed using HRC v1.1 as a reference on the Michigan imputation server^103^. In all HRC imputed datasets, variants with imputation info score <0.3 were removed. For the WGS datasets, we removed indels and poorly genotyped SNPs having VQSR tranche <99.0, genotype quality <20, inbreeding coefficient <-0.3 and >5% missingness, setting genotype calls with allelic depth <10 and allelic balance <0.2 or >0.8 as missing. WGS datasets were not imputed with HRC. Considering the small size of some of the datasets, we decided to focus further analysis on variants with MAF >1% and Hardy-Weinberg p-value >0.0001.

In each dataset, we removed genetically similar individuals by removing individuals with pihat >0.125, as calculated with PLINK 2.0^104^. Additionally, we merged genotypes with those from 1kgp, pruned genotypes with --indep-pairwise 50 5 0.2 in PLINK, and performed PCA on the sample correlation matrix. We performed k-nearest neighbors (k=7) on the first two PCs, using the known ancestry labels in 1kgp, to assign an ancestry to each genotyped sample. The majority of included samples was of EUR descent: 5,138 samples had an EUR assignment, 805 samples had an AFR assignment, and 573 samples were assigned to the other ethnicities (**Supplementary Table 1,** Figure 2B).

For the purpose of eQTL analysis, we next assessed links between RNA-seq and genotype samples and noted that some individuals had multiple RNA-seq samples (e.g. from multiple brain regions) or multiple genotype samples (e.g. from different genotyping platforms). In total, we were able to determine 7,644 links between RNA-seq samples and genotype samples (**Supplementary Table 1**), reflecting 3,525 unique EUR individuals, 624 unique AFR individuals and 510 unique individuals assigned to other ethnicities. We then grouped linked RNA-seq samples based on ethnicity and tissue group to prevent possible biases on eQTL results. For those individuals with multiple linked RNA-seq samples, we selected a sample at random within these groups. Within each tissue and ethnicity group, we then selected unique genotype samples across datasets in such a way to maximize sample size per genotype dataset. For the eQTL analysis per tissue, we only considered those datasets having more than 30 unique linked samples available, and for which at least two independent datasets were available. Using these criteria for sample and dataset selection, we were able to create 7 eQTL discovery datasets: Basal ganglia-EUR (n=208), Cerebellum-EUR (n=492), Cortex-EUR (n=2,970), Cortex-AFR (n=420), Hippocampus-EUR (n=168) and Spinal cord-EUR (n=108; **Supplementary Table 1,** Figure 2C).

### eQTL analysis

Our dataset consists of different tissues and ethnicities, and samples have been collected in different institutes using different protocols. Consequently, combining these datasets to perform eQTL analysis is complicated, due to possible biases each of these factors may introduce. To resolve this issue, we opted to perform an eQTL meta-analysis within each of the defined eQTL discovery datasets. To reduce the effect of possible gene expression outliers, we calculated Spearman’s rank correlation coefficients for each eQTL in each dataset separately, and then meta-analyzed the resulting coefficients using a sample size weighted Z-score method, as described previously^14^. While we acknowledge that this method may provide less statistical power than the commonly used linear regression, we chose this method to provide conservative effect estimates. To identify *cis*-eQTLs, we tested SNPs located within 1 Mb of the transcription start site, while for the identification of *trans*-eQTLs, we required this distance to be at least 5 Mb. For both analyses, we selected variants having a MAF>1%, and a Hardy-Weinberg p-value >0.0001. Using the GENCODE v32 annotation, we were able to quantify 58,243 genes, of which 19,373 are protein coding. While non-coding genes have been implicated to be important for brain function^105^, these genes generally have poor genomic and functional annotations, meaning that it is often unknown in which pathway they function, and that there is uncertainty about their genomic sequence. We therefore focused our eQTL analyses on protein coding genes.

To correct for multiple testing, we reperformed the *cis*- and *trans*-eQTL analyses, while permuting the sample labels 10 times. Using the permuted p-values, we created empirical null distributions and determined a false discovery rate (FDR) as the proportion of unpermuted observations over the permuted observations and considered associations with FDR<0.05 as significant. To provide a more stringent FDR estimate for our *cis*-eQTL results, we limited FDR estimation to the top associations per gene, as described previously^14^. We note that our FDR estimate is evaluated on a genome-wide level, rather than per gene, and consequently FDR estimates stabilize after a few permutations^106^.

Since *cis*-eQTL loci are known to often harbor multiple independent associations, we performed an iterative conditional analysis, where for each iteration, we regressed the top association per gene from the previous associations, and re-performed the *cis*-eQTL analysis until no additional associations at FDR<0.05 could be identified.

Since a genome-wide *trans*-eQTL analysis would result in a large multiple testing burden considering the billions of potential tests, we limited this analysis to a set of 130,968 variants with a known interpretation. This set constituted of variants that were either previously associated with traits, having a GWAS p-value <5×10^-8^ in the IEU OpenGWAS database^107^ and EBI GWAS catalog^108^ on May 3^rd^, 2020, and additional neurological traits (see **Supplementary Table 17**) or were showing an association with FDR<0.05 in any of our discovery *cis*-eQTL analyses (including secondary, tertiary and quandary associations identified in the iterative conditional analysis). *Cis*-eQTLs in Cortex-EUR were highly concordant when replicated in Cortex-AFR (Figure 3C). Consequently, to maximize the sample size and statistical power, we meta-analyzed Cortex-EUR and Cortex-AFR datasets together. However, for the *trans*-eQTL analysis we omitted ENA, to prevent bias by genotypes called from RNA-seq samples. Additionally, For the *trans*-eQTL analysis, we did not correct the gene expression data for 10 PCs, since *trans*-eQTLs can be driven by cell proportion differences^17^, and many of the first 10 PCs in the *MetaBrain* dataset were correlated with estimated cell type proportions (**Supplementary Figure 32**). To test for *trans*-eQTLs, we assessed those combinations of SNPs and genes where the SNP-TSS distance was >5 Mb, or where gene and SNP were on different chromosomes. We note that we did not evaluate eQTLs where the SNP-TSS distance was >1 Mb and <5 Mb, which potentially excludes detection of long-range *cis*-eQTLs or short-range *trans*-eQTLs. We expect however, that this excludes only a limited number of eQTLs, since we observed that this distance was <31Kb for 50% of *cis*-eQTLs (Figure 3B), indicating most *cis*-eQTLs are short-ranged. Additionally, we reasoned that the >5 Mb cutoff would prevent identification of false-positive *trans*-eQTLs due to long-range LD.

### Estimation of cell type proportions and identification of cell type mediated eQTLs

By leveraging cell type specific gene expression collected through scRNA-seq, a bulk tissue sample can be modelled as a parts-based representation of the distinct cell types it consists of. In such a model, the weights of each part (i.e. cell type proportions) can be determined by deconvolution. In the deconvolution of the *MetaBrain* bulk expression data we used a single-cell derived signature matrix including the five major cell types in the brain: neurons, oligodendrocytes, macrophages, endothelial cells and astrocytes. This signature matrix was generated in the context of the CellMap project (Zhengyu Ouyang *et al*.; manuscript in preparation). In short, we created pseudo-bulk expression profiles by extracting gene expression values for specific cell types of interest from annotated single cell and single nuclei expression matrices. Using differential expression analysis and applying several rounds of training and testing, we selected 1,166 differentially expressed genes and calculated the average read counts per cell type. We then filtered out genes that had no variation in expression, leaving a total of 1,132 genes. We extracted the corresponding TMM normalized gene counts of these signature genes for all European cortex samples in *MetaBrain*. After correcting the counts for cohort effects using OLS, but not for any other technical covariates, we applied log2 transformation on both the signature matrix as well as the bulk gene count matrix. Subsequently we applied non-negative least squares (NNLS)^109^ using SciPy (version 1.4.1)^110^ to model the bulk expression as a parts-based representation of the single-nucleus derived signature matrix. First introduced by Lawson and Hanson^109^, NNLS method is the basis of numerous deconvolution methods to date. In short, NNLS attempt to find a non-negative weight (coefficient) for each of the cell types that, when summed together, minimizes the least-squares distance to the observed gene counts. Lastly, we transformed the resulting coefficients into cell type proportions by dividing them over the sum of coefficients for each sample. The resulting cell proportions are then used to identify cell type mediated eQTL effects. For this we applied Decon-eQTL^22^ (version 1.4; default parameters) in order to systematically test for significant interaction between each cell type proportion and genotype, while also controlling for the effect on expression of the other cell types. The resulting p-values are then correct for multiple testing using the Benjamini-Hochberg method on a per-cell-type basis.

### Cell type specific ROSMAP single-nucleus datasets

In order further confirm cell type specific eQTL effects, we used the ROSMAP single-nucleus data, encompassing 80,660 single-nucleus transcriptomes from the prefrontal cortex of 48 individuals with varying degrees of Alzheimer’s disease pathology^111^. We used Seurat version 3.2.2^112^ to analyze the data. First, we removed the genes that did not pass filtering as described previously^111^ leaving us with 16,866 genes and 70,634 cells for further analysis. After this, we normalized the expression matrix on a per individual per cell type basis using sctransform^113^ and visualized the normalized expression matrix using UMAP dimensionality reduction^114^. We observed that cell types, as defined by Mathys *et al*^115^., for the majority cluster together (**Supplementary Figures 33 and 34**). We then created expression matrices for each broad cell type (excitatory neurons, oligodendrocytes, inhibitory neurons, astrocytes, oligodendrocyte precursor cells, microglia, pericytes and endothelial cells) by calculating the average expression per gene and per individual basis. We then used these cell-type datasets for eQTL mapping using the same procedure as the bulk data. To correct for multiple testing, we confined the analysis to only test for primary *cis*- and *trans*-eQTLs found in *MetaBrain* cortex, while also permuting the sample labels 100 times. Lastly, we calculated the Spearman correlation between gene expression levels and genotypes and their 95% confidence intervals^116^.

### Single SNP Mendelian Randomization analysis

Mendelian Randomization (MR) was conducted between the Cortex-EUR eQTLs and 31 neurological traits (21 neurological disease outcomes, 2 quantitative traits and 8 brain volume outcomes) (**Supplementary Table 11**). Cortex-EUR eQTLs at genome-wide significant (p<5×10^-8^) were selected and then LD clumped to obtain independent SNPs to form our set of instruments. LD clumping was carried out using the ld_clump() function in the ieugwasr package^117^ on the default settings (10,000 Kb clumping window with r^2^ cut-off of 0.001 using the 1000 Genomes EUR reference panel). SNP associations for each of the eQTL instruments were then looked up in the outcome GWASs of interest. If the SNP could not be found in the outcome GWAS using a direct lookup of the dbSNP rsid, then a proxy search was performed to extract the next closest SNP available in terms of pairwise LD, providing minimum r^2^ threshold of 0.8 with the instrument. Outcome GWAS lookup and proxy search was performed using the associations() function in the ieugwasr package. To ensure correct orientation of effect alleles between the eQTL instrument and outcome GWAS associations, the SNP effects were harmonized using the harmonise_data() function in TwoSampleMR^87^. Action 2 was selected which assumes that the alleles are forward stranded in the GWASs (i.e. no filtering or re-orientation of alleles according to frequency was conducted on the palindromic SNPs). Single SNP MR was then performed on the harmonized SNP summary statistics using the mr_singlesnp() function in TwoSampleMR. Single SNP MR step computes a Wald ratio, which estimates the change in risk for the outcome per unit change in gene expression, explained through the effect allele of the instrumenting SNP. We reported all the MR findings that passed a p-value threshold of 5×10^-5^, but note that the Bonferroni-corrected p=0.05 threshold for multiple testing correction is p=1.865×10^-7^. We did not implement multi-SNP analysis (such as the Inverse Variance Weighted method), because there are a small number of instrumenting SNPs available per gene, which could result in unreliable pooled MR estimates for genes.

### Colocalization

Following the MR analysis, colocalization analysis was performed on the MR findings that passed the suggestive threshold to determine if the eQTL and trait shared the same underlying signal. We ran colocalization^36^ using both the default parameters (p1=p2=10^-4^ and p12=10^-5^) and parameters based on the number of SNPs in the region (p1=p2=1/(number of SNPs in the region) and p12=p1/10). We considered the two traits, eQTL and GWAS outcome to colocalize if either of the two parameters yielded PP4>0.7. Additionally, colocalization was systematically analyzed against one trait to compare to robustness of the Cortex-EUR eQTLs with existing cortex eQTL data sets (see **Supplementary Note**).

## URLs

**Picard**: http://broadinstitute.github.io/picard/

**dbGAP**: https://dbgap.ncbi.nlm.nih.gov

**European Nucleotide Archive**: http://www.ebi.ac.uk/ena

**ieugwasr package**: https://mrcieu.github.io/ieugwasr/

**TwoSampleMR**: https://mrcieu.github.io/TwoSampleMR/

## Accessions

**TargetALS**^12^ TargetALS data was pushed directly from the NY Genome center to our sftp server.

**CMC**^118^ CMC data was downloaded from https://www.synapse.org/ using synapse client (https://python-docs.synapse.org/build/html/index.html). Accession code: syn2759792

**GTEx**^86^ GTEx was downloaded from SRA using fastq-dump of the SRA toolkit (http://www.ncbi.nlm.nih.gov/Traces/sra/sra.cgi?cmd=show&f=software&m=software&s=software). Access has been requested and granted through dbGaP.

**Braineac**^7^ Braineac data has been pushed to our ftp server by Biogen.

**AMP-AD**^5^ AMP-AD data has been downloaded from synapse^13^. Accession code: syn2580853. snRNA-seq was collected using Synapse accession code: syn18485175. IHC data: https://github.com/ellispatrick/CortexCellDeconv/tree/master/CellTypeDeconvAnalysis/Data

**ENA**^13^ ENA data has been downloaded from the European Nucleotide Archive. The identifiers of the 76 included studies and 2021 brain samples are listed in Supplementary Table 29.

**CMC_HBCC:** CMC_HBCC data was downloaded from https://www.synapse.org/ using synapse client (https://python-docs.synapse.org/build/html/index.html). Accession code: syn10623034

**BrainSeq** BrainSeq data was downloaded from https://www.synapse.org/ using synapse client (https://python-docs.synapse.org/build/html/index.html). Accession code: syn12299750

**UCLA_ASD** UCLA_ASD data was downloaded from https://www.synapse.org/ using synapse client (https://python-docs.synapse.org/build/html/index.html). Accession code: syn4587609

**BrainGVEx** BrainGVEx data was downloaded from https://www.synapse.org/ using synapse client (https://python-docs.synapse.org/build/html/index.html). Accession code: syn4590909

**BipSeq** BipSeq data was downloaded from https://www.synapse.org/ using synapse client (https://python-docs.synapse.org/build/html/index.html). Accession code: syn5844980

**GTEx** GTEx data was downloaded from dbgap. Accession code: phs000424.v7.p2

**NABEC** NABEC data was downloaded from dbgap. Accession code: phs001301.v1.p1

**CellMap** single-cell and single-nuclei RNA-seq datasets were downloaded from Gene Expression Omnibus (GEO), BioProject, the European Genome-phenome Archive (EGA) and the Allan Brain Atlas. Accession codes: GSE97930, GSE126836, GSE103723, GSE104276, PRJNA544731, PRJNA434002, phs000424, phs001836.

## Acknowledgements

We thank the donors of the brain tissues underlying the RNA-seq data used for this study and their families for their willingness to donate samples for research. We would like to thank the Center for Information Technology of the University of Groningen for their support and for providing access to the Peregrine high-performance computing cluster, as well as the UMCG Genomics Coordination center, the UG Center for Information Technology and their sponsors BBMRI-NL and TarGet for storage and compute infrastructure. We would like to greatly thank all researchers involved with the following projects for making their data available for use. E.A.T., Y.H., C.-Y.C, E.E.M., M.I.Z. and H.R. are employed by Biogen. D.B. and T.R.G. are supported by funding from the UK Medical Research Council (MRC Integrative Epidemiology Unit at the University of Bristol, MC_UU_00011/4) and a sponsored research collaboration with Biogen. L.F. is supported by grants from the Dutch Research Council (ZonMW-VIDI 917.14.374 to L.F.), by an ERC Starting Grant, grant agreement 637640 (ImmRisk), by an Oncode Senior Investigator grant and a sponsored research collaboration with Biogen. This project has received funding from the European Research Council (ERC) under the European Union’s Horizon 2020 research and innovation programme (grant agreement n° 772376 - EScORIAL. The authors thank the Biogen CellMap team (Z. Ouyang, N. Bourgeois, E. Lyashenko, P. Cundiff, K. Li, X. Zhang, F. Casey, S. Engle, R. Kleiman, B. Zhang and M. Zavodszky) for the expertise and advice provided toward deriving the cell-type specific expression profiles.

## ROSMAP

The results published here are in whole or in part based on data obtained from the AMP-AD Knowledge Portal (<u>doi:10.7303/syn2580853</u>) Study data were provided by the Rush Alzheimer’s Disease Center, Rush University Medical Center, Chicago. Data collection was supported through funding by NIA grants P30AG10161, R01AG15819, R01AG17917, R01AG30146, R01AG36836, U01AG32984, U01AG46152, the Illinois Department of Public Health, and the Translational Genomics Research Institute.

Genotype data: <u>doi:10.1038/mp.2017.20.</u> RNAseq: <u>doi:10.1038/s41593-018-0154-9</u>. snRNA-seq: doi:10.7303/syn18485175

## Mayo

The results published here are in whole or in part based on data obtained from the AMP-AD Knowledge Portal (<u>doi:10.7303/syn2580853</u>). Study data were provided by the following sources: The Mayo Clinic Alzheimer’s Disease Genetic Studies, led by Dr. Nilufer Taner and Dr. Steven G. Younkin, Mayo Clinic, Jacksonville, FL using samples from the Mayo Clinic Study of Aging, the Mayo Clinic Alzheimer’s Disease Research Center, and the Mayo Clinic Brain Bank. Data collection was supported through funding by NIA grants P50 AG016574, R01 AG032990, U01 AG046139, R01 AG018023, U01 AG006576, U01 AG006786, R01 AG025711, R01 AG017216, R01 AG003949, NINDS grant R01 NS080820, CurePSP Foundation, and support from Mayo Foundation. Study data includes samples collected through the Sun Health Research Institute Brain and Body Donation Program of Sun City, Arizona. The Brain and Body Donation Program is supported by the National Institute of Neurological Disorders and Stroke (U24 NS072026 National Brain and Tissue Resource for Parkinsons Disease and Related Disorders), the National Institute on Aging (P30 AG19610 Arizona Alzheimer’s Disease Core Center), the Arizona Department of Health Services (contract 211002, Arizona Alzheimer’s Research Center), the Arizona Biomedical Research Commission (contracts 4001, 0011, 05-901 and 1001 to the Arizona Parkinson’s Disease Consortium) and the Michael J. Fox Foundation for Parkinson’s Research. <u>doi:10.1038/sdata.2016.89</u>

## MSBB

The results published here are in whole or in part based on data obtained from the AMP-AD Knowledge Portal (<u>doi:10.7303/syn2580853</u>). These data were generated from postmortem brain tissue collected through the Mount Sinai VA Medical Center Brain Bank and were provided by Dr. Eric Schadt from Mount Sinai School of Medicine.

## CMC

Data were generated as part of the CommonMind Consortium supported by funding from Takeda Pharmaceuticals Company Limited, F. Hoffman-La Roche Ltd and NIH grants R01MH085542, R01MH093725, P50MH066392, P50MH080405, R01MH097276, RO1-MH-075916, P50M096891, P50MH084053S1, R37MH057881, AG02219, AG05138, MH06692, R01MH110921, R01MH109677, R01MH109897, U01MH103392, and contract HHSN271201300031C through IRP NIMH. Brain tissue for the study was obtained from the following brain bank collections: The Mount Sinai NIH Brain and Tissue Repository, the University of Pennsylvania Alzheimer’s Disease Core Center, the University of Pittsburgh NeuroBioBank and Brain and Tissue Repositories, and the NIMH Human Brain Collection Core. CMC Leadership: Panos Roussos, Joseph Buxbaum, Andrew Chess, Schahram Akbarian, Vahram Haroutunian (Icahn School of Medicine at Mount Sinai), Bernie Devlin, David Lewis (University of Pittsburgh), Raquel Gur, Chang-Gyu Hahn (University of Pennsylvania), Enrico Domenici (University of Trento), Mette A. Peters, Solveig Sieberts (Sage Bionetworks), Thomas Lehner, Stefano Marenco, Barbara K. Lipska (NIMH).

## GTEx

The Genotype-Tissue Expression (GTEx) Project was supported by the Common Fund of the Office of the Director of the National Institutes of Health (commonfund.nih.gov/GTEx). Additional funds were provided by the NCI, NHGRI, NHLBI, NIDA, NIMH, and NINDS. Donors were enrolled at Biospecimen Source Sites funded by NCI\Leidos Biomedical Research, Inc. subcontracts to the National Disease Research Interchange (10XS170), Roswell Park Cancer Institute (10XS171), and Science Care, Inc. (×10S172). The Laboratory, Data Analysis, and Coordinating Center (LDACC) was funded through a contract (HHSN268201000029C) to the The Broad Institute, Inc. Biorepository operations were funded through a Leidos Biomedical Research, Inc. subcontract to Van Andel Research Institute (10ST1035). Additional data repository and project management were provided by Leidos Biomedical Research, Inc.(HHSN261200800001E). The Brain Bank was supported supplements to University of Miami grant DA006227. Statistical Methods development grants were made to the University of Geneva (MH090941 & MH101814), the University of Chicago (MH090951, MH090937, MH101825, & MH101820), the University of North Carolina - Chapel Hill (MH090936), North Carolina State University (MH101819), Harvard University (MH090948), Stanford University (MH101782), Washington University (MH101810), and to the University of Pennsylvania (MH101822). The datasets used for the analyses described in this manuscript were obtained from dbGaP at http://www.ncbi.nlm.nih.gov/gap through dbGaP accession number phs000424.v7.p2 on 02/27/2020.

## NABEC

Data was collected from dbGAP accession phs001301.v1.p1, which was generated by J. R. Gibbs, M. van der Brug, D. Hernandez, B. Traynor, M. Nalls, S-L. Lai, S. Arepalli, A. Dillman, I. Rafferty, J. Troncoso, R. Johnson, H. R. Zielke, L. Ferrucci, D. Longo, M.R. Cookson, and A.B. Singleton. The NABEC dataset was generated at National Institute on Aging, Bethesda, MD, USA, Institute of Neurology, University College London, London, UK, The Scripps Research Institute, Jupiter, FL, USA, Johns Hopkins University, Baltimore, MD, USA, and the University of Maryland Medical School, Baltimore, MD, USA. NABEC was funded by Z01 AG000949-02. National Institutes of Health, Bethesda, MD, USA and Z01 AG000015-49. National Institutes of Health, Bethesda, MD, USA.

## TargetALS

This data set was generated and supported by the following: Target ALS Human Postmortem Tissue Core, New York Genome Center for Genomics of Neurodegenerative Disease, Amyotrophic Lateral Sclerosis Association and TOW Foundation.

## Braineac

Data was collected from doi.org/10.1038/s41467-020-14483-x, which was generated by Mina Ryten, David Zhang, and Karishma D’Sa, Sebastian Guelfi and Regina Reynolds. Mina Ryten, David Zhang, and Karishma D’Sa were supported by the UK Medical Research Council (MRC) through the award of Tenure-track Clinician Scientist Fellowship to Mina Ryten (MR/N008324/1). Sebastian Guelfi was supported by Alzheimer’s Research UK through the award of a PhD Fellowship (ARUK-PhD2014-16). Regina Reynolds was supported through the award of a Leonard Wolfson Doctoral Training Fellowship in Neurodegeneration. All RNA sequencing data performed as part of this study were generated by the commercial company AROS Applied Biotechnology A/S (Denmark).

We also would like to thank Guelfi *et al*.^119^ for the use of their data.

## European Nucleotide Archive

We would like to thank all donors and their families, principal investigators and their funding bodies for each of the projects included from the European Nucleotide Archive.

## UCLA ASD, Bipseq, BrainGVEx and LIBD

Data were generated as part of the PsychENCODE Consortium supported by: U01MH103339, U01MH103365, U01MH103392, U01MH103340, U01MH103346, R01MH105472, R01MH094714, R01MH105898, R21MH102791, R21MH105881, R21MH103877, and P50MH106934 awarded to: Schahram Akbarian (Icahn School of Medicine at Mount Sinai), Gregory Crawford (Duke), Stella Dracheva (Icahn School of Medicine at Mount Sinai), Peggy Farnham (USC), Mark Gerstein (Yale), Daniel Geschwind (UCLA), Thomas M. Hyde (LIBD), Andrew Jaffe (LIBD), James A. Knowles (USC), Chunyu Liu (UIC), Dalila Pinto (Icahn School of Medicine at Mount Sinai), Nenad Sestan (Yale), Pamela Sklar (Icahn School of Medicine at Mount Sinai), Matthew State (UCSF), Patrick Sullivan (UNC), Flora Vaccarino (Yale), Sherman Weissman (Yale), Kevin White (UChicago) and Peter Zandi (JHU).

## Author contributions

N.K., O.E.G, and H.W. processed the RNA-seq and genotype data. N.K. and H.W. were responsible for data management. N.K. and H.W. were responsible for the *cis-*eQTL analysis. H.W. was responsible for the *trans*-eQTL analysis. M.V., Z.O. and M.I.Z. were responsible for the cell type proportion prediction. N.K. and M.V. were responsible for the cell type interaction analysis. S.D. was responsible for the selection of brain samples from ENA. D.B., Y.H., C.-Y.C., E.E.M, T.R.G. and E.A.T. were responsible for MR and colocalization analysis and interpretation. P.D., O.B.B. and L.F. were responsible for the Downstreamer analysis. L.F., E.A.T. and H.R. acquired funding and supervised the study. N.K., E.A.T., M.V., D.B, Y.H., C.-Y.C., O.B.B., H.R., L.F. and H.W. drafted the manuscript. All authors have proof-read the manuscript.

## Supplementary Figure Legends

**Supplementary Figure 1. European Nucleotide Archive brain sample selection. (A)** Principal component (PC) analysis on the expression data of 74,052 samples included in the SkyMap database shows clustering on tissue type but also many outliers with high PC1 scores. **(B)** Coloring on single and paired-end sequencing shows no clear clustering. **(C)** Coloring single cell identifies the samples with high PC1 scores as single-cell samples. **(D)** Mean % reads mapped, number of reads, and max reads per bin of PC1. **(E)** Re-calculation of PCs on all samples with PC score <0 in panel A-D, after covariate correction. **(F)** Brain and Tissue score calculated by correlating expression of known tissue and brain samples to each of the PCs. **(G)** As panel F, cancer score was calculated by correlating expression of known cancer genes to all PCs.

**Supplementary Figure 2. RNA-seq alignment QC.** The two main RNA-seq QC metrics used for filtering samples. (**A**) Percentage coding bases colored by dataset and (**B**) percentage of reads aligned colored per dataset. Red dotted line is the threshold for filtering (10% for coding bases and 60% for percentage reads aligned respectively). Triangles are samples filtered out by any of the RNA-seq QC metrics.

**Supplementary Figure 3. Sample filtering by PCA.** Principal component analysis (PCA) plot before normalization and covariate removal. For all plots the red line indicates 4 standard deviations from the mean and red dots are samples to be filtered out. **(A)** PCA on all samples after removing alignment QC outliers. **(B)** PCA on samples after removal of outlier samples from A. **(C)** PCA on samples after removal of outlier samples of A and B.

**Supplementary Figure 4. PCA before and after covariate correction.** (**A**) PC1 and PC2 on normalized expression data before covariate correction, colored on dataset. (**B**) PC1 and PC2 on normalized expression data after covariate correction.

**Supplementary Figure 5. Assigning ethnicity through principal component analysis.** For each of the included datasets principal component (PC) scores are calculated on their genotypes. Samples are clustered with the 1000 genome samples (left). Right panels show dataset genotype samples without 1000g samples on the right projected on the same PCs. Using k-nearest neighbors clustering, samples are assigned an ethnicity based on their closeness to the 1000g samples of a population.

**Supplementary Figure 6. eQTL Z-score comparison between datasets.** The pairwise spearman correlation and concordance of direction of the eQTL Z-scores between all cohorts, and between each cohort and the meta-analysis Z-score. As two examples, (**A**) shows the Z-score comparison between Cortex-EUR eQTL datasets EUR-LIBD_h650 and EUR-UCLA_ASD, and (**B**) shows the Z-score comparison between the meta-analysis Z-score and the Cortex-EUR cohort EUR-AMPAD-ROSMAP-V2. (**C**) shows the correlation for each pairwise combination of cohorts between each other (small dots), and with the meta-analysis Z-scores (large dots). (**D**) shows the directional concordance for each pairwise combination of cohorts between each other (small dots), and with the meta-analysis Z-scores (large dots). The dots in (**C**) and (**D**) that correspond to the (**A**) and (**B**) plots are shown by the grey dottes lines.

**Supplementary Figure 7. Reads mapping on patch chromosome version of MAPT.** Number of reads mapped to the MAPT gene located on the primary assembly (ENSG00000186868) and the MAPT genes located on the patch chromosomes (ENSG00000276155 and ENSG00000277956). Each dot is an individual, and the color shows if they are homozygous reference (0/0), heterozygous (0/1), or homozygous alternative (1/1) for a SNP (rs34619181) located in the MAPT gene. Left plot compares counts mapped to ENSG00000186868 (ref) to those mapped to ENSG00000276155 (patch), middle plot compares ENSG00000186868 (ref) and ENSG00000277956 (patch), right plot compares ENSG00000276155 (patch) and ENSG00000277956 (patch).

**Supplementary Figure 8. EQTL z-scores in the MAPT locus.** Z-scores (y-axis) of the MAPT locus (x-axis) for all the datasets used in the Cortex-EUR meta-analysis. Left upper plot shows the meta-analysis Z-score. Blue dots are the SNPs that are in high LD with the top SNP.

**Supplementary Figure 9. Colocalization locus plot for MAPT**. Y-axis shows the colocalization log10(-p-value). X-axis shows the position of the SNPs (dots). Color is the LD with rs56240678.

**Supplementary Figure 10. (A)** Mean of log_2_ of the expression (x-axis) and standard deviation of the log_2_ of expression for primary, secondary, tertiary, and quaternary eQTL genes. eQTLs that have only one independent SNP effect have higher mean expression but lower standard deviation than genes with multiple independent effects. **(B)** g:profiler enrichment for all genes with a single independent eQTL effect. **(C)** g:profiler enrichment for all genes with multiple independent eQTL effects.

**Supplementary Figure 11. Properties of cerebellum specific eQTLs.** (**A**) UpSet plot of the number of eQTL genes per brain region for European datasets. (**B**) The distribution of log2(TMM+1) expression in cortex (x-axis) and cerebellum (y-axis) of the 846 eQTL genes that were only significant in cerebellum. Blue line is the minima of the bimodal distribution and is used as cut-off point in panel **C** (**C**) The expression in cortex (x-axis) and cerebellum (y-axis) of the 846 eQTL genes that were only significant eQTLs in cerebellum. The blue line is the cut-off from panel **B**. (**D**) The expression (dots) and standard deviation (lines) of the transcription factors that are enriched for binding to transcription sites around the 662 genes for cortex (x-axis) and cerebellum (y-axis). The 5 transcription factors that are labelled are lower expressed in cortex and higher expressed in cerebellum.

**Supplementary Figure 12. Cortex primary eQTL replication in GTEx.** The replication between primary *cis*-eQTLs of Cortex-EUR (discovery) with all the GTEx tissues (replication). The x-axis is the number of eQTLs that is significant in both discovery and replication, and the y-axis is the percentage that shows the same direction of effect.

**Supplementary Figure 13.** Comparison of meta-analysis Z-scores for eQTLs detected in the different *MetaBrain* datasets (x-axis), and eQTLgen (y-axis).

**Supplementary Figure 14. Distribution of predicted cell proportions.** The distribution of the predicted cell proportions (x-axis) for cortex and cerebellum samples (y-axis).

**Supplementary Figure 15. Cell type proportions per brain region are comparable, with the exception of the spinal cord.** Visualization of the cell type proportions with one row per cell type and colors indicating brain region. (**A**) Density plot where the x-axis shows the predicted cell type proportion, and the y-axis shows the frequency. (**B**) Boxplot of the predicted cell type proportion. Boxes represent the 25^th^ and 75^th^ percentiles and internal line represents the median. The whiskers represent 1.5 multiplied by the inter-quartile range. Outliers are shown as individual points.

**Supplementary Figure 16. Cell type fractions per brain tissue shows little differences with the exception of the spinal cord.** Visualization of the cell type proportions with one row per brain region and colors indicating cell types. (**A**) Density plot where the x-axis shows the predicted cell type proportion, and the y-axis shows the frequency. (**B**) Boxplot of the predicted cell type proportion. Boxes represent the 25^th^ and 75^th^ percentiles and internal line represents the median. The whiskers represent 1.5 multiplied by the inter-quartile range. Outliers are shown as individual points.

**Supplementary Figure 17. Cell type mediated eQTLs in cerebellum are mostly mediated by astrocytes and macrophages.** The number of cell type interacting eQTLs for cerebellum deconvoluted cell types. We did not identify eQTLs that were shared between cell types.

**Supplementary Figure 18. Replication of cortex *cis*-eQTLs in snRNA-seq data.** Each figure in this plot represents a comparison between bulk RNA-seq (y-axis) and single-nucleus RNA-seq (x-axis). Each dot represents one *cis*-eQTL, and the legend shows the Pearson correlation coefficient. Each column is a comparison between equivalent (and where not possible; similar) cell types in both datasets. Each row illustrates a different filtering on which eQTLs are shown and/or a different value on the y-axis. The x-axis always denotes the overall z-score of the eQTL effect in the single nucleus dataset of that respective column. (**A**) Meta-analysis eQTL z-score (y-axis) in Cortex-EUR bulk RNA-seq data, no filtering is applied. (**B**) Meta-analysis eQTL z-score (y-axis) in Cortex-EUR bulk data, eQTLs are filtered based on the Decon-QTL Benjamini-Hochberg corrected p-value <0.05 in each respective column. (**C**) same as row **B** but now showing the log betas of the interaction model on the y-axis. (**D**) Meta-analysis eQTL z-score (y-axis) in bulk data for eQTLs that are significantly replicating in each respective dataset. Dots are colored if they are significantly cell type mediated (BH FDR<0.05) by the respective cell type in bulk data. (**E)** y-axis shows the log betas of the interaction model (y-axis) and filtering eQTLs on both significantly replicating in each respective dataset, as well as being significantly cell type mediated in bulk data.

**Supplementary Figure 19. Bulk interacting eQTLs replicating in single-nucleus ROSMAP.** Replication of cell type interaction eQTLs for *STMN4* **(A)**, *FAM221A* **(B)**, *NKAIN1* **(C)** and *SCL25A27* **(D)**. First column: Boxplots of the eQTL effect in Cortex-EUR bulk RNA-seq. Second column: Cell type interacting eQTL effect in Cortex-EUR bulk RNA-seq. The x-axis shows the estimated cell type proportion, the y-axis shows the gene expression, each dot represents a sample, and the colors indicate the SNP genotype, with yellow being the minor allele. Values under the alleles are Spearman correlation coefficients. Third column: Forest plot of the spearman coefficient with effect direction relative to the minor allele when replicating the eQTL effect in ROSMAP single nucleus data (n=38). Error bars indicate 95% confidence interval. Each row denotes a cell type specific dataset: excitatory neurons (EX), oligodendrocytes (OLI), inhibitory neurons (IN), astrocytes (AST), oligodendrocyte precursor cells (OPC), microglia (MIC), pericytes (PER) and endothelial cells (END). The bold cell type corresponds to the cell type that showed an interaction effect in bulk RNA-seq. Fourth column: Cell type interacting eQTL effect in ROSMAP single-nucleus RNA-seq (n=38) of the bold highlighted cell type in the third colum.

**Supplementary Figure 20. Mendelian Randomization summary.** Each plot is for a different trait (Intelligence, Intracranial volume, Putamen volume, Years of schooling, Alzheimer’s disease, Amyotrophic Lateral Sclerosis, Depression (broad), Frontotemporal Dementia, Parkinson’s disease, Bipolar disorder, Generalized epilepsy, juvenile myoclonic epilepsy, multiple sclerosis and schizophrenia). For each SNP the effect allele (EA) is given, the eQTL beta of the EA on the given gene, the odds ratio (disease traits) or beta (quantitative traits) of the EA on the phenotype, and the Wald ratio p-value of the mendelian randomization analysis.

**Supplementary Figure 21**. Colocalization regional plots for five suggestive MR findings in Cortex-EUR that were replicated in eQTLGen with allelic discordance. Regional plots were made for five MR findings (*CASS4* for Alzheimer’s disease, *TMEM170B* for intelligence, *GATAD2A* for schizophrenia and years of schooling, and *ZCWPW1* for years of schooling) in Cortex-EUR (top), eQTLGen (middle) and outcome GWAS (bottom) to show colocalization. These five findings all passed suggestive threshold (p<5×10^-5^) in Cortex-EUR, with eQTL effects replicated in eQTLGen (p<0.05), showed colocalization for both Cortex-EUR and eQTLGen but opposite directions of effect.

**Supplementary Figure 22**. Colocalization regional plots for two suggestive MR findings for multiple sclerosis that showed opposite directions of effect between Cortex-EUR and eQTLGen. Regional plots were made for two suggestive MR findings for MS (*KMT5A*, *RNF19B*), both of which were suggestive signals in Cortex-EUR as well as eQTLGen (p<5×10^-5^). Opposite directions of effect were observed between Cortex-EUR and eQTLGen but colocalization was only found in Cortex-EUR.

**Supplementary Figure 23**. Scatterplots comparing MR effects for multiple sclerosis derived using instruments from the metabrain versus eQTLGen studies. The top panel shows the WR comparison on the same gene but with the different SNP instruments selected by each study (matching on the top WR finding if gene instrumented with multiple SNPs in the study) and the bottom panel the WR comparison between *MetaBrain* instruments and eQTLGen matching on both the same gene and SNP instrument. Genes which showed opposite direction of WR effect between *MetaBrain* and eQTLGen are colored in red and the genes with the same direction in blue.

**Supplementary Figure 24.** Log10 of median **e**xpression of brain and blood tissue samples in GTEx for 28 multiple sclerosis genes for which there are no significant eQTLgen instruments in brain and blood.

**Supplementary Figure 25. Cell type proportions in Alzheimer’s disease patients.** Predicted cell count proportions for the AMP-AD samples that were used in the Cortex-EUR eQTL analysis for individuals with Alzheimer’s disease and non-neurological controls. Each dot is the predicted cell proportion for one sample. Numbers under the boxplots indicate the number of samples plotted. Values above the line are p-values from a t-test between groups.

**Supplementary Figure 26. Forest plots for rs1990622 *trans*-eQTLs.** Forest plots for each of the *trans*-eQTL genes associated with rs1990622. Each plot shows the *trans*-eQTL beta and 95% confidence interval for each of the included datasets and the meta-analysis. Effect directions are relative to the A allele of rs1990622. Sizes of dots are relative to sample size of each dataset. *Trans*-eQTL effects are most pronounced in AMP-AD datasets.

**Supplementary Figure 27. Summary of 7p21.3 locus trans-eQTLs. (A)** Forest plots showing effect sizes for rs1990622 (yellow; beta and 95% confidence interval) for *cis*-eQTL gene *THSD7A*, *trans*-eQTL gene *CALB2*, and association of rs1990622 with estimated neuron proportion. Right panel shows average estimated neuron proportions per dataset (blue violin plots). EQTL and neuron proportion associations are most pronounced in AMP-AD datasets, while average neuron proportions are comparable. **(B)** *Trans*-eQTL meta-analysis Z-scores for rs11974335, rs10950398 and rs1990622 (x-axis), and the correlation of those *trans*-eQTL genes with predicted neuron proportion (y-axis) are highly correlated**. (C)** Comparison of *trans*-eQTL Z-scores between Alzheimer’s disease patients (x-axis) and neurotypical controls (y-axis) shows that eQTL Z-scores are higher in patients.

**Supplementary Figure 28. Replication of cortex *trans*-eQTLs in single-nucleus data.** Each figure in this plot represents a comparison between bulk RNA-seq (y-axis) and single-nucleus RNA-seq (x-axis). Each dot represents one *trans*-eQTL, and the legend shows the Pearson correlation coefficient. Each column is a comparison between equivalent (and where not possible; similar) cell types in both datasets. Each row illustrates a different filtering on which eQTLs are shown and/or a different value on the y-axis. The x-axis always denotes the overall z-score of the eQTL effect in the single nucleus dataset of that respective column. (**A**) Meta-analysis eQTL z-score (y-axis) in Cortex-EUR bulk RNA-seq data, no filtering is applied. (**B**) Meta-analysis eQTL z-score (y-axis) in Cortex-EUR bulk data, eQTLs are filtered based on the Decon-QTL Benjamini-Hochberg corrected p-value <0.05 in each respective column. (**C**) same as row **B** but now showing the log betas of the interaction model on the y-axis. (**D**) Meta-analysis eQTL z-score (y-axis) in bulk data for eQTLs that are significantly replicating in each respective dataset. Dots are colored if they are significantly cell type mediated (BH FDR<0.05) by the respective cell type in bulk data. (**E)** y-axis shows the log betas of the interaction model (y-axis) and filtering eQTLs on both significantly replicating in each respective dataset, as well as being significantly cell type mediated in bulk data.

**Supplementary figure 29. Comparison of AUC distribution for different eigenvector cut-offs.** The quality of the gene network that we built for *MetaBrain* is measured by an AUC for each gene derived from a leave-one-out procedure. One of the parameters to build the network is the number of eigenvectors to use after PCA over the gene correlation matrix. Here we show for the 6 annotation categories (KEGG, REACTOME, GO Biological Process, GO Molecular Function, GO Cellular Component, and HPO) the AUC mean (dot) and standard deviation (lines) at different eigenvector cut-offs. The red dot and line indicate the eigenvector cut-off that was used for that annotation category.

**Supplementary Figure 30. Heatmaps of the Pearson correlation of the AUC values between different eigenvector cut-offs.** Correlation was calculated between the different eigenvector cutoffs for the 6 annotation categories.

**Supplementary Figure 31. (A)** UMAP representation of heterogeneous gene network. Immune and blood cell types show increased gene expression levels for genes prioritized using *Downstreamer* for multiple sclerosis, while decreased expression is observed in brain related tissues. **(B)** Within MetaBrain, those same genes show lower expression in cortex, but higher expression in spinal cord and cerebellum.

**Supplementary Figure 32. Spearman correlation heatmap of predicted cell fractions versus principal components calculated using all *MetaBrain* samples.** A heatmap showing the first fifty principal components as the columns and the five cell types for which we predicted proportions as rows. Each cell is colored based on the spearman correlation coefficients. Blue denotes a negative correlation, red a positive correlation and white denotes no correlation.

**Supplementary Figure 33. SnRNA-seq visualization by cell type.** UMAP dimensionality reduction plot of 39 snRNA-seq samples from ROSMAP. Each dot represents a single cell (n=70,634). The dots are colored by their corresponding cell type: excitatory neurons (EX), oligodendrocytes (OLI), inhibitory neurons (IN), astrocytes (AST), oligodendrocyte precursor cells (OPC), microglia (MIC), pericytes (PER) and endothelial cells (END).

**Supplementary Figure 34. SnRNA-seq visualization by cell type.** UMAP dimensionality reduction plot of 39 snRNA-seq samples from ROSMAP. Each dot represents a single cell (n=70,634). The dots are colored by their corresponding cell type subcluster: excitatory neurons (EX), oligodendrocytes (OLI), inhibitory neurons (IN), astrocytes (AST), oligodendrocyte precursor cells (OPC), microglia (MIC), pericytes (PER) and endothelial cells (END).

## Supplementary Table descriptions

**Supplementary table 1. Number of samples and individuals.**

**Sheet Genotype QC:** The number of genotype individuals and samples pre-QC (**column C-H**) and post-QC (column I-N) for the different RNA-seq (**column A**) and genotype (**column B**) datasets. Columns are: **PreQC**: Number of initial genotype samples processed for QC. **PostQC**: Number of genotype samples left after QC filtering. **RNA-seq dataset**: Name of the complete dataset. **Genotype dataset**: Name of the genotype dataset. Some datasets have multiple genotype platforms, or multiple smaller datasets that are part of the larger RNA-seq dataset. **Individuals:** The number of individuals per dataset. **EUR**: Number of genotype samples per dataset of individuals of European population. **AFR**: Number of genotype samples per dataset of individuals of African population. **EAS**: Number of genotype samples per dataset of individuals of East-Asian population. **SAS**: Number of genotype samples per dataset of individuals of South-Asian population. **AMR:** Number of genotype samples per dataset of individuals of Ad Mixed American population.

Sheet **RNA-QC**: The number of RNA-seq samples at different steps of QC and for different brain regions. Cells A2-F18 have the number of samples at different QC steps. Columns are: **Dataset**: dataset name. **Number of RNA-seq samples**: Number of RNA-seq samples processed to go through QC. **Alignment QC**: Number of RNA-seq samples left after filtering on alignment QC (e.g. percent reads aligned). **RNA-seq PCA outliers - step 1**: Number of RNA-seq samples left after filtering samples >4SD from mean of PC1. **RNA-seq PCA outliers - step 2**: Number of samples left after recalculating PCA and again removing samples >4SD fom mean of PC1.

**Covariate removal:** Number of samples left after covariate removal. **RNA Tissue grouping:** the meta-data across different datasets uses different granularity of tissue annotation. Tissues were grouped accordingly.

**Sheet Sample Links**: RNA-seq samples linked to genotype samples. Left top: numbers of RNA-seq sample linked to a genotype sample per dataset, per population. Top right: number of unique individuals per dataset per population. Middle: number of uniquely linked individuals per dataset, per population and per tissue group. Bottom: numbers of individuals used from each dataset and population for *cis*- and *trans*-eQTL analysis.

**Supplementary table 2. C*is*-eQTL summary statistics.**

*Cis*-eQTL summary statistics listing index variant per gene (FDR<0.05). One sheet per eQTL discovery dataset. Genomic positions are GRCh38. eQTL Rank: whether the eQTL is a primary, secondary, tertiary, quaternary, or higher eQTL.

**Supplementary table 3. Number of *cis-* and *trans-*eQTLs.** For each dataset the number of *cis-* and *trans*-eQTL SNPs, genes, and SNP-gene combinations found at FDR<0.05. Columns are: **Basalganglia, Cerebellum, Cortex, Hippocampus, Spinalcord**: the five different brain regions for which eQTL calling was done. **EUR:** Number of eQTLs with samples from European population. **AFR**: Number of eQTLs with samples from African population. **EAS**: Number of eQTLs with samples from East-Asian population. **EUR+AFR, wo ENA, no PCA**: Number of eQTLs with samples from EUR and AFR populations, excluding samples from the ENA cohorts, and using gene expression levels that were not corrected for principal components.

**Supplementary table 4. Gene set enrichment summary statistics for primary and higher rank eQTLs.** Gene set enrichment summary statistics generated using g:Profiler for genes having a primary eQTL effect (sheet Primary eQTL), and those also having a secondary eQTL (sheet Non-primary eQTL).

**Supplementary table 5.** Gene set enrichment summary statistics generated using g:Profiler for genes having an eQTL effect in cerebellum.

**Supplementary table 6. GTEx *cis-*eQTL replication.** Replication between *cis*-eQTLs of different *MetaBrain* regions and all GTEx tissues. Discovery was performed in each *MetaBrain* dataset while excluding GTEx, and then replicated in each GTEx tissue. **Tested eQTLs:** those eQTLs that were also present in the GTEx dataset. **Proportion shared and FDR<0.05**: proportion of tested eQTLs that was also significant in GTEx. **Concordant and FDR<0.05**: number of tested eQTLs that was also significant and for which the allelic direction was concordant. **Concordance**: proportion of concordant tested and significant eQTLs.

**Supplementary table 7. eQTLgen *cis*-eQTL replication.** *MetaBrain cis-*eQTLs (FDR<0.05) as discovery cohort and eQTLgen eQTLs as replication cohort. Top table: FDR<0.05 in MetaBrain discovery only (FDR<1 in eQTLGen). Bottom table: FDR<0.05 in both MetaBrain and eQTLgen datasets. **Shared**: number of shared eQTLs. **Concordant**: number of shared eQTLs that has the same allelic direction of effect. **Concordant over total**: proportion of concordant eQTLs over the total number of eQTLs discovered. **Concordant over shared:** proportion of concordant eQTLs over number of shared eQTLs.

**Supplementary table 8. Cell type deconvolution summary statistics. Sheet cortex:** All Decon-eQTL results for cortex. **Sheet cerebellum:** All Decon-eQTL results for cerebellum. Columns for both sheets are: **Gene**: deconvoluted eQTL gene ensebl ID. **Gene symbol**: deconvolution eQTL gene symbol. **SNP**: deconvoluted eQTL SNP. **Alleles**: SNP alleles. **Effect Allele**: the allele to which the betas are directed. **Columns ending with p-value**: p-value for the cell-type interaction. **Columns ending with beta**: beta for the cell-type proportion term. **Columns ending with beta:GT**: beta for the genotype x cell-type interaction term.

**Supplementary table 9 Replication of the MetaBrain cortex primary *cis*-ieQTLs in ROSMAP single-nucleus data.** For each of the deconvoluted cell-types, the FDR and betas are listed. For each of the cell types in the single nucleus data, the FDR and eQTL Z-scores are listed. All betas andZ-scores are relative to the Effect Allele.

**Supplementary table 10. eQTL SNPs in linkage disequilibrium with GWAS SNPs.** The GWAS SNPs that are in high linkage disequilibrium (LD) with the *cis*-eQTL SNPs. Each sheet is a different *metabrain* eQTL datasets from EUR populations. The sheet Included Traits lists GWAS traits that were tested. Columns are: **eQTL rank**: the rank of conditional eQTLs (1=primary, 2=secondary, etc). **GWASID**: GWAS ID of the GWAS SNP. **Trait**: Name of the GWAS trait. **Index variant**: the GWAS variant. **Index Variant P**: GWAS p-value. **Index Variant Alleles**: Alleles of the GWAS variant. **Index Variant Effect**: GWAS effect. **Linked EQTL SNP**: the eQTL SNP. LD(rsq): the LD r^2^. **LinkedEQTLGenes**: the eQTL genes that the linked SNP affects. **Linked EQTL Gene Symbols**: HGNC name of the linked genes. **Linked EQTL Alleles**: Alleles of the eQTL SNP. **Linked EQTL Effect Allele**: The allele that is related to the effect direction. **Linked EQTL Zscores**: Z-scores of the eQTL effect. **Linked EQTL P**: p-value of the eQTL effect. **GWAS Cluster Size**: Number of GWAS SNPs in LD with Index Variant. SNPs In **GWAS Cluster**: SNPs that are in LD with the Index Variant.

**Supplementary table 11. List of traits used in Mendelian randomization and colocalization analysis.**

**Supplementary table 12 eQTL SNPs which showed evidence of genetic colocalization with tested brain-related traits. ID, Chromosome, Position, SNP, Effect Allele, Non Effect Allele:** Position of instrumenting SNP with effect allele used during the harmonization procedure. **Proxy used, Proxy SNP:** whether proxy lookup had to be performed to find SNP in outcome GWAS and the rsid of the proxy used. **MetaBrain SNP effects:** gene name and summary statistics for the instrument-exposure SNP association (*MetaBrain* eQTL). **Outcome SNP effects:** outcome name (neurological trait) and summary statistics for the harmonized instrument-outcome SNP association. **MR effects:** single SNP Wald ratio effect between the instrumented eQTL and neurological outcome. **Coloc results:** colocalization probability of both traits sharing the same causal variant in the region. **Decon-QTL results: eQTL SNP:** the SNP that was tested for cell type mediated effects. In some cases a SNP which is in high LD with the instrument SNP is used for Decon-QTL. **LD R-squared:** the LD between SNP and eQTL SNP. Columns listing Decon-QTL results: **beta:** the beta of the interaction term in the Decon-QTL model with respect to the Effect Allele column. **FDR:** the Benjamini-Hochberg corrected interaction p-value. **Mendelian Disorders:** overlap of genes with Development Disorder Genotype - Phenotype Database (DDG2P) and OrphaNet.

**Supplementary Table 13. Colocalization results for latest AD GWAS loci with *MetaBrain* Cortex-EUR primary eQTLs (columns A to P** were adapted from Schwartzentruber *et al*. for comparisons and **columns Q to Y** are *MetaBrain* findings. **Category - 1:** previously identified and replicated in *MetaBrain* Cortex-EUR, **2:** novel results found by *MetaBrain* Cortex-EUR, **3:** previously identified but not replicated in *MetaBrain* Cortex-EUR.

**Supplementary table 14. Mendelian Randomization comparison between MetaBrain and eQTLGen on multiple sclerosis outcome.** (a) Wald Ratio comparison on the same gene using different SNP instruments. For this analysis, the Wald Ratio effects for the top hit eQTL for each gene within each study were compared. (b) Wald Ratio comparison on the same gene fixing on the same eQTL instrument between studies. For this analysis, the eQTLGen Wald Ratios were re-derived using the second Taylor expansion error term on the same SNP instruments as MetaBrain.

**Supplementary table 15. Colocalization of MR suggestive hits with high LD but allelic discordance. This** table displays the colocalization results for 31 suggestive MR findings from Cortex-EUR with eQTL instruments replicated in eQTLGen (p<0.05) but allelic discordance (opposite directionalities of alleles). Highlighted rows are findings with colocalization in both Cortex-EUR and eQTLGen.

**Supplementary table 16. Comparison of MR suggestive hits for MS between metaBrain and eQTLGen.** This table displays 157 suggestive MR signals for multiple sclerosis in Cortex-EUR and the replication MR and colocalization results of corresponding genes in eQTLGen.

**Supplementary table 17. *Trans*-eQTL summary statistics.** Sheet Trans-eQTLs: all trans-eQTLs detected in this study (FDR<0.05). **Percentage cross-mapping:** percentage of the gene that can be mapped within 5Mb of the *trans*-eQTL SNP. Sheet *Trans*-eQTLs no crossmap: *trans*-eQTLs that remain significant after cross-mapping eQTLs have been removed. Sheet *Trans*-eQTLs with cis per trait: in this sheet, *trans*-eQTLs are annotated with *cis*-eQTLs for the same SNP, and subsequently split per trait annotation for the SNP. Consequently, a single trans-eQTL may be represented by multiple rows. Sheet Convergent *trans*-eQTLs: genes on which multiple independent loci have a *trans*-eQTL, split per annotated trait. Sheet TraitsAndNrOfSNPs: list of traits included in the analysis, and the number of included SNPs per trait.

**Supplementary table 18. Summary statistics for associations between SNPs and predicted cell-type proportions.** Sheet Cortex-EUR: associations (FDR<0.05) while limiting to Cortex-EUR samples. Sheet Cortex-EUR+AFR-woENA: associations (FDR<0.05) for the analysis including AFR samples, but excluding ENA samples.

**Supplementary table 19. Differences in predicted neuron proportions between included datasets.** T-test p-values comparing neuron proportions for pairwise comparisons between the datasets included in the *trans*-eQTL analysis.

**Supplementary table 20. Gene-cell count correlations and 7p21.3 *trans*-eQTL Z-scores.** *Trans*-eQTL Z-scores for three SNPs (rs11974335, rs10950398, and rs1990622), and correlations of the *trans*-eQTL genes with predicted neuron proportions.

**Supplementary table 21. Gene set enrichments for 7p21.3 *trans*-eQTL genes.** Gene set enrichments calculated using g:Profiler. Sheet downregulated genes: gene set enrichments for genes that show downregulation due to the 7p21.3 trans-eQTL effect alleles. Sheet upregulated genes: gene set enrichments for genes that show upregulation due to the 7p21.3 trans-eQTL effect alleles.

**Supplementary table 22. Replication of the MetaBrain cortex primary *trans*-ieQTLs in ROSMAP single-nucleus data.** For each of the deconvoluted cell-types, the FDR and betas are listed. For each of the cell types in the single nucleus data, the FDR and eQTL Z-scores are listed. All betas and Z-scores are relative to the Effect Allele.

**Supplementary table 23. Downstreamer results for amyotrophic lateral sclerosis in EUR and Asian populations.** Sheet overview: lists set of ontologies tested for this phenotype. Sheet GenePrioritization_MetaBrain: gene prioritization performed in all *MetaBrain* samples. Sheet GenePrioritization_MetaBrainCortexOnly: gene prioritization performed in *MetaBrain* cortex samples. GenePrioritization_MetaBrainCerebellumOnly: gene prioritization performed in *MetaBrain* cerebellum samples. Sheets Reactome_MetaBrain, GO_BP_MetaBrain, GO_CC_MetaBrain, GO_MF_MetaBrain, KEGG_MetaBrain, and HPO_MetaBrain: gene set enrichments for coregulated genes identified using Downstreamer. Sheets Expression_MetaBrain, Expression_HCA, and GtexV8_relative: expression enrichment using all MetaBrain samples, Human Cell Atlas, and GTEx v8.

**Supplementary table 24. Downstreamer results for Parkinson’s disease.** Sheet overview: lists set of ontologies tested for this phenotype. Sheet GenePrioritization_MetaBrain: gene prioritization performed in all *MetaBrain* samples. Sheet GenePrioritization_MetaBrainCortexOnly: gene prioritization performed in *MetaBrain* cortex samples. GenePrioritization_MetaBrainCerebellumOnly: gene prioritization performed in *MetaBrain* cerebellum samples. Sheets Reactome_MetaBrain, GO_BP_MetaBrain, GO_CC_MetaBrain, GO_MF_MetaBrain, KEGG_MetaBrain, and HPO_MetaBrain: gene set enrichments for coregulated genes identified using Downstreamer. Sheets Expression_MetaBrain, Expression_HCA, and GtexV8_relative: expression enrichment using all MetaBrain samples, Human Cell Atlas, and GTEx v8.

**Supplementary table 25. Downstreamer results for schizophrenia.** Sheet overview: lists set of ontologies tested for this phenotype. Sheet GenePrioritization_MetaBrain: gene prioritization performed in all *MetaBrain* samples. Sheet GenePrioritization_MetaBrainCortexOnly: gene prioritization performed in *MetaBrain* cortex samples.

GenePrioritization_MetaBrainCerebellumOnly: gene prioritization performed in *MetaBrain* cerebellum samples. Sheets Reactome_MetaBrain, GO_BP_MetaBrain, GO_CC_MetaBrain, GO_MF_MetaBrain, KEGG_MetaBrain, and HPO_MetaBrain: gene set enrichments for coregulated genes identified using Downstreamer. Sheets Expression_MetaBrain, Expression_HCA, and GtexV8_relative: expression enrichment using all MetaBrain samples, Human Cell Atlas, and GTEx v8.

**Supplementary table 26. Downstreamer results for Alzheimer’s disease.** Sheet overview: lists set of ontologies tested for this phenotype. Sheet GenePrioritization_MetaBrain: gene prioritization performed in all *MetaBrain* samples. Sheet GenePrioritization_MetaBrainCortexOnly: gene prioritization performed in *MetaBrain* cortex samples. GenePrioritization_MetaBrainCerebellumOnly: gene prioritization performed in *MetaBrain* cerebellum samples. Sheets Reactome_MetaBrain, GO_BP_MetaBrain, GO_CC_MetaBrain, GO_MF_MetaBrain, KEGG_MetaBrain, and HPO_MetaBrain: gene set enrichments for coregulated genes identified using Downstreamer. Sheets Expression_MetaBrain, Expression_HCA, and GtexV8_relative: expression enrichment using all MetaBrain samples, Human Cell Atlas, and GTEx v8.

**Supplementary table 27. Downstreamer results for multiple sclerosis.** Sheet overview: lists set of ontologies tested for this phenotype. Sheet GenePrioritization_MetaBrain: gene prioritization performed in all *MetaBrain* samples. Sheet GenePrioritization_MetaBrainCortexOnly: gene prioritization performed in *MetaBrain* cortex samples. GenePrioritization_MetaBrainCerebellumOnly: gene prioritization performed in *MetaBrain* cerebellum samples. Sheets Reactome_MetaBrain, GO_BP_MetaBrain, GO_CC_MetaBrain, GO_MF_MetaBrain, KEGG_MetaBrain, and HPO_MetaBrain: gene set enrichments for coregulated genes identified using Downstreamer. Sheets Expression_MetaBrain, Expression_HCA, and GtexV8_relative: expression enrichment using all MetaBrain samples, Human Cell Atlas, and GTEx v8.

**Supplementary table 28. Downstreamer results for amyotrophic lateral sclerosis in EUR population.** Sheet overview: lists set of ontologies tested for this phenotype. Sheet GenePrioritization_MetaBrain: gene prioritization performed in all *MetaBrain* samples. Sheet GenePrioritization_MetaBrainCortexOnly: gene prioritization performed in *MetaBrain* cortex samples. GenePrioritization_MetaBrainCerebellumOnly: gene prioritization performed in *MetaBrain* cerebellum samples. Sheets Reactome_MetaBrain, GO_BP_MetaBrain, GO_CC_MetaBrain, GO_MF_MetaBrain, KEGG_MetaBrain, and HPO_MetaBrain: gene set enrichments for coregulated genes identified using Downstreamer. Sheets Expression_MetaBrain, Expression_HCA, and GtexV8_relative: expression enrichment using all MetaBrain samples, Human Cell Atlas, and GTEx v8.

**Supplementary table 29. ENA accession IDs.** List of study accession IDs collected from European Nucleotide Archive. Columns are: **study_accession:** ID of the study in ENA. **run_accession:** ID of all the ENA runs included in this study (before quality control)

## References

1. Vos, T. et al. Global burden of 369 diseases and injuries in 204 countries and territories, 1990–2019: a systematic analysis for the Global Burden of Disease Study 2019. The Lancet 396, 1204–1222 (2020).

2. World Alzheimer Report 2018 - The state of the art of dementia research: New frontiers. NEW Front. 48.

3. Donovan, M. K. R., D’Antonio-Chronowska, A., D’Antonio, M. & Frazer, K. A. Cellular deconvolution of GTEx tissues powers discovery of disease and cell-type associated regulatory variants. Nat. Commun. 11, 955 (2020).

4. Wang, D. et al. Comprehensive functional genomic resource and integrative model for the human brain. Science 362, (2018).

5. Raj, T. et al. Integrative transcriptome analyses of the aging brain implicate altered splicing in Alzheimer’s disease susceptibility. Nat. Genet. 50, 1584–1592 (2018).

6. Hodes, R. J. & Buckholtz, N. Accelerating Medicines Partnership: Alzheimer’s Disease (AMP-AD) Knowledge Portal Aids Alzheimer’s Drug Discovery through Open Data Sharing. Expert Opin. Ther. Targets 20, 389–391 (2016).

7. Ramasamy, A. et al. Genetic variability in the regulation of gene expression in ten regions of the human brain. Nat. Neurosci. 17, 1418–1428 (2014).

8. Consortium*, T. P. Revealing the brain’s molecular architecture. Science 362, 1262–1263 (2018).

9. Fromer, M. et al. Gene expression elucidates functional impact of polygenic risk for schizophrenia. Nat. Neurosci. 19, 1442–1453 (2016).

10. BrainSeq: A Human Brain Genomics Consortium. BrainSeq: Neurogenomics to Drive Novel Target Discovery for Neuropsychiatric Disorders. Neuron 88, 1078–1083 (2015).

11. Gibbs, J. R. et al. Abundant Quantitative Trait Loci Exist for DNA Methylation and Gene Expression in Human Brain. PLOS Genet. 6, e1000952 (2010).

12. Prudencio, M. et al. Distinct brain transcriptome profiles in C9orf72-associated and sporadic ALS. Nat. Neurosci. 18, 1175–1182 (2015).

13. Leinonen, R. et al. The European Nucleotide Archive. Nucleic Acids Res. 39, D28–D31 (2011).

14. Võsa, U. et al. Unraveling the polygenic architecture of complex traits using blood eQTL metaanalysis. bioRxiv 447367 (2018) doi:10.1101/447367.

15. Dobbyn, A. et al. Landscape of Conditional eQTL in Dorsolateral Prefrontal Cortex and Co-localization with Schizophrenia GWAS. Am. J. Hum. Genet. 102, 1169–1184 (2018).

16. Wingender, E., Dietze, P., Karas, H. & Knüppel, R. TRANSFAC: a database on transcription factors and their DNA binding sites. Nucleic Acids Res. 24, 238–241 (1996).

17. Võsa, U. et al. Unraveling the polygenic architecture of complex traits using blood eQTL metaanalysis. bioRxiv 447367 (2018) doi:10.1101/447367.

18. Foley, C. N. et al. A fast and efficient colocalization algorithm for identifying shared genetic risk factors across multiple traits. Nat. Commun. 12, 764 (2021).

19. Fu, J. et al. Unraveling the Regulatory Mechanisms Underlying Tissue-Dependent Genetic Variation of Gene Expression. PLOS Genet. 8, e1002431 (2012).

20. Fu, J. et al. Unraveling the Regulatory Mechanisms Underlying Tissue-Dependent Genetic Variation of Gene Expression. PLOS Genet. 8, e1002431 (2012).

21. Glastonbury, C. A., Couto Alves, A., El-Sayed Moustafa, J. S. & Small, K. S. Cell-Type Heterogeneity in Adipose Tissue Is Associated with Complex Traits and Reveals Disease-Relevant Cell-Specific eQTLs. Am. J. Hum. Genet. 104, 1013–1024 (2019).

22. Raúl Aguirre-Gamboa et al. Deconvolution of bulk blood eQTL effects into immune cell subpopulations. BMC Bioinformatics 21, 243 (2020).

23. Zhengyu, O., et al. CellMap: Characterizing the type and composition of iPSC-derived cell lines from bulk RNA-seq data. Manuscr. Prep.

24. Bahney, J. & Von Bartheld, C. S. The Cellular Composition and Glia-Neuron Ratio in the Spinal Cord of a Human and a Non-Human Primate: Comparison with other Species and Brain Regions. Anat. Rec. Hoboken NJ 2007 301, 697–710 (2018).

25. Patrick, E. et al. Deconvolving the contributions of cell-type heterogeneity on cortical gene expression. PLOS Comput. Biol. 16, e1008120 (2020).

26. Herculano-Houzel, S. The human brain in numbers: a linearly scaled-up primate brain. Front. Hum. Neurosci. 3, (2009).

27. von Bartheld, C. S., Bahney, J. & Herculano-Houzel, S. The Search for True Numbers of Neurons and Glial Cells in the Human Brain: A Review of 150 Years of Cell Counting. J. Comp. Neurol. 524, 3865–3895 (2016).

28. Mathys, H. et al. Single-cell transcriptomic analysis of Alzheimer’s disease. Nature 570, 332–337 (2019).

29. Ng, B. et al. Using Transcriptomic Hidden Variables to Infer Context-Specific Genotype Effects in the Brain. Am. J. Hum. Genet. 105, 562–572 (2019).

30. Zhao, B. et al. Large-scale GWAS reveals genetic architecture of brain white matter microstructure and genetic overlap with cognitive and mental health traits (n = 17,706). Mol. Psychiatry 1–13 (2019) doi:10.1038/s41380-019-0569-z.

31. Smorodchenko, A. et al. Comparative analysis of uncoupling protein 4 distribution in various tissues under physiological conditions and during development. Biochim. Biophys. Acta BBA - Biomembr. 1788, 2309–2319 (2009).

32. Liu, D. et al. Mitochondrial UCP4 mediates an adaptive shift in energy metabolism and increases the resistance of neurons to metabolic and oxidative stress. Neuromolecular Med. 8, 389–414 (2006).

33. Yasuno, K. et al. Synergistic association of mitochondrial uncoupling protein (UCP) genes with schizophrenia. Am. J. Med. Genet. Part B Neuropsychiatr. Genet. Off. Publ. Int. Soc. Psychiatr. Genet. 144B, 250–253 (2007).

34. Ho, P. W. et al. Mitochondrial neuronal uncoupling proteins: a target for potential disease-modification in Parkinson’s disease. Transl. Neurodegener. 1, 3 (2012).

35. Ramsden, D. B. et al. Human neuronal uncoupling proteins 4 and 5 (UCP4 and UCP5): structural properties, regulation, and physiological role in protection against oxidative stress and mitochondrial dysfunction. Brain Behav. 2, 468–478 (2012).

36. Giambartolomei, C. et al. Bayesian test for colocalisation between pairs of genetic association studies using summary statistics. PLoS Genet. 10, e1004383 (2014).

37. Consortium*†, I. M. S. G. Multiple sclerosis genomic map implicates peripheral immune cells and microglia in susceptibility. Science 365, (2019).

38. International Multiple Sclerosis Genetics, C. et al. Genetic risk and a primary role for cell-mediated immune mechanisms in multiple sclerosis. Nature 476, 214–9 (2011).

39. Consortium*†, I. M. S. G. Multiple sclerosis genomic map implicates peripheral immune cells and microglia in susceptibility. Science 365, (2019).

40. Jones, G., Prosser, D. E. & Kaufmann, M. 25-Hydroxyvitamin D-24-hydroxylase (CYP24A1): its important role in the degradation of vitamin D. Arch Biochem Biophys 523, 9–18 (2012).

41. Schlingmann, K. P. et al. Mutations in CYP24A1 and idiopathic infantile hypercalcemia. N Engl J Med 365, 410–21 (2011).

42. Cappellani, D. et al. Hereditary Hypercalcemia Caused by a Homozygous Pathogenic Variant in the CYP24A1 Gene: A Case Report and Review of the Literature. Case Rep Endocrinol 2019, 4982621 (2019).

43. Mpandzou, G., Aït Ben Haddou, E., Regragui, W., Benomar, A. & Yahyaoui, M. Vitamin D deficiency and its role in neurological conditions: A review. Rev. Neurol. (Paris) 172, 109–122 (2016).

44. Agnello, L. et al. CYP27A1, CYP24A1, and RXR-alpha Polymorphisms, Vitamin D, and Multiple Sclerosis: a Pilot Study. J Mol Neurosci 66, 77–84 (2018).

45. Pierrot-Deseilligny, C. & Souberbielle, J. C. Is hypovitaminosis D one of the environmental risk factors for multiple sclerosis? Brain 133, 1869–88 (2010).

46. Rhead, B. et al. Mendelian randomization shows a causal effect of low vitamin D on multiple sclerosis risk. Neurol. Genet. 2, e97 (2016).

47. Jacobs, B. M., Noyce, A. J., Giovannoni, G. & Dobson, R. BMI and low vitamin D are causal factors for multiple sclerosis: A Mendelian Randomization study. Neurol. Neuroimmunol. Neuroinflammation 7, (2020).

48. Jiang, X., Ge, T. & Chen, C.-Y. The causal role of circulating vitamin D concentrations in human complex traits and diseases: a large-scale Mendelian randomization study. Sci. Rep. 11, 184 (2021).

49. Ramasamy, A. et al. Genetic evidence for a pathogenic role for the vitamin D3 metabolizing enzyme CYP24A1 in multiple sclerosis. Mult. Scler. Relat. Disord. 3, 211–219 (2014).

50. van Luijn, M. M. et al. Multiple sclerosis-associated CLEC16A controls HLA class II expression via late endosome biogenesis. Brain J. Neurol. 138, 1531–1547 (2015).

51. Ferrari, R. et al. Frontotemporal dementia and its subtypes: a genome-wide association study. Lancet Neurol 13, 686–99 (2014).

52. Wray, N. R. et al. Genome-wide association analyses identify 44 risk variants and refine the genetic architecture of major depression. Nat. Genet. 50, 668–681 (2018).

53. Li, Z. et al. Genetic variants associated with Alzheimer’s disease confer different cerebral cortex cell-type population structure. Genome Med. 10, 43 (2018).

54. Park, Y. et al. Single-cell deconvolution of 3,000 post-mortem brain samples for eQTL and GWAS dissection in mental disorders. bioRxiv 2021.01.21.426000 (2021) doi:10.1101/2021.01.21.426000.

55. Li, Z. et al. The TMEM106B FTLD-protective variant, rs1990621, is also associated with increased neuronal proportion. Acta Neuropathol. (Berl.) 139, 45–61 (2020).

56. Raudvere, U. et al. g:Profiler: a web server for functional enrichment analysis and conversions of gene lists (2019 update). Nucleic Acids Res. 47, W191–W198 (2019).

57. Sankaran, V. G. et al. Human Fetal Hemoglobin Expression Is Regulated by the Developmental Stage-Specific Repressor BCL11A. Science 322, 1839–1842 (2008).

58. Jiang, J. et al. cMYB is involved in the regulation of fetal hemoglobin production in adults. Blood 108, 1077–1083 (2006).

59. Métais, J.-Y. et al. Genome editing of HBG1 and HBG2 to induce fetal hemoglobin. Blood Adv. 3, 3379–3392 (2019).

60. J, D. & A, B. Dalfampridine: a brief review of its mechanism of action and efficacy as a treatment to improve walking in patients with multiple sclerosis. Curr. Med. Res. Opin. 27, 1415–1423 (2011).

61. Al-Owais, M. M. et al. Multiple mechanisms mediating carbon monoxide inhibition of the voltage-gated K + channel Kv1.5. Cell Death Dis. 8, e3163–e3163 (2017).

62. Rus, H. et al. The voltage-gated potassium channel Kv1.3 is highly expressed on inflammatory infiltrates in multiple sclerosis brain. Proc. Natl. Acad. Sci. 102, 11094–11099 (2005).

63. Noordam, R. et al. Multi-ancestry sleep-by-SNP interaction analysis in 126,926 individuals reveals lipid loci stratified by sleep duration. Nat. Commun. 10, 5121 (2019).

64. Gallois, A. et al. A comprehensive study of metabolite genetics reveals strong pleiotropy and heterogeneity across time and context. Nat. Commun. 10, 4788 (2019).

65. Sun, B. B. et al. Genomic atlas of the human plasma proteome. Nature 558, 73–79 (2018).

66. Suhre, K. et al. Connecting genetic risk to disease end points through the human blood plasma proteome. Nat. Commun. 8, 14357 (2017).

67. Tin, A. et al. Target genes, variants, tissues and transcriptional pathways influencing human serum urate levels. Nat. Genet. 51, 1459–1474 (2019).

68. Kichaev, G. et al. Leveraging Polygenic Functional Enrichment to Improve GWAS Power. Am. J. Hum. Genet. 104, 65–75 (2019).

69. Pers, T. H. et al. Biological interpretation of genome-wide association studies using predicted gene functions. Nat. Commun. 6, 5890 (2015).

70. Deelen, P. et al. Improving the diagnostic yield of exome-sequencing by predicting gene–phenotype associations using large-scale gene expression analysis. Nat. Commun. 10, 1–13 (2019).

71. Ripke, S. et al. Biological insights from 108 schizophrenia-associated genetic loci. Nature 511, 421–427 (2014).

72. Nalls, M. A. et al. Identification of novel risk loci, causal insights, and heritable risk for Parkinson’s disease: a meta-analysis of genome-wide association studies. Lancet Neurol 18, 1091–1102 (2019).

73. Kunkle, B. W. et al. Genetic meta-analysis of diagnosed Alzheimer’s disease identifies new risk loci and implicates Aβ, tau, immunity and lipid processing. Nat. Genet. 51, 414–430 (2019).

74. Conroy, J. et al. A novel locus for episodic ataxia:UBR4 the likely candidate. Eur. J. Hum. Genet. EJHG 22, 505–510 (2014).

75. Leal, S. S. & Gomes, C. M. Calcium dysregulation links ALS defective proteins and motor neuron selective vulnerability. Front. Cell. Neurosci. 9, 225 (2015).

76. Kalinowska-Lyszczarz, A. & Losy, J. The Role of Neurotrophins in Multiple Sclerosis— Pathological and Clinical Implications. Int. J. Mol. Sci. 13, 13713–13725 (2012).

77. Kerschensteiner, M. et al. Activated human T cells, B cells, and monocytes produce brain-derived neurotrophic factor in vitro and in inflammatory brain lesions: a neuroprotective role of inflammation? J. Exp. Med. 189, 865–870 (1999).

78. De Santi, L. et al. Neuroinflammation and neuroprotection: an update on (future) neurotrophin-related strategies in multiple sclerosis treatment. Curr. Med. Chem. 18, 1775–1784 (2011).

79. Baecher-Allan, C., Kaskow, B. J. & Weiner, H. L. Multiple Sclerosis: Mechanisms and Immunotherapy. Neuron 97, 742–768 (2018).

80. Redondo, J. et al. Purkinje Cell Pathology and Loss in Multiple Sclerosis Cerebellum. Brain Pathol. Zurich Switz. 25, 692–700 (2015).

81. Wang, X. & Goldstein, D. B. Enhancer Domains Predict Gene Pathogenicity and Inform Gene Discovery in Complex Disease. Am. J. Hum. Genet. 106, 215–233 (2020).

82. Wijst, M. G. P. van der et al. Single-cell RNA sequencing identifies celltype-specific cis-eQTLs and co-expression QTLs. Nat. Genet. 50, 493–497 (2018).

83. Population-scale single-cell RNA-seq profiling across dopaminergic neuron differentiation | bioRxiv. https://www.biorxiv.org/content/10.1101/2020.05.21.103820v1.

84. Consortium, S. W. G. of the P. G., Ripke, S., Walters, J. T. & O’Donovan, M. C. Mapping genomic loci prioritises genes and implicates synaptic biology in schizophrenia. medRxiv 2020.09.12.20192922 (2020) doi:10.1101/2020.09.12.20192922.

85. Davenport, E. E. et al. Discovering in vivo cytokine-eQTL interactions from a lupus clinical trial. Genome Biol. 19, 168 (2018).

86. Genetic effects on gene expression across human tissues. Nature 550, 204–213 (2017).

87. Hemani, G. et al. The MR-Base platform supports systematic causal inference across the human phenome. eLife 7, (2018).

88. Hemani, G., Bowden, J. & Davey Smith, G. Evaluating the potential role of pleiotropy in Mendelian randomization studies. Hum. Mol. Genet. 27, R195–R208 (2018).

89. Eliciting priors and relaxing the single causal variant assumption in colocalisation analyses. https://journals.plos.org/plosgenetics/article?id=10.1371/journal.pgen.1008720.

90. Team, T. S. E. synapseclient: A client for Synapse, a collaborative compute space that allows scientists to share and analyze data together.

91. Tsui, B., Dow, M., Skola, D. & Carter, H. Extracting allelic read counts from 250,000 human sequencing runs in Sequence Read Archive. bioRxiv 386441 (2018) doi:10.1101/386441.

92. Swertz, M. A. et al. The MOLGENIS toolkit: rapid prototyping of biosoftware at the push of a button. BMC Bioinformatics 11, S12 (2010).

93. Frankish, A. et al. GENCODE reference annotation for the human and mouse genomes. Nucleic Acids Res. 47, D766–D773 (2019).

94. Dobin, A. et al. STAR: ultrafast universal RNA-seq aligner. Bioinformatics 29, 15–21 (2013).

95. Anders, S., Pyl, P. T. & Huber, W. HTSeq—a Python framework to work with high-throughput sequencing data. Bioinformatics 31, 166–169 (2015).

96. Robinson, M. D. & Oshlack, A. A scaling normalization method for differential expression analysis of RNA-seq data. Genome Biol. 11, R25 (2010).

97. Robinson, M. D., McCarthy, D. J. & Smyth, G. K. edgeR: a Bioconductor package for differential expression analysis of digital gene expression data. Bioinformatics 26, 139–140 (2010).

98. R Core Team. R: A language and environment for statistical computing. (R Foundation for Statistical Computing, 2017).

99. Babraham Bioinformatics - FastQC A Quality Control tool for High Throughput Sequence Data. https://www.bioinformatics.babraham.ac.uk/projects/fastqc/.

100. Broad Institute. Picard Tools. (2019).

101. McCarthy, S. et al. A reference panel of 64,976 haplotypes for genotype imputation. Nat. Genet. 48, 1279–1283 (2016).

102. Deelen, P. et al. Genotype harmonizer: automatic strand alignment and format conversion for genotype data integration. BMC Res. Notes 7, 901 (2014).

103. Das, S. et al. Next-generation genotype imputation service and methods. Nat. Genet. 48, 1284–1287 (2016).

104. Chang, C. C. et al. Second-generation PLINK: rising to the challenge of larger and richer datasets. GigaScience 4, (2015).

105. Roberts, T. C., Morris, K. V. & Wood, M. J. A. The role of long non-coding RNAs in neurodevelopment, brain function and neurological disease. Philos. Trans. R. Soc. B Biol. Sci. 369, (2014).

106. Westra, H.-J. et al. Systematic identification of trans-eQTLs as putative drivers of known disease associations. Nat. Genet. 45, 1238–1243 (2013).

107. Lyon, M. et al. The variant call format provides efficient and robust storage of GWAS summary statistics. bioRxiv 2020.05.29.115824 (2020) doi:10.1101/2020.05.29.115824.

108. Buniello, A. et al. The NHGRI-EBI GWAS Catalog of published genome-wide association studies, targeted arrays and summary statistics 2019. Nucleic Acids Res. 47, D1005– D1012 (2019).

109. Lawson, C. L. & Hanson, R. J. Solving Least Squares Problems. (Society for Industrial and Applied Mathematics, 1995). doi:10.1137/1.9781611971217.

110. Virtanen, P. et al. SciPy 1.0: fundamental algorithms for scientific computing in Python. Nat. Methods 17, 261–272 (2020).

111. Mathys, H. et al. Single-cell transcriptomic analysis of Alzheimer’s disease. Nature 570, 332–337 (2019).

112. Stuart, T. et al. Comprehensive Integration of Single-Cell Data. Cell 177, 1888–1902.e21 (2019).

113. Hafemeister, C. & Satija, R. Normalization and variance stabilization of single-cell RNA-seq data using regularized negative binomial regression. Genome Biol. 20, 296 (2019).

114. McInnes, L., Healy, J. & Melville, J. UMAP: Uniform Manifold Approximation and Projection for Dimension Reduction. ArXiv180203426 Cs Stat (2020).

115. Mathys, H. et al. Single-cell transcriptomic analysis of Alzheimer’s disease. Nature 570, 332–337 (2019).

116. Bonett, D. G. & Wright, T. A. Sample size requirements for estimating pearson, kendall and spearman correlations. Psychometrika 65, 23–28 (2000).

117. Elsworth, B. et al. The MRC IEU OpenGWAS data infrastructure. bioRxiv 2020.08.10.244293 (2020) doi:10.1101/2020.08.10.244293.

118. Ng, B. et al. An xQTL map integrates the genetic architecture of the human brain’s transcriptome and epigenome. Nat. Neurosci. 20, 1418–1426 (2017).

119. Guelfi, S. et al. Regulatory sites for splicing in human basal ganglia are enriched for disease-relevant information. Nat. Commun. 11, 1041 (2020).

