## Supplementary Note for "Brain expression quantitative trait locus and network analysis reveals downstream effects and putative drivers for brain-related diseases"

### Selecting brain RNA-seq samples from European Nucleotide Archive

The European Nucleotide Archive (ENA) contains many bulk RNA-seq samples from many different tissues. We aimed to select the majority of brain related samples from this database. To this end, we made use of the SkyMap database, which consists of read counts for each sample in ENA, quantified with Kallisto v0.44.0 to GRCh build 38. We accessed this database through Synapse (<https://www.synapse.org/#!Synapse:syn16556230>) and selected 74,052 samples where at least 70% of the reads were mapped for 34,739 genes. We log2 transformed and quantile normalized the read counts and performed principal component analysis on the sample covariance matrix in order to detect outlier samples, and observed clustering of tissues on PC2, no clear clustering of single end vs paired end reads, and clear clustering of single cell versus bulk RNA-seq on PC1 (**Supplementary Figure 1A**, **B and C**). We set a threshold of 0 on PC1 to select 60,163 bulk RNA-seq samples. We noticed a number of samples with perfectly correlated read counts. These are likely the same samples that have been uploaded to the public domain multiple times and, as a result, have been included multiple times in SkyMap^1^. We removed 1,244 perfectly correlated (duplicate) samples to remain with 58,919 samples.

In a previous analysis, we found that the following technical covariates all significantly correlated to our sample PC scores for multiple PCs (p-value<0.01): read length, paired/single end, total reads in the dataset and percentage mapping read. This indicates that these technical factors affect the co-expression detected in the dataset, if not removed. Consequently, we next removed these covariates using linear regression, but decided not to correct for GC content per gene as this may also have biological meaning.

After the covariate correction, we recalculated the principal components, this time over the gene correlation matrix, and calculated PC scores for each sample for all PCs (**Supplementary Figure 1D**). We used the PC eigencoefficients to perform coregulation analysis using, limiting the analysis to 664 PCs having a Cronbach’s alpha ≥ 0.7 (which is an often-used threshold to select reliable PCs). The resulting PC showed clear clustering of tissues. Next, we determined which samples were tissues and which were cell lines by using a prediction algorithm: for each PC the PC scores per sample we correlated the binary vector describing if a sample is a tissue or not, resulting in a ‘tissue representation’ score indicative of how well that PC captures the signal differentiating tissues from non-tissue samples. Next, we correlated these ‘tissue representation’ scores of each PC to the PC scores for a sample for each PC. The resulting score indicates how likely this sample is a tissue (higher is more likely). In a similar manner we calculated “Brain scores” (**Supplementary Figure 1E**). We used a cutoff of >2 for the resulting tissue score, selecting 2,446 samples. However, we observed a large cluster of 449 samples indicated as cell lines, which were from a single study including ERCC Spike in controls. As such we decided to remove these samples, leaving 1,997 samples. We finally investigated whether these samples included cancer samples, by calculating a ‘cancer score’, and observed 292 samples which could be indicated as brain cancer samples. To maximize our sample size, we opted to include those samples (**Supplementary Figure 1F**). The selected samples were then downloaded from ENA using their SRA identifiers.

### Genotyping and imputation of ENA samples

Since we intended to include the ENA samples also in the eQTL analysis, yet genotype data was not available for these samples, we called genotypes using the RNA-seq reads. In a previous publication, we have shown that this can be performed with relatively high accuracy, suitable for eQTL analysis^2^.

Prior to genotype calling, we first aligned the samples using STAR^15^ (version 2.6.1c) with GENCODE^13^ v32 primary assembly as a reference, while omitting patch sequences from the reference. During this process, some samples failed to align, leaving 1,761 samples for genotype calling. After converting aligned SAM files to sorted BAM files, we used several GATK^3^ (version 4.0.8.1) steps for genotyping. In order we used MarkDuplicates to flag duplicated alignments, SplitNCigarReads to split reads into exon segments and clip reads overhanging introns, IndelRealignment to locally realign reads to minimize mismatching bases, BQSR to detect systematic errors made by the sequencer when it estimated the quality score of each base call, and HaplotypeCallerGvcf for estimating most likely genotypes and allele frequencies. Per interval in storage.googleapis.com/gatk-test-data/intervals/hg38.even.handcurated.20k.chr*.intervals (where chr* is chr1, chr2, etc), we created a genomic database using GATK’s GenomicsDBImport. Finally, we performed joined genotype calling per interval, using GATK’s GenotypeGVCF. Using this procedure, we called a total of 28,798,830 variants.

Since genes have varying read depths, genotype calls for many of the variants will have high missingness. We therefore first filtered out variants with >50% missingness using vcftools^4^ v0.1.14, selecting calls for biallelic SNPs with minor allele frequency (MAF) >0.1%, a minimal read depth (DP) of 10, and a minimal genotype quality (GQ) of 20. After this filter, 160,175 SNPs remained. Next, we evaluated missingness per sample, using the plink v1.9 --missing option and detected 315 samples with >50% missing genotype calls. We then repeated the vcftools filter step while omitting these 315 samples, leaving 210,498 SNPs. To identify the ethnicity of the included samples, we next performed Principal Component Analysis (PCA) on the genotypes, after merging overlapping SNPs with 1000 genomes phase 3 v5a, and pruning with plink v1.9 --indep-pairwise 50 5 0.2. We evaluated the first two Principal Components (PCs; **Supplementary Figure 5**) and assigned sample ethnicity labels by performing k-nearest neighbors (k=7) on these PCS. We observed a large cluster of 388 samples to the left of the samples labeled EUR. Upon closer inspection, these samples were from SRA project accession PRJNA208369, which is a study in the reliability of RNA-seq and includes spike in RNA sequences. We therefore removed those samples, leaving 1,060 samples for further processing.

The resulting genotypes still contain many missing values, while the joint genotyping approach of GATK provides genotype probabilities for many of the missing calls. We therefore replaced the missing calls by calculating genotyping likelihoods using the Beagle v4.1.11mar19.69c gtgl command, using the plink GRCh38 genome map as a reference genetic map. Then, we prepared the dataset for imputation by first lifting over the genotypes using GATK v4.1.4.1 LiftOverVCF using UCSC’s hg38ToHg19 liftover chain file, and the hg19 genome build as a reference. Finally, we matched the allele encoding of the lifted over genotypes using the HRC.r1-1.GRCh37.wgs.mac5.sites.vcf.gz sites list and used the Michigan imputation server to impute using HRC v1.1 as a reference.

### MAPT and the influence of patch sequences in GRCh38 on RNA-seq quantification

The eQTL association in the MAPT region is somewhat controversial: the xqtl browser^5^ (<http://mostafavilab.stat.ubc.ca/xqtl/>), which is constructed using ROSMAP and CMC studies, finds very strong effects on the MAPT locus. The MAPT eQTL has also been found in the newer and much larger PsychENCODE study (n~1,400; <http://resource.psychencode.org/>), but curiously cannot be found in the newest GTEx release (<https://www.gtexportal.org/>) nor in the latest meta-analysis from Sieberts *et al*.^6^, where ROSMAP is included as well (summary statistics are found here: <https://www.synapse.org/#!Synapse:syn16984815>).

One of the reasons this can happen is because the MAPT region is represented by multiple contigs/patch/alt sequences. Each of these sequences contains a full copy of the MAPT gene (i.e., ENSG00000276155 and ENSG00000277956), which confuses alignment and gene quantification software when these patch sequences are kept in the genome reference (they were excluded in the latest GTEx release). As a consequence, read counts for these alternate gene copies correlate very strongly with genotype, introducing an eQTL (**Supplementary Figure 7**).

Genetic association at the MAPT locus is inconsistent^7^. Even after re-aligning to the primary genome to exclude the patch chromosomes, there is a lot of heterogeneity in signal left between datasets. The haplotypic signal is present after meta-analysis, but mostly comes from CMC and ENA, with the effect in other datasets being much lower (**Supplementary Figure 8**). Additionally, co-localization at the MAPT locus is difficult due to the MAPT haplotypes with many SNPs in high LD in the region (**Supplementary Figure 9**). Genetic association at the MAPT locus is inconsistent^7^. Even after re-aligning to the primary genome to exclude the patch chromosomes, there is a lot of heterogeneity in signal left between datasets. The haplotypic signal is present after meta-analysis, but mostly comes from CMC and ENA, with the effect in other datasets being much lower (**Supplementary Figure 8**). Additionally, co-localization at the MAPT locus is difficult due to the MAPT haplotypes with many SNPs in high LD in the region (**Supplementary Figure 9**).

### Properties of primary, secondary, tertiary and quaternary *cis*-eQTLs

In blood, we have shown recently genes with the highest expression are most often *cis*-eQTL genes, cis-eQTL SNPs are generally located within 50Kb of the transcription start site (TSS) and genes without a *cis*-eQTL have low tolerance for loss of function mutations^8^. In contrast, after we ranked genes by average expression in Cortex-EUR, we observed that genes within the second decile are most likely cis-eQTLs, rather than those in the 10th decile. Nevertheless, as in blood, we observed that the median expression level of primary eQTL genes (median log_2_(TMM) expression=2.511) was slightly, but significantly, higher than that of genes without a *cis*-eQTL (median expression=2.505, Wilcoxon p-value: 9.96x10^-12^; **Figure 3B**). Comparing genes having only a primary eQTL association (median expression=2.596) with those having a secondary eQTL, we observed that genes with a secondary eQTL generally had a lower median expression level (median log_2_(TMM) expression=2.388; Wilcoxon p-value=1.3x10^-25^). Genes with additional independent eQTL associations showed a further decrease in median expression level (**Figure 3B**).

When evaluating the distance to TSS, we observed that the distance between the eQTL variant and the transcription start site (TSS) shifts further away with additional independent eQTL associations: primary eQTLs had a median distance of 31kb, secondary eQTLs a median distance of 42kb, tertiary eQTLs a median distance of 53Kb, and quaternary eQTLs a median distance of 76kb (**Figure 3B, middle**). This shift in distance between eQTL variant and TSS was smaller than that of previously published results in cortex^9^, potentially due to the larger sample size of *MetaBrain* resulting in an increase in power to detect secondary associations. Like observed previously in blood, genes with low tolerance for loss of function mutations, as indicated by the pLI score, were less likely to be eQTL genes^8^: primary eQTL having lower pLI scores than those without (*χ ^2^* p=6.35x10^-147^). This effect was smaller but still significant for secondary (*χ ^2^* p= 4.4x10^-43^) and tertiary eQTLs (*χ ^2^* p=3.7x10^-06^), but not for quaternary eQTLs (*χ^2^* p>0.05, **Figure 3B, right**).

Functional enrichment showed little difference between the types of eQTLs, with both primary and non-primary genes being enriched for autosomal recessive disorders (HPO). However, primary genes did have strong enrichments for protein binding (GO molecular function), intracellular and cytoplasm (GO cellular component) (**Supplementary Figure 10C**)**,** while non-primary genes were enriched for catalytic activity (GO molecular function). Finally, primary eQTL genes had many more TRANSFAC^10^ transcription factor motifs enriched. See **Supplementary Table 4** for a full list of significant enrichments.

### Binned expression of eQTL genes

It has been shown that in blood. with a large sample size, 95% of highly expressed genes have at least one independent eQTL effect, and that highly expressed genes that are not an eQTL have low tolerance to loss of function variants^11^. To test if this is also true for genes that are highly expressed in cortex, we calculated the mean and standard deviation over the log10(TMM normalized expression+1) table (before quantile normalization and covariate correction) for all samples that were included in the Cortex-EUR eQTL dataset. We then subset the genes to only those that were tested for being an eQTL at each iteration (For primary eQTLs: only protein coding genes, for secondary eQTLs: only significant primary eQTL genes, for tertiary eQTLs: only significant secondary eQTL genes, for quaternary eQTLs: only significant tertiary eQTL genes). For each iteration, we divided the genes up in 10 bins, ranking the genes from low to high expression. For each bin, we calculated the proportion of genes that had a significant eQTL, and observed that the 3^rd^ decile had the most (**Supplementary Figure 10A**). To test if this was due to the fact that the genes in the bins with highly expressed genes had a lower standard deviation, we also determined the standard deviation per bin, and compared this to the standard deviation per bin of protein coding genes in BIOS^1^, a blood dataset (**Supplementary Figure 10B**). Comparing between bins, we did not see difference in standard deviation for higher expressed bins in either *MetaBrain* or BIOS.

### Cis-eQTL concordance between tissues and ethnicities

Tissue specific^12^, brain region specific^6^, and to a lesser extent, population specific^13^ eQTLs have been observed in previous reports. Our dataset allowed us to evaluate the influence of these factors on eQTL concordance, by comparing the discovered eQTLs between different brain tissues and ethnicities.

We examined eQTLs discovered in the Cortex-EUR and Cortex-AFR datasets and had access to a smaller East-Asian dataset (Cortex-EAS; n=208, limited to ENA dataset). Due to limited sample size, we observed fewer eQTLs in AFR (5,440) and EAS datasets (231). However, concordance in allelic direction was high (>95.67%) if the eQTL was significant (FDR<0.05) in both datasets (**Figure 3C**). Due to the limited sample size and genotype density of Cortex-EAS, concordance rates were generally lower for that population. These results indicate that eQTLs from the same tissue generally share the same allelic direction across populations and that sample size strongly determines how many *cis-*eQTLs can be found and replicate significantly in another population.

We next compared the different brain region datasets for the EUR population. Again, we observed high concordance of allelic directionality (>94.58%; **Figure 3C**). Cerebellum had overall lower concordance with other brain regions (94.58%, 96.96%, 96.95%, 96.33% for Cortex-EUR, Basal ganglia, Hippocampus, and Spinal cord, respectively, **Figure 3C**). We reasoned this might be independent of sample size, since cerebellum has the second largest sample size in *MetaBrain*. Rather, this suggests a difference in genetic regulation between cerebellum and other tissues of the brain.

Although the concordance of the shared eQTL effects was high between brain regions, a substantial number of eQTL genes were only found in one region. For example, for Cortex-EUR, we identified 5,514 unique eQTL genes, and 846 were unique for Cerebellum-EUR (**Supplementary Figure 11A**). For 690 of these genes (82%), the expression in cerebellum was higher than in Cortex-EUR (**Supplementary Figure** **11B**).

Nevertheless, 681 of these 846 genes had a high expression in Cortex-EUR (**Supplementary Figure 11C**). Consequently, we reasoned that these genes are regulated by different transcription factors (TF) in cerebellum and cortex. We observed that they were indeed enriched for binding sites of many TFs, nine of which (*EOMES, TFAP2B, IRX1, IRX5, HES7, ETV2, ELF3, and RUNX3)* are highly expressed in cerebellum (log_2_(TMM+1) >4) and lower (log_2_(TMM+1) <4) in cortex (**Supplementary Figure 11D**). This difference in TF expression might explain why these eQTLs are found in the Cerebellum dataset but not in the larger sample size Cortex dataset.

We next evaluated whether the high rate of sharing between tissues was specific to *MetaBrain*: we repeated the Cortex-EUR discovery while omitting GTEx samples, and subsequently attempted replication in the different GTEx tissues^14^. Cerebral and cortex regions of the brain had highest directional concordance with Cortex-EUR eQTLs (>99%), while concordance in cerebellum was more comparable to other, non-brain, tissues (**Figure 3D, Supplementary Figure 12; Supplementary Table 6**). The overall allelic concordance was highest in GTEx brain samples (>90%), and the lowest in testis (84%) and whole blood (85%). We next evaluated whether the lower concordance in blood was due to the limited sample size of GTEx, by determining the number of concordant Cortex-EUR eQTLs in eQTLgen, a large blood-based dataset (n=31,684). This resulted in an allelic concordance of only 76% (**Supplementary Figure 13**): thus 24% of the shared eQTLs between blood and brain show opposite allelic effects. Since the procedures for eQTL mapping were identical between *MetaBrain* and eQTLgen, we conclude that when eQTL cohort sample-sizes become large it becomes apparent that there is strong tissue-specific regulation, which can explain these opposite allelic effects^15^**.**

### *Cis*-eQTL LD overlap

We evaluated whether the detected *cis*-eQTLs could be linked to neurological traits and diseases, using three different approaches: we determined linkage disequilibrium (LD) overlap, we performed a Mendelian Randomization (MR) approach and we performed statistical colocalization.

First, we investigated LD overlap between *cis*-eQTL SNPs showing the strongest association per gene (i.e. the index eSNP) and 3,279 variants previously identified in 989 GWAS for neurological traits. For this analysis we focused on *MetaBrain* *cis*-eQTL discovery datasets from EUR populations, since most GWAS studies are limited to that population. For each included GWAS trait, we first determined which variants were in LD with any of the *cis*-eQTL index SNPs (R^2^>0.8, calculated using European individuals in the CMC dataset), when the distance between GWAS SNP and eQTL SNP was less than 5Mb. Next, for each GWAS, we clumped the index variants when they were within 1Mb of each other and when they had an LD R^2^>0.2. For each of the clumped GWAS regions, we reported the linked eQTL index SNPs in **Supplementary Table 10**.

Focusing on the Cortex-EUR primary *cis*-eQTLs, 313 (10%) of the trait SNPs were in LD (r^2^>0.8, distance between SNPs <1Mb, LD calculations performed in EUR subset of CMC) with at least one eSNP, linking a total of 242 eQTL genes. Six traits had 10 or more variants in high LD, including years of schooling (21 variants, 8%), multiple sclerosis (MS; 19 variants, 13%), cognitive performance (17 variants; 13%), intelligence (16 variants; 11%), schizophrenia (11 variants; 16%), and neuroticism (10 variants, 11%). Investigating LD overlap in Cerebellum-EUR, we observed smaller numbers: 185 trait variants were in LD with an index eSNP, linking a total of 131 genes. The only trait having 10 or more variants linked in Cerebellum-EUR was cognitive performance (10 variants; 8%). In Hippocampus-EUR, Spinalcord-EUR and Basalganglia-EUR, no traits had 10 or more variants in LD with an index eSNP (**Supplementary Table 10**). Including secondary, tertiary, and quaternary ranked eQTLs marginally improved LD overlap in Cortex-EUR: 393 (12%) unique trait SNPs were in LD with at least 1 index eSNP. In Cortex-EUR, 239 out of 11,803 primary eQTL SNPs were linked to at least one GWAS SNP, while 44 out of 7,047 secondary, tertiary or quaternary eQTL SNPs were linked to at least one GWAS SNP. Consequently, we observed that primary eQTL SNPs were 3.3-fold more likely to be in LD with an GWAS SNP compared to secondary, tertiary and quaternary eQTL SNPs (Fisher exact test p-value: 6.2x10^-16^).

### Enrichment analysis

#### Enrichment of cerebellum vs cortex specific *cis-*eQTL genes

We identified 5,514 *cis*-eQTL genes unique to Cortex-EUR, and 846 for Cerebellum-EUR (**Supplementary Figure 11A**). The large number of genes unique to Cortex-EUR is likely due to the much larger sample size of this dataset compared to the other *MetaBrain* eQTL discovery datasets: we expect that if the sample size of the other datasets would have been larger, the number of significant genes for those datasets would increase, and likely also the overlap of significant genes between datasets. Interestingly, however, we did observe many genes unique to Cerebellum-EUR, which consequently is less likely due to the sample size of this dataset.

We therefore focused on the Cerebellum-EUR unique genes. For some of these genes, it is likely that they are a Cerebellum-EUR specific eQTL because they are expressed in cerebellum and not in cortex.

We observed that the expression levels of these 846 genes in cortex was binomially distributed, and assigned 184 genes on the left side of the minima of this binomial distribution as low-expressed cortex genes, and 681 genes on the right as high-expressed cortex genes (**Supplementary Figure 11B**). We observed that of the 864 genes, 690 (79.77%) had higher expression levels in cerebellum (**Supplementary Figure 11C**, blue line indicates the cut-off between low and high-expressed cortex genes). We took the 681 genes that are unique eQTL genes for Cerebellum-EUR, but high-expressed in cortex and used g:Profiler^16^ to perform functional enrichment (**Supplementary Table 5**).

This analysis included TRANSFAC^10^ enrichment of transcription factor sites around the eQTL genes. We used these enrichments to determine whether genes that are highly expressed in cortex but only have a *cis*-eQTL in cerebellum show a difference in enrichment for specific transcription factor binding sites. We therefore extracted all transcription factors from the g:Profiler table and because transcription factors can be enriched multiple times for different binding sites, we deduplicated the results. We converted the TRANSAC IDs to ENSEMBL IDs using GeneCards^17^ and plotted the expression of each transcription factor in cerebellum and cortex (**Supplementary Figure 11D**). Five of these transcription factors (*EOMES, TFAP2B, TFAP2A, IRX1* and *IRX5*) are lowly expressed in cortex and highly expressed in cerebellum, and these could further explain why some of their target genes are eQTLs in cerebellum and not in cortex, while being expressed in both.

**Enrichment of primary vs non-primary *cis-*eQTL genes**

Enrichment of primary vs non-primary *cis*-eQTLs was done using g:Profiler^16^ with default parameters.

#### Gene co-regulation network

For the gene co-regulation network, gene expression was quantified using Kallisto^18^ (version 0.43.1) to be in line with the gene expression quantification for the gene co-regulation network as described by Deelen *et al*.^19^ CRAM files created during RNA-seq processing for the eQTL analysis were converted to FASTQ files using SAMtools^20^ (version 1.9). Quantification was done against the Ensembl^21^ v98 transcriptome with the patch chromosomes removed. The index was built using default options, with k-mer size = 31. For both paired-end and single-end samples the --bias option was used. For the paired-end samples all other options used default values. For single-end samples --fragment-length=200 and --sd=20 were used, for all other options the default values were used.

After transcript quantification, transcript counts were summed to gene counts using GENCODE^22^ v32 primary assembly GTF for transcript to gene mapping. Genes that showed zero variance over all samples were removed for further analysis. Raw counts were quantile normalized before running PCA analysis to identify and remove possible outliers, but no outliers were detected, likely because only samples that passed the quality control for the eQTL analysis were included. Subsequently, raw counts were normalized by the median of ratios method described in DESeq^23^ before technical covariates identified for the eQTL analysis were removed. From the DESeq^23^ normalized and covariate removed expression data a gene-gene Pearson correlation matrix was calculated and eigenvector and eigenvalues were calculated on this matrix using eigenvalue decomposition. The eigenvectors were centered and scaled and a gene-gene Pearson correlation matrix was calculated on the eigenvector matrix.

For 6 types of gene set databases (KEGG, REACTOME, HPO, GO Molecular Function, GO Biological Process, and GO Cellular Component) coregulation and gene set predictions were calculated with GeneNetwork. The gene set predictions use the number of eigenvectors to include as a parameter. To determine the optimal number of eigenvectors to use we repeated the gene set predictions for n eigenvectors between 25-1000 with steps of 25, and 1000-2000 with steps of 200, and for each gene set database selected the number of eigenvectors with the highest mean AUC (**Supplementary Figure 28**). Heatmaps of the Pearson correlation of the AUC values between different steps shows that there is not much difference in AUC’s once a certain number of eigenvectors has been reached (generally around 100-225 eigenvectors, **Supplementary Figure 29**). The AUC was calculated using a leave-one-out procedure described previously^19^. Given the optimal number of eigenvectors, we created for all genes (coding and non-coding) enrichment scores for each of the 6 databases.

### Mendelian randomization (MR)

#### Methods

##### Annotating suggestive MR findings

Methods for performing MR and colocalization are found in the main manuscript. In addition, we also compared the suggestive MR findings with two catalogues of Mendelian diseases, Development Disorder Genotype - Phenotype Database (DDG2P) and OrphaNet. Because we focused on neurological traits for MR, only the developmental disorders that affected Brain/Cognition in DDG2P, as specified in the “organ specificity list” column, were annotated^24^ (data freeze 2020-12-16). We looked for genes that appear to follow an allelic series^25^.

To annotate with OrphaNet data, the November 2020 release of XML files were downloaded from their GitHub repository^26^, and Python package xmltodict^27^ was used to aid in data parsing. Information was extracted from the following files and combined on OrphaNet ID via pandas^28^ rare disease epidemiology, rare diseases natural history, genes associated with rare diseases and classifications of rare diseases.

#### Results

Suggested MR findings (Wald ratio p<5x10^-5^) for the disease trait outcomes and brain volume outcomes are reported in **Supplementary Table 12**. The p-value threshold for significance after adjusting for 268,030 tests performed is 1.865x10^-7^.

##### Alzheimer’s disease

There were 32 significant Wald ratios for Alzheimer’s disease (AD, of which 10 passed colocalization (PP4>0.7). Higher expression of five genes (*CR1*, *TSPAN14*, *ZNF668*, *CCDC6* and *APH1B*) and lower expression of five genes (*CASS4*, *PRSS36*, *ZNF646*, *KAT8* and *ACE*) were associated with increased AD risk.

*CR1* is a top-ranked AD risk gene expressed by microglia and involved in Ab clearance. Previous studies have identified CR1-B/S allele, coding the longer isoform with additional C3b binding site, associated with increased AD risk^29,30^. Some evidence suggested increased activity of *CR1* can potentially inhibit complement cascade activated by Ab and lead to enhanced Ab deposition^31,32^. Consistently, we found higher expression of *CR1* associated with increased risk of AD (Wald ratio or WR=0.15, p=1.4x10^-23^). However, the molecular underpinnings of CR1’s role in AD pathogenesis are still not yet clear.

*TSPAN14* resides in a recently identified AD risk locus^33^, it belongs to the TspanC8 family of tetraspanins and interacts with ADAM10, which is encoded by another AD risk gene and mediates proteolytic cleavage of more than 40 substrates including APP and TREM2^34^. Overexpression of *TSPAN14* has been shown to increase cell surface expression of ADAM10^35^. Our MR and colocalization analysis using rs10749609 as an eQTL instrument variable found higher expression of *TSPAN14* as putatively causal for AD (MR Wald ratio=0.17, p=6.7x10^-9^; Coloc PP4>0.95). *ADAM10* was also borderline significant in our MR analysis (p=2.67x10^-7^) for AD, but its Cortex-EUR *cis*-eQTL did not colocalize with the AD GWAS locus (Coloc PP4=0.08). Missense mutations in *ADAM10* that attenuate its α-secretase activity shift APP processing toward β-secretase-mediated cleavage, increase Aβ plaque load and reactive gliosis^36^. However, it is not well understood how cleavage activity of APP by ADAM10 is impacted by expression of *TSPAN14*, though some *in vitro* evidence showed decreased APP-ADAM10 cleavage by stably expressing *TSPAN14* in U2OS-N1 cells^35^. ADAM10 also mediates cleavage of TREM2, another microglial gene long-established in AD pathology^37^. Both human and preclinical data supported the association between TREM2 deficiency and risk of AD^37^, and scRNA sequencing data also revealed more pronounced expression of *TSPAN14* and *ADAM10* in microglia than neurons^38^. Lower expression of *TREM2* is suggestively associated with increased AD risk (Wald ratio=-0.28, p=2.6x10^-7^), however, the Cortex-EUR eQTL only colocalized with a secondary GWAS signal after conditioning on the primary signal which was driven by rare variants (MAF<0.01) (**Supplementary Table 13**). Enhanced shedding of TREM2 has been discovered for a Han Chinese late onset AD-associated *TREM2* coding variant, while reduced TREM2 shedding has been found when ADAM10 was inhibited^39,40^. Though the mechanism of TSPAN14 regulated TREM2-ADAM10 shedding remains unclear, it has been shown that overexpression of *TSPAN14* increases the surface expression of ADAM10^35^. Further experiments are needed to better understand how the effect of higher expression of *TSPAN14* affects the cleavage of TREM2 by ADAM10, which could potentially alter TREM2‐dependent phagocytosis and microglial function.

*ACE* encodes for the angiotensin-converting enzyme that can convert angiotensin I to angiotensin II, where the latter can constrict blood vessels and increase blood pressure^41^. ACE-inhibitors are widely used as hypertension medication, but ACE also converts Ab42 to Ab40 and has been implicated in AD^42^. Some evidence showed inhibiting ACE increases amyloid deposition in AD mice models and some hypertensive patients taking ACE inhibitors exhibited worsen decline in cognitive function^43^. However, other studies found lower AD risk for those who took ACE inhibitors^44,45^ and null effects of ACE inhibition on Ab levels *in vivo*^46,47^. A recent report found rare *ACE* coding variants and developed a knock-in mouse model for one missense mutation (p.R1279Q), and observed increased ACE protein and activity, together with EEG disruption, memory impairment, neuroinflammation and hippocampal neurodegeneration, but no effect on Ab^48^. In addition, the adverse outcomes of *ACE* p.R1279Q can be rescued by brain-penetrant medications inhibiting ACE which seems conflicting with our MR findings suggesting lower expression of *ACE* leads to increased AD risk (WR=-0.15, p=1.4x10^-7^).

*APH1B* contributes to γ-secretase step of APP leading to Ab accumulation. It has been suggested that targeting Aph1B γ-secretase complex can potentially lower Aβ peptide production in human AD^49^. However, a recent study tested the impact of a missense variant in *APH1B* (rs117618017, in high LD with top *MetaBrain* eQTL rs75763893, EUR r^2^=0.97) observed no effect on the γ-secretase processing of established substrates compared with cells expressing wild-type *APH1B*^50^.

*CASS4* has been discovered as AD susceptibility gene^51^ and the risk SNP (rs7274581) has also been associated with core neuropathologic features of AD including neuritic plaques and neurofibrillary tangles in a large cohort with brain autopsies^52^. We found lower expression of *CASS4* associated with increased AD risk (WR=-0.42, p=6.5x10^-10^) and shared causal variant between *MetaBrain* eQTL and GWAS signal (PP4=0.99), but the molecular mechanism of *CASS4* in AD pathology remains unclear.

At the 16p11.2 locus harboring *ZNF646*, *ZNF668*, *KAT8* and *PRSS36*, these genes were also identified with MR and colocalization evidence. However, high LD in this region made it difficult to ascertain the causal gene and the biological involvements of these genes in AD remain unclear. We also identified *CCDC6* which is a tumor-suppressor gene and its fusion with RET has been found in multiple cancers^53^ *CCDC6* is functionally involved in apoptosis mediated by ATM upon DNA damage.

We also evaluated 60 of the 69 AD associated eQTL instruments for an interaction effect with cell type proportions. In total, 7 eQTL instruments where significant ieQTLs, 5 with macrophages (*CD33*, *FCER1G*, *SIGLEC11*, *HLA-DQA2* and *HLA-DQB1*), 1 with oligodendrocytes (*SLC25A48*) and 1 with neurons and oligodendrocytes (*NFYA*). Of these, 4 eQTL instruments passed the significance threshold only for colocalization (*CD33*, *FCER1G*, *SIGLEC11* and *SLC25A48*).

##### Attention deficit/hyperactivity disorder (ADHD)

There were no significant Wald ratios for attention deficit/hyperactivity disorder (ADHD). There were five suggestive signals (p<5x10^-5^) for ADHD that pass colocalization, suggesting increased expression of *ARID5B* (WR=0.31, p=3.80 x10^-5^), *GMPPB* (WR=0.10, p=1.36x10^-5^), *SIRPD* (WR=0.32, p=1.53x10^-5^), *KIFC2* (WR=0.37, p=4.92 x10^-5^) and *PIDD1* (WR=0.21, p=7.13x10^-6^) increased risk for ADHD. None of these loci were genome-wide significant in the outcome GWAS, so the lack of power could be a contributing factor for the sub-significant findings^54^.

We also evaluated 10 of the 10 ADHD associated eQTL instruments for an interaction effect with cell type proportions. In total, 2 eQTL instruments where significant ieQTLs, 1 with endothelial cells (*TIE1*) and 1 with oligodendrocytes (*ELOVL1*). Of these, 1 eQTL instrument passed the significance threshold only for MR (*ELOVL1*).

##### Amyotrophic lateral sclerosis

There were two significant Wald ratio findings for amyotrophic lateral sclerosis (ALS) that passed colocalization, *SCFD1* (WR=0.092, p=5.31x10^-15^) and *G2E3* (WR=0.092, p=5.70x10^-15^). The genes are adjacent and the eQTL instruments are in high LD (EUR r^2^=0.99), so it would be difficult to discriminate the underlying biology at this locus. The *SCFD1* gene encodes for the Sm protein Sly1, and facilitates the SNARE complex formation^55^. The SNARE proteins facilitate neurotransmitter release and they have a protective role against neurodegeneration^56^. *G2E3* is a nucleo-cytoplasmic shuttling protein with a HECT domain that controls the protein localization. It has been hypothesized to play a role in cell cycle regulation and response to DNA damage^57^.

We also evaluated 23 of the 25 ALS associated eQTL instruments for an interaction effect with cell type proportions. In total, 4 eQTL instruments where significant ieQTLs, 1 with astrocytes (*TNFSF13*) and 3 with neurons (*SCFD1*, *RESP18* and *DHRS11*). Of these, 1 eQTL instrument passed the significance threshold for both MR and colocalization (*SCFD1*) and 2 eQTL instruments passed the significance threshold only for colocalization (*RESP18* and *TNFSF13*).

##### Autism spectrum disorder

There were no significant Wald ratio findings for autism spectrum disorder (ASD). There were two suggestive signals for ASD that pass colocalization, suggesting increased expression of *GABBR1* (beta=0.30, p=2.19x10^-6^) and decreased expression of *PPP1R3B* (WR=-0.38, p=8.91x10^-6^) increased risk for ASD.

We also evaluated 8 of the 12 autism associated eQTL instruments for an interaction effect with cell type proportions. In total, 1 eQTL instrument was a significant ieQTL; interacting with macrophages and oligodendrocytes (*ARL17B*). This ieQTL did, however, not overlap with significant MR and colocalization signals.

##### Bipolar disorder

There were 8 significant Wald ratios for bipolar disorder, of which 6 passed colocalization (PP4>0.7). We find that downregulating *LMAN2L* (WR=-0.32, p=1.93x10^-8^) and *DCLK3* (WR=-0.53, p=4.79x10^-14^) increase risk for bipolar disorder and upregulating *GNL3* (WR=0.34, p=1.07x10^-8^), *HAPLN4* (WR=0.34, p=3.07x10^-9^), *CILP2* (WR=0.52, p=3.68x10^-8^) and *TM6SF2* (WR=0.36, p=9.75x10^-8^) increase risk for bipolar disorder.

*LMAN2L* encodes a transmembrane protein belonging to the L-type lectin group of type 1 membrane proteins. The LMAN2L protein is located at the endoplasmic reticulum and function in the mammalian early secretory pathway as a cargo receptor in the transport of glycoproteins. Homozygous mutation *LMAN2L* is known to cause autosomal recessive mental retardation^58^ and this gene has also been previously implicated with bipolar disorder due to its interaction with *ANK3*^59^.

*DCLK3* is a member of the doublecortin family and encodes serine/threonine kinase-domains that show substantial homology to Ca2+/calmodulin-dependent (Cam) protein kinases that involves in regulate cyclic AMP (cAMP) signaling. Broadly, members of the doublecortin family involve in neuronal migration, neurogenesis and eye receptor development and are associated with subcortical band heterotopia, lissencephaly, epilepsy, developmental dyslexia and retinitis pigmentosa.

*GNL3*, G protein nucleolar 3, is important for stem cell proliferation and involved in maintaining stem cell self-renewal. It is also suggested that *GNL3* encodes for a protein that interacts with p53 and involves in cell death caused by overexpression, as well as tumorigenesis^60^. *GNL3* is associated with schizophrenia, bipolar disorder^61^ and osteoarthritis^62^.

There is very little information on the gene function of *HAPLN4*, hyaluronan and proteoglycan link protein 4. There are GWAS associations for bipolar disorder, lipid levels (LDL, triglyceride, total cholesterol;^63^) and metabolites near this gene^64^ . Gene Ontology (GO) annotations related to *HAPLN4* include extracellular matrix structural constituent and hyaluronic acid binding.

*CILP2*, cartilage intermediate layer protein 2, is associated with cognitive function^65^, multiple lipid phenotypes including LDL, triglycerides^66,67^, as well as blood cell traits such as monocyte count and red cell distribution width^68^. Gene Ontology (GO) annotations related to CILP2 include carbohydrate binding and nucleotide diphosphatase activity.

*TM6SF2*, transmembrane 6 superfamily member 2, is significantly associated with type 2 diabetes^69^, triglycerides, LDL^70^, total cholesterol^67^, but interestingly it is not significantly associated with bipolar disorder with a genome-wide significant p-value.

Interestingly, *CCILP2* and *TM6SF2* are also significant MR findings for schizophrenia.

We also evaluated 51 of the 52 bipolar disorder associated eQTL instruments for an interaction effect with cell type proportions. In total, 5 eQTL instruments where significant ieQTLs; all of which interacted with neurons (*TRANK1*, *RHEBL1*, *MCHR1*, *MED24* and *CDHR1*). Of these, 1 eQTL instrument passed the significance threshold only for MR (*TRANK1*) and 4 eQTL instruments passed the significance threshold only for colocalization (*RHEBL1*, *MCHR1*, *MED24* and *CDHR1*).

##### Epilepsy

There were 3 significant Wald ratios for epilepsy, of which 2 passed colocalization (PP4>0.7). MR analysis was performed for several epilepsy outcomes. Epilepsy (all documented cases) was the all-encompassing disease outcome, and additional epilepsy subtype outcomes were also analyzed, including generalized epilepsy (all documented cases), focal epilepsy (all documented cases), focal epilepsy (documented lesion negative), juvenile absence epilepsy, childhood absence epilepsy, focal epilepsy (documented hippocampal sclerosis), focal epilepsy (documented lesion other than hippocampal sclerosis), generalized epilepsy with tonic-clonic seizures and juvenile myoclonic epilepsy^71^. There were two significant Wald ratio findings that passed colocalization across the epilepsy outcomes. The first finding suggests that downregulation of CDK5RAP3 increases risk for generalized epilepsy, all documented cases (WR=-0.21, p=6.79x10^-9^). CDK5RAP3 encodes for the catalytic CDK5 submit of neuronal CDC2-like kinase. It has been hypothesized to be involved in neuronal differentiation^72^. This gene appears to be highlight upregulated in epileptic patients (>11-fold upregulation observed in a small cohort of 8 epileptic and 5 healthy children). In a larger group of children, including 32 epileptic patients that were treated with vaporic acid for 12 months, common drug to treat seizures, decreased CDK5RAP3 was observed in these patients^73^. The second finding suggests that upregulating HSD3B7 increases risk for juvenile myoclonic epilepsy (WR=0.065, p=3.97x10^-9^). HSD3B7 is thought to be a metabolic enzyme that participated in bile acid synthesis^74^.

We also evaluated 27 of the 32 epilepsy associated eQTL instruments for an interaction effect with cell type proportions, none of which were significant.

##### Frontotemporal dementia

There was one significant Wald ratio finding for all subtypes of frontotemporal dementia (FTD) that also passed colocalization. It suggest that upregulating BTNL2 (WR=0.76, p=2.20x10^-10^) increased risk for FTD. This gene is located near the major histocompatibility complex (MHC) region, and the underlying biology of this locus is thought to be driven by the critical immunological genes in the region^75^. BTNL2 is known to inhibit T cell activation^76^, so the direction of effect found in the MR analysis is confusing. A protein truncating splicing variant in BTNL2 that reduces protein function has been associated with an inflammatory disease known as sarcoidosis^77^, where small subset of the patients present with rapidly progressing dementia^78^. No signal was found for frontotemporal dementia, TDP-43 subtype GWAS outcome, likely due to the low resolution of the GWAS summary statistics.

We also evaluated 2 of the 2 frontotemporal dementia associated eQTL instruments for an interaction effect with cell type proportions, none of which were significant.

##### Major depressive disorder

There were two significant Wald ratio findings for Depression (broad) or major depressive disorder that also passed colocalization. Upregulating *NEGR1* (WR=0.030, p=3.74x10^-8^) and *SLC12A5* (WR=0.040, p=8.71x10^-8^) increase risk for major depressive disorder.

In addition to depression, *NEGR1*, neuronal growth regulator 1, is associated with neurociticism^79^, insomnia^80^, schizophrenia^81^, cognitive function^82–84^, late onset Alzheimer's disease^85^, BMI^86^, and lupus^87^ in GWAS. The pleiotropic association between *NEGR1* and psychiatric traits and BMI and obesity-related traits suggests a neuronal component of obesity^88^.

*SLC12A5*, solute carrier family 12 member 5, encodes the neuronal KCC2 channel that is the major extruder of intracellular chloride in mature neurons. SLC12A5 is exclusively expressed in the central nervous system. Homozygous or compound heterozygous mutation in the *SLC12A5* causes early infantile epileptic encephalopathy (OMIM: 616645) and heterozygous mutation in the *SLC12A5* gene increases the susceptibility of generalized epilepsy (OMIM: 616685). In addition to depression, *SLC12A5* is associated with neuroticism^89^ and chronotype^90^ in GWAS. Of note, *SLC12A5* did not show genome-wide significant association in the depression GWAS analyzed in this study, and *SLC12A5* is also a significant MR finding for multiple sclerosis.

We also evaluated 26 of the 27 depression associated eQTL instruments for an interaction effect with cell type proportions. In total, 2 eQTL instruments where significant ieQTLs; both interacting with macrophages (*BTN3A2* and *HLA-DMB*). None of these ieQTLs, however, overlapped with significant MR and colocalization signals.

##### Multiple Sclerosis

Through our *MetaBrain* eQTL MR approach followed by colocalization filtering, we found 20 genes that are significant and supported by GWAS-eQTL colocalization for multiple sclerosis (MS). MR analysis suggested higher expression of eleven genes (*HLA-DRB1*, *SLC12A5*, *CYP24A1*, *TRAF3*, *IFITM1*, *TBX6*, *MYO19*, *TSFM*, *NPEPPS*, *TSPAN31* and *SCO2*) and lower expression of nine genes (*MPV17L2*, *CCDC155*, *IFITM3*, *TTC34*, *EEF1AKMT3*, *RNFT1*, *IL7*, *MYNN*, *KMT5A* and *CLECL1*) in the cortex causally associated with greater risk in MS.

The HLA genes were the earliest established MS susceptibility genes^91–93^ and the HLA-DRB1*15:01 allele was a major MS risk factor^94^. Previous studies found the MS risk allele associated with higher HLA-DRB1 expression^95,96^, and hypomethylation of HLA-DRB1 could be mediating the expression change and lead to increased disease risk^97^. Our MR analysis also suggested a putative causal effect of increased HLA-DRB1 expression level (WR=0.81, p=1.4x10^-13^) in the etiology of MS.

For *SLC12A5* we observed a conflicting directionality where loss of function mutations were found previously to be pathogenic for pediatric epilepsy^98,99^, but our MR results showed increased expression associated with MS risk (WR=0.71, p=1.9x10^-12^). Epileptic genes are generally tightly regulated so it is possible to observe detrimental effects in both directions. Interestingly, this locus also harbors *CD40*, which is another strong candidate gene for MS risk but CD40 locus did not colocalize with the cortex *cis*-eQTL. A secondary signal for *CD40* likely exists at this locus but was undetected based on current data, due to potential cell-type specific effects where *SLC12A5* is more neuronal but *CD40* acts through microglia, phagocytes and endothelial cells^100^.

*CYP24A1* is discussed in the main results.

*TRAF3* encodes one of the tumor necrosis factor receptor (TNFR) associated factors binding one of the TNFR superfamily member CD40, and TRAF3 plays an inhibitory role in CD40 signaling in B lymphocytes^101^. Although it remains mechanistically unclear how genetically determined higher expression of *TRAF3* contributes to MS risk (WR=0.37, p=1x10^-8^), TRAF3 deficiency was shown to disrupt the signaling pathway of Epstein Barr virus (EBV)-encoded CD40 mimic, Latent Membrane Protein 1 (*LMP1*), which is necessary for EBV-associated lymphoproliferation and potentially MS etiology^102^.

*IFITM1* and *IFITM3* are interferon (IFN)-induced transmembrane proteins and are involved in IFN signaling pathways. IFN-β was the first therapy approved that could change the course of MS. However, we observed opposite directionalities between *IFITM1* (WR=0.49, p=1.8x10^-8^) vs. IFITM3 (WR=-0.44, p=2.3x10^-8^) in MS risk. The biological mechanisms for IFITMs in MS remain unclear but a recent report suggested IFITM1 increases the infectivity of EBV^103^, further evidence is needed to understand the involvement of IFITMs in MS etiology.

We also found two genes: *TBX6* and *CCDC155*, though the biological understanding of their involvement in MS is lacking, with colocalization evidence for cis-methylation QTL (cis-mQTL) in CD4+ T cells derived from MS patients^104^. For the TBX locus, the MS risk allele (rs3809627-C) is associated with hypomethylation and consistently we observed higher expression of TBX6 is associated with increased MS risk (WR=0.23, p=3.2x10^-8^). For the CCDC155 locus, the MS risk allele (rs1465697-T) is associated with hypermethylation and we also observed lower expression of *CCDC155* associated with MS risk (WR=-0.62, p=9x10^-10^).

*MYO19* and *MPV17L2* are both associated with mitochondria^105,106^. Although the functional mechanism of how these two genes contribute to MS remain unclear, mitochondria dysfunction can contribute to neurodegeneration and axonal loss in MS^101,107^.

We found lower expression of *TTC34* associated with higher risk in MS (WR=-0.51, p=2.0x10^-15^), but the biological mechanisms of *TTC34* involvement in MS etiology remain unclear. A few reported relevance of *TTC34* in autoimmune diseases such as systemic lupus erythematosus (SLE)^108,109^ and type I diabetes^110^ .

We found three genes with significant MR associations and positive colocalization signals from the 12q14.1 locus, *EEF1AKMT3*, *TSFM* and *TSPAN31*. Lower expression of *EEF1AKMT3* (WR=-0.15, p=5.1x10^-11^) and higher expression of *TSFM* (WR=0.21, p=1.6x10^-10^) and TSPAN31 (WR=0.65, p=1.7x10^-9^) were found putatively causal for increasing MS risk. *EEF1AKMT3* encodes a lysine methyltransferase targeting methylation of Lys-165 in EEF1A in an aminoacyl-tRNA- and GTP-dependent manner^111,112^. *EEF1A* encodes eukaryotic elongation factor 1A (with two isoforms, EEF1A1 and EEF1A2) and plays an important role in regulation of protein elongation and synthesis^113^. Lys-165 methylation of EEF1A-coupled with upregulation of *EEF1AKMT3* can be increased by various types of stress including ER-stress^111^ and EEF1A overexpression under ER-stress has been found to prevent apoptosis^114^. Previous reports showed evidence on the potential involvement of EEF1A and apoptosis pathway in neurodegeneration and PD^115–117^. In humans, mutations in EEF1A2 have been linked with multiple neurological deficits including developmental delay, autistic behaviors and epilepsy^118–120^, and dysregulation of EEF1A is also observed in AD^121,122^. *TSFM* encodes a mitochondrial translation elongation factor and plays an important role in mitochondrial protein translation. Mutations in TSFM have been associated with rare infantile mitochondrial disorders with various clinical manifestations including fatal encephalomyopathy, cardiomyopathy, neuropathy and childhood onset ataxia^123–125^. Although the biological mechanisms of *TSFM* in MS etiology are not well-understood, the involvement of mitochondrial genes and pathways have been described in MS^126^, including three (*TSFM*, *MYO19* and *MPV17L2*) that were found by our MR analysis. *TSPAN31* is the natural antisense transcript of cyclin dependent kinase 4 (CDK4) and regulates the expression of *CDK4* mRNA and protein^127^. Higher expression of *TSPAN31* can potentially down-regulate *CDK4* and prevent cell proliferation, and CDK4/6-inihibitors have been developed as a therapeutic strategy to treat multiple cancers^128^. Some evidence has been shown for the potential involvement of CDK4 in regulating immune cells where CDK4/6-inhibition promotes T-cell activity^129,130^. However, others have also reported potential treatment effect of CDK4/6-inhibitors for autoimmune diseases such as RA without obvious effect on immune response^131^.

*NPEPPS*, also known as puromycin-sensitive aminopeptidase, encodes for a protein that hydrolyzes physiological endogenous peptides such as dynorphins and enkephalins, and contributes to the degradation of these peptides in the brain^132^. Interestingly, NPEPPS has been reported to degrade tau protein and inhibit tau-induced neurodegeneration^133,134^, as well as regulate SOD1 via proteolysis in ALS^135^. However, we found higher gene expression of *NPEPPS* leading to higher risk in MS (WR=0.62, p=3.0x10^-10^).

*RNFT1*, or Ring finger protein transmembrane 1, is an E3 ubiquitin-protein ligase participating in the ubiquitin proteasome system (UPS) that induces protein degradation. This protein also has zinc-binding activities^136^. Giordana *et al.* observed abnormally strong colocalization of ubiquitin in the myelinated white matter of all six MS patients analyzed, suggesting that MS patients may suffer from UPS deficits^137^.

Interleukin 7 (*IL7*) regulates naïve and memory CD8+ T cells and has been known to be a critical gene for early T-cell development^138^. This gene has been suggested to play a role in autoimmunity. A decrease systemic IL-7 and soluble IL-7Rα was observed in MS patients^139^, and our MR findings also suggest that abnormalities in the IL-7 pathway could increase risk for MS.

*SCO2*, synthesis of cytochrome c oxidase (COX), is a COX assembly gene that modulates proton transfer across the inner mitochondrial membrane. Loss of function mutations of *SCO2* also cause an autosomal recessive form of fatal cardioencephalomyopathy and COX deficiency^140^, Loss of function mutations in the functional catalytic domain of SCO2 causes an autosomal dominant form of myopia^141^. Several reports have suggested a link between mitochondrial dysfunction and multiple sclerosis^107^.

Myoneurin (*MYNN*) encodes for a protein with a classic C2H2 zinc finger motif and a BTB/POZ protein-protein interaction domain. There is limited information published about the function of this protein, though based on its protein domains, it is hypothesized to play a regulatory function in gene expression and transcriptional activation and repression^142^.
Lysine methyltransferase 5A (*KMT5A*), also known as SETD8, encodes for a lysine methyltransferase for histone H4 and influences transcriptional regulation, heterochromatin formation, genomic stability, cell cycle progression, and development^143^. It is associated schizophrenia, intelligence^84^, insomnia^80^, lymphocyte, eosinophil and neutrophil count^144^. This MR analysis finds that decreasing *KMT5A* expression levels drives risk for MS and schizophrenia. Interestingly, the MR analysis also suggests that decreasing KMT5A levels increase years of schooling and fluid intelligence. This highlights the important role of epigenetic mechanisms in neurological traits^145,146^.

We also evaluated 122 of the 157 MS associated eQTL instruments for an interaction effect with cell type proportions. In total, 16 eQTL instruments where significant ieQTLs, 7 with macrophages (*HLA-DRB1*, *HLA-DQA2*, *HLA-DOA*, *CLECL1*, *IFITM1*, *HLA-DPB1* and *IER3*), 5 with neurons (*CYP24A1*, *RNFT1*, *RMI2*, *CASQ1* and *DHRS11*) and 4 with oligodendrocytes (*ZFP57*, *SFTA2*, *PSORS1C1* and *HIST1H3E*). Of these, 4 eQTL instruments passed the significance threshold for both MR and colocalization (*CLECL1*, *CYP24A1*, *RNFT1* and *IFITM1*), 8 eQTL instruments passed the significance threshold only for MR (*HLA-DRB1*, *HLA-DQA2*, *ZFP57*, *HLA-DOA*, *SFTA2*, *RMI2*, *HLA-DPB1* and *IER3*) and 1 eQTL instrument passed the significance threshold only for colocalization (*CASQ1*).

##### Parkinson’s disease

There were 6 significant WR findings for Parkinson’s disease (PD) that passed colocalization (PP4>0.7) that suggest increasing expression of *GPMNB* (WR=0.12, p=2.26x 10^-8^), *RAB29* (WR=0.29, p=5.15x10^-9^) and *SCARB2* (WR=0.38, p=1.77x10^-8^) and decreased expression of *CD38* (WR=-0.29, p=1.20x10^-13^), *LRRC37A2* (WR=-0.22, p=5.14 10^-16^) and *HSD3B7* (WR=-0.67, p=1.90x10^-10^) increased risk for disease.

Transmembrane glycoprotein NMB (*GPMNB*) is also known as osteoactivin and was first characterized in a melanoma cell line (PMID: 7814155) and causes a Mendelian form of primary localized cutaneous amyloidosis. It has recently become a potential gene of interest for neurodegenerative disorders. Mouse models of AD and patient samples suggest that *GPMNB* is only expressed in the brain under neurodegenerative conditions and show that the GPNMB protein co-localizes with a distinct population of IBA1-positive, activated microglia cells that cluster around amyloid plaques^147^. In PD, it has been shown that GPMNB was elevated in the substantia nigra of PD patients and in wild type mice with CBE-induced GCase lysosomal dysfunction^148^.

The finding that increased *RAB29* expression increases risk for PD risk is interesting, because Rab29 controls LRRK2 activation, localization and possibly phosphorylation as well. *LRRK2* mutations cause the most common autosomal dominant form of Parkinson’s disease and this gene plays a role in the endolysosomal regulation. Pathogenic *LRRK2* mutations are more readily recruit to the Golgi and activated by Rab29^149,150^.

Scavenger Receptor Class B Member 2 (*SCARB2*) is a type III glycoprotein located in the endosomal/lysosomal cell compartment and regulates endolysosomal transport. Autophagy dysregulation has been repeatedly reported as a driving factor for Parkinson’s disease^151^. This gene also causes an autosomal recessive form of progressive myoclonic epilepsy-4 (EPM4), whereby the mutations found in three unrelated probands resulted in the lack of *SCARB2* protein^152^.

*CD38*, also known as cyclic ADP-ribose hydrolase, plays a role in regulating microglia through microglial activation and activation-induced cell death^153^. This enzyme is also responsible for nicotinamide adenine dinucleotide (NAD) degradation, and NAD levels are important in age-related metabolic decline and have been observed to decrease with age^154^.

*LRRC37A2* encodes for the Leucine Rich Repeat Containing 38 Member A2 protein, one of the genes in a novel gene family of hominoid lineage^155^. This gene is also part of a duplication segment flanking the MAPT inversion.^156^

Interestingly, while decreased *HSD3B7* levels could increase risk for PD, we had noted above that upregulating this gene increases risk for juvenile myoclonic epilepsy. The latest PD GWAS publication in eQTLGen also reported significant SMR findings for this gene was also reported in blood (eQTLGen^8^) and in brain^157^ eQTL dataset in opposite directions of effect. The results did not pass the HEIDI p-value threshold, suggesting a potential pleiotropic association invalidating the MR finding. Thus, this finding should be interpreted with caution. We were underpowered to detect associations with Parkinson’s disease due to the lack of power with the publicly available summary statistics^158^.

We also evaluated 37 of the 48 PD associated eQTL instruments for an interaction effect with cell type proportions. In total, 8 eQTL instruments where significant ieQTLs, 3 with astrocytes (*CD38*, *ADORA2B* and *ZNF391*), 1 with macrophages (*GPNMB*), 3 with neurons (*LRRC37A2*, *NUP42* and *CCDC189*) and 1 with macrophages and oligodendrocytes (*ARL17B*). Of these, 2 eQTL instruments passed the significance threshold for both MR and colocalization (*CD38* and *GPNMB*), 2 eQTL instruments passed the significance threshold only for MR (*ARL17B* and *LRRC37A2*) and 2 eQTL instruments passed the significance threshold only for colocalization (*ADORA2B* and *ZNF391*).

##### Schizophrenia

There are 20 schizophrenia MR findings that colocalize (PP4>0.7): *RERE*, *SF3B1*, *FTCDNL1*, *CNTN4*, *DCLK3*, *GLYCTK*, *GNL3*, *THOC7*, *PCCB*, *CLCN3*, *CLIC1*, *MDK*, *KMT5A*, *RGS6*, *FES*, *FURIN*, *INO80E*, *ZNF823*, *MAU2* and *GATAD2A*.

*CLIC1*, chloride intracellular channel 1, is a member of the p64 family, which includes chloride channel proteins that regulate fundamental cellular processes including stabilization of cell membrane potential, transepithelial transport, maintenance of intracellular pH, and regulation of cell volume. *CLIC1* has been reported to be a key regulator of Ca(2+) signaling in cancer survival^159^, and is associated with various cancers including gastric cancer^160^, gallbladder cancer^161^, glioblastoma^162^), liver cancer^163^, and colon cancer^164^. *CLIC1* is also associated with schizophrenia, depression^89^, strep throat^165^ and mean platelet volume^68^ in GWAS.

*FTCDNL1* encodes for formiminotransferase cyclodeaminase N-terminal like, and this protein has been related to transferase activity and folic acid binding in Gene ontology and is associated with schizophrenia, osteoporosis^166^, fracture and type 2 diabetes^167^ in GWAS.

*FURIN*, also known as furin paired basic amino acid cleaving enzyme, encodes a type 1 membrane bound protease that belongs to the subtilisin-like proprotein convertase family. The protease processes protein and peptide precursors trafficking through regulated or constitutive branches of the secretory pathway and is expressed in many tissues, including neuroendocrine and brain. *FURIN* is associated with epilepsy through regulating GABA-A receptors-mediated inhibitory synaptic transmission^168^. More recently, it is also found that *FURIN* plays a crucial role in SARS-CoV-2 spike protein cleavage, which mediates the virus entry into cells^169^. It is also associated with schizophrenia, autism^170^, insomnia^80^, risk-taking behavior^171^, as well as hypertension/blood pressure^80^, coronary artery disease^172^ and parental longevity in GWAS^173^.

*SF3B1*, splicing factor 3b subunit 1, encodes subunit 1 of the splicing factor 3b protein complex, which is part of the U2 small nuclear ribonucleoproteins complex (U2 snRNP) that has an essential role in the selection of the precursor mRNA branch-site adenosine (the nucleophile for the first step of splicing)^174^. It has been shown that inhibition of *SF3B1* reduced cell proliferation, induced apoptosis, and resulted in cell cycle arrest human gastric cancer cells in vitro^175^. The somatic mutations in *SF3B1* have been associated with prolactinomas^176^ and chronic lymphocytic leukemia^177^. *SF3B1* is also associated with schizophrenia and depression^178^ in GWAS.

*PCCB*, propionyl-CoA carboxylase subunit beta, encodes the beta subunit of the propionyl-CoA carboxylase (PCC) enzyme, which is involved in the catabolism of propionyl-CoA. *PCCB* defects are known to be a cause of propionic acidemia type II (PA-2) (OMIM: 606054), intellectual disability (OMIM: 617635) and are associated with autism in propionic acidemia patients^179^. In GWAS, *PCCB* is associated psychiatric traits including schizophrenia, cognitive performance^65^, neuroticism^180^; anthropometric traits including height^181^, body mass index^182^, and waist-hip ratio^182^ and metabolic traits including circulating fibrinogen levels^183^, C-reactive protein levels^184^, HDL cholesterol^185^ and triglycerides^70^.

*CNTN4*, contactin 4, encodes a member of the contactin family of immunoglobulins, which are axon-associated cell adhesion molecules that function in neuronal network formation and plasticity. *CNTN4* is known to be associated with chromosome 3p deletion syndrome. In GWAS, *CNTN4* variants are associated schizophrenia, autism, intelligence, amyotrophic lateral sclerosis, as well as gallbladder cancer^186^ and basophil count^68^.

*DCKL3* is described in the Bipolar disease section.

*MDK*, or midkine, encodes a retinoic acid-responsive, heparin-binding growth factor expressed in various cell types during embryogenesis. MDK protein promotes angiogenesis, cell growth, and cell migration in particular during tumorigenesis. It is associated with schizophrenia, autism^170^ and smoking initiation^187^ in GWAS and has been targeted as a therapeutic for a variety of different diseases, such as non-small cell lung cancer^188^ and hepatocellular carcinoma^189^.

*THOC7*, THO complex 7, encodes a protein that is part of the THO complex, which together with ALY and UAP56 forms the human TREX complex. The TREX complex is recruited to spliced mRNAs by a transcription-independent mechanism and is recruited in a splicing- and cap-dependent manner to a region near the 5' end of the mRNA where it functions in mRNA export to the cytoplasm via the TAP/NFX1 pathway. In addition to schizophrenia, *THOC7* is also associated with neutrophil count in GWAS^144^.

*FES*, also known as FES proto-oncogene, tyrosine kinase, encodes the human cellular counterpart of a feline sarcoma retrovirus protein with transforming capabilities. The gene product has tyrosine-specific protein kinase activity that is required for maintenance of cellular transformation. *FES* is associated cardiovascular disease^190^, blood pressure^191^, and insomnia^80^ in addition to schizophrenia.

*KMT5A* is described in the multiple sclerosis section.

*INO80E*, INO80 complex subunit E, is associated with schizophrenia, body mass index^192^ and depression related symptom^1^.

*GNL3* is described in the schizophrenia section.

*RGS6*, regulator of G protein signaling 6, is a member of the RGS (regulator of G protein signaling) family. Proteins in the RGS family are defined by the presence of an RGS domain that confers the GTPase-activating activity of these proteins toward G alpha subunits. The RGS proteins have been associated with schizophrenia, cognitive function^65^, motor neuron diseases^193,194^, blood pressure^195^, resting heart rate^196^ , small cell lung carcinoma^197^, C-reactive protein level^184^ in GWAS.

*GATAD2A*, GATA zinc finger domain containing 2A, encodes a protein as a subunit of the methyl-CpG-binding protein-1 complex (MeCP1). MeCP1 deacetylates methylated nucleosomes to repress gene expression^198^. *GATAD2A* is associated with schizophrenia and thought to have pleiotropic effects on breast cancer risk^199^. It's also associated with type 2 diabetes^200^.

*RERE*, also known as arginine-glutamic acid dipeptide repeats, encodes a member of the atrophin family of arginine-glutamic acid (RE) dipeptide repeat-containing proteins. The *RERE* gene is a nuclear receptor coregulator that positively regulates retinoic acid signaling. *RERE* is known to be associated with neurodevelopmental disorder (OMIM: 616975), which is an autosomal dominant syndrome characterized by developmental delay, intellectual disability, and behavioral disorders, such as autism spectrum disorders^201^. *RERE* is also associated with a wide range of phenotypes including cognitive function^65,202^, neuroticism^203^, depression^79,204^, smoking^187^, well-being^89^, white blood cell count^68,144^, ophthalmology measure^205,206^ and myopia^207^, blood pressure^208^, heel bone mineral density^209^ and asthma^210,211^.

Little is known about *ZNF823*, zinc finger protein 823 and its gene function. It has been associated with metabolite levels^212^.

*GLYCTK*, glycerate kinase, encodes a member of the glycerate kinase type-2 family and encoded enzyme catalyzes the phosphorylation of (R)-glycerate. GLYCTK did not show genome-wide significant association in schizophrenia GWAS analyzed in this study. It is known to be associated with D-glyceric aciduria (glycerate kinase deficiency; OMIM: 220120). The rare autosomal recessive metabolic disorder has highly variable phenotypes from severe phenotypes like encephalopathy, severe mental retardation, seizures, microcephaly, to mild phenotypes such as mild speech delay or even normal development^213^.

*CLCN3*, chloride voltage-gated channel 3, did not show genome-wide significant association in schizophrenia GWAS analyzed in this study; however, it was associated in other schizophrenia GWAS^214^. *CLCN3* encodes a member of the voltage-gated chloride channel (ClC) family, which is present in all cell types and localized in plasma membranes and in intracellular vesicles. Besides schizophrenia, *CLCN3* is also associated with Parkinson's disease^158^.

*MAU2*, also known as MAU2 sister chromatid cohesion factor, is involved plays a role in sister chromatid cohesion, which is essential for segregation of homologous chromosomes during meiosis I and for repair of DNA double-strand breaks during G2 phase. *MAU2* did not show genome-wide significant association in schizophrenia GWAS analyzed in this study. *MAU2* is associated with bipolar disorder^215^ as well as lipid and blood biomarkers, such as triglycerides^67^, LDL^70^, plateletcrit and white blood cell count^68^.

We also evaluated 141 of the 160 schizophrenia associated eQTL instruments for an interaction effect with cell type proportions. In total, 21 eQTL instruments where significant ieQTLs, 1 with astrocytes (*TNFRSF13C*), 1 with endothelial cells (*MICB*), 5 with macrophages (*BTN3A2*, *C4A*, *HLA-DMB*, *RPS17* and *HLA-DQB1*), 7 with neurons (*CNNM2*, *TYW5*, *SNX19*, *PCDHA8*, *CPEB1*, *ENDOG* and *METTL21A*), 6 with oligodendrocytes (*COL11A2*, *PSORS1C1*, *ANKRD44*, *FOXN2*, *CORO7* and *SFTA2*) and 1 with astrocytes and neurons (*AS3MT*). Of these, 6 eQTL instruments passed the significance threshold only for MR (*BTN3A2*, *C4A*, *HLA-DMB*, *CNNM2*, *COL11A2* and *TYW5*) and 7 eQTL instruments passed the significance threshold only for colocalization (*RPS17*, *PCDHA8*, *FOXN2*, *CPEB1*, *CORO7*, *ENDOG* and *METTL21A*).

##### Years of schooling and cognitive function

There are 31 findings for cognitive function and 55 MR findings for years of schooling that passed colocalization (PP4>0.7, **Supplementary Table 12**). This is the first report of MR with brain cis-eQTL instruments for the cognitive function and years of schooling outcomes^65,83^. Lee *et al.* only reported brain eQTLs from GTEx, and there were only 2 loci that had eQTLs for *PITPNM2* at 12q24.31 and two genes at 22q13.1, *TAB1* and *MGAT3* (Table S16 in Lee *et al*^65^).

We investigated the pathways enriched for highly co-regulated genes of the findings that passed MR and colocalization using the *MetaBrain* GeneNetwork browser. The top HPO pathway enrichments for cognitive function were Cerebellar hypoplasia (p=4.7x10^-8^) and Aplasia/Hypoplasia of the cerebellum (p=6.8x10^-7^), suggesting an important role of the cerebellum in cognitive function. Cerebellum volume and function has long been linked to intelligence^216–218^, and three of the top HPO enrichment terms for intelligence were related to cerebellar hypoplasia or abnormalities in the cerebellum. The top HPO enrichment terms for years of schooling included Abnormal nervous system physiology (p=1.2x10^-5^), Abnormality of the nervous system (p=6.5x10^-5^) and Abnormal muscle fiber protein expression (p=1.0x10^-4^). The top two findings suggest that contributions from important neurodevelopmental genes could factor into the years of schooling trait. Interestingly, the third ranked pathway enrichment is a muscle phenotype, and motor delay is a common co-morbidity with intellectual disability/developmental delay^219^.

There are 8 genes with significant findings for both traits, *CYB561D1*, *KIFC2*, *KMT5A*, *NPIPB9*, *RECQL4*, *RHEBL1*, *SYPL2* and *TUFM*. All of these findings were significant in the years of schooling outcome, but *KIFC2* and *RECQL4* were just below the GWAS significance threshold for cognitive function. There is shared genetic basis between years of schooling and cognitive function, and they are genetically correlated with the cerebral cortical morphology^220^. Moreover, the genes common to these two traits may influence cognitive ability. The top two HPO enrichment terms were Cerebellar hypoplasia (p=3.0x10^-4^) and Aplasia/Hypoplasia of the cerebellum (p=1.1x10^-3^), underscoring the importance of these common set of 8 genes in cognitive ability.

We also evaluated 532 of the 578 years of schooling and cognitive function associated eQTL instruments for an interaction effect with cell type proportions. In total, 51 eQTL instruments where significant ieQTLs, 2 with astrocytes (*SNORC* and *EEF1AKMT2*), 6 with endothelial cells (*CCDC32*, *TIE1*, *ELOA3D*, *PRTG*, *ELOA3B* and *UPK1A*), 5 with macrophages (*BTN3A2*, *ERAP2*, *FCER1A*, *HIST1H3C* and *HLA-DRB5*), 22 with neurons (*RHEBL1*, *AKTIP*, *PCDHA8*, *AFF3*, *DHRS11*, *FAM180B*, *ZNF584*, *KIF1BP*, *POM121L2*, *CCDC65*, *SLC39A4*, *RMC1*, *RHEBL1*, *AFF3*, *TRANK1*, *FIBP*, *WDR92*, *CNNM2*, *AKTIP*, *SNX32*, *LY6D* and *DHRS11*), 11 with oligodendrocytes (*NPIPB9*, *TEX14*, *SH3BGR*, *SEPTIN10*, *NPIPB9*, *LRRIQ3*, *HAUS4*, *ZCWPW1*, *ELOVL1*, *PXK* and *AIF1L*), 3 with neurons and oligodendrocytes (*CEP192*, *PDE1A* and *CEP192*) and 2 with macrophages and oligodendrocytes (*ARL17B* and *ARL17B*). Of these, 10 eQTL instruments passed the significance threshold for both MR and colocalization (*NPIPB9*, *RHEBL1*, *FAM180B*, *NPIPB9*, *RMC1*, *RHEBL1*, *LRRIQ3*, *ZCWPW1*, *TRANK1* and *ELOA3D*), 12 eQTL instruments passed the significance threshold only for MR (*AKTIP*, *TEX14*, *PCDHA8*, *AFF3*, *DHRS11*, *ARL17B*, *AFF3*, *HAUS4*, *FCER1A*, *TIE1*, *ELOVL1* and *FIBP*) and 9 eQTL instruments passed the significance threshold only for colocalization (*CEP192*, *KIF1BP*, *SEPTIN10*, *PRTG*, *ELOA3B*, *EEF1AKMT2*, *CEP192*, *UPK1A* and *PXK*).

##### Brain volume

MR analysis was performed for several brain volume outcomes for amygdala, caudate, hippocampus, intercranial, nucleus accumbens, pallidum, putamen and thalamus^221^. There were 20 brain volume findings with significant Wald ratio findings, but only 3 intracranial volume and 2 putamen volume findings also passed colocalization. Downregulation of *KANSL1* (WR=-15,484, p=8.75x10^-8^), *SPPL2C* (WR=-51,071, p=4.38x10^-8^) and *STH* (WR=-66,867, p=3.79x10^-8^) increased intercranial volume. These genes and eQTL instruments are part of the *MAPT* extended haplotype, but there was no significant cis-eQTL in *MetaBrain* for *MAPT*. The H1 haplotype appeared to correspond with decreased brain volume, concordant with previous reports^221,222^. Upregulating *DCC* (WR=155, p=1.27x10^-7^) increased putamen volume. *DCC* is upregulated in the first two trimesters of pregnancy, suggesting that this gene could be important in prenatal brain development^223^. There was also a significant WR finding where decreased *DCC* expression related with years of schooling.

We also evaluated 18 of the 20 brain volume associated eQTL instruments for an interaction effect with cell type proportions. In total, 3 eQTL instruments where significant ieQTLs, 1 with endothelial cells (*MYLK2*), 1 with oligodendrocytes (*SEPTIN12*) and 1 with macrophages and oligodendrocytes (*ARL17B*). Of these, 1 eQTL instrument passed the significance threshold only for colocalization (*SEPTIN12*).

##### Mendelian disease overlap

Of the significant WR findings that pass colocalization, 15 gene-indication pairs overlap a gene annotated in the DDG2P database of Mendelian neurodevelopmental disorders, of which 6 overlapped with brain-related disease traits, including depression, multiple sclerosis and schizophrenia (**Supplementary Table 12**). Two findings are in the same direction of effect with Mendelian diseases, *PCCB* and *MAU2*. Loss of function mutations in *PCCB* cause autosomal recessive propionic acidemia^224^, a metabolic disorder caused by deficiency of propionyl-CoA carboxylase, causing accumulation of toxic compounds in the blood, including propionyl-CoA, propionic acid, ketones and ammonia^225^. Interestingly, this disorder is associated with visual hallucinations and psychosis, and the psychiatric symptoms last longer than the metabolic imbalance^226^, suggesting that metabolic imbalance arising from *PCCB* deficits could also be driving a neuropsychiatric clinical presentation, as suggested by the WR findings that decreased *PCCB* expression increases SCZ risk. We also found that decreased *MAU2* increases risk for SCZ, and loss of function mutations in this gene causes MAU2 neurodevelopmental disorder with Cornelia de Lange Syndrome clinical presentation^227^, in the same direction of effect. *MAU2* and *NIPBL*, the primary gene causing Cornelia de Lange Syndrome, form the cohesin loading complex and play an important role in initiating the binding of cohesin onto DNA. It has been suggested that decreased cohesion in the brain leads to defective synapse development and some neuropsychiatric presentations^228^, so neurodevelopmental and neuropsychiatric traits may both be affected by dysregulation of cohesion. Increasing *SLC12A5* expression is implicated in both depression and multiple sclerosis disease risk, although the Mendelian diseases caused by this gene, autosomal dominant epilepsy and autosomal recessive developmental and epileptic encephalopathy, are likely due to loss of function variants^99^. This suggests that *SLC12A5* is highly dosage sensitive. Similarly, increased expression of *RERE* appears to increase risk for schizophrenia, whereas this gene is in the critical region of 1p36 deletion syndrome^229^ and haploinsufficiency of *RERE* mimics deficits found in 1p36 deletion syndrome^201^.

### Systematic colocalization comparison of AD risk GWAS and cortex eQTLs

#### Method

For Alzheimer’s disease (AD), we first sought out to compare colocalization of *MetaBrain* Cortex-EUR *cis*-eQTLs with the latest AD GWAS findings by Schwartzentruber *et al.* in which smaller cortex or microglia eQTL datasets were used including brainseq^230^, ROSMAP^5^, xQTL-eQTL^5^, AMP-AD including CMC^231^ and primary microglia^139,232^. We examined 36 genome-wide significant AD loci excluding the *APOE* locus. Nine loci were found with multiple independent signals so conditional analysis was done using GCTA-COJO with the 1000 Genome European LD reference panel. Colocalization was then tested for all independent signals against primary Cortex-EUR cis-eQTLs. Our analysis was split into two parts: first, we aimed at replicating 781 pair-wise brain eQTL colocalizations examined in Schwartzentruber *et al.*^33^; second, we aimed at including all cis-eQTL protein-coding genes around AD GWAS loci additionally identified in *MetaBrain* Cortex-EUR, by extracting all significant *cis*-eQTLs (FDR<0.05) within 500Kb of the GWAS lead SNPs. The R package, coloc, was used to estimate the posterior probability of a shared causal variant between AD GWAS loci and Cortex-EUR *cis*-eQTLs. For each GWAS lead SNP, a +/- 500Kb region was constructed and colocalization was tested using the default priors.

#### Results

In total, 1,025 pairwise colocalizations were tested (**Supplementary Table 13**). Of which, 29 findings colocalized (PP4>0.7). We replicated (PP4>0.7) the majority of AD genes including *FCER1G*, *CR1*, *TREM2*, *CD2AP*, *CCDC6*, *TSPAN14*, *APH1B*, *KAT8*, *PRSS36*, *ZNF668*, *ACE*, and *CD33*. Noticeably we found *CASS4* cortex eQTL colocalization which was only reported in microglia by Schwartzentruber *et al.*^33^ In addition, we also identified several novel colocalization signals including *KLHDC9* at the *ADAMTS4* locus (PP4=0.77), *EPHA1* (PP4=0.71) and *TAS2R60* (PP4=0.97) at the *EPHA1* locus, and *EED* at the *PICALM* locus (PP4=0.87). We also didn’t find colocalization for several genes including *ABCA7*, *SLC39A13*, *SPPL2A* which were identified in Schwartzentruber *et al.*^33^ using ROSMAP data, although our results had much denser SNP coverage at these loci with much stronger eQTL signals. Further evidence is needed to interpret these loci and the potential causal genes. For *BIN1*, we didn’t find any significant cortex eQTL and previous colocalization evidence was also based on microglia data.

### Colocalization of top MR hits with opposite effect directionalities between *MetaBrain* and eQTLGen

We identified 1,192 top MR hits using *MetaBrain* Cortex-EUR (**Supplementary Table 12**) that passed the suggestive threshold (p<5x10^-5^). After comparing the eQTL effect between *MetaBrain* Cortex-EUR and eQTLgen for the MR instruments, we found 624 MR top hits of which the *MetaBrain* Cortex-EUR instruments were significant (p<0.05) in eQTLgen, but 146 MR hits showed allelic discordance between *MetaBrain* Cortex-EUR and eQTLgen. We then checked if the *MetaBrain* Cortex-EUR instruments are in high LD with the top eQTLgen eQTL for the allelic discordant MR hits and found 31 with *MetaBrain* instruments in high LD with the top eQTLGen eQTL (r^2^>0.8). (**Supplementary Table 15**) Colocalization was then checked for these 31 MR hits in both *MetaBrain* and eQTLgen to identify potentially shared causal genes between cortex and blood but with opposite directionalities for neurological traits. Colocalization methods were described in the Methods section of the main text. As a result, we found 11 MR top hits that colocalized with both *MetaBrain* Cortex-EUR and eQTLgen, and 5 of them also passed the MR Bonferroni correction: *CASS4* for AD, *TMEM170B* for intelligence, *GATAD2A* for SCZ and years of schooling and *ZCWPW1* for years of schooling (**Supplementary Figure 21**).

### MR comparison between *MetaBrain* and eQTLGen in multiple sclerosis

Multiple sclerosis (MS) is hypothesized to be largely mediated by the immune system as an outcome of an inflammatory insult to the central nervous system and peripheral tissues. As both blood and brain cell types could play an important role in the etiology of the disease, we compared the MR results using cis-eQTL instruments in both *MetaBrain* Cortex-EUR and eQTLgen blood tissue, primarily to assess whether or not there were significant MR findings that show discrepant directions of effect between the tissue types.

For each data set, we selected genome-wide significant eQTLs at p<5x10^-8^ cut-off and LD clumped to form a set of independent eQTLs for each study. The *MetaBrain* instruments consisted of 10,510 eQTLs across 8,949 genes and the eQTLGen instruments consisted of 41,157 eQTLs across 16,189 genes. We looked up the SNP effect in the MS GWAS for these instruments, harmonised the effects and performed MR to obtain a set of Wald ratios (WR) estimating the causal effect between gene expression and MS pertaining to the eQTL in each study. The WR measures the change in MS risk per unit change in gene expression through the effect allele of the instrumenting eQTL. A positive WR would indicate that increased gene expression is related to increased MS risk, whereas a negative WR would indicate that decreased gene expression is related to increased multiple sclerosis risk. Therefore, if the WRs agree between studies, then there is agreement on the direction of the expression effect acting on MS risk (i.e., whether promoting or inhibiting the gene would increase MS risk). In total, the *MetaBrain* WR set consisted of 9,392 WRs across 8,295 genes and the eQTLGen WR set consisted of 34,044 WRs across 16,567 genes.

We first assessed whether the MR results agreed between *MetaBrain* and eQTLGen for each gene, comparing the WRs on the different SNP instruments selected within each study. This represents the naïve analysis where, for example, a researcher tries to infer genes causally related to MS by conducting MR relying on eQTLs from blood only (i.e., blind to brain specific tissue effects). For genes which were instrumented by more than one SNP, we selected the top hit WR (lowest p-value) to do the comparison on, removing the trans-chromosomal eQTLGen instruments which were not on the same chromosome as the *MetaBrain* instruments. This reduced the *MetaBrain* WR set by 12% from 9,392 to 8,295 WRs and the eQTLGen instrument set by 82% from 34,044 to 5,919 WRs unique to the gene. We then compared the *MetaBrain* instruments against eQTLGen and found that the proportion of the WR effects which agreed to be very low: 2,291 (38.7%) of the 5,919 shared instrumented genes showing WRs with opposite direction between the studies (**Supplementary Table 14A, Supplementary Figure 23, top panel**). We observed no major change in the WR agreement among genes which showed robust association with multiple sclerosis (at WR p-value <5x10^-5^). Within the 103 genes associated in *MetaBrain* 36 genes (35.0%) had an opposite WR to eQTLGen and within the 152 genes associated in eQTLGen 56 genes (36.8%) had an opposite WR to *MetaBrain*.

Taking a closer look at the 157 top MR hits (**Supplementary Table 12**) identified for MS in *MetaBrain* Cortex-EUR, we identified a total of 28 genes that did not have a significant *cis*-eQTL in blood tissue, of which 6 genes (*SLC12A5*, *CCDC155*, *MYNN*, *HIST1H1D*, *PIGW* and *ATG16L2*) also passed colocalization in Cortex-EUR (**Supplementary Table 16**). Interestingly, we also identified 32 genes where MR in Cortex-EUR and eQTLGen showed both suggestive signal (p<5x10^-5^) but opposite directionalities suggesting potentially tissue-dependent putative causal genes. Furthermore, two of them (*KMT5A* and *RNF19B*) also passed colocalization in Cortex-EUR, although neither passed colocalization in eQTLGen. The lack of colocalization signals in eQTLGen, however, can be due to the violation of single causal variant in the region (**Supplementary Figure 22**). Conditional eQTLs are not available from eQTLGen to adequately address this issue.

We then compared MR results on the same SNP instrument to account for the discrepant WRs that could be due to the *MetaBrain* and eQTLGen eQTLs not being in LD (i.e., the instruments selected for each study are not sharing the same causal variant in the region). In the previous comparison, only 178 (3%) of the WRs had been derived from the same SNP instrument. This would represent the scenario where, for example, a researcher is able to incorporate information on brain tissue eQTL effects to improve specificity of blood eQTL based MR analysis to identify the causal genes in MS. To do the comparison we looked up the full *MetaBrain* instrument set (all 9,392 eQTLs) in the eQTLGen study and re-performed the MR between these eQTLGen effects and MS. We found a shared instrument for 7,274 (77.4%) of the 9,932 *MetaBrain* eQTLs within eQTLGen which we could conduct the MR on. As the eQTLs from the lookup in eQTLGen will not always be significant at the p<5x10^-8^ level this could result in poorly calibrated standard errors for the WR. Therefore, to account for the standard error in the eQTL instrument, we computed the standard errors in the WRs with the second term approximation in the Taylor expansion included (see section: Taylor expansion for the Wald ratio standard error). Although fixing on the same SNP instrument improved the WR agreement between the studies, the discordancy remained relatively high (**Supplementary Table 14B**): 1,891 (26%) from the 7,274 WRs showed an opposite direction of effect. However, in contrast to the previous comparison done on the same genes, the WR agreement did improve for genes associated with multiple sclerosis (WR p-value <5x10^-5^) when also restricted to the same instruments. Within the 124 genes associated in *MetaBrain* 19 genes (15.3%) had opposite WRs to eQTLGen and within the 75 genes associated in eQTLGen 8 genes (10.6%) had opposite WRs to the *MetaBrain* study.

Due to our WR comparison being limited to the set of eQTL instruments which intersect with the MS GWAS only, we also performed a comparison on the eQTL effect sizes across all the instruments available. As we anticipated, the findings from this analysis were consistent with the MR comparison with a considerable proportion of the eQTL effects showing opposing expression effect directions. We found that 7,986 of the 10,510 *MetaBrain* instruments (76.0%) were present in eQTLGen, of which 2,066 eQTLs (25.9%) had opposite direction of expression effect. Of which, 5,269 (66.0%) of these eQTLs reached p<5x10^-8^ cut-off used to select instruments, and 1,019 eQTLs (19.3%) had opposing expression effect in *MetaBrain*. In the reverse lookup, 23,968 of the 41,157 eQTLGen instruments (58.2%) were present in *MetaBrain*, of which 7,826 eQTLs (32.7%) had opposing expression effect. 10,987 (45.8%) of these eQTLs reached p<5x10^-8^, of which 2,432 (22.1%) showed opposing expression effect in eQTLGen.

Of the 135 genes with MR findings in Cortex-EUR for MS, we identified 28 genes without a significant eQTLGen instrument, including 3 genes (*SLC12A5, CCDC155* and *MYNN*) for which we found both MR significance and colocalization in *MetaBrain* (**Supplementary Table 16**). For 25 MS genes, we were able to compare gene expression levels in blood and different brain regions using GTEx samples. We downloaded median gene expression from <https://gtexportal.org/home/datasets> (GTEx_Analysis_2017-06-05_v8_RNASeQCv1.1.9_gene_median_tpm.gct.gz). The majority (n=16; 64%) had almost no expression in blood, including SLC12A5, 3 (12%) had lower expression in blood than in cortical tissues, 2 (8%) had comparable expression in blood and cortical tissues (CCDC155 and MYNN) and 4 (16%) had higher expression in blood than in cortical tissues (**Supplementary Figure 24**).

### Taylor expansion for the Wald ratio standard error

To account for the error in both the instrument-exposure and instrument-outcome when comparing WR on the same SNP instrument we used the two term Taylor expansion for computing the standard error (SE) of the WR. Let WR_XY_ denote the Wald ratio between the exposure (X, eQTL) and outcome (Y, multiple sclerosis) for a SNP instrument. Let B_XY_ and SE_XY_ denote the effect size and standard error of the eQTL effect and B_ZY_ and SE_ZY_ denote the same for the outcome SNP effect.

For the primary MR analysis in our study, we used the first term expansion only which accounts for the variance in the outcome SNP effect. Note: the contribution of the eQTL error to the WR SE will be almost negligible as highly significant eQTLs are selected (p<5x10^-8^ in this study).

In the first term approximation, the Wald ratio is estimated as:

$${WR}_{XY}= \frac{\beta_{ZY}}{\beta_{ZX}}$$

with variance:

$${Var\left( {WR}_{XY} \right)}_{1}= \frac{{{SE}_{ZY}}^{2}}{{\beta_{ZX}}^{2}}$$

However, we can also expand the series by another term to account for the variance in the instrument-exposure (eQTL) effect:

$${Var\left( {WR}_{XY} \right)}_{2}= {Var\left( {WR}_{XY} \right)}_{1}+ \frac{{\beta_{ZY}}^{2}{{SE}_{ZX}}^{2}}{{\beta_{ZX}}^{4}}$$

and obtain the standard error for this two-term expansion:

$${SE\left( {WR}_{XY} \right)}_{2}= \sqrt{{Var\left( {WR}_{XY} \right)}_{2}}$$

### Removing cross-mapping artefacts from *trans*-eQTL results

When a *trans*-eQTL gene has similar paralogous genes in the close proximity (<5Mb) of a given eQTL SNP, the apparent *trans*-eQTL effect may actually reflect a much stronger *cis*-eQTL effect, which might be caused by reads mapping to multiple positions in the genome^8^. While this should be corrected by not counting the RNA-seq reads assigned to multiple genomic features, there might still be some non-detected cases.

To remove such false positive *trans*-eQTLs, we created sets of 35bp "reads" from the human reference genome (GENCODE^22^ v32) for each significant (FDR<0.05) *trans*-eQTL gene. To span the gene sequence, we used a shifting window approach, with each consecutive window shifting 2bp, while also generating reads spanning exon-exon boundaries. We then created 10Mb sequences centered around each significant *trans*-eQTL SNP. Then, we mapped the reads generated for the gene to the 10Mb SNP region using BWA-mem^233^ v0.7.15 for each *trans*-eQTL SNP-gene pair. We note that we did not explicitly require the reads to map to genes in order to account for potentially unannotated genes and pseudogenes in the genome. Finally, for each *trans*-eQTL, we determined the number of base pairs mapped, and divided this by the number of base pairs generated for the gene, resulting in a proportion of the gene mapped within 5Mb of the SNP.

As *trans*-eQTLs with high proportions of genes mapping within the SNP region are more likely actually *cis*-eQTL effects, we opted to use 5% of gene mapping to the vicinity of SNP as a threshold to declare *trans*-eQTL to be potentially caused by cross-mapping. To correct the multiple testing threshold for the identified potential false positive *trans*-eQTLs, we then repeated the FDR estimation, leaving out the identified cross-mapping *trans*-eQTLs.

### *Trans*-eQTLs in the 7p21.3 locus

2,252 (87%) of the observed *trans*-eQTL genes were affected by a set of 45 variants that were located at 7p21.3 (**Figure 6A**). This locus is associated with many traits, including brain-related phenotypes, such as frontotemporal dementia and major depressive disorder (**Supplementary Table 17**). The majority (81%) of these *trans*-eQTLs were associated with two SNPs, rs11974335 and rs10950398 (R^2^=0.98), which are both located in an intron of *TMEM106B* and were associated with 1,025 and 790 genes, respectively (**Figure 6C**). While we did not observe a *TMEM106B* *cis*-eQTL for any variant in the locus, we did observe *cis*-eQTLs for 22 other genes (**Supplementary Table 17**). Of note, this included rs1990622 (R^2^=0.96 with rs11974335), an index variant for frontotemporal lobar degeneration^234^ (FTLD) located downstream of *TMEM106B*, which had a *cis*-eQTL on *THSD7A* (SNP–TSS distance >411Kb; **Figure 6C**). The FTLD risk allele rs1990622-A decreased expression of this gene, which is suggested to be involved in neuro-angiogenesis^235^ (**Supplementary Figure 26A**). rs1990622 was associated with 35 trans-eQTL genes, of which 31 were negatively regulated by the FTLD risk allele (**Supplementary Figure 25**). Downregulated genes included genes involved in calcium transport such as *CALB2*, and *CBLN1*, and potassium transport, such as *KCND3*, *KCHN5* and *KCDT2* (**Figure 6C, Supplementary Figure 26A**). Collectively, upregulated *trans*-eQTL genes for rs11974335, rs10950398 and rs1990622 were enriched for neuron related processes, such as synaptic signaling (p=1.3x10^-28^) and nervous system development (p=2.9x10^-21^), while downregulated genes were enriched for gliogenesis (p=1.6x10^-8^) and oligodendrocyte differentiation (p=3.1x10^-6^; **Supplementary Table 21**).

While rs11974335 and rs10950398, rs1990622 are in high LD (R^2^>0.9), we observed only a limited number of *trans*-eQTL genes for rs1990622. rs11974335 and rs10950398 were tested in only 6 datasets with WGS derived genotypes, while rs1990622 was tested in 20 datasets, suggesting that heterogeneity between datasets decreased the number of significant *trans*-eQTLs for rs1990622 (**Figure 6C**). Indeed, *trans*-eQTLs for rs1990622 were highly heterogeneous across the included datasets, being most pronounced in the AMP-AD and UCLA_ASD datasets, and less so in other datasets of comparable sample size, such as CMC (**Supplementary Figure 25**). As another potential source for heterogeneity, we considered that many of the observed *trans*-eQTLs might be driven by differences in cell type proportions. We therefore associated the five predicted cell type proportions with genotypes. We observed 52 associations from 35 SNPs in Cortex-EUR samples, and 67 associations from 49 SNPs when AFR datasets were included, but ENA excluded (FDR<0.05; **Supplementary Table 18**). Respectively 31 (Cortex-EUR) and 11 SNPs (Cortex-EUR+AFR) were located in in the 7p21.3 locus. In the Cortex-EUR samples, this included rs1990622, which showed a decrease in neuron proportions for the FTLD risk allele, matching previous reports (**Supplementary Figure 26A**)^236,237^. This SNP showed a similar association when AFR samples were included, but was not significant (FDR=0.08). Like the observed *trans*-eQTLs, these neuron proportion associations were most pronounced in the AMP-AD datasets. However, when comparing average neuron proportions in AMP-AD with those in other datasets, we did not observe many significant differences (**Supplementary Figure 26A**; **Supplementary Table 19**). Nevertheless, we observed a strong relationship between the *trans*-eQTL Z-scores and the correlation of *trans*-eQTL gene expression levels and neuron proportions (R^2^>0.85; **Supplementary Figure 26B; Supplementary Table 20**). Comparing Alzheimer cases versus controls in the AMP-AD dataset, we observed that neuron proportions were significantly lower (**Supplementary Figure 24**) and that *trans*-eQTL Z-scores were higher (**Supplementary Figure 26C**). This indicates that the observed *trans*-eQTLs and neuron associations may be in part driven Alzheimer dependent loss of neurons, driven by the 7p21.3 locus. While our analysis cannot disentangle the neuron loss from the Alzheimer association at rs1990622, it does suggest that this locus affects many downstream genes in Alzheimer patients.

### Comparison of predicted cell count proportions between AD patients and neurological controls

To assign Alzheimer’s disease status for AMP-AD samples we used either CERAD or cogdx score, whichever was available. For cogdx, individuals with a score of 4 or 5 were assigned Alzheimer’s disease status, and individuals with a score of 1 as non-neurological control. For CERAD, individuals with a score of 3 were assigned Alzheimer’s disease status, and individuals with score of 1 as non-neurological control. Individuals with other gogdx or CERAD scores were not included in this comparison.

### Gene prioritization using Downstreamer

#### Overview of Downstreamer methodology

In short, Downstreamer associates a gene level prioritization score (GWAS gene Z-scores) to a gene-gene co-regulation matrix to find genes that have many connections at the expression level to genes inside GWAS loci (core genes). In addition, Downstreamer can identify pathway enrichments by switching the co-regulation matrix for pathway annotations. Downstreamer implements a strategy that can do these associations while accounting for linkage disequilibrium (LD) structure and chromosomal organization. Downstreamer operates in two steps: In the first step, the GWAS gene Z-scores are calculated for the GWAS trait and a null distribution. In the second step, the GWAS gene Z-scores are associated with the phenotypes outlined above. The details on these steps are outlined in the sections below.

#### Downstreamer step 1: Calculation of GWAS gene Z-scores

The primary step in the Downstreamer converts the GWAS summary statistics from p-values per variant to an aggregate p-value per gene (gene p-value) while accounting for local LD structure. This aggregate gene level p-value represents the GWAS signal potentially attributable to that gene.

First, we applied genomic control to correct for inflation in the GWAS signal. We then integrated the procedure from the PASCAL^238^ method to Downstreamer so that we can aggregate variant p-values into a gene p-value while accounting for the LD structure. We aggregated all variants within a 25Kb window around the start and end of a gene using the non-Finnish European samples of the 1000 Genomes (1000G) Project Phase 3 to calculate LD^239^. We calculated these GWAS gene p-values for all 20,327 protein-coding genes (Ensembl^21^ release v75). The gene p-values were then converted to Z-scores for use in subsequent analysis. These are referred to as GWAS gene Z-scores.

To account for the long-range effects of haplotype structure which results in genes getting a similar GWAS gene Z-score, we use a generalized least squares (GLS) regression model for all regressions done in Downstreamer. The GLS model takes a correlation matrix that models this gene-gene correlation.

To calculate this correlation matrix, we first simulated 10,000 random phenotypes by drawing phenotypes from a normal distribution and then associating them to the genotypes of the 1000G Phase 3 non-Finnish European samples. Here, we only use the overlapping variants between the real traits and the permuted GWASs to avoid biases introduced by genotyping platforms or imputation. We then calculate the GWAS gene Z-scores for each of the 10,000 simulated GWAS signals as described above. Next, we calculate the Pearson correlations between the GWAS gene Z-scores. As simulated GWAS signals are random and independent of each other, any remaining correlation between GWAS gene Z-scores reflects the underlying LD patterns and chromosomal organization of genes.

We simulated an additional 10,000 GWASs as described above to empirically determine enrichment p-values, and finally, we used an additional 100 simulations to estimate the false discovery rate (FDR) of Downstreamer associations.

#### Downstreamer step 2: Association of GWAS Z-scores with phenotypes

##### Pre-processing of GWAS gene Z-scores and pruning of highly correlated genes

For each GWAS, both real and simulated, the GWAS Z-scores were re-scaled to fit a normal distribution to ensure that outliers would not have disproportionate weights. Limitations in the PASCAL methodology result in ties at the minimum significance level of 1x10^-12^ for highly significant genes, so we use the minimum SNP p-value from the GWAS to identify the most significant gene and resolve the tie. We then used a linear model to correct for gene length, as longer genes will typically harbor more SNPs.

Sometimes, two (or more) genes will be so close to one another that their GWAS gene Z-scores are highly correlated, violating the assumptions of the linear model. Thus, genes with a Pearson correlation r ≥ 0.8 in the 10,000 GWAS permutations were collapsed into 'meta-genes' and treated as one gene. Meta-gene Z-scores were averaged across the input Z-scores. The GWAS Z-scores of the meta genes were scaled (mean=0, standard deviation=1).

##### Generalized least squares model to calculate pathway enrichment and core gene scores

We used a GLS regression to associate the GWAS gene Z-scores to the pathway Z-scores and co-regulation Z-scores. These two analyses result in the pathway enrichments and core gene prioritizations, respectively. We used the gene-gene correlation matrix derived from the 10,000 permutations as a measure of conditional covariance of the error term (𝛀) in the GLS to account for the relationships between genes due to LD and proximity. The pseudo-inverse of 𝛀 is used as a substitute for 𝛀-1

The formula of the GLS is as follows:

β=(XTΩ-1X)-1XTΩ-1y

Where β is the estimated effect size of pathway, term or gene from the co-regulation matrix, Ω is the gene-gene correlation matrix, X is the design matrix of real GWAS Z-scores and y is the vector of gene Z-scores per pathway, term or gene from the co-regulation matrix. As we standardized the predictors, we did not include an intercept in the design matrix and X only contains one column with the real GWAS gene Z-scores. We estimated the betas for the 10,000 random GWASs in the same way and subsequently used them to estimate the empirical p-value for β.

##### Pathway and gene set gene prediction

To identify pathway and disease enrichments, we used the following databases: Human Phenotype Ontology (HPO)^240^, Kyoto Encyclopedia of Genes and Genomes (KEGG)^241^, Reactome^242^ and Gene Ontology^243^ (GO) Biological Process, Cellular Component and Molecular Function. We have previously predicted how much each gene contributes to these gene sets, resulting in a Z-score per pathway or term per gene^19^. In a parallel step, genes were collapsed into meta-genes to ensure compatibility with the GWAS gene Z-scores, following the same procedure as in the GWAS pre-processing. Meta-gene Z-scores were calculated as the Z-score sum divided by the square root of the number of genes. Finally, all pathway Z-scores were scaled (mean=0, standard deviation=1).

##### Co-regulation matrix

To calculate core scores, we used a previously generated co-regulation matrix that is based on a large multi-tissue gene network^19^. This network was generated using the publicly available RNA-seq samples that were downloaded from the European Nucleotide Archive (https://www.ebi.ac.uk/ena). After QC, 56,435 genes and 31,499 samples covering a wide range of human cell-types and tissues remained. We performed a PCA on this dataset and selected 165 components representing 50% of the variation that offered the best prediction of gene function. We then selected the protein coding genes and centered and scaled the eigenvectors for these 165 components (mean=0, standard deviation=1) such that each component was given equal weight. The first components mostly describe tissue differences^244^, so this normalization ensures that tissue-specific-patterns do not disproportionately drive the co-regulation matrix. The co-regulation matrix is defined as the Pearson correlation between the genes from the scaled eigenvector matrix. The diagonal of the co-regulation matrix was set to zero to eliminate the disproportionate effect of the gene to its gene p-value. Pearson r values were converted to Z-scores.

### Downstreamer analysis in schizophrenia

The Downstreamer analysis, that uses co-expression to identify genes that co-expressed with genes in GWAS loci, prioritized 184 Bonferroni significant (p-value ≤ 2.55×10^-6^) genes for Schizophrenia (**Supplementary Table 23**). Ten of these prioritized genes are located within 250kb of a genome wide significant GWAS hit, the remaining 174 genes are not directly identified by GWAS. An enrichment analysis revealed that 57 (31%) of these prioritized genes are known to cause Mendelian forms of Intellectual disability and/or global developmental delay (Enrichment p-values: HP:0001249: 3.76×10^-17^ and HP:0001263: 1.58×10^-14^) (**Supplementary Table 23**). Another interesting enrichment was for the chromatin organization Reactome pathway, 26 (14%) of the prioritized genes are annotated to this pathway (p-value: 1.89×10^-20^). Of these 26 chromatin organization genes there were 18 (69%) that are overlapping with Intellectual disability and global developmental delay genes. For rare neurodevelopmental disorders it is known that the causative gene is often involved in chromatin organization^245^. For schizophrenia there are also already indications of the importance of chromatin modeling, for instance the childhood onset of schizophrenia that is caused by a damaging variant in the chromatin remodeling *CHD2* gene^246^. Here it is worthwhile to mention *CHD2* is one of the 184 genes that we predict to be important for schizophrenia. The overlap of our prioritized genes with neurodevelopmental disorders fits the hypothesis that schizophrenia is, or partly is, a neurodevelopmental disorder^247^ and implicates altered chromatin organization as one of the causative mechanisms for schizophrenia.

### Software

R
python
Java

**R packages**
ggrepel^248^,ggplot^249^,ggpubr^250^,viridis^251^,lattice^252^,gridextra^253^,data.table^254^,dplyr^255^,readxl^256^,scales^257^,GGally^258^,edgeR^259^,ggExtra^260^,gtable^261^,matrixStats^262^,naniar^263^,plyr^264^,reshape2^265^,stringr^266^,tidyr^267^,tidyverse^268^,topGO^269^,g:Profiler^16^

**Python packages**
pandas^28^**,** seaborn^270^**,** matplotlib^271^**,** scipy^272^**,** numpy^273^**,** statsmodels**.**api^274^**,** tabix^275,276^**,** sklearn^277^**,**upsetplot^278,279^**,**wget^280^**,**beautifultable^281^**,**sqlite3^282^

### References

1.	Tsui, B., Dow, M., Skola, D. & Carter, H. Extracting allelic read counts from 250,000 human sequencing runs in Sequence Read Archive. *bioRxiv* 386441 (2018) doi:10.1101/386441.

2.	Deelen, P. *et al.* Calling genotypes from public RNA-sequencing data enables identification of genetic variants that affect gene-expression levels. *Genome Medicine* **7**, 30 (2015).

3.	McKenna, A. *et al.* The Genome Analysis Toolkit: A MapReduce framework for analyzing next-generation DNA sequencing data. *Genome Res.* **20**, 1297–1303 (2010).

4.	Danecek, P. *et al.* The variant call format and VCFtools. *Bioinformatics* **27**, 2156–2158 (2011).

5.	Ng, B. *et al.* An xQTL map integrates the genetic architecture of the human brain’s transcriptome and epigenome. *Nat Neurosci* **20**, 1418–1426 (2017).

6.	Sieberts, S. K. *et al.* Large eQTL meta-analysis reveals differing patterns between cerebral cortical and cerebellar brain regions. *bioRxiv* 638544 (2019) doi:10.1101/638544.

7.	Allen, M. *et al.* Association of MAPT haplotypes with Alzheimer’s disease risk and MAPT brain gene expression levels. *Alzheimer’s Research & Therapy* **6**, 39 (2014).

8.	Võsa, U. *et al.* Unraveling the polygenic architecture of complex traits using blood eQTL metaanalysis. *bioRxiv* 447367 (2018) doi:10.1101/447367.

9.	Dobbyn, A. *et al.* Landscape of Conditional eQTL in Dorsolateral Prefrontal Cortex and Co-localization with Schizophrenia GWAS. *Am J Hum Genet* **102**, 1169–1184 (2018).

10.	Wingender, E., Dietze, P., Karas, H. & Knüppel, R. TRANSFAC: a database on transcription factors and their DNA binding sites. *Nucleic Acids Res* **24**, 238–241 (1996).

11.	Võsa, U. *et al.* Unraveling the polygenic architecture of complex traits using blood eQTL metaanalysis. *bioRxiv* 447367 (2018) doi:10.1101/447367.

12.	Genetic effects on gene expression across human tissues. *Nature* **550**, 204–213 (2017).

13.	Shang, L. *et al.* Genetic Architecture of Gene Expression in European and African Americans: An eQTL Mapping Study in GENOA. *The American Journal of Human Genetics* **106**, 496–512 (2020).

14.	Consortium, T. Gte. The GTEx Consortium atlas of genetic regulatory effects across human tissues. *Science* **369**, 1318–1330 (2020).

15.	Fu, J. *et al.* Unraveling the Regulatory Mechanisms Underlying Tissue-Dependent Genetic Variation of Gene Expression. *PLOS Genetics* **8**, e1002431 (2012).

16.	Raudvere, U. *et al.* g:Profiler: a web server for functional enrichment analysis and conversions of gene lists (2019 update). *Nucleic Acids Research* **47**, W191–W198 (2019).

17.	Stelzer, G. *et al.* The GeneCards Suite: From Gene Data Mining to Disease Genome Sequence Analyses. *Current Protocols in Bioinformatics* **54**, 1.30.1-1.30.33 (2016).

18.	Bray, N. L., Pimentel, H., Melsted, P. & Pachter, L. Near-optimal probabilistic RNA-seq quantification. *Nat. Biotechnol.* **34**, 525–527 (2016).

19.	Deelen, P. *et al.* Improving the diagnostic yield of exome- sequencing by predicting gene–phenotype associations using large-scale gene expression analysis. *Nat Commun* **10**, 1–13 (2019).

20.	Li, H. *et al.* The Sequence Alignment/Map format and SAMtools. *Bioinformatics* **25**, 2078–2079 (2009).

21.	Yates, A. D. *et al.* Ensembl 2020. *Nucleic Acids Res* **48**, D682–D688 (2020).

22.	Frankish, A. *et al.* GENCODE reference annotation for the human and mouse genomes. *Nucleic Acids Res* **47**, D766–D773 (2019).

23.	Love, M. I., Huber, W. & Anders, S. Moderated estimation of fold change and dispersion for RNA-seq data with DESeq2. *Genome Biology* **15**, 550 (2014).

24.	gene2phenotype. https://www.ebi.ac.uk/gene2phenotype/downloads.

25.	Plenge, R. M. Priority index for human genetics and drug discovery. *Nat Genet* **51**, 1073–1075 (2019).

26.	Hanauer, M. *Orphanet/Orphadata_aggregated*. (2021).

27.	Blech, M. *martinblech/xmltodict*. (2021).

28.	McKinney, W. Data Structures for Statistical Computing in Python. *Proceedings of the 9th Python in Science Conference* 56–61 (2010) doi:10.25080/Majora-92bf1922-00a.

29.	Brouwers, N. *et al.* Alzheimer risk associated with a copy number variation in the complement receptor 1 increasing C3b/C4b binding sites. *Molecular psychiatry* **17**, 223–33 (2012).

30.	Kucukkilic, E. *et al.* Complement receptor 1 gene (CR1) intragenic duplication and risk of Alzheimer’s disease. *Human genetics* **137**, 305–314 (2018).

31.	Dunkelberger, J. R. & Song, W. C. Complement and its role in innate and adaptive immune responses. *Cell research* **20**, 34–50 (2010).

32.	Maier, M. *et al.* Complement C3 deficiency leads to accelerated amyloid beta plaque deposition and neurodegeneration and modulation of the microglia/macrophage phenotype in amyloid precursor protein transgenic mice. *The Journal of neuroscience : the official journal of the Society for Neuroscience* **28**, 6333–41 (2008).

33.	Schwartzentruber, J. *et al.* Genome-wide meta-analysis, fine-mapping and integrative prioritization implicate new Alzheimer’s disease risk genes. *Nature Genetics* 1–11 (2021) doi:10.1038/s41588-020-00776-w.

34.	Matthews, A. L. *et al.* Regulation of Leukocytes by TspanC8 Tetraspanins and the ‘Molecular Scissor’ ADAM10. *Frontiers in immunology* **9**, 1451 (2018).

35.	Jouannet, S. *et al.* TspanC8 tetraspanins differentially regulate the cleavage of ADAM10 substrates, Notch activation and ADAM10 membrane compartmentalization. *Cellular and molecular life sciences : CMLS* **73**, 1895–915 (2016).

36.	Suh, J. *et al.* ADAM10 missense mutations potentiate β-amyloid accumulation by impairing prodomain chaperone function. *Neuron* **80**, 385–401 (2013).

37.	Ulland, T. K. & Colonna, M. TREM2 - a key player in microglial biology and Alzheimer disease. *Nature reviews. Neurology* **14**, 667–675 (2018).

38.	Li, Q. *et al.* Developmental Heterogeneity of Microglia and Brain Myeloid Cells Revealed by Deep Single-Cell RNA Sequencing. *Neuron* **101**, 207-223.e10 (2019).

39.	Schlepckow, K. *et al.* An Alzheimer-associated TREM2 variant occurs at the ADAM cleavage site and affects shedding and phagocytic function. *EMBO molecular medicine* **9**, 1356–1365 (2017).

40.	Thornton, P. *et al.* TREM2 shedding by cleavage at the H157-S158 bond is accelerated for the Alzheimer’s disease-associated H157Y variant. *EMBO Mol Med* **9**, 1366–1378 (2017).

41.	Bernstein, K. E. *et al.* A modern understanding of the traditional and nontraditional biological functions of angiotensin-converting enzyme. *Pharmacological reviews* **65**, 1–46 (2013).

42.	Zou, K. *et al.* Angiotensin-converting enzyme converts amyloid beta-protein 1-42 (Abeta(1-42)) to Abeta(1-40), and its inhibition enhances brain Abeta deposition. *The Journal of neuroscience : the official journal of the Society for Neuroscience* **27**, 8628–35 (2007).

43.	Liu, S. *et al.* A clinical dose of angiotensin-converting enzyme (ACE) inhibitor and heterozygous ACE deletion exacerbate Alzheimer’s disease pathology in mice. *The Journal of biological chemistry* **294**, 9760–9770 (2019).

44.	Quitterer, U. & AbdAlla, S. Improvements of symptoms of Alzheimer`s disease by inhibition of the angiotensin system. *Pharmacological research* **154**, 104230 (2020).

45.	Ding, J. *et al.* Antihypertensive medications and risk for incident dementia and Alzheimer’s disease: a meta-analysis of individual participant data from prospective cohort studies. *The Lancet. Neurology* **19**, 61–70 (2020).

46.	Eckman, E. A. *et al.* Regulation of steady-state beta-amyloid levels in the brain by neprilysin and endothelin-converting enzyme but not angiotensin-converting enzyme. *The Journal of biological chemistry* **281**, 30471–8 (2006).

47.	Hemming, M. L., Selkoe, D. J. & Farris, W. Effects of prolonged angiotensin-converting enzyme inhibitor treatment on amyloid beta-protein metabolism in mouse models of Alzheimer disease. *Neurobiology of disease* **26**, 273–81 (2007).

48.	Cuddy, L. K. *et al.* Aβ-accelerated neurodegeneration caused by Alzheimer’s-associated ACE variant R1279Q is rescued by angiotensin system inhibition in mice. *Science translational medicine* **12**, (2020).

49.	Serneels, L. *et al.* gamma-Secretase heterogeneity in the Aph1 subunit: relevance for Alzheimer’s disease. *Science (New York, N.Y.)* **324**, 639–42 (2009).

50.	Zhang, X. *et al.* Negative evidence for a role of APH1B T27I variant in Alzheimer’s disease. *Human molecular genetics* **29**, 955–966 (2020).

51.	Lambert, J. C. *et al.* Meta-analysis of 74,046 individuals identifies 11 new susceptibility loci for Alzheimer’s disease. *Nature genetics* **45**, 1452–8 (2013).

52.	Beecham, G. W. *et al.* Genome-wide association meta-analysis of neuropathologic features of Alzheimer’s disease and related dementias. *PLoS genetics* **10**, e1004606 (2014).

53.	Laxmi, A., Gupta, P. & Gupta, J. CCDC6, a gene product in fusion with different protoncogenes, as a potential chemotherapeutic target. *Cancer biomarkers : section A of Disease markers* **24**, 383–393 (2019).

54.	Demontis, D. *et al.* Discovery of the first genome-wide significant risk loci for attention deficit/hyperactivity disorder. *Nat Genet* **51**, 63–75 (2019).

55.	Demircioglu, F. E., Burkhardt, P. & Fasshauer, D. The SM protein Sly1 accelerates assembly of the ER-Golgi SNARE complex. *Proc Natl Acad Sci U S A* **111**, 13828–33 (2014).

56.	Burgoyne, R. D. & Morgan, A. Chaperoning the SNAREs: a role in preventing neurodegeneration? *Nat Cell Biol* **13**, 8–9 (2011).

57.	Brooks, W. S., Banerjee, S. & Crawford, D. F. G2E3 is a nucleo-cytoplasmic shuttling protein with DNA damage responsive localization. *Exp Cell Res* **313**, 665–76 (2007).

58.	Rafiullah, R. *et al.* Homozygous missense mutation in the LMAN2L gene segregates with intellectual disability in a large consanguineous Pakistani family. *Journal of Medical Genetics* **53**, 138–144 (2016).

59.	Lim, C. H. *et al.* Genetic association of LMAN2L gene in schizophrenia and bipolar disorder and its interaction with ANK3 gene polymorphism. *Prog Neuropsychopharmacol Biol Psychiatry* **54**, 157–162 (2014).

60.	Tsai, R. Y. L. & McKay, R. D. G. A nucleolar mechanism controlling cell proliferation in stem cells and cancer cells. *Genes Dev* **16**, 2991–3003 (2002).

61.	Goes, F. S. *et al.* Genome-wide association of mood-incongruent psychotic bipolar disorder. *Transl Psychiatry* **2**, e180 (2012).

62.	Styrkarsdottir, U. *et al.* Meta-analysis of Icelandic and UK data sets identifies missense variants in SMO, IL11, COL11A1 and 13 more new loci associated with osteoarthritis. *Nat Genet* **50**, 1681–1687 (2018).

63.	Southam, L. *et al.* Whole genome sequencing and imputation in isolated populations identify genetic associations with medically-relevant complex traits. *Nat Commun* **8**, 15606 (2017).

64.	Kettunen, J. *et al.* Genome-wide study for circulating metabolites identifies 62 loci and reveals novel systemic effects of LPA. *Nat Commun* **7**, 11122 (2016).

65.	Lee, J. J. *et al.* Gene discovery and polygenic prediction from a genome-wide association study of educational attainment in 1.1 million individuals. *Nat Genet* **50**, 1112–1121 (2018).

66.	Kathiresan, S. *et al.* Six new loci associated with blood low-density lipoprotein cholesterol, high-density lipoprotein cholesterol or triglycerides in humans. *Nat Genet* **40**, 189–197 (2008).

67.	Wojcik, G. L. *et al.* Genetic analyses of diverse populations improves discovery for complex traits. *Nature* **570**, 514–518 (2019).

68.	Astle, W. J. *et al.* The Allelic Landscape of Human Blood Cell Trait Variation and Links to Common Complex Disease. *Cell* **167**, 1415-1429.e19 (2016).

69.	Vujkovic, M. *et al.* Discovery of 318 new risk loci for type 2 diabetes and related vascular outcomes among 1.4 million participants in a multi-ancestry meta-analysis. *Nat Genet* **52**, 680–691 (2020).

70.	Hoffmann, T. J. *et al.* A large electronic-health-record-based genome-wide study of serum lipids. *Nat Genet* **50**, 401–413 (2018).

71.	International League Against Epilepsy Consortium on Complex, E. Genome-wide mega-analysis identifies 16 loci and highlights diverse biological mechanisms in the common epilepsies. *Nat Commun* **9**, 5269 (2018).

72.	Lee, K. Y., Qi, Z., Yu, Y. P. & Wang, J. H. Neuronal Cdc2-like kinases: neuron-specific forms of Cdk5. *Int J Biochem Cell Biol* **29**, 951–8 (1997).

73.	Floriano-Sanchez, E. *et al.* Differential Gene Expression Profile Induced by Valproic Acid (VPA) in Pediatric Epileptic Patients. *Genes (Basel)* **9**, (2018).

74.	Schwarz, M. *et al.* The bile acid synthetic gene 3beta-hydroxy-Delta(5)-C(27)-steroid oxidoreductase is mutated in progressive intrahepatic cholestasis. *J Clin Invest* **106**, 1175–84 (2000).

75.	Ferrari, R. *et al.* Frontotemporal dementia and its subtypes: a genome-wide association study. *Lancet Neurol* **13**, 686–99 (2014).

76.	Nguyen, T., Liu, X. K., Zhang, Y. & Dong, C. BTNL2, a butyrophilin-like molecule that functions to inhibit T cell activation. *J Immunol* **176**, 7354–60 (2006).

77.	Valentonyte, R. *et al.* Sarcoidosis is associated with a truncating splice site mutation in BTNL2. *Nat Genet* **37**, 357–64 (2005).

78.	Fortes, G. C. C. *et al.* Rapidly progressive dementia due to neurosarcoidosis. *Dement Neuropsychol* **7**, 428–434 (2013).

79.	Nagel, M., Watanabe, K., Stringer, S., Posthuma, D. & van der Sluis, S. Item-level analyses reveal genetic heterogeneity in neuroticism. *Nat Commun* **9**, 905 (2018).

80.	Jansen, P. R. *et al.* Genome-wide analysis of insomnia in 1,331,010 individuals identifies new risk loci and functional pathways. *Nat Genet* **51**, 394–403 (2019).

81.	Goes, F. S. *et al.* Genome-wide association study of schizophrenia in Ashkenazi Jews. *Am J Med Genet B Neuropsychiatr Genet* **168**, 649–659 (2015).

82.	Sherva, R. *et al.* Genome-wide association study of rate of cognitive decline in Alzheimer’s disease patients identifies novel genes and pathways. *Alzheimers Dement* **16**, 1134–1145 (2020).

83.	Savage, J. E. *et al.* Genome-wide association meta-analysis in 269,867 individuals identifies new genetic and functional links to intelligence. *Nat Genet* **50**, 912–919 (2018).

84.	Davies, G. *et al.* Study of 300,486 individuals identifies 148 independent genetic loci influencing general cognitive function. *Nat Commun* **9**, 2098 (2018).

85.	Kunkle, B. W. *et al.* Genetic meta-analysis of diagnosed Alzheimer’s disease identifies new risk loci and implicates Aβ, tau, immunity and lipid processing. *Nature genetics* **51**, 414–430 (2019).

86.	Zhu, Z. *et al.* Shared genetic and experimental links between obesity-related traits and asthma subtypes in UK Biobank. *J Allergy Clin Immunol* **145**, 537–549 (2020).

87.	Hom, G. *et al.* Association of systemic lupus erythematosus with C8orf13-BLK and ITGAM-ITGAX. *N Engl J Med* **358**, 900–909 (2008).

88.	Mägi, R. *et al.* Contribution of 32 GWAS-identified common variants to severe obesity in European adults referred for bariatric surgery. *PLoS One* **8**, e70735 (2013).

89.	Baselmans, B. M. L. *et al.* Multivariate genome-wide analyses of the well-being spectrum. *Nat Genet* **51**, 445–451 (2019).

90.	Jones, S. E. *et al.* Genome-wide association analyses of chronotype in 697,828 individuals provides insights into circadian rhythms. *Nat Commun* **10**, 343 (2019).

91.	Jersild, C., Svejgaard, A. & Fog, T. HL-A antigens and multiple sclerosis. *Lancet (London, England)* **1**, 1240–1 (1972).

92.	Ligers, A. *et al.* Evidence of linkage with HLA-DR in DRB1*15-negative families with multiple sclerosis. *American journal of human genetics* **69**, 900–3 (2001).

93.	International Multiple Sclerosis Genetics, C. *et al.* Risk alleles for multiple sclerosis identified by a genomewide study. *N Engl J Med* **357**, 851–62 (2007).

94.	International Multiple Sclerosis Genetics, C. *et al.* Genetic risk and a primary role for cell-mediated immune mechanisms in multiple sclerosis. *Nature* **476**, 214–9 (2011).

95.	Alcina, A. *et al.* Multiple sclerosis risk variant HLA-DRB1*1501 associates with high expression of DRB1 gene in different human populations. *PLoS One* **7**, e29819 (2012).

96.	Apperson, M. L. *et al.* Genome wide differences of gene expression associated with HLA-DRB1 genotype in multiple sclerosis: a pilot study. *Journal of neuroimmunology* **257**, 90–6 (2013).

97.	Kular, L. *et al.* DNA methylation as a mediator of HLA-DRB1*15:01 and a protective variant in multiple sclerosis. *Nat Commun* **9**, 2397 (2018).

98.	Puskarjov, M. *et al.* A variant of KCC2 from patients with febrile seizures impairs neuronal Cl- extrusion and dendritic spine formation. *EMBO reports* **15**, 723–9 (2014).

99.	Stödberg, T. *et al.* Mutations in SLC12A5 in epilepsy of infancy with migrating focal seizures. *Nature communications* **6**, 8038 (2015).

100.	Zhang, Y. *et al.* Purification and Characterization of Progenitor and Mature Human Astrocytes Reveals Transcriptional and Functional Differences with Mouse. *Neuron* **89**, 37–53 (2016).

101.	Bishop, G. A., Stunz, L. L. & Hostager, B. S. TRAF3 as a Multifaceted Regulator of B Lymphocyte Survival and Activation. *Frontiers in immunology* **9**, 2161 (2018).

102.	Xie, P., Hostager, B. S. & Bishop, G. A. Requirement for TRAF3 in signaling by LMP1 but not CD40 in B lymphocytes. *The Journal of experimental medicine* **199**, 661–71 (2004).

103.	Hussein, H. A. M. & Akula, S. M. miRNA-36 inhibits KSHV, EBV, HSV-2 infection of cells via stifling expression of interferon induced transmembrane protein 1 (IFITM1). *Sci Rep* **7**, 17972 (2017).

104.	Roostaei, T. *et al.* Impact of genetic susceptibility to multiple sclerosis on the T cell epigenome: proximal and distal effects. *bioRxiv* 2020.07.11.198721 (2020) doi:10.1101/2020.07.11.198721.

105.	Quintero, O. A. *et al.* Human Myo19 is a novel myosin that associates with mitochondria. *Current biology : CB* **19**, 2008–13 (2009).

106.	Dalla Rosa, I. *et al.* MPV17L2 is required for ribosome assembly in mitochondria. *Nucleic acids research* **42**, 8500–15 (2014).

107.	Campbell, G. & Mahad, D. J. Mitochondrial dysfunction and axon degeneration in progressive multiple sclerosis. *FEBS Lett* **592**, 1113–1121 (2018).

108.	Lanata, C. M. *et al.* Genetic contributions to lupus nephritis in a multi-ethnic cohort of systemic lupus erythematous patients. *PloS one* **13**, e0199003 (2018).

109.	Ross, K. A. Coherent somatic mutation in autoimmune disease. *PloS one* **9**, e101093 (2014).

110.	Sharma, A. *et al.* Identification of non-HLA genes associated with development of islet autoimmunity and type 1 diabetes in the prospective TEDDY cohort. *Journal of autoimmunity* **89**, 90–100 (2018).

111.	Malecki, J. *et al.* The novel lysine specific methyltransferase METTL21B affects mRNA translation through inducible and dynamic methylation of Lys-165 in human eukaryotic elongation factor 1 alpha (eEF1A). *Nucleic acids research* **45**, 4370–4389 (2017).

112.	Hamey, J. J., Wienert, B., Quinlan, K. G. R. & Wilkins, M. R. METTL21B Is a Novel Human Lysine Methyltransferase of Translation Elongation Factor 1A: Discovery by CRISPR/Cas9 Knockout. *Molecular & cellular proteomics : MCP* **16**, 2229–2242 (2017).

113.	Knight, J. R. P. *et al.* Control of translation elongation in health and disease. *Disease models & mechanisms* **13**, (2020).

114.	Talapatra, S., Wagner, J. D. & Thompson, C. B. Elongation factor-1 alpha is a selective regulator of growth factor withdrawal and ER stress-induced apoptosis. *Cell death and differentiation* **9**, 856–61 (2002).

115.	Chalorak, P., Dharmasaroja, P. & Meemon, K. Downregulation of eEF1A/EFT3-4 Enhances Dopaminergic Neurodegeneration After 6-OHDA Exposure in C. elegans Model. *Frontiers in neuroscience* **14**, 303 (2020).

116.	Prommahom, A. & Dharmasaroja, P. Effects of eEF1A2 knockdown on autophagy in an MPP(+)-induced cellular model of Parkinson’s disease. *Neuroscience research* (2020) doi:10.1016/j.neures.2020.03.013.

117.	Garcia-Esparcia, P. *et al.* Altered machinery of protein synthesis is region- and stage-dependent and is associated with α-synuclein oligomers in Parkinson’s disease. *Acta neuropathologica communications* **3**, 76 (2015).

118.	Cao, S. *et al.* Homozygous EEF1A2 mutation causes dilated cardiomyopathy, failure to thrive, global developmental delay, epilepsy and early death. *Human molecular genetics* **26**, 3545–3552 (2017).

119.	Lam, W. W. *et al.* Novel de novo EEF1A2 missense mutations causing epilepsy and intellectual disability. *Molecular genetics & genomic medicine* **4**, 465–74 (2016).

120.	Nakajima, J. *et al.* De novo EEF1A2 mutations in patients with characteristic facial features, intellectual disability, autistic behaviors and epilepsy. *Clinical genetics* **87**, 356–61 (2015).

121.	Beckelman, B. C. *et al.* Dysregulation of Elongation Factor 1A Expression is Correlated with Synaptic Plasticity Impairments in Alzheimer’s Disease. *Journal of Alzheimer’s disease : JAD* **54**, 669–78 (2016).

122.	Beckelman, B. C., Zhou, X., Keene, C. D. & Ma, T. Impaired Eukaryotic Elongation Factor 1A Expression in Alzheimer’s Disease. *Neuro-degenerative diseases* **16**, 39–43 (2016).

123.	Ahola, S. *et al.* Mitochondrial EFTs defects in juvenile-onset Leigh disease, ataxia, neuropathy, and optic atrophy. *Neurology* **83**, 743–51 (2014).

124.	Emperador, S. *et al.* Molecular-genetic characterization and rescue of a TSFM mutation causing childhood-onset ataxia and nonobstructive cardiomyopathy. *European journal of human genetics : EJHG* **25**, 153–156 (2016).

125.	Smeitink, J. A. *et al.* Distinct clinical phenotypes associated with a mutation in the mitochondrial translation elongation factor EFTs. *American journal of human genetics* **79**, 869–77 (2006).

126.	Barcelos, I. P. de, Troxell, R. M. & Graves, J. S. Mitochondrial Dysfunction and Multiple Sclerosis. *Biology (Basel)* **8**, (2019).

127.	Wang, J. *et al.* TSPAN31 is a critical regulator on transduction of survival and apoptotic signals in hepatocellular carcinoma cells. *FEBS letters* **591**, 2905–2918 (2017).

128.	Gao, X., Leone, G. W. & Wang, H. Cyclin D-CDK4/6 functions in cancer. *Advances in cancer research* **148**, 147–169 (2020).

129.	Deng, J. *et al.* CDK4/6 Inhibition Augments Antitumor Immunity by Enhancing T-cell Activation. *Cancer discovery* **8**, 216–233 (2018).

130.	Goel, S. *et al.* CDK4/6 inhibition triggers anti-tumour immunity. *Nature* **548**, 471–475 (2017).

131.	Sekine, C. *et al.* Successful treatment of animal models of rheumatoid arthritis with small-molecule cyclin-dependent kinase inhibitors. *Journal of immunology (Baltimore, Md. : 1950)* **180**, 1954–61 (2008).

132.	Safavi, A. & Hersh, L. B. Degradation of dynorphin-related peptides by the puromycin-sensitive aminopeptidase and aminopeptidase M. *Journal of neurochemistry* **65**, 389–95 (1995).

133.	Kudo, L. C. *et al.* Puromycin-sensitive aminopeptidase (PSA/NPEPPS) impedes development of neuropathology in hPSA/TAU(P301L) double-transgenic mice. *Human molecular genetics* **20**, 1820–33 (2011).

134.	Karsten, S. L. *et al.* A genomic screen for modifiers of tauopathy identifies puromycin-sensitive aminopeptidase as an inhibitor of tau-induced neurodegeneration. *Neuron* **51**, 549–60 (2006).

135.	Ren, G. *et al.* Cu, Zn-superoxide dismutase 1 (SOD1) is a novel target of Puromycin-sensitive aminopeptidase (PSA/NPEPPS): PSA/NPEPPS is a possible modifier of amyotrophic lateral sclerosis. *Molecular neurodegeneration* **6**, 29 (2011).

136.	Lin, Y.-H. *et al.* Identification of ten novel genes involved in human spermatogenesis by microarray analysis of testicular tissue. *Fertil Steril* **86**, 1650–1658 (2006).

137.	Giordana, M. T., Richiardi, P., Trevisan, E., Boghi, A. & Palmucci, L. Abnormal ubiquitination of axons in normally myelinated white matter in multiple sclerosis brain. *Neuropathol Appl Neurobiol* **28**, 35–41 (2002).

138.	O’Connor, A. M., Crawley, A. M. & Angel, J. B. Interleukin-7 enhances memory CD8(+) T-cell recall responses in health but its activity is impaired in human immunodeficiency virus infection. *Immunology* **131**, 525–536 (2010).

139.	Kreft, K. L. *et al.* Decreased systemic IL-7 and soluble IL-7Rα in multiple sclerosis patients. *Genes Immun* **13**, 587–592 (2012).

140.	Papadopoulou, L. C. *et al.* Fatal infantile cardioencephalomyopathy with COX deficiency and mutations in SCO2, a COX assembly gene. *Nat Genet* **23**, 333–337 (1999).

141.	Tran-Viet, K.-N. *et al.* Mutations in SCO2 are associated with autosomal-dominant high-grade myopia. *Am J Hum Genet* **92**, 820–826 (2013).

142.	Alliel, P. M. *et al.* Myoneurin, a novel member of the BTB/POZ-zinc finger family highly expressed in human muscle. *Biochem Biophys Res Commun* **273**, 385–391 (2000).

143.	Yang, F. *et al.* SET8 promotes epithelial-mesenchymal transition and confers TWIST dual transcriptional activities. *EMBO J* **31**, 110–123 (2012).

144.	Chen, M.-H. *et al.* Trans-ethnic and Ancestry-Specific Blood-Cell Genetics in 746,667 Individuals from 5 Global Populations. *Cell* **182**, 1198-1213.e14 (2020).

145.	Landgrave-Gómez, J., Mercado-Gómez, O. & Guevara-Guzmán, R. Epigenetic mechanisms in neurological and neurodegenerative diseases. *Front Cell Neurosci* **9**, 58 (2015).

146.	Kuehner, J. N., Bruggeman, E. C., Wen, Z. & Yao, B. Epigenetic Regulations in Neuropsychiatric Disorders. *Front Genet* **10**, 268 (2019).

147.	Hüttenrauch, M. *et al.* Glycoprotein NMB: a novel Alzheimer’s disease associated marker expressed in a subset of activated microglia. *Acta Neuropathol Commun* **6**, 108 (2018).

148.	Moloney, E. B., Moskites, A., Ferrari, E. J., Isacson, O. & Hallett, P. J. The glycoprotein GPNMB is selectively elevated in the substantia nigra of Parkinson’s disease patients and increases after lysosomal stress. *Neurobiol Dis* **120**, 1–11 (2018).

149.	Purlyte, E. *et al.* Rab29 activation of the Parkinson’s disease-associated LRRK2 kinase. *EMBO J* **37**, 1–18 (2018).

150.	Kuwahara, T. & Iwatsubo, T. The Emerging Functions of LRRK2 and Rab GTPases in the Endolysosomal System. *Front Neurosci* **14**, 227 (2020).

151.	Dehay, B. *et al.* Pathogenic lysosomal depletion in Parkinson’s disease. *J Neurosci* **30**, 12535–12544 (2010).

152.	Berkovic, S. F. *et al.* Array-based gene discovery with three unrelated subjects shows SCARB2/LIMP-2 deficiency causes myoclonus epilepsy and glomerulosclerosis. *Am J Hum Genet* **82**, 673–684 (2008).

153.	Mayo, L. *et al.* Dual role of CD38 in microglial activation and activation-induced cell death. *J Immunol* **181**, 92–103 (2008).

154.	Camacho-Pereira, J. *et al.* CD38 Dictates Age-Related NAD Decline and Mitochondrial Dysfunction through an SIRT3-Dependent Mechanism. *Cell Metab* **23**, 1127–1139 (2016).

155.	Giannuzzi, G. *et al.* Evolutionary dynamism of the primate LRRC37 gene family. *Genome Res* **23**, 46–59 (2013).

156.	Zody, M. C. *et al.* Evolutionary toggling of the MAPT 17q21.31 inversion region. *Nat Genet* **40**, 1076–1083 (2008).

157.	Qi, T. *et al.* Identifying gene targets for brain-related traits using transcriptomic and methylomic data from blood. *Nat Commun* **9**, (2018).

158.	Nalls, M. A. *et al.* Identification of novel risk loci, causal insights, and heritable risk for Parkinson’s disease: a meta-analysis of genome-wide association studies. *Lancet Neurol* **18**, 1091–1102 (2019).

159.	Lee, J.-R. *et al.* The inhibition of chloride intracellular channel 1 enhances Ca2+ and reactive oxygen species signaling in A549 human lung cancer cells. *Exp Mol Med* **51**, 81 (2019).

160.	Zhao, K. *et al.* Exosome-mediated transfer of CLIC1 contributes to the vincristine-resistance in gastric cancer. *Mol Cell Biochem* **462**, 97–105 (2019).

161.	He, Y.-M. *et al.* Effect of CLIC1 gene silencing on proliferation, migration, invasion and apoptosis of human gallbladder cancer cells. *J Cell Mol Med* **22**, 2569–2579 (2018).

162.	Peretti, M. *et al.* Mutual Influence of ROS, pH, and CLIC1 Membrane Protein in the Regulation of G1-S Phase Progression in Human Glioblastoma Stem Cells. *Mol Cancer Ther* **17**, 2451–2461 (2018).

163.	Yue, X., Cui, Y., You, Q., Lu, Y. & Zhang, J. MicroRNA‑124 negatively regulates chloride intracellular channel 1 to suppress the migration and invasion of liver cancer cells. *Oncol Rep* **42**, 1380–1390 (2019).

164.	Zhu, D., Chen, C., Xia, Y., Kong, L.-Y. & Luo, J. A Purified Resin Glycoside Fraction from Pharbitidis Semen Induces Paraptosis by Activating Chloride Intracellular Channel-1 in Human Colon Cancer Cells. *Integr Cancer Ther* **18**, 1534735418822120 (2019).

165.	Tian, C. *et al.* Genome-wide association and HLA region fine-mapping studies identify susceptibility loci for multiple common infections. *Nat Commun* **8**, 599 (2017).

166.	Kou, I. *et al.* Common variants in a novel gene, FONG on chromosome 2q33.1 confer risk of osteoporosis in Japanese. *PLoS One* **6**, e19641 (2011).

167.	Anderson, D. *et al.* First genome-wide association study in an Australian aboriginal population provides insights into genetic risk factors for body mass index and type 2 diabetes. *PLoS One* **10**, e0119333 (2015).

168.	Yang, Y. *et al.* Transgenic overexpression of furin increases epileptic susceptibility. *Cell Death Dis* **9**, 1058 (2018).

169.	Shang, J. *et al.* Cell entry mechanisms of SARS-CoV-2. *Proc Natl Acad Sci U S A* **117**, 11727–11734 (2020).

170.	Autism Spectrum Disorders Working Group of The Psychiatric Genomics Consortium. Meta-analysis of GWAS of over 16,000 individuals with autism spectrum disorder highlights a novel locus at 10q24.32 and a significant overlap with schizophrenia. *Mol Autism* **8**, 21 (2017).

171.	Karlsson Linnér, R. *et al.* Genome-wide association analyses of risk tolerance and risky behaviors in over 1 million individuals identify hundreds of loci and shared genetic influences. *Nat Genet* **51**, 245–257 (2019).

172.	Matsunaga, H. *et al.* Transethnic Meta-Analysis of Genome-Wide Association Studies Identifies Three New Loci and Characterizes Population-Specific Differences for Coronary Artery Disease. *Circ Genom Precis Med* **13**, e002670 (2020).

173.	Pilling, L. C. *et al.* Human longevity: 25 genetic loci associated in 389,166 UK biobank participants. *Aging (Albany NY)* **9**, 2504–2520 (2017).

174.	Zhang, Z. *et al.* Molecular architecture of the human 17S U2 snRNP. *Nature* **583**, 310–313 (2020).

175.	Zhang, Y. *et al.* Inhibition of Splicing Factor 3b Subunit 1 (SF3B1) Reduced Cell Proliferation, Induced Apoptosis and Resulted in Cell Cycle Arrest by Regulating Homeobox A10 (HOXA10) Splicing in AGS and MKN28 Human Gastric Cancer Cells. *Med Sci Monit* **26**, e919460 (2020).

176.	Li, C. *et al.* Somatic SF3B1 hotspot mutation in prolactinomas. *Nat Commun* **11**, 2506 (2020).

177.	Tang, A. D. *et al.* Full-length transcript characterization of SF3B1 mutation in chronic lymphocytic leukemia reveals downregulation of retained introns. *Nat Commun* **11**, 1438 (2020).

178.	Hyde, C. L. *et al.* Identification of 15 genetic loci associated with risk of major depression in individuals of European descent. *Nat Genet* **48**, 1031–1036 (2016).

179.	Witters, P. *et al.* Autism in patients with propionic acidemia. *Mol Genet Metab* **119**, 317–321 (2016).

180.	Nagel, M. *et al.* Meta-analysis of genome-wide association studies for neuroticism in 449,484 individuals identifies novel genetic loci and pathways. *Nat Genet* **50**, 920–927 (2018).

181.	Lango Allen, H. *et al.* Hundreds of variants clustered in genomic loci and biological pathways affect human height. *Nature* **467**, 832–838 (2010).

182.	Pulit, S. L. *et al.* Meta-analysis of genome-wide association studies for body fat distribution in 694 649 individuals of European ancestry. *Hum Mol Genet* **28**, 166–174 (2019).

183.	Sabater-Lleal, M. *et al.* Multiethnic meta-analysis of genome-wide association studies in >100 000 subjects identifies 23 fibrinogen-associated Loci but no strong evidence of a causal association between circulating fibrinogen and cardiovascular disease. *Circulation* **128**, 1310–1324 (2013).

184.	Ligthart, S. *et al.* Genome Analyses of >200,000 Individuals Identify 58 Loci for Chronic Inflammation and Highlight Pathways that Link Inflammation and Complex Disorders. *Am J Hum Genet* **103**, 691–706 (2018).

185.	Surakka, I. *et al.* The impact of low-frequency and rare variants on lipid levels. *Nat Genet* **47**, 589–597 (2015).

186.	Cha, P.-C. *et al.* A genome-wide association study identifies SNP in DCC is associated with gallbladder cancer in the Japanese population. *J Hum Genet* **57**, 235–237 (2012).

187.	Liu, M. *et al.* Association studies of up to 1.2 million individuals yield new insights into the genetic etiology of tobacco and alcohol use. *Nat Genet* **51**, 237–244 (2019).

188.	Zhang, F. *et al.* Clinical value of jointly detection pleural fluid Midkine, pleural fluid adenosine deaminase, and pleural fluid carbohydrate antigen 125 in the identification of nonsmall cell lung cancer-associated malignant pleural effusion. *J Clin Lab Anal* **32**, e22576 (2018).

189.	Mashaly, A. H., Anwar, R., Ebrahim, M. A., Eissa, L. A. & El Shishtawy, M. M. Diagnostic and Prognostic Value of Talin-1 and Midkine as Tumor Markers in Hepatocellular Carcinoma in Egyptian Patients. *Asian Pac J Cancer Prev* **19**, 1503–1508 (2018).

190.	Kichaev, G. *et al.* Leveraging Polygenic Functional Enrichment to Improve GWAS Power. *Am J Hum Genet* **104**, 65–75 (2019).

191.	Wain, L. V. *et al.* Genome-wide association study identifies six new loci influencing pulse pressure and mean arterial pressure. *Nat Genet* **43**, 1005–1011 (2011).

192.	Locke, A. E. *et al.* Genetic studies of body mass index yield new insights for obesity biology. *Nature* **518**, 197–206 (2015).

193.	Xie, T. *et al.* Genome-wide association study combining pathway analysis for typical sporadic amyotrophic lateral sclerosis in Chinese Han populations. *Neurobiol Aging* **35**, 1778.e9-1778.e23 (2014).

194.	Weiss, R. B. *et al.* Long-range genomic regulators of THBS1 and LTBP4 modify disease severity in duchenne muscular dystrophy. *Ann Neurol* **84**, 234–245 (2018).

195.	Evangelou, E. *et al.* Genetic analysis of over 1 million people identifies 535 new loci associated with blood pressure traits. *Nat Genet* **50**, 1412–1425 (2018).

196.	Ramírez, J. *et al.* Thirty loci identified for heart rate response to exercise and recovery implicate autonomic nervous system. *Nat Commun* **9**, 1947 (2018).

197.	McKay, J. D. *et al.* Large-scale association analysis identifies new lung cancer susceptibility loci and heterogeneity in genetic susceptibility across histological subtypes. *Nat Genet* **49**, 1126–1132 (2017).

198.	Brackertz, M., Boeke, J., Zhang, R. & Renkawitz, R. Two highly related p66 proteins comprise a new family of potent transcriptional repressors interacting with MBD2 and MBD3. *J Biol Chem* **277**, 40958–40966 (2002).

199.	Lu, D. *et al.* A shared genetic contribution to breast cancer and schizophrenia. *Nat Commun* **11**, 4637 (2020).

200.	Zhao, W. *et al.* Identification of new susceptibility loci for type 2 diabetes and shared etiological pathways with coronary heart disease. *Nat Genet* **49**, 1450–1457 (2017).

201.	Fregeau, B. *et al.* De Novo Mutations of RERE Cause a Genetic Syndrome with Features that Overlap Those Associated with Proximal 1p36 Deletions. *Am J Hum Genet* **98**, 963–970 (2016).

202.	Lam, M. *et al.* Pleiotropic Meta-Analysis of Cognition, Education, and Schizophrenia Differentiates Roles of Early Neurodevelopmental and Adult Synaptic Pathways. *Am J Hum Genet* **105**, 334–350 (2019).

203.	Turley, P. *et al.* Multi-trait analysis of genome-wide association summary statistics using MTAG. *Nat Genet* **50**, 229–237 (2018).

204.	Howard, D. M. *et al.* Genome-wide meta-analysis of depression identifies 102 independent variants and highlights the importance of the prefrontal brain regions. *Nat Neurosci* **22**, 343–352 (2019).

205.	Craig, J. E. *et al.* Multitrait analysis of glaucoma identifies new risk loci and enables polygenic prediction of disease susceptibility and progression. *Nat Genet* **52**, 160–166 (2020).

206.	Springelkamp, H. *et al.* New insights into the genetics of primary open-angle glaucoma based on meta-analyses of intraocular pressure and optic disc characteristics. *Hum Mol Genet* **26**, 438–453 (2017).

207.	Hysi, P. G. *et al.* Meta-analysis of 542,934 subjects of European ancestry identifies new genes and mechanisms predisposing to refractive error and myopia. *Nat Genet* **52**, 401–407 (2020).

208.	Giri, A. *et al.* Trans-ethnic association study of blood pressure determinants in over 750,000 individuals. *Nat Genet* **51**, 51–62 (2019).

209.	Morris, J. A. *et al.* An atlas of genetic influences on osteoporosis in humans and mice. *Nat Genet* **51**, 258–266 (2019).

210.	Zhu, Z. *et al.* Shared genetics of asthma and mental health disorders: a large-scale genome-wide cross-trait analysis. *Eur Respir J* **54**, (2019).

211.	Han, Y. *et al.* Genome-wide analysis highlights contribution of immune system pathways to the genetic architecture of asthma. *Nat Commun* **11**, 1776 (2020).

212.	Rhee, E. P. *et al.* A genome-wide association study of the human metabolome in a community-based cohort. *Cell Metab* **18**, 130–143 (2013).

213.	Sass, J. O. *et al.* D-glyceric aciduria is caused by genetic deficiency of D-glycerate kinase (GLYCTK). *Hum Mutat* **31**, 1280–1285 (2010).

214.	Pardiñas, A. F. *et al.* Common schizophrenia alleles are enriched in mutation-intolerant genes and in regions under strong background selection. *Nat Genet* **50**, 381–389 (2018).

215.	Mühleisen, T. W. *et al.* Genome-wide association study reveals two new risk loci for bipolar disorder. *Nat Commun* **5**, 3339 (2014).

216.	Paradiso, S., Andreasen, N. C., O’Leary, D. S., Arndt, S. & Robinson, R. G. Cerebellar size and cognition: correlations with IQ, verbal memory and motor dexterity. *Neuropsychiatry Neuropsychol Behav Neurol* **10**, 1–8 (1997).

217.	Parmeggiani, A., Posar, A., Scaduto, M. C., Chiodo, S. & Giovanardi-Rossi, P. Epilepsy, intelligence, and psychiatric disorders in patients with cerebellar hypoplasia. *J Child Neurol* **18**, 1–4 (2003).

218.	Yoon, Y. B. *et al.* Brain Structural Networks Associated with Intelligence and Visuomotor Ability. *Sci Rep* **7**, 2177 (2017).

219.	Jeste, S. S. The Neurology of Autism Spectrum Disorders. *Curr Opin Neurol* **24**, 132–139 (2011).

220.	Ge, T. *et al.* The Shared Genetic Basis of Educational Attainment and Cerebral Cortical Morphology. *Cereb Cortex* **29**, 3471–3481 (2019).

221.	Hibar, D. P. *et al.* Common genetic variants influence human subcortical brain structures. *Nature* **520**, 224–9 (2015).

222.	Canu, E. *et al.* H1 haplotype of the MAPT gene is associated with lower regional gray matter volume in healthy carriers. *Eur J Hum Genet* **17**, 287–94 (2009).

223.	Kang, H. J. *et al.* Spatio-temporal transcriptome of the human brain. *Nature* **478**, 483–9 (2011).

224.	Tahara, T., Kraus, J. P. & Rosenberg, L. E. An unusual insertion/deletion in the gene encoding the beta-subunit of propionyl-CoA carboxylase is a frequent mutation in Caucasian propionic acidemia. *Proc Natl Acad Sci U S A* **87**, 1372–1376 (1990).

225.	Wolf, B. *et al.* Propionic acidemia: A clinical update. *The Journal of Pediatrics* **99**, 835–846 (1981).

226.	Dejean de la Bâtie, C. *et al.* Acute psychosis in propionic acidemia: 2 case reports. *J Child Neurol* **29**, 274–279 (2014).

227.	Parenti, I. *et al.* MAU2 and NIPBL Variants Impair the Heterodimerization of the Cohesin Loader Subunits and Cause Cornelia de Lange Syndrome. *Cell Rep* **31**, 107647 (2020).

228.	Fujita, Y. *et al.* Decreased cohesin in the brain leads to defective synapse development and anxiety-related behavior. *J Exp Med* **214**, 1431–1452 (2017).

229.	Zaveri, H. P. *et al.* Identification of critical regions and candidate genes for cardiovascular malformations and cardiomyopathy associated with deletions of chromosome 1p36. *PLoS One* **9**, e85600 (2014).

230.	Jaffe, A. E. *et al.* Developmental and genetic regulation of the human cortex transcriptome illuminate schizophrenia pathogenesis. *Nature Neuroscience* **21**, 1117–1125 (2018).

231.	Sieberts, S. K. *et al.* Large eQTL meta-analysis reveals differing patterns between cerebral cortical and cerebellar brain regions. *Scientific Data* **7**, 340 (2020).

232.	Young, A. M. *et al.* A map of transcriptional heterogeneity and regulatory variation in human microglia. *bioRxiv* 2019.12.20.874099 (2019) doi:10.1101/2019.12.20.874099.

233.	Li, H. Aligning sequence reads, clone sequences and assembly contigs with BWA-MEM. *arXiv:1303.3997 [q-bio]* (2013).

234.	Li, Z. *et al.* Genetic variants associated with Alzheimer’s disease confer different cerebral cortex cell-type population structure. *Genome Medicine* **10**, 43 (2018).

235.	Kuo, M.-W., Wang, C.-H., Wu, H.-C., Chang, S.-J. & Chuang, Y.-J. Soluble THSD7A Is an N-Glycoprotein That Promotes Endothelial Cell Migration and Tube Formation in Angiogenesis. *PLOS ONE* **6**, e29000 (2011).

236.	Park, Y. *et al.* Single-cell deconvolution of 3,000 post-mortem brain samples for eQTL and GWAS dissection in mental disorders. *bioRxiv* 2021.01.21.426000 (2021) doi:10.1101/2021.01.21.426000.

237.	Li, Z. *et al.* The TMEM106B FTLD-protective variant, rs1990621, is also associated with increased neuronal proportion. *Acta Neuropathol* **139**, 45–61 (2020).

238.	Lamparter, D., Marbach, D., Rueedi, R., Kutalik, Z. & Bergmann, S. Fast and Rigorous Computation of Gene and Pathway Scores from SNP-Based Summary Statistics. *PLOS Computational Biology* **12**, e1004714 (2016).

239.	The 1000 Genomes Project Consortium. A global reference for human genetic variation. *Nature* **526**, 68–74 (2015).

240.	Köhler, S. *et al.* Expansion of the Human Phenotype Ontology (HPO) knowledge base and resources. *Nucleic Acids Res.* **47**, D1018–D1027 (2019).

241.	Kanehisa, M. & Goto, S. KEGG: kyoto encyclopedia of genes and genomes. *Nucleic Acids Res.* **28**, 27–30 (2000).

242.	Jassal, B. *et al.* The reactome pathway knowledgebase. *Nucleic Acids Res.* **48**, D498–D503 (2020).

243.	The Gene Ontology Resource: 20 years and still GOing strong. *Nucleic Acids Res* **47**, D330–D338 (2019).

244.	Deelen, P. *et al.* Improving the diagnostic yield of exome- sequencing by predicting gene–phenotype associations using large-scale gene expression analysis. *Nat Commun* **10**, 1–13 (2019).

245.	Gabriele, M., Lopez Tobon, A., D’Agostino, G. & Testa, G. The chromatin basis of neurodevelopmental disorders: Rethinking dysfunction along the molecular and temporal axes. *Progress in Neuro-Psychopharmacology and Biological Psychiatry* **84**, 306–327 (2018).

246.	Poisson, A. *et al.* Chromatin remodeling dysfunction extends the etiological spectrum of schizophrenia: a case report. *BMC Med Genet* **21**, (2020).

247.	What is schizophrenia: A neurodevelopmental or neurodegenerative disorder or a combination of both? A critical analysis. https://www.ncbi.nlm.nih.gov/pmc/articles/PMC2824976/.

248.	Slowikowski, K. *et al.* *ggrepel: Automatically Position Non-Overlapping Text Labels with ‘ggplot2’*. (2021).

249.	Wickham, H. *ggplot2: Elegant Graphics for Data Analysis*. (Springer-Verlag, 2009). doi:10.1007/978-0-387-98141-3.

250.	Kassambara, A. *ggpubr: ‘ggplot2’ Based Publication Ready Plots*. (2020).

251.	Garnier, S., Ross, N., Rudis, B., Sciaini, M. & Scherer, C. *viridis: Default Color Maps from ‘matplotlib’*. (2018).

252.	Sarkar, D. *Lattice: Multivariate Data Visualization with R*. (Springer-Verlag, 2008). doi:10.1007/978-0-387-75969-2.

253.	Auguie, B. & Antonov, A. *gridExtra: Miscellaneous Functions for ‘Grid’ Graphics*. (2017).

254.	Dowle, M. *et al.* *data.table: Extension of ‘data.frame’*. (2021).

255.	Wickham, H., François, R., Henry, L., Müller, K. & RStudio. *dplyr: A Grammar of Data Manipulation*. (2021).

256.	Wickham, H. *et al.* *readxl: Read Excel Files*. (2019).

257.	Wickham, H., Seidel, D. & RStudio. *scales: Scale Functions for Visualization*. (2020).

258.	Schloerke, B. *et al.* *GGally: Extension to ‘ggplot2’*. (2021).

259.	Robinson, M. D., McCarthy, D. J. & Smyth, G. K. edgeR: a Bioconductor package for differential expression analysis of digital gene expression data. *Bioinformatics* **26**, 139–140 (2010).

260.	Attali, D. & Baker, C. *ggExtra: Add Marginal Histograms to ‘ggplot2’, and More ‘ggplot2’ Enhancements*. (2019).

261.	Wickham, H., Pedersen, T. L. & RStudio. *gtable: Arrange ‘Grobs’ in Tables*. (2019).

262.	Bengtsson, H. *et al.* *matrixStats: Functions that Apply to Rows and Columns of Matrices (and to Vectors)*. (2021).

263.	Tierney, N. *et al.* *naniar: Data Structures, Summaries, and Visualisations for Missing Data*. (2020).

264.	Wickham, H. The Split-Apply-Combine Strategy for Data Analysis. *Journal of Statistical Software* **40**, 1–29 (2011).

265.	Wickham, H. Reshaping Data with the reshape Package. *Journal of Statistical Software* **21**, 1–20 (2007).

266.	Wickham, H. & RStudio. *stringr: Simple, Consistent Wrappers for Common String Operations*. (2019).

267.	Wickham, H. & RStudio. *tidyr: Tidy Messy Data*. (2020).

268.	Welcome to the Tidyverse. https://tidyverse.tidyverse.org/articles/paper.html.

269.	Alexa, A. & Rahnenfuhrer, J. *topGO: Enrichment Analysis for Gene Ontology*. (Bioconductor version: Development (3.13), 2021). doi:10.18129/B9.bioc.topGO.

270.	Waskom, M. *et al.* *mwaskom/seaborn: v0.11.1 (December 2020)*. (Zenodo, 2020). doi:10.5281/ZENODO.592845.

271.	Hunter, J. D. Matplotlib: A 2D Graphics Environment. *Comput. Sci. Eng.* **9**, 90–95 (2007).

272.	Virtanen, P. *et al.* SciPy 1.0: fundamental algorithms for scientific computing in Python. *Nature Methods* **17**, 261–272 (2020).

273.	Harris, C. R. *et al.* Array programming with NumPy. *Nature* **585**, 357–362 (2020).

274.	Seabold, S. & Perktold, J. Statsmodels: Econometric and Statistical Modeling with Python. in 92–96 (2010). doi:10.25080/Majora-92bf1922-011.

275.	Li, H. Tabix: fast retrieval of sequence features from generic TAB-delimited files. *Bioinformatics* **27**, 718–719 (2011).

276.	Slowikowski, K. *slowkow/pytabix*. (2020).

277.	Pedregosa, F. *et al.* Scikit-learn: Machine Learning in Python. *MACHINE LEARNING IN PYTHON* 6.

278.	Lex, A., Gehlenborg, N., Strobelt, H., Vuillemot, R. & Pfister, H. UpSet: Visualization of Intersecting Sets. *IEEE Transactions on Visualization and Computer Graphics* **20**, 1983–1992 (2014).

279.	Nothman, J. *UpSetPlot: Draw Lex et al.’s UpSet plots with Pandas and Matplotlib*.

280.	*wget: pure python download utility*.

281.	Singh, P. *beautifultable: Print text tables for terminals*.

282.	Leifer, C. *pysqlite3: DB-API 2.0 interface for Sqlite 3.x*.
