## Supplementary Figure 5 for "Brain expression quantitative trait locus and network analysis reveals downstream effects and putative drivers for brain-related diseases"

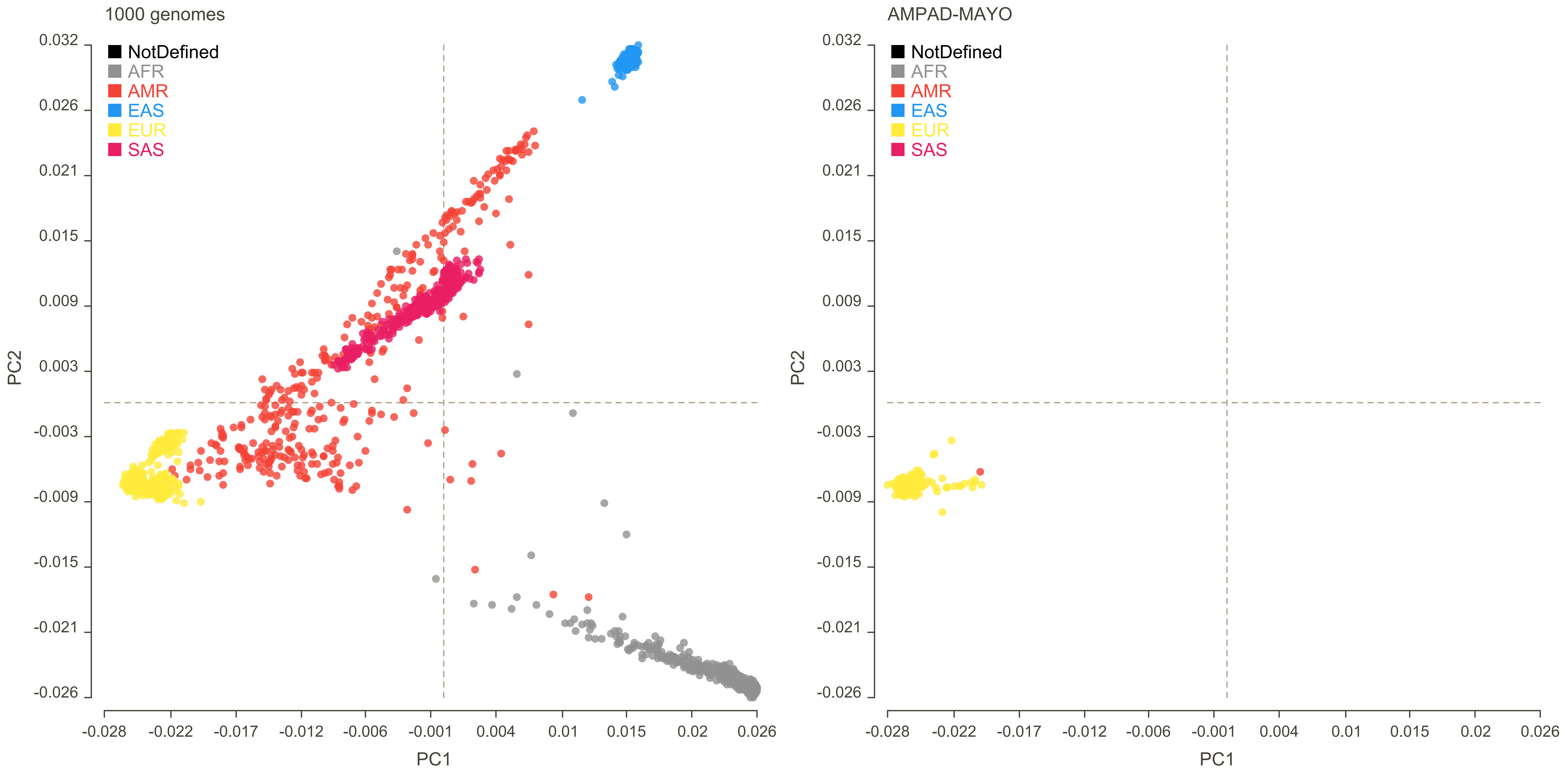
1000 genomes
NotDefined
AFR
AMR
EAS
EUR
SAS
PC2
PC1
0.032
0.026
0.021
0.015
0.009
0.003
-0.003
-0.009
-0.015
-0.021
-0.026
-0.028
-0.022
-0.017
-0.012
-0.006
-0.001
0.004
0.01
0.015
0.02
0.026
AMPAD-MAYO
NotDefined
AFR
AMR
EAS
EUR
SAS
PC2
PC1

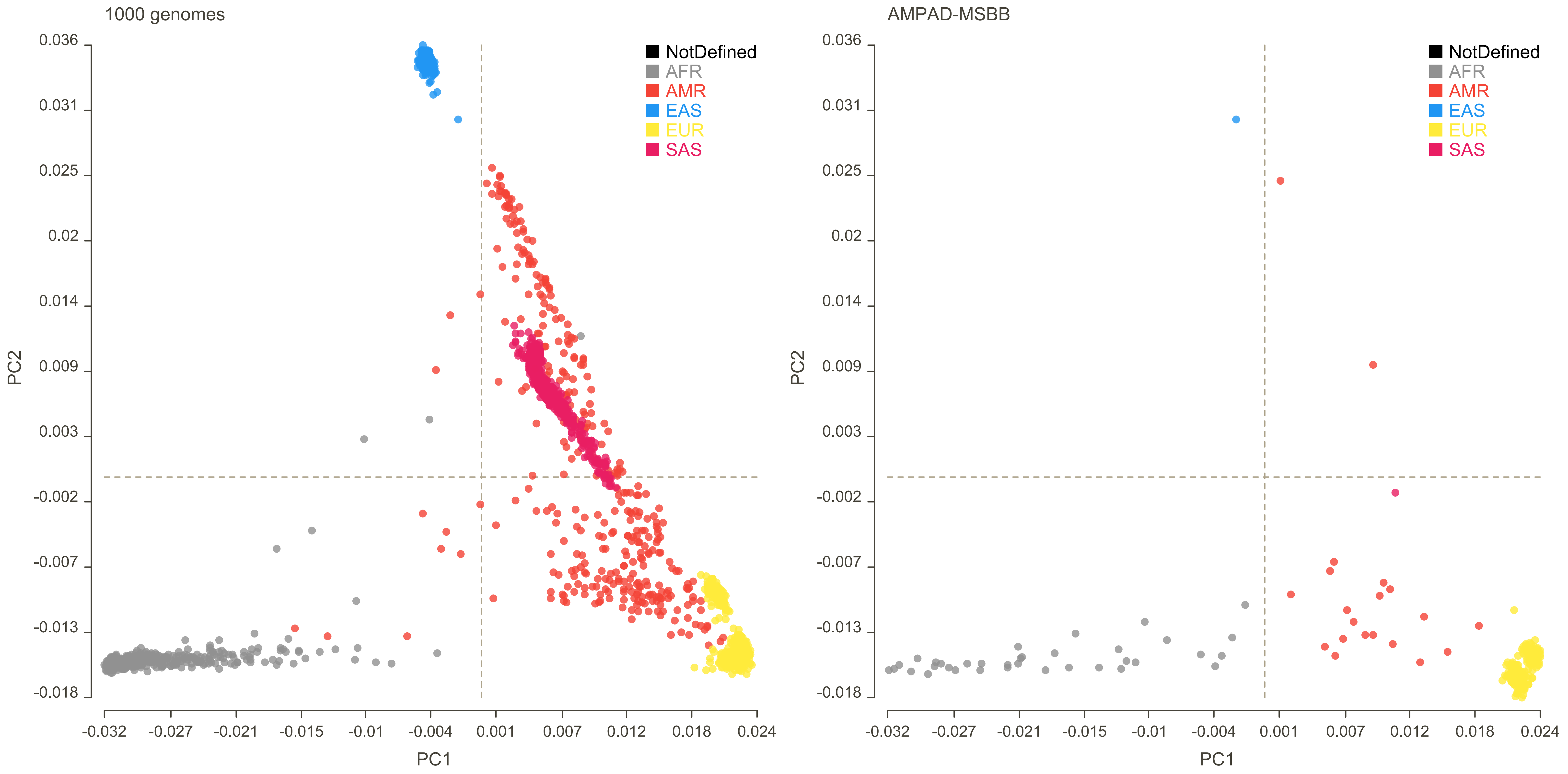

1000 genomes
AMPAD-MSBB
NotDefined
AFR
AMR
EAS
EUR
SAS
PC1
PC2
0.036
0.031
0.025
0.02
0.014
0.009
0.003
-0.002
-0.007
-0.013
-0.018
-0.032
-0.027
-0.021
-0.015
-0.01
-0.004
0.001
0.007
0.012
0.018
0.024

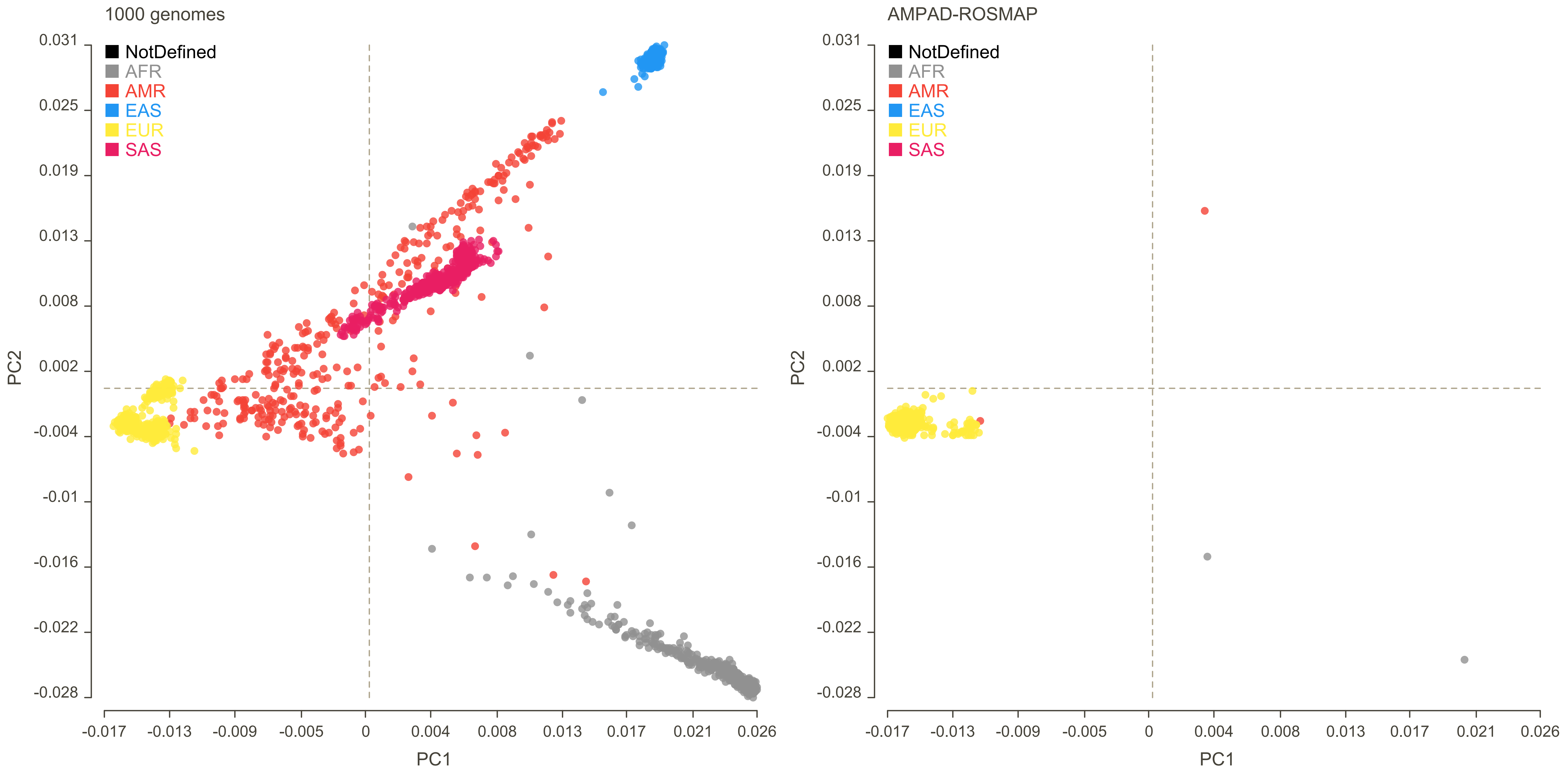

1000 genomes
AMPAD-ROSMAP
NotDefined
AFR
AMR
EAS
EUR
SAS
PC1
PC2
0.031
0.025
0.019
0.013
0.008
0.002
-0.004
-0.01
-0.016
-0.022
-0.028
-0.017
-0.013
-0.009
-0.005
0
0.004
0.008
0.013
0.017
0.021
0.026

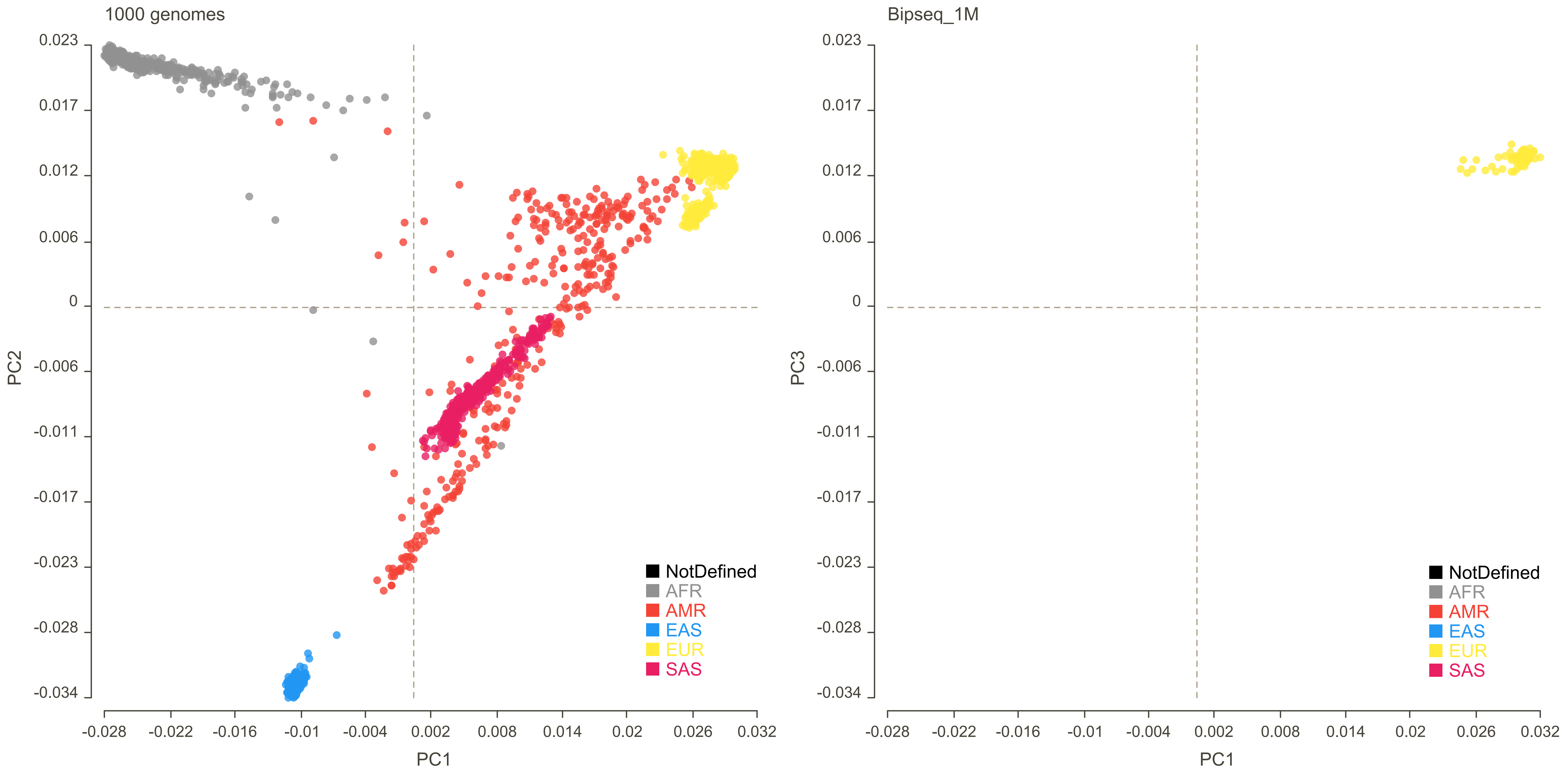
1000 genomes
0.023
0.017
0.012
0.006
0
-0.006
-0.011
-0.017
-0.023
-0.028
-0.034
PC2
-0.028
-0.022
-0.016
-0.01
-0.004
0.002
0.008
0.014
0.02
0.026
0.032
PC1
NotDefined
AFR
AMR
EAS
EUR
SAS
Bipseq_1M
0.023
0.017
0.012
0.006
0
-0.006
-0.011
-0.017
-0.023
-0.028
-0.034
PC3
-0.028
-0.022
-0.016
-0.01
-0.004
0.002
0.008
0.014
0.02
0.026
0.032
PC1
NotDefined
AFR
AMR
EAS
EUR
SAS

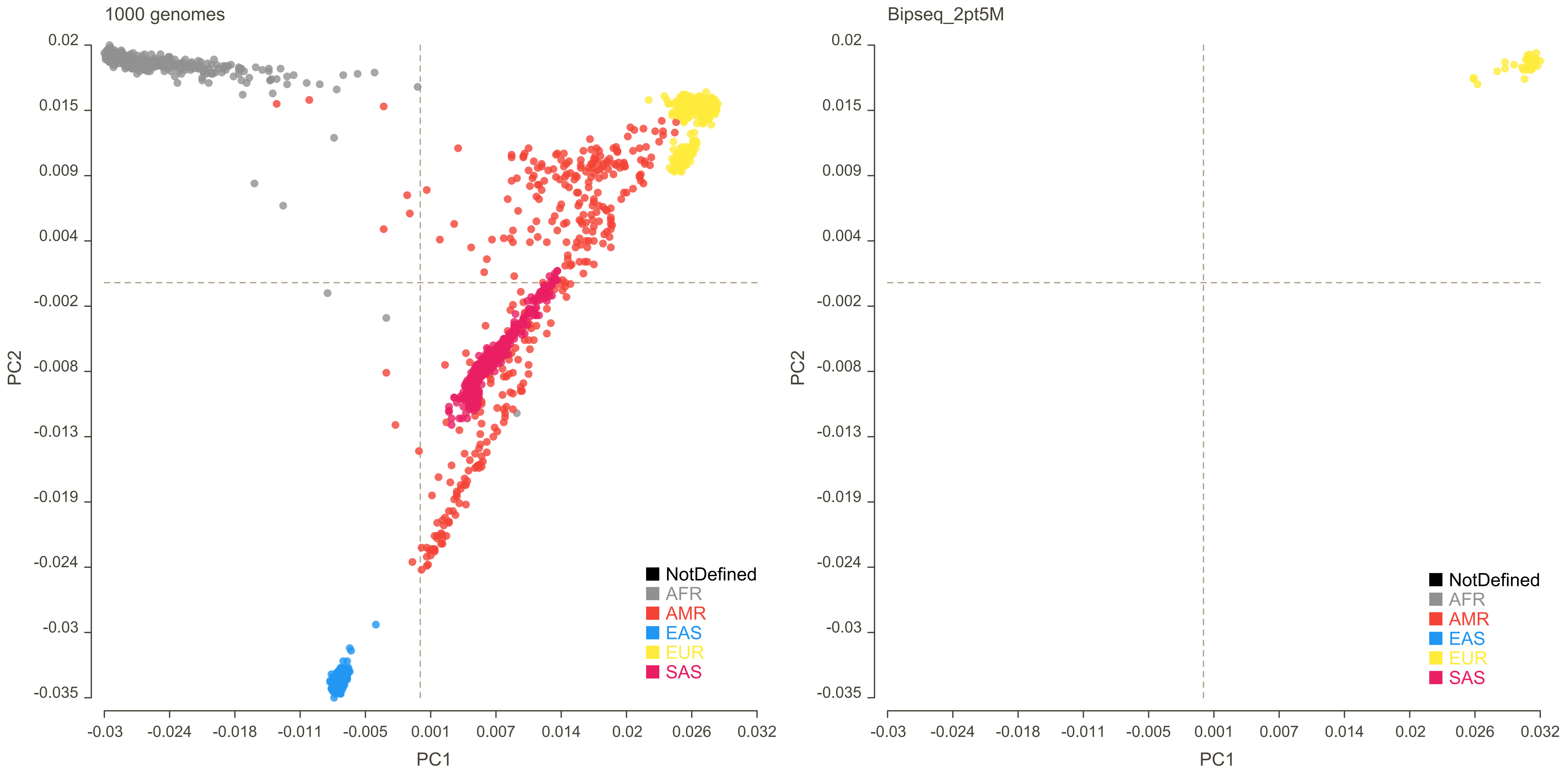

1000 genomes
0.02
0.015
0.009
0.004
-0.002
-0.008
-0.013
-0.019
-0.024
-0.03
-0.035
PC2
-0.03
-0.024
-0.018
-0.011
-0.005
0.001
0.007
0.014
0.02
0.026
0.032
PC1
NotDefined
AFR
AMR
EAS
EUR
SAS
Bipseq_2pt5M
0.02
0.015
0.009
0.004
-0.002
-0.008
-0.013
-0.019
-0.024
-0.03
-0.035
PC2
-0.03
-0.024
-0.018
-0.011
-0.005
0.001
0.007
0.014
0.02
0.026
0.032
PC1
NotDefined
AFR
AMR
EAS
EUR
SAS

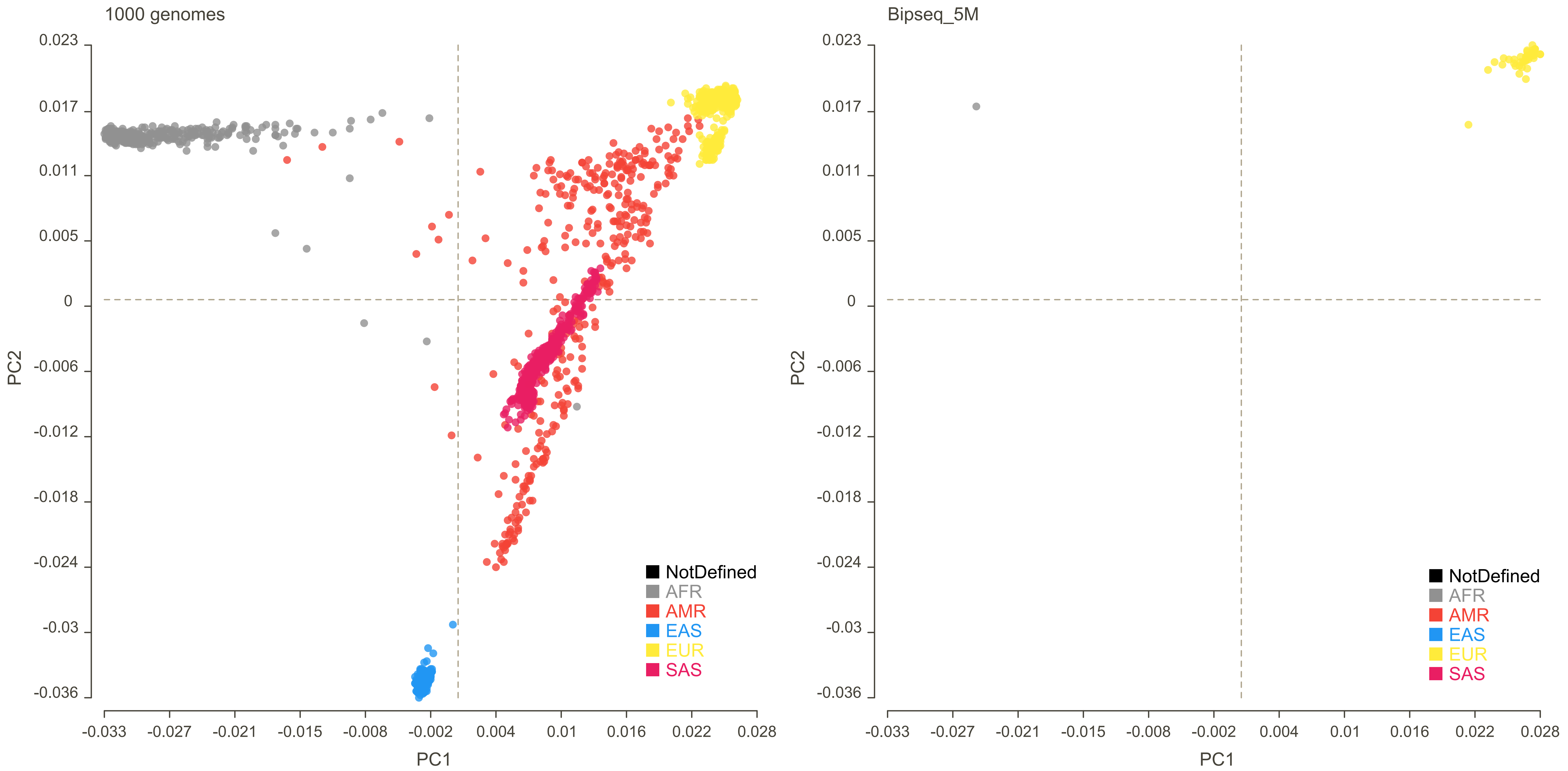
1000 genomes
Bipseq_5M
PC1
PC2
0.023
0.017
0.011
0.005
0
-0.006
-0.012
-0.018
-0.024
-0.03
-0.036
-0.033
-0.027
-0.021
-0.015
-0.008
-0.002
0.004
0.01
0.016
0.022
0.028
NotDefined
AFR
AMR
EAS
EUR
SAS

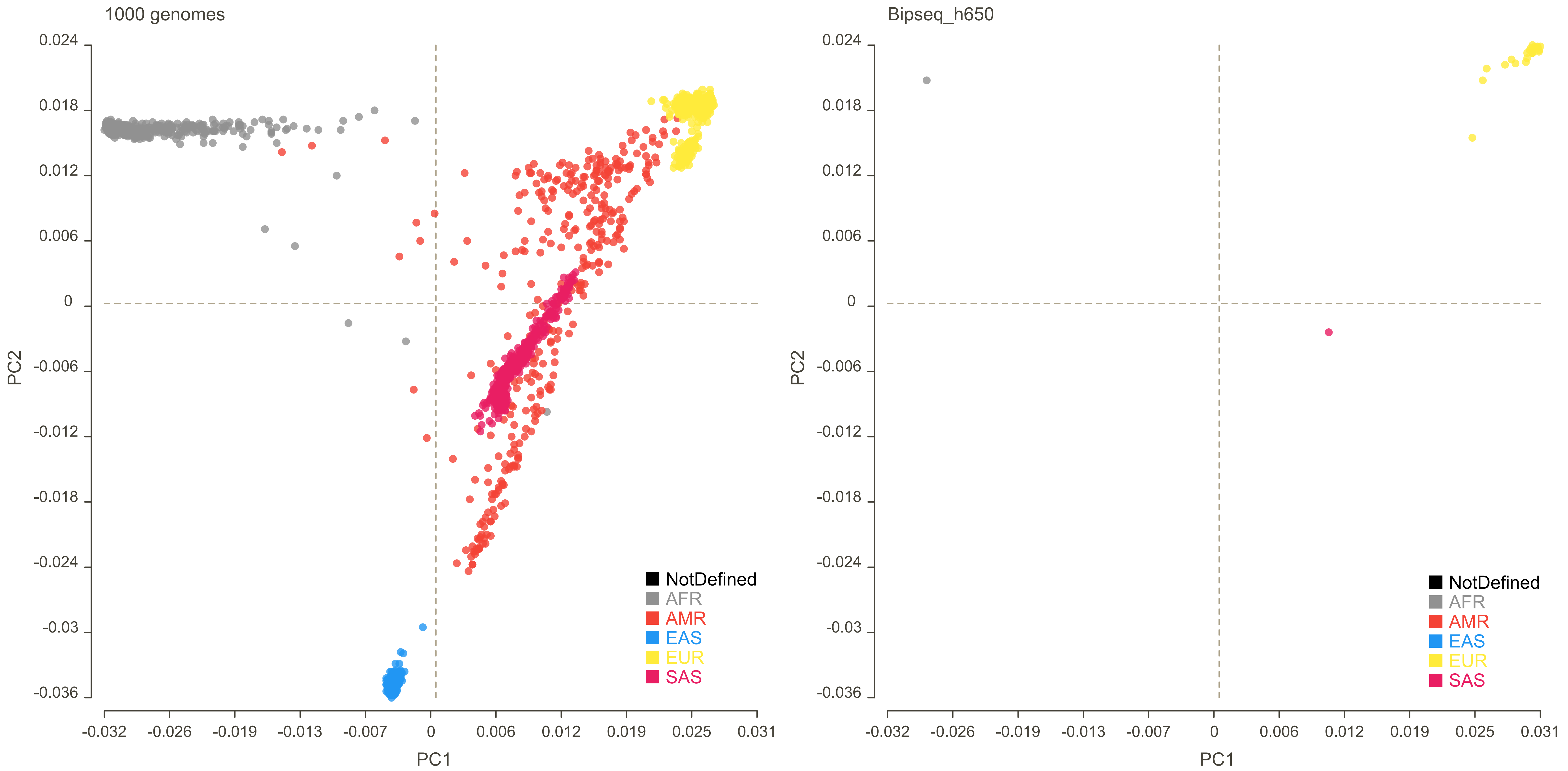

1000 genomes
Bipseq_h650
0.024
0.018
0.012
0.006
0
-0.006
-0.012
-0.018
-0.024
-0.03
-0.036
PC2
-0.032
-0.026
-0.019
-0.013
-0.007
0
0.006
0.012
0.019
0.025
0.031
PC1
NotDefined
AFR
AMR
EAS
EUR
SAS

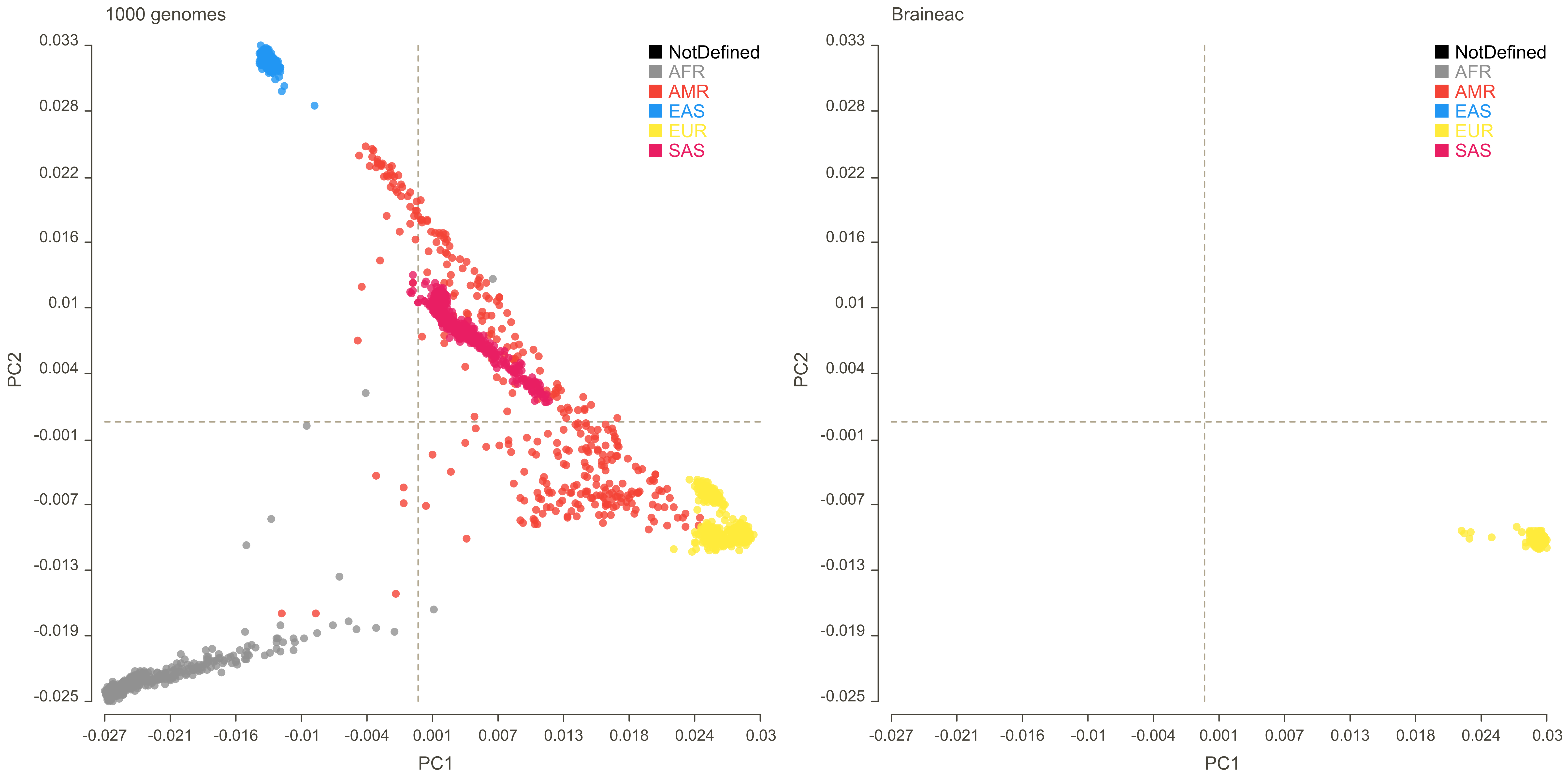

1000 genomes
Braineac
NotDefined
AFR
AMR
EAS
EUR
SAS
PC2
PC1
0.033
0.028
0.022
0.016
0.01
0.004
-0.001
-0.007
-0.013
-0.019
-0.025
-0.027
-0.021
-0.016
-0.01
-0.004
0.001
0.007
0.013
0.018
0.024
0.03

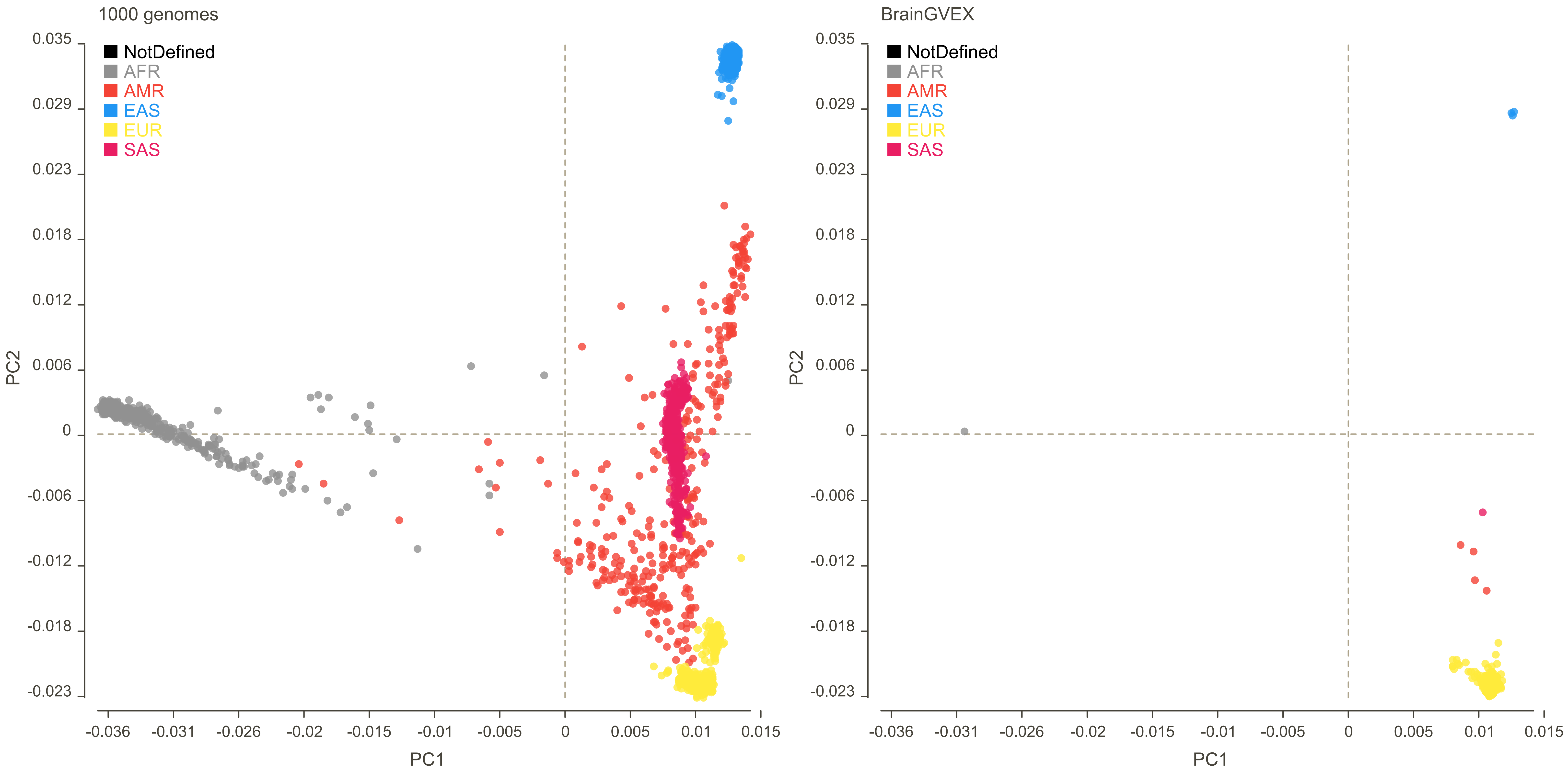
1000 genomes
BrainGVEX
NotDefined
AFR
AMR
EAS
EUR
SAS
PC2
PC1
0.035
0.029
0.023
0.018
0.012
0.006
0
-0.006
-0.012
-0.018
-0.023
-0.036
-0.031
-0.026
-0.02
-0.015
-0.01
-0.005
0
0.005
0.01
0.015

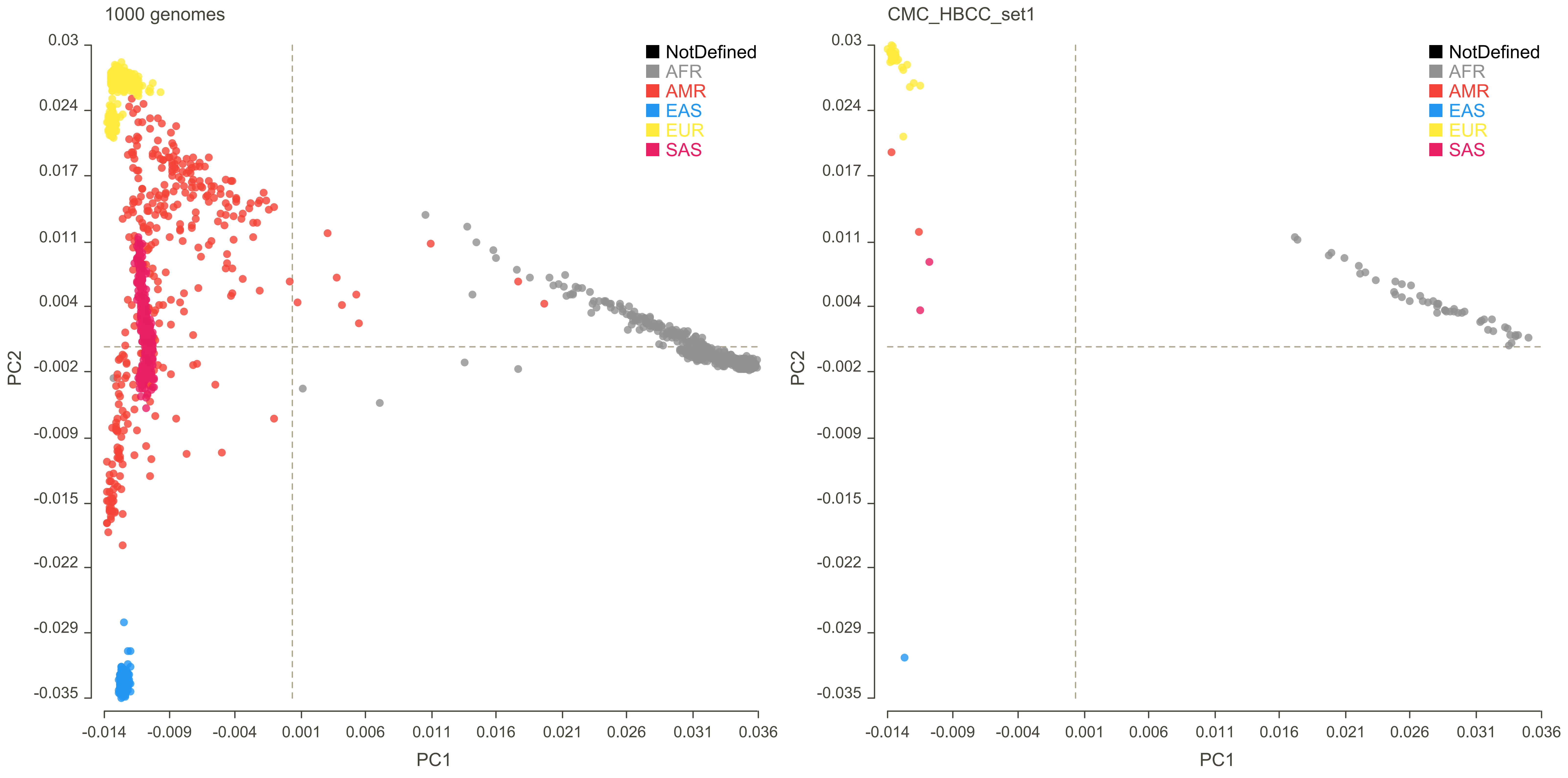

1000 genomes
CMC_HBCC_set1
NotDefined
AFR
AMR
EAS
EUR
SAS
PC2
PC1
0.03
0.024
0.017
0.011
0.004
-0.002
-0.009
-0.015
-0.022
-0.029
-0.035
-0.014
-0.009
-0.004
0.001
0.006
0.011
0.016
0.021
0.026
0.031
0.036

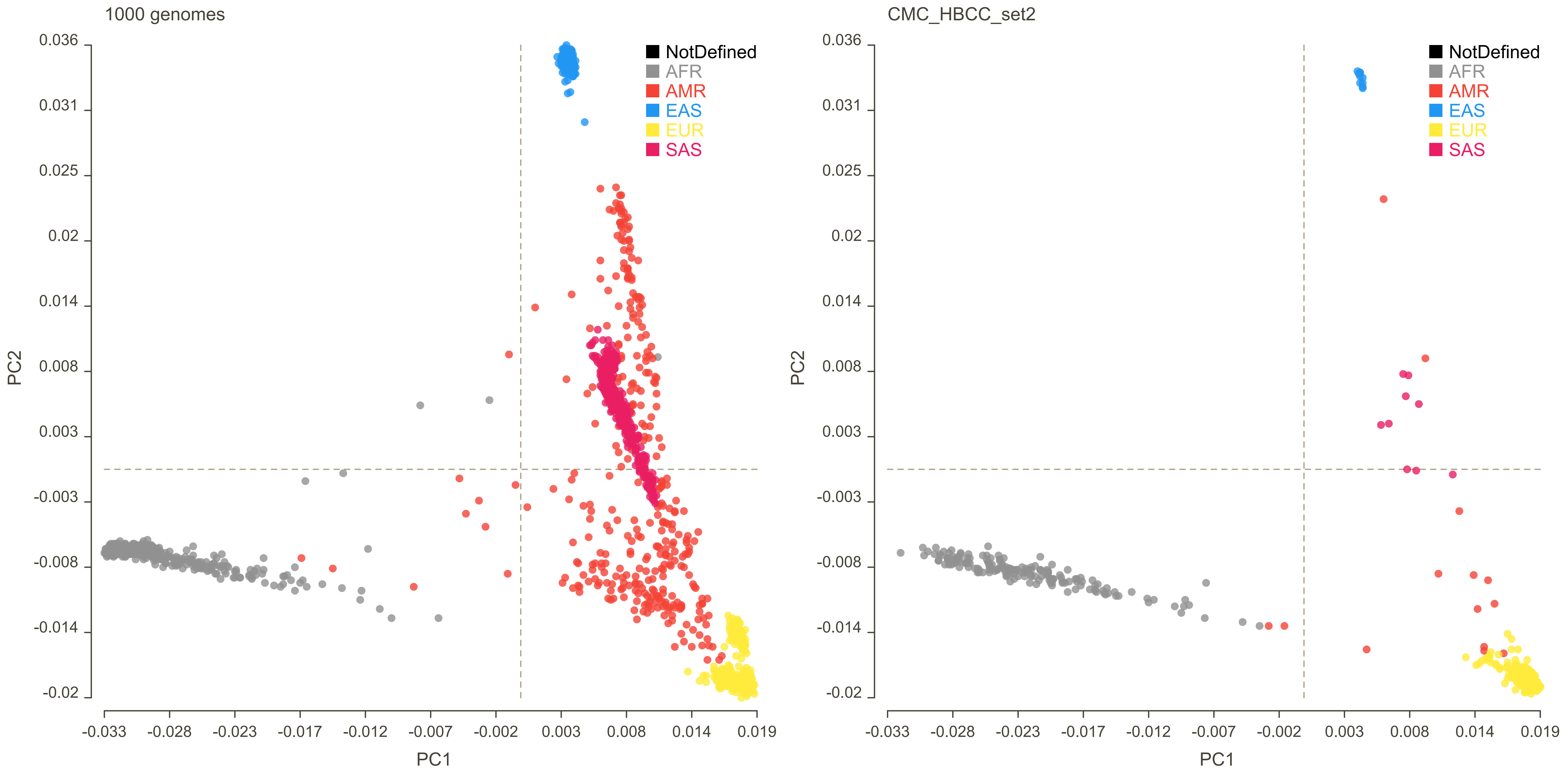

1000 genomes
CMC_HBCC_set2
NotDefined
AFR
AMR
EAS
EUR
SAS
PC2
PC1
0.036
0.031
0.025
0.02
0.014
0.008
0.003
-0.003
-0.008
-0.014
-0.02
-0.033
-0.028
-0.023
-0.017
-0.012
-0.007
-0.002
0.003
0.008
0.014
0.019

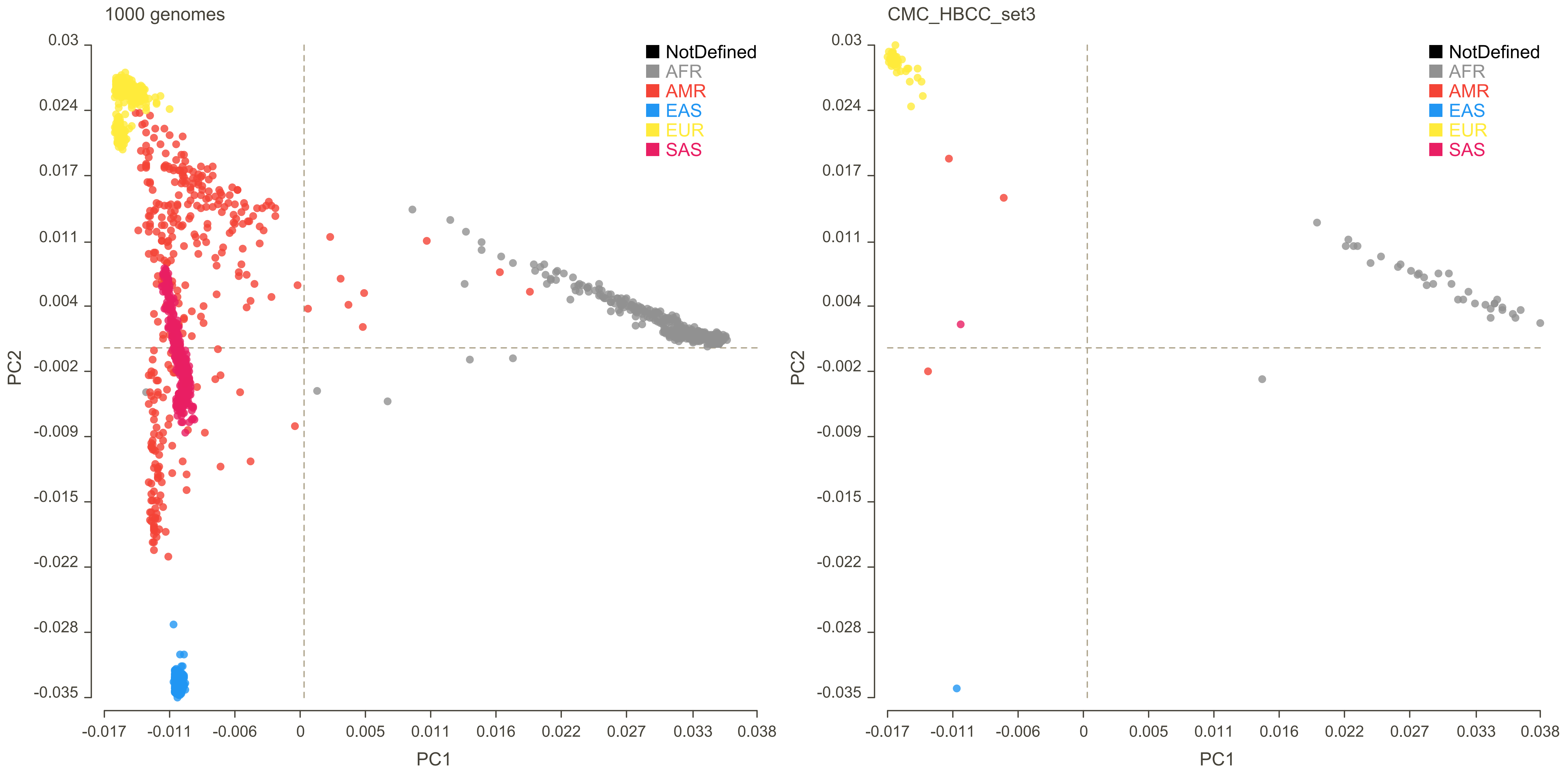
1000 genomes
CMC_HBCC_set3
NotDefined
AFR
AMR
EAS
EUR
SAS
PC1
PC2
0.03
0.024
0.017
0.011
0.004
-0.002
-0.009
-0.015
-0.022
-0.028
-0.035
-0.017
-0.011
-0.006
0
0.005
0.011
0.016
0.022
0.027
0.033
0.038

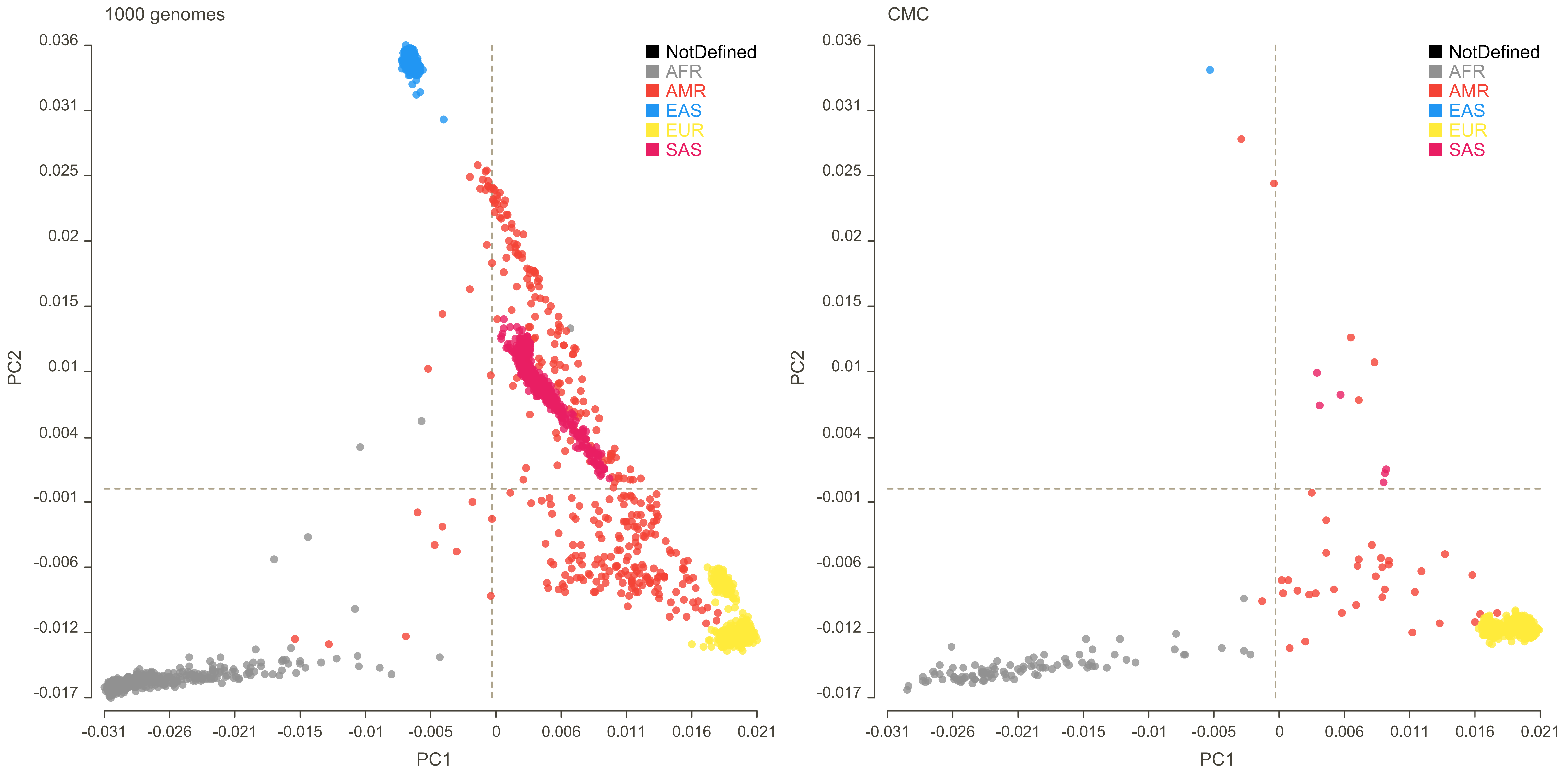
1000 genomes
CMC
NotDefined
AFR
AMR
EAS
EUR
SAS
PC2
PC1
0.036
0.031
0.025
0.02
0.015
0.01
0.004
-0.001
-0.006
-0.012
-0.017
-0.031
-0.026
-0.021
-0.015
-0.01
-0.005
0
0.006
0.011
0.016
0.021

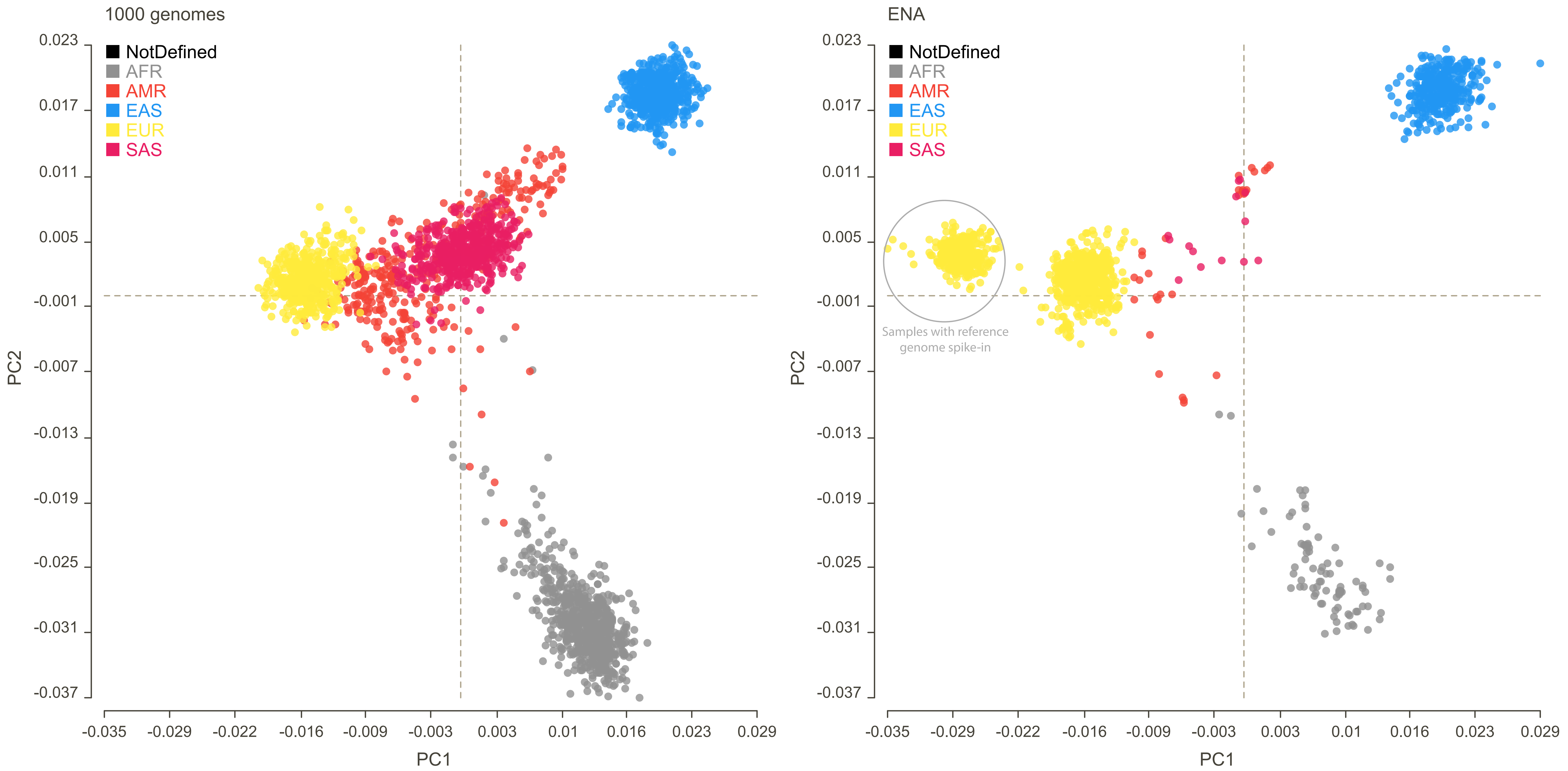
1000 genomes
ENA
NotDefined
AFR
AMR
EAS
EUR
SAS
PC2
PC1
0.023
0.017
0.011
0.005
-0.001
-0.007
-0.013
-0.019
-0.025
-0.031
-0.037
-0.035
-0.029
-0.022
-0.016
-0.009
-0.003
0.003
0.01
0.016
0.023
0.029
Samples with reference genome spike-in

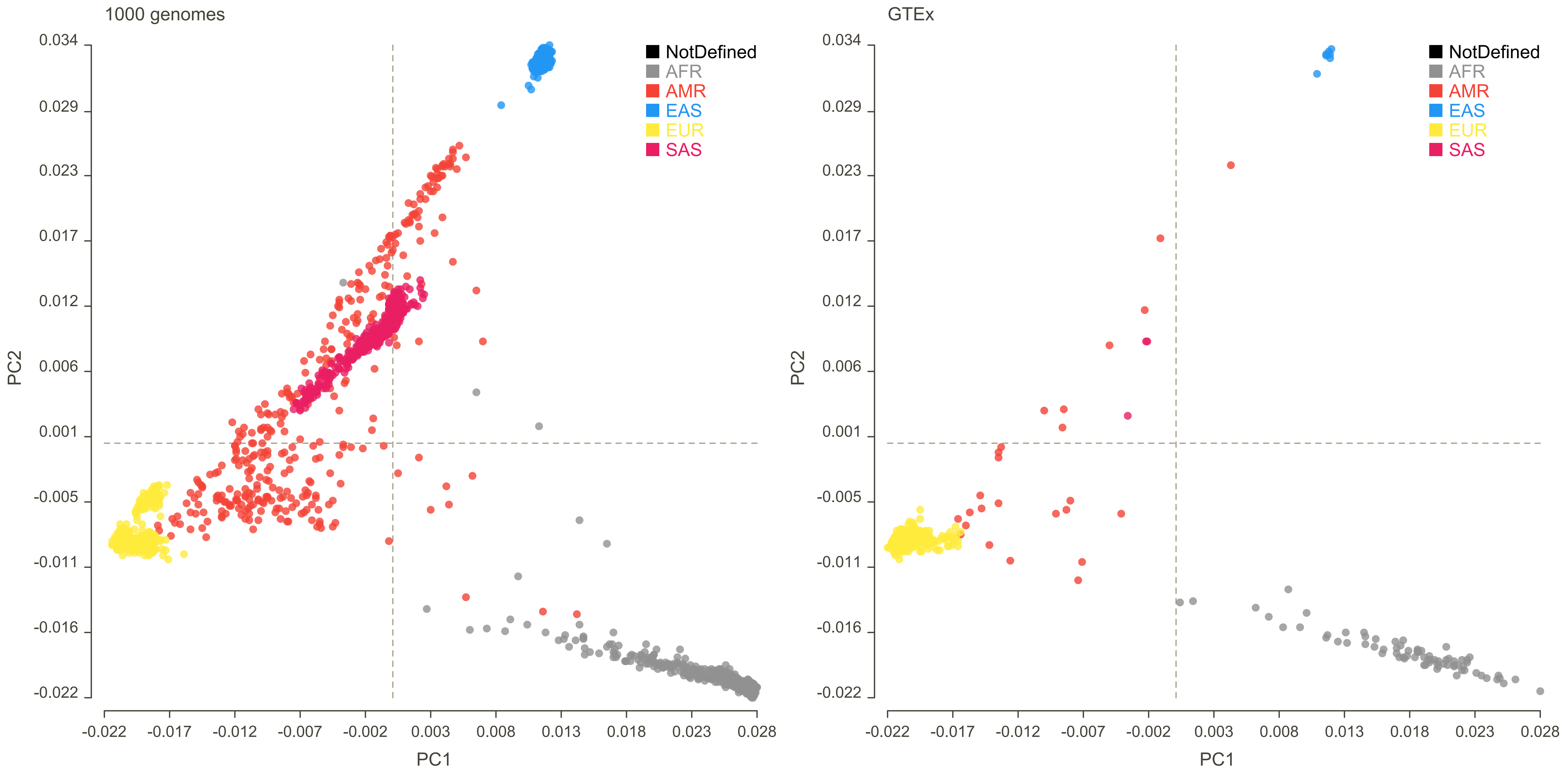
1000 genomes
GTEx
NotDefined
AFR
AMR
EAS
EUR
SAS
PC1
PC2
0.034
0.029
0.023
0.017
0.012
0.006
0.001
-0.005
-0.011
-0.016
-0.022
-0.022
-0.017
-0.012
-0.007
-0.002
0.003
0.008
0.013
0.018
0.023
0.028

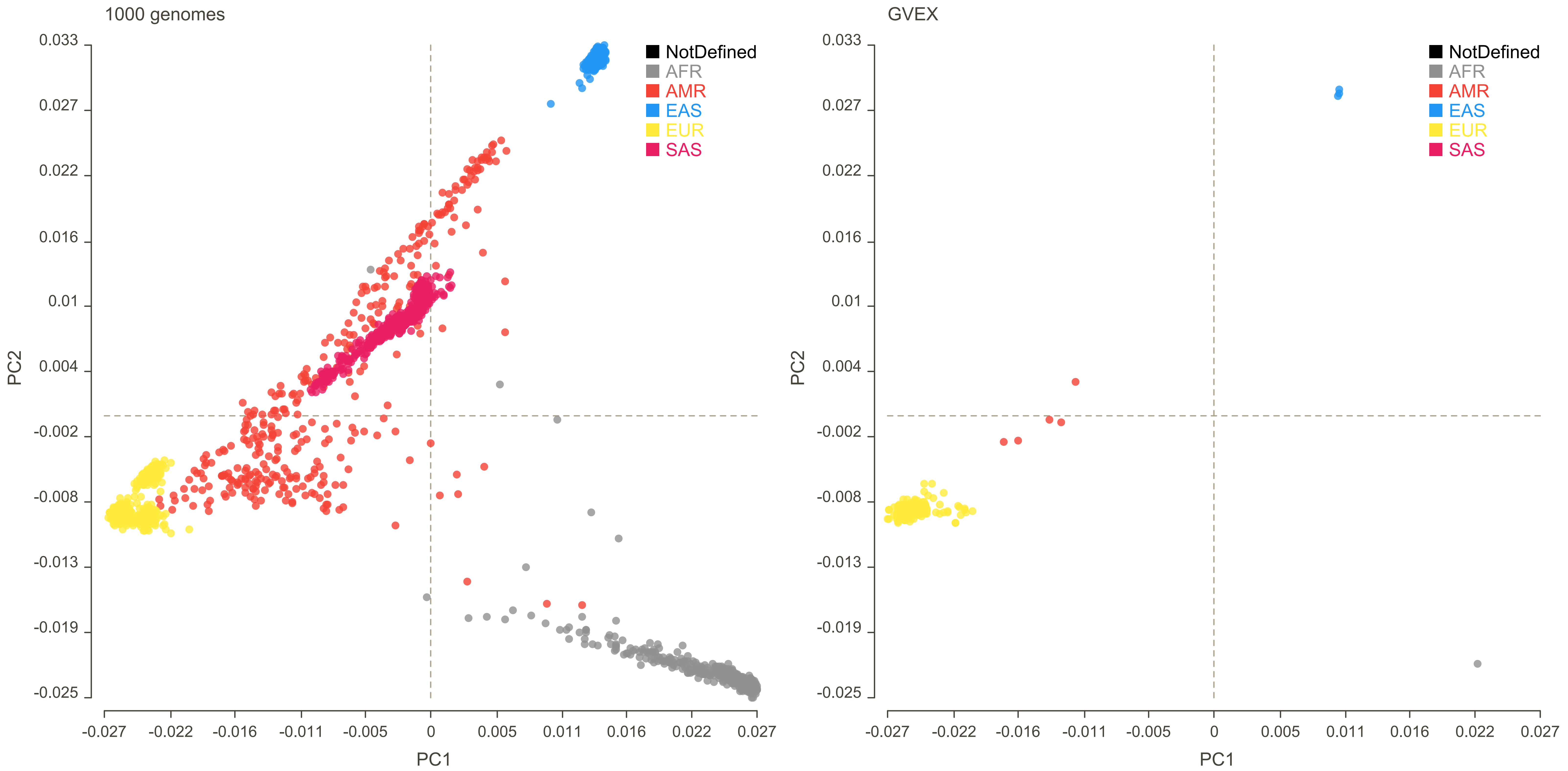
1000 genomes
GVEX
NotDefined
AFR
AMR
EAS
EUR
SAS
PC1
PC2
0.033
0.027
0.022
0.016
0.01
0.004
-0.002
-0.008
-0.013
-0.019
-0.025
-0.027
-0.022
-0.016
-0.011
-0.005
0
0.005
0.011
0.016
0.022
0.027

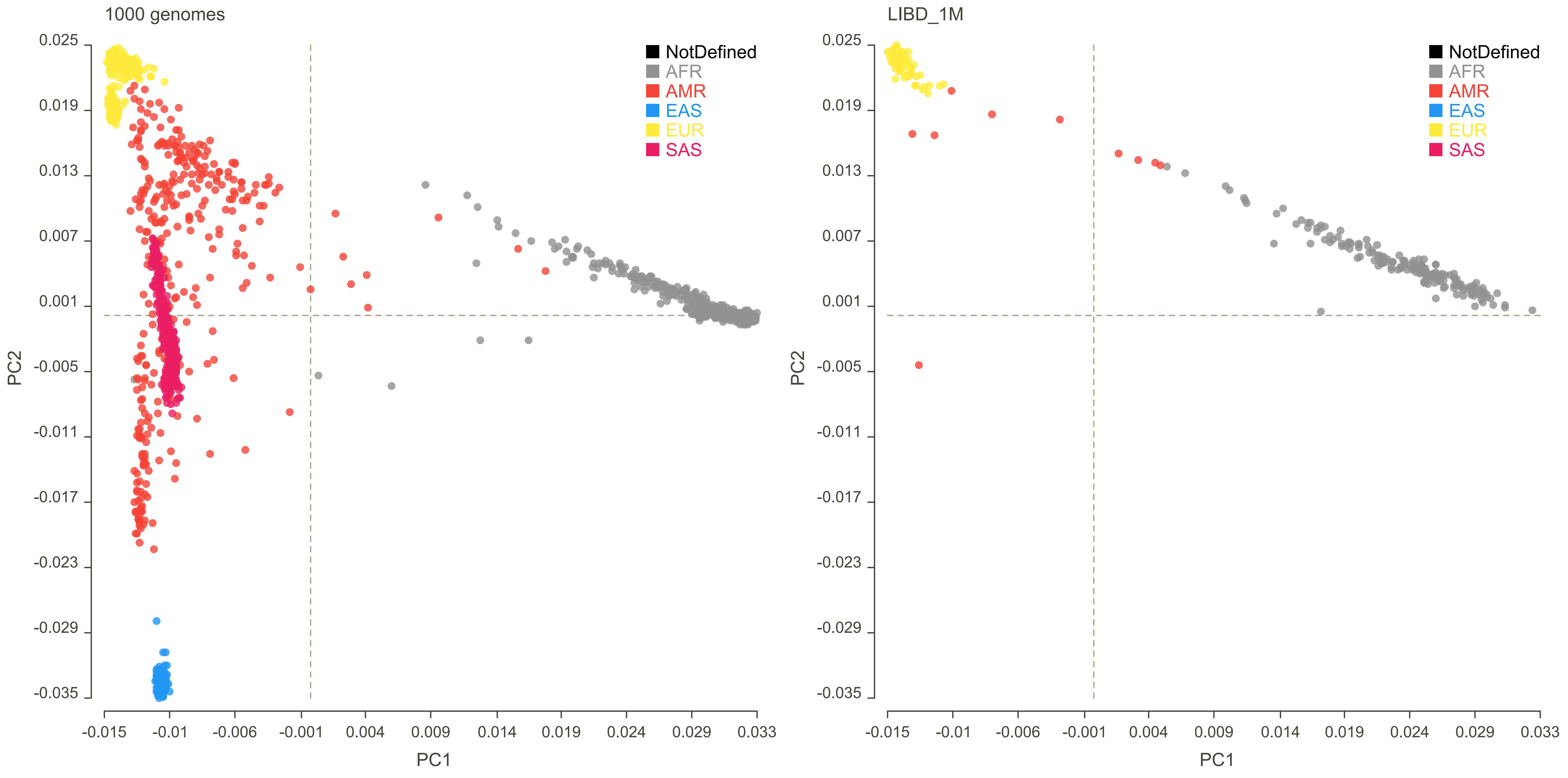
1000 genomes
LIBD_1M
NotDefined
AFR
AMR
EAS
EUR
SAS
PC1
PC2
0.025
0.019
0.013
0.007
0.001
-0.005
-0.011
-0.017
-0.023
-0.029
-0.035
-0.015
-0.01
-0.006
-0.001
0.004
0.009
0.014
0.019
0.024
0.029
0.033

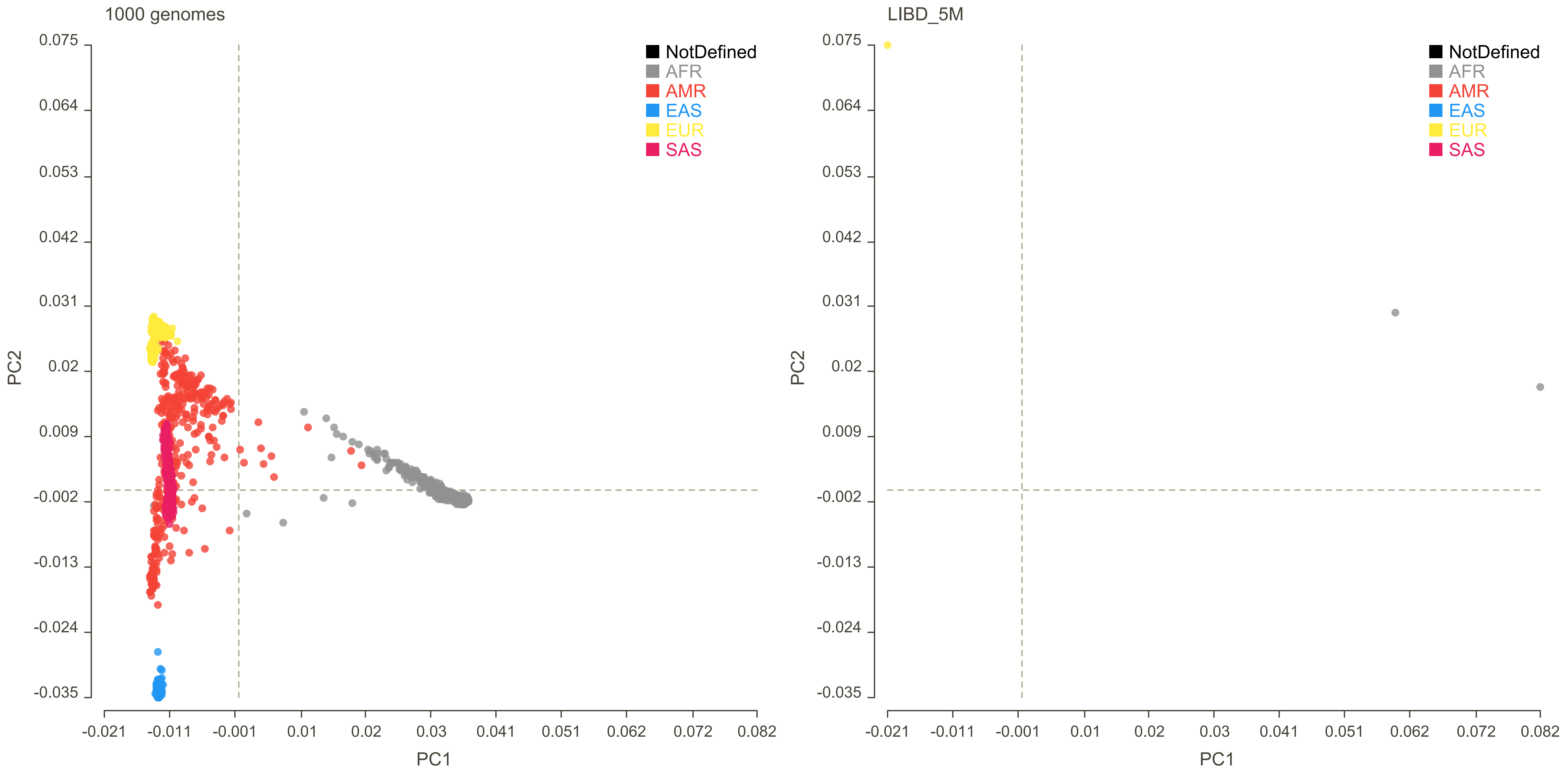

1000 genomes
LIBD_5M
NotDefined
AFR
AMR
EAS
EUR
SAS
PC1
PC2
0.075
0.064
0.053
0.042
0.031
0.02
0.009
-0.002
-0.013
-0.024
-0.035
-0.021
-0.011
-0.001
0.01
0.02
0.03
0.041
0.051
0.062
0.072
0.082

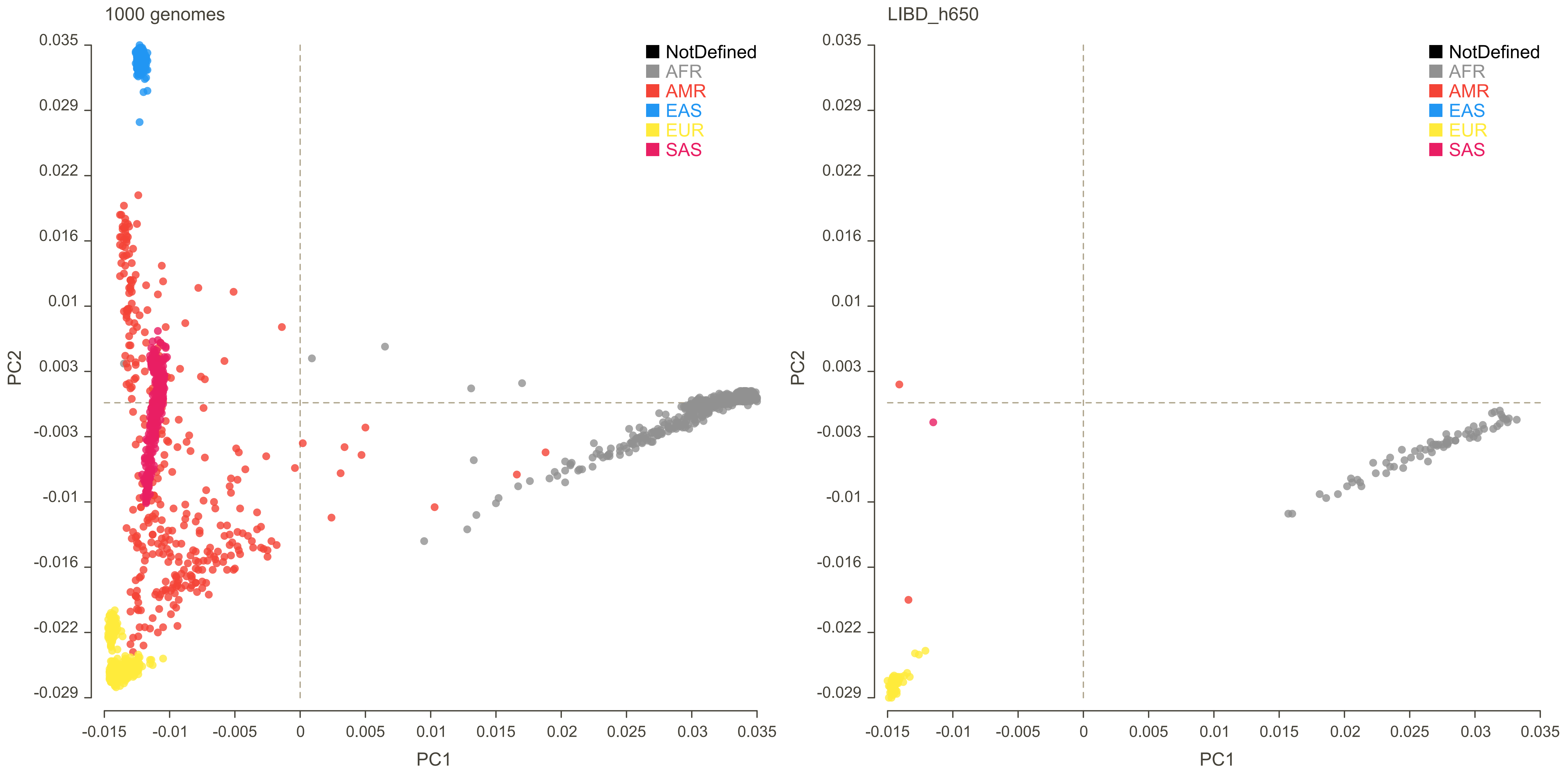

1000 genomes
LIBD_h650
NotDefined
AFR
AMR
EAS
EUR
SAS
PC1
PC2
0.035
0.029
0.022
0.016
0.01
0.003
-0.003
-0.01
-0.016
-0.022
-0.029
-0.015
-0.01
-0.005
0
0.005
0.01
0.015
0.02
0.025
0.03
0.035

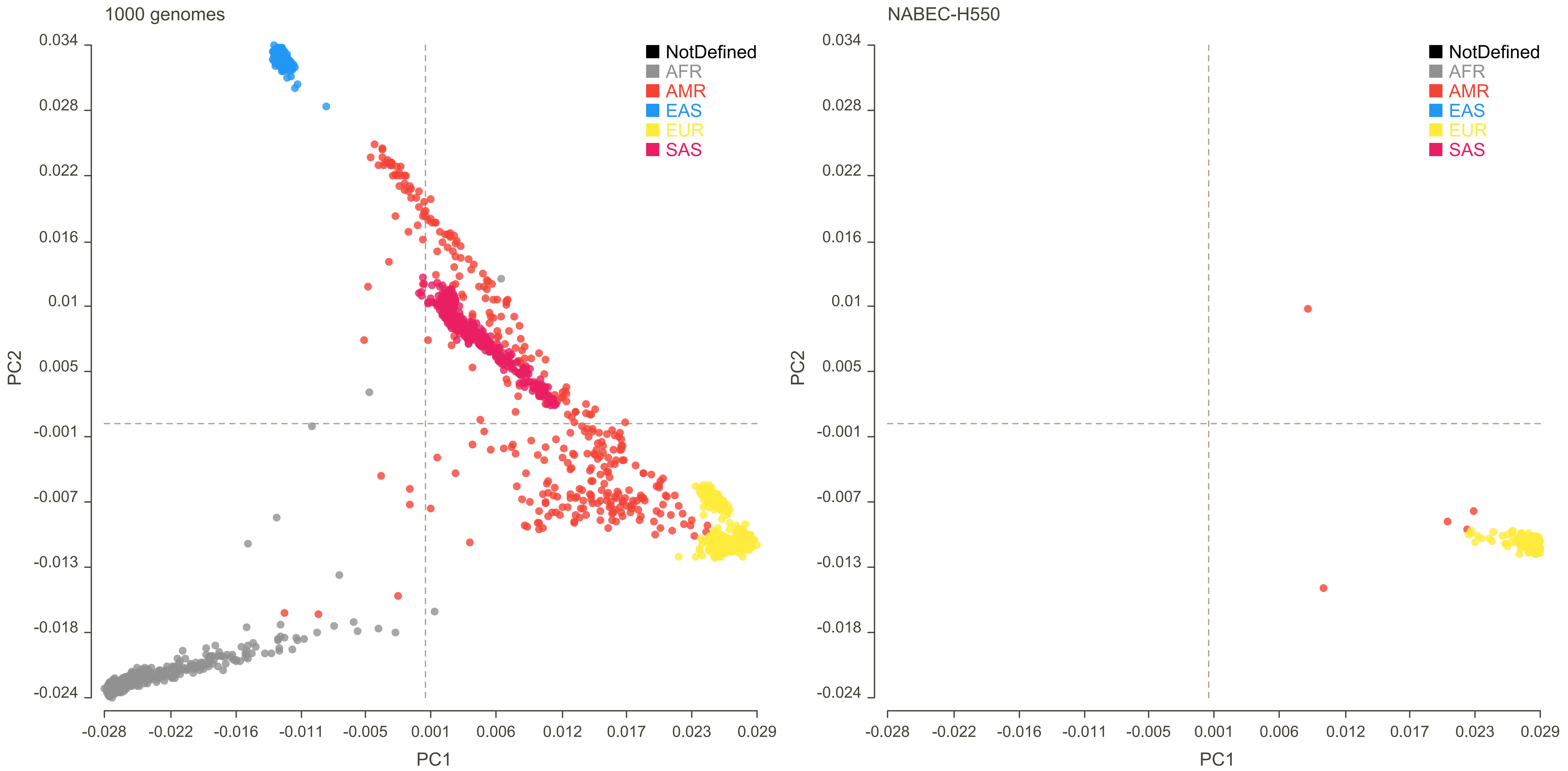
1000 genomes
NABEC-H550
NotDefined
AFR
AMR
EAS
EUR
SAS
PC1
PC2
0.034
0.028
0.022
0.016
0.01
0.005
-0.001
-0.007
-0.013
-0.018
-0.024
-0.028
-0.022
-0.016
-0.011
-0.005
0.001
0.006
0.012
0.017
0.023
0.029

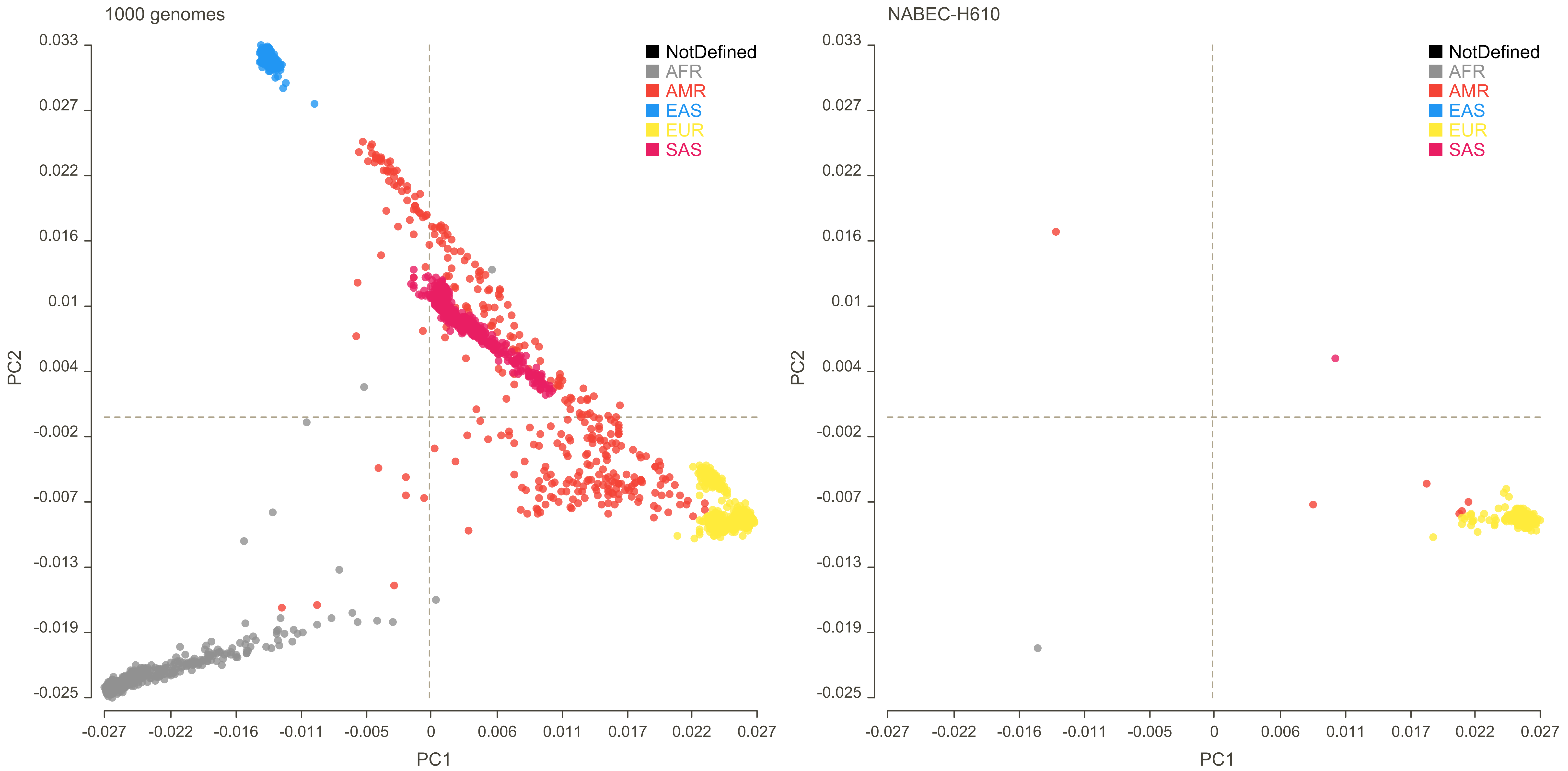
1000 genomes
NABEC-H610
NotDefined
AFR
AMR
EAS
EUR
SAS
PC1
PC2
0.033
0.027
0.022
0.016
0.01
0.004
-0.002
-0.007
-0.013
-0.019
-0.025
-0.027
-0.022
-0.016
-0.011
-0.005
0
0.006
0.011
0.017
0.022
0.027

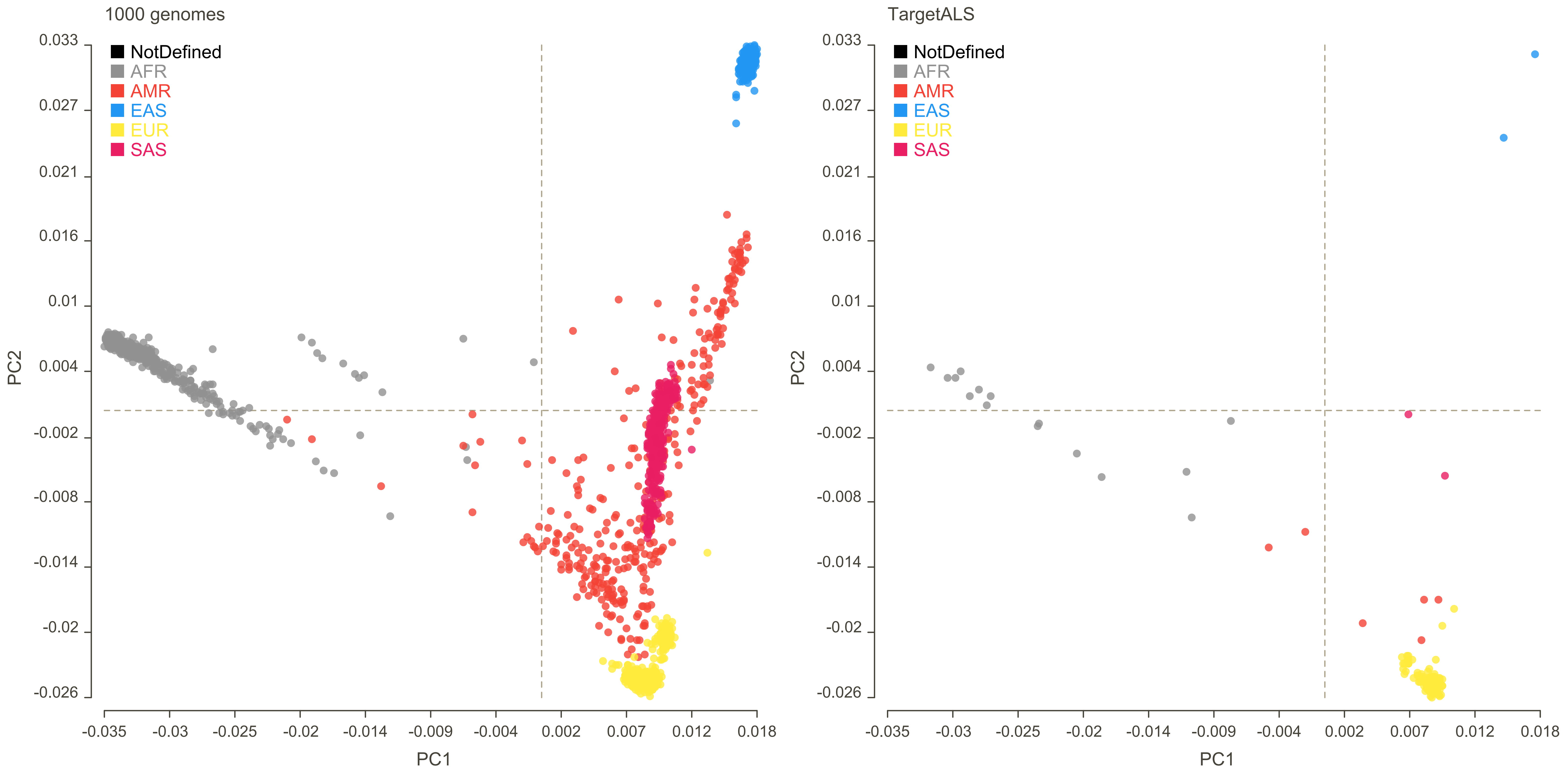
1000 genomes
TargetALS
NotDefined
AFR
AMR
EAS
EUR
SAS
PC1
PC2
0.033
0.027
0.021
0.016
0.01
0.004
-0.002
-0.008
-0.014
-0.02
-0.026
-0.035
-0.03
-0.025
-0.02
-0.014
-0.009
-0.004
0.002
0.007
0.012
0.018

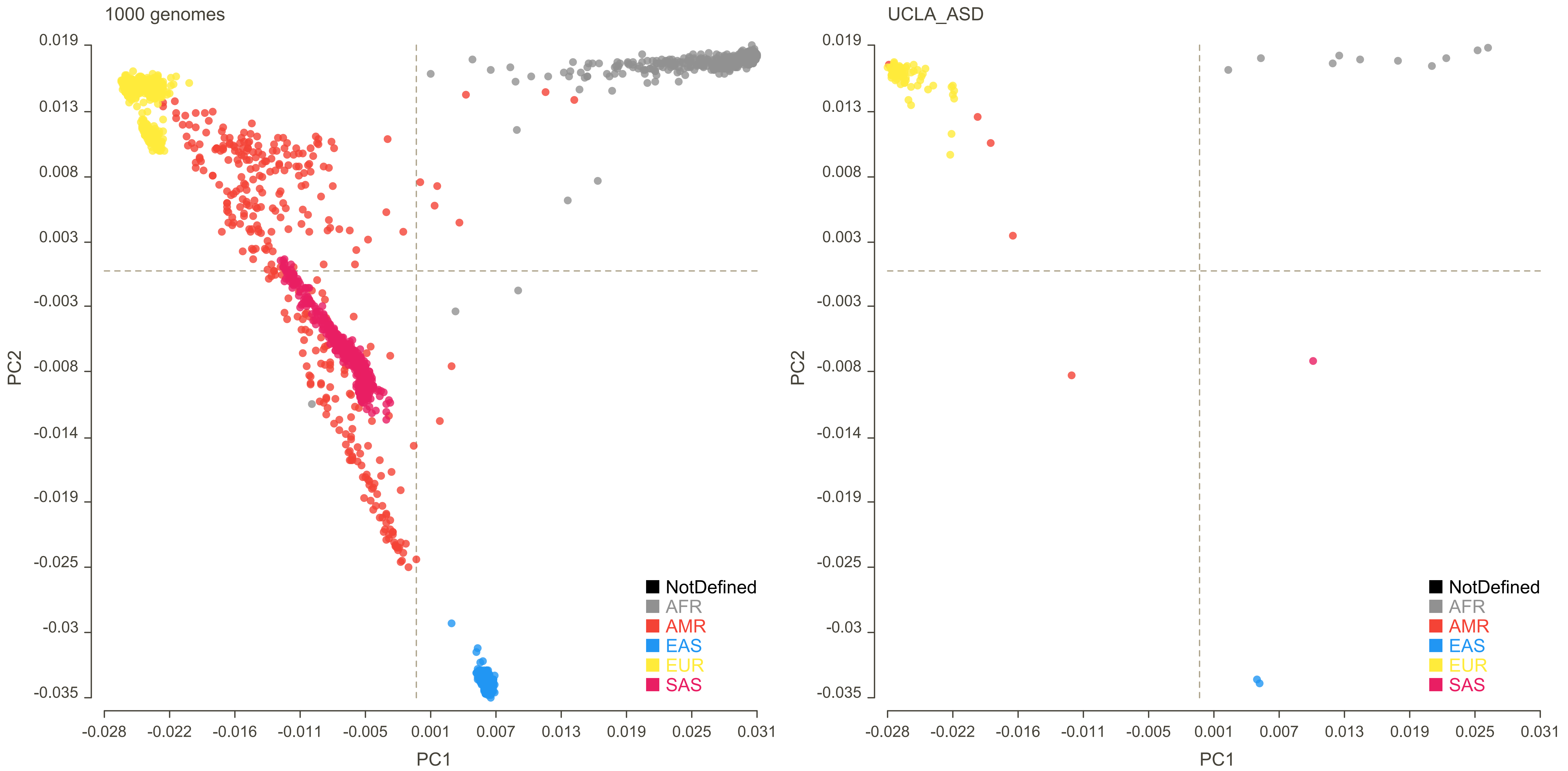
1000 genomes
UCLA_ASD
PC1
PC2
0.019
0.013
0.008
0.003
-0.003
-0.008
-0.014
-0.019
-0.025
-0.03
-0.035
-0.028
-0.022
-0.016
-0.011
-0.005
0.001
0.007
0.013
0.019
0.025
0.031
NotDefined
AFR
AMR
EAS
EUR
SAS
