## Supplementary Figure 21 for "Brain expression quantitative trait locus and network analysis reveals downstream effects and putative drivers for brain-related diseases"

A. CASS4 - Alzheimer's disease (AD)

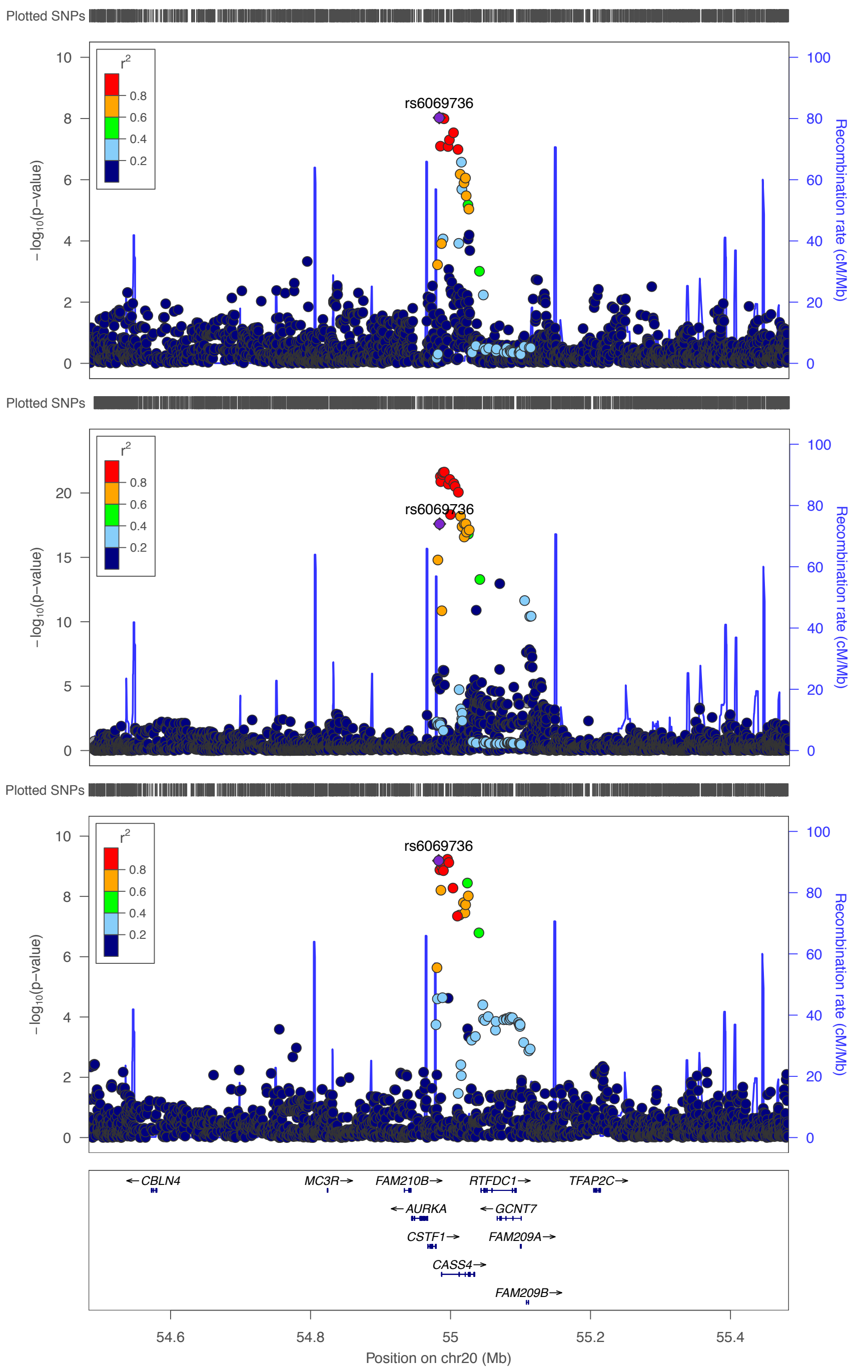


| Source | SNP | EA/NEA | Beta | P |
|---|---|---|---|---|
| metaBrain | rs6069736 | T/C | 0.252 | $9\times10^{-9}$ |
| eQTLGen | rs6069736 | T/C | -0.128 | $3\times10^{-18}$ |
| AD | rs6069736 | T/C | -0.105 | $7\times10^{-10}$ |

B. TMEM170B - Intelligence

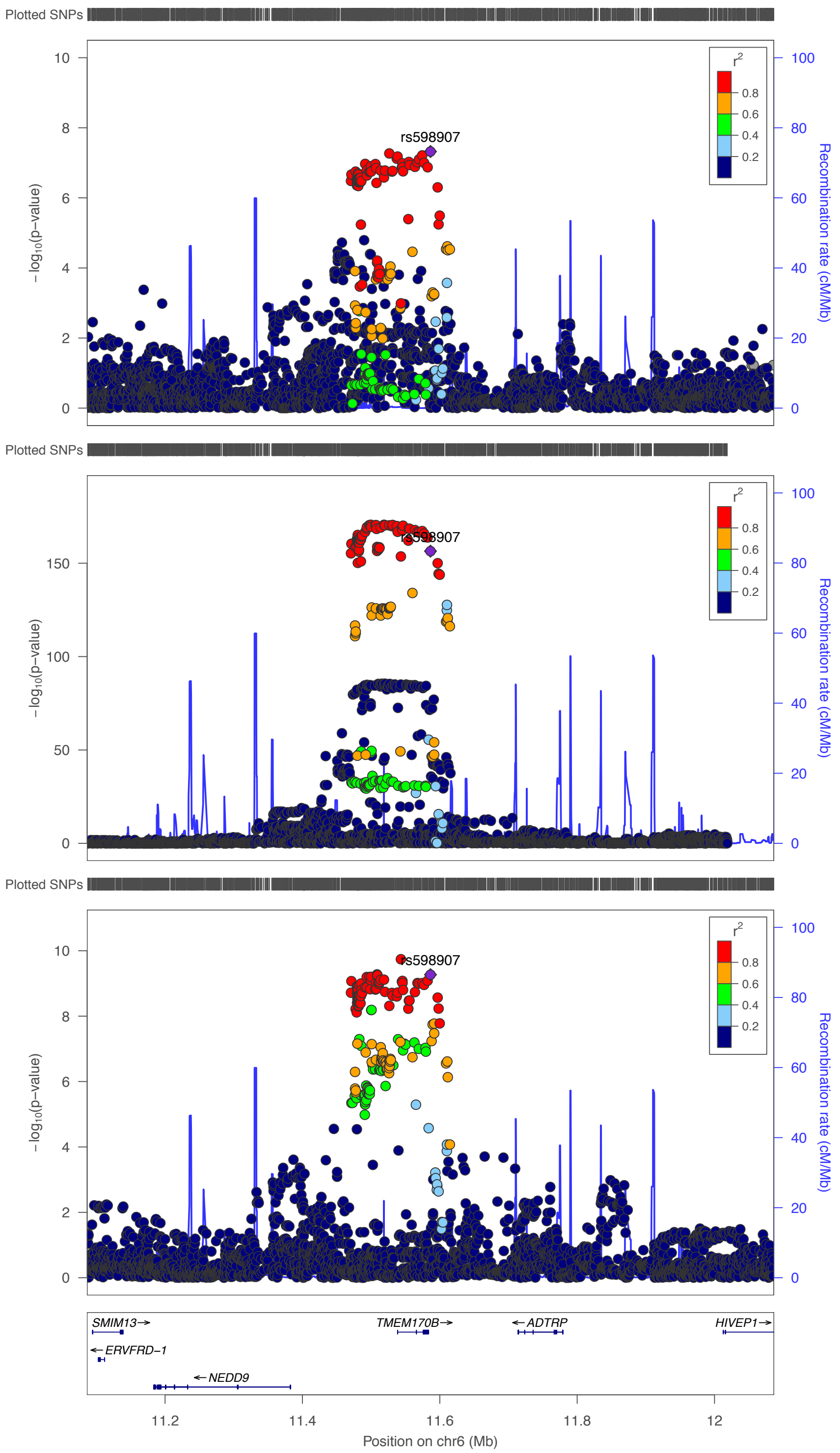


| Source | SNP | EA/NEA | Beta | P |
|---|---|---|---|---|
| metaBrain | rs598907 | C/T | 0.158 | $5\times10^{-8}$ |
| eQTLGen | rs598907 | C/T | -0.292 | $4\times10^{-157}$ |
| Intelligence | rs598907 | C/T | 0.019 | $5\times10^{-10}$ |

C. GATAD2A - Schizophrenia

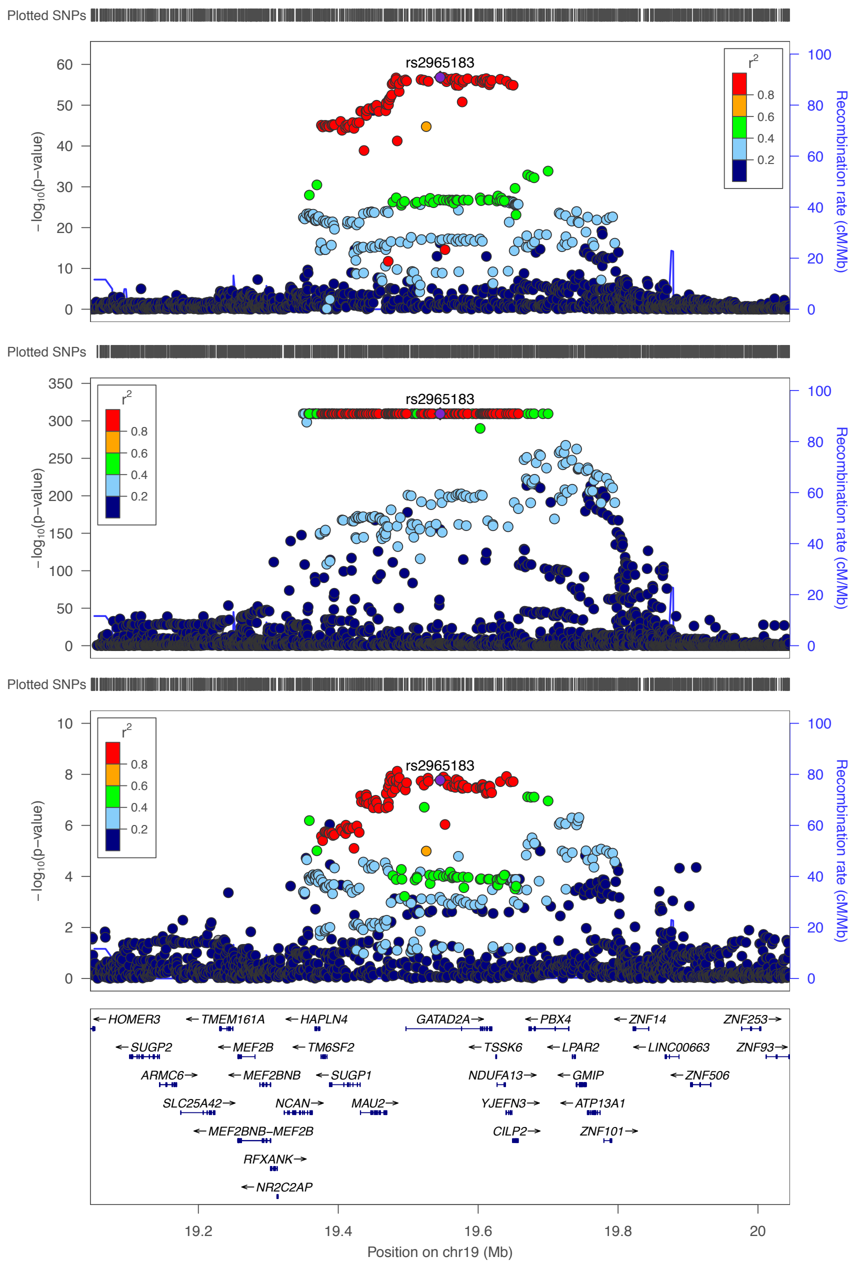


| Source | SNP | EA/NEA | Beta | P |
|---|---|---|---|---|
| metaBrain | rs2965183 | A/G | -0.416 | $1\times10^{-57}$ |
| eQTLGen | rs2965183 | A/G | 0.478 | $3\times10^{-310}$ |
| Schizophrenia | rs2965183 | A/G | 0.064 | $2\times10^{-8}$ |

D. GATAD2A - Years of schooling

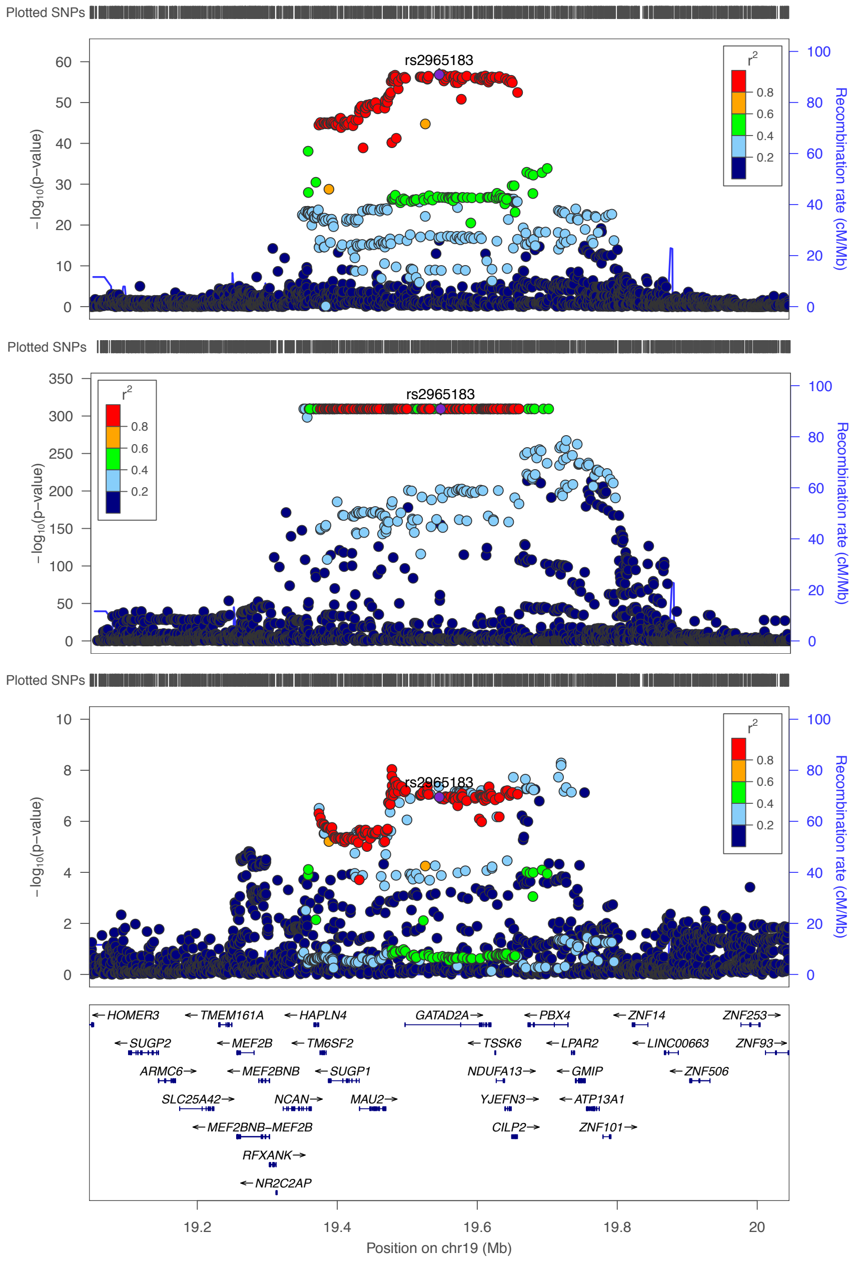


| Source | SNP | EA/NEA | Beta | P |
|---|---|---|---|---|
| metaBrain | rs2965183 | A/G | -0.416 | $1\times10^{-57}$ |
| eQTLGen | rs2965183 | A/G | 0.478 | $3\times10^{-310}$ |
| Years of schooling | rs2965183 | A/G | -0.009 | $1\times10^{-7}$ |

E. ZCWPW1 - Years of schooling

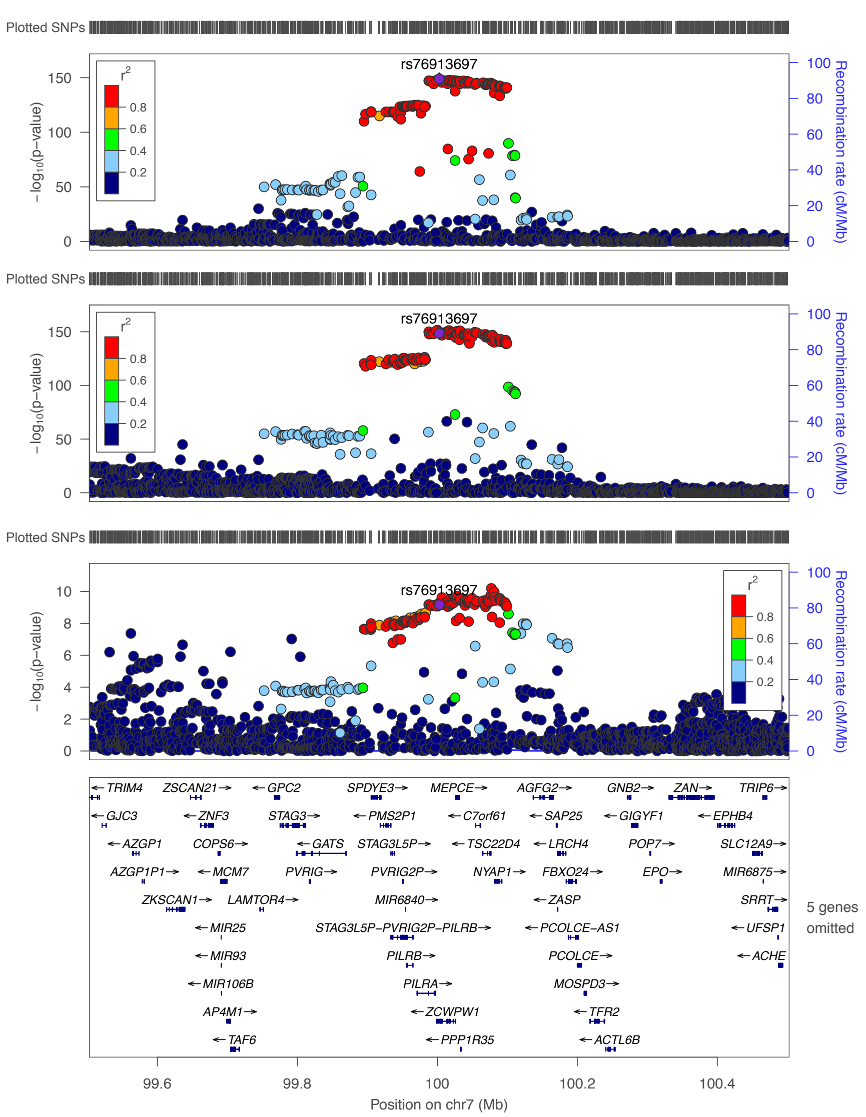


| Source | SNP | EA/NEA | Beta | P |
|---|---|---|---|---|
| metaBrain | rs76913697 | G/A | 0.772 | $1\times10^{-149}$ |
| eQTLGen | rs76913697 | G/A | -0.261 | $2\times10^{-149}$ |
| Years of schooling | rs76913697 | G/A | -0.013 | $7\times10^{-10}$ |
